# The dynamics of protein-RNA interfaces using all-atom molecular dynamics simulations

**DOI:** 10.1101/2023.11.07.565982

**Authors:** Afra Sabei, Cécilia Hognon, Juliette Martin, Elisa Frezza

**Affiliations:** Université Paris Cité, CiTCoM, CNRS, F-75006 Paris, France; Univ Lyon, Université Claude Bernard Lyon 1, CNRS, UMR 5086 MMSB, Lyon, France

## Abstract

Facing the current challenges raised by human health diseases requires the understanding of cell machinery at a molecular level. The interplay between proteins and RNA is key for any physiological phenomenon, as well protein-RNA interactions. To understand these interactions many experimental techniques have been developed, spanning a very wide range of spatial and temporal resolutions. In particular, the knowledge of tridimensional structures of protein-RNA complexes provides structural, mechanical and dynamical pieces of information essential to understand their functions. To get insights into the dynamics of protein-RNA complexes, we carried out all-atom molecular dynamics simulations in explicit solvent on nine different protein-RNA complexes with different functions and interface size by taking into account the bound and unbound forms. First, we characterized structural changes upon binding and for the RNA part the change in the puckering. Second, we extensively analyzed the interfaces, their dynamics and structural properties, and the structural waters involved in the binding, as well as the contacts mediated by them. Based on our analysis, the interfaces rearranged during the simulation time showing alternative and stable residue-residue contacts with respect to the experimental structure.

## Introduction

Protein-RNA interactions are crucial for any cellular function.^1^ RNA-binding proteins are key to RNA export, processing, splicing, localization, translation, degradation, and the regulation of all these steps.^2,3^ The dysfunction of such complexes is implicated in many humans and animal pathologies.^4,5^ For example, the formation of ribonucleoprotein particles (RNPs) including mRNAs is key for the post-transcriptional regulation of gene expression.^6,7^ As another example, many viruses manipulate the translation initiation complexes to ensure their replication, by recruiting the host ribosomes to translate their mRNA.^8^ Hence, understanding and predicting protein-RNA interactions is a crucial step to develop new therapeutic approaches and improve our knowledge of cell machineries.

Despite their significance, the specific mechanisms of protein-RNA interactions are not completely understood. Our ability to predict such mechanisms is hindered by several factors, among which the conformational changes undergone by the structures during the binding process and the lack of available tridimensional complex structures are to be mentioned.^9,10^ In fact, obtaining high-resolution tridimensional structures through high-resolution techniques is still a challenging task as shown by the relatively small number of structures deposited in the Nucleic Acid Data Bank (NDB).^11,12^ The limited number of known complex structures is nevertheless constrained by the inherent flexibility of proteins and/or RNA and the size of the complexes. The ability of binding proteins to change their conformation is often made possible by RNA flexibility, which is a crucial component in the binding process. Among other techniques, NMR spectroscopy is largely used and one of the most powerful methods for studying RNA’s and protein-RNA structure and conformational dynamics in solution. Despite that, the current application of NMR methodology imposes a constraint on the size of the molecules, typically limiting them to approximately 150 nucleotides (nt), achieved through the utilization of site-selective labeling strategies.^13–15^ In parallel, biochemical reactivity data on like SHAPE (selective 2’-hydroxyl acylation analyzed by primer extension) have become popular, but their application in the prediction of protein-RNA complexes is still limited since the change of reactivity upon binding depends on different factors like the change of flexibility of the ribose.^16–18^

Another key aspect is the proper characterization of the role of energetic, structural, and thermodynamic factors in the specificity and mechanism of complexation. Such factors are still beyond our understanding for a number of reasons. Thermodynamic data are very sparse and often obtained in different conditions, and the determination of the actual affinity strongly depends on the method used and on the strength of the interactions that are being measured.^19,20^ Therefore, most analyses have focused on structural explanations, and only a limited number of complexes have been analyzed in an attempt to characterize the common features of the interactions.^21,22^ In the dataset developed by Krüger and co-workers, 322 complexes are analyzed and their structures present a large diversity of interface size and properties.^23^ It has been shown that there are important differences in RNA-protein complexes as compared to DNA-protein complexes, most notably related to the nature of contacting surfaces of the two molecules.^21–23^ For example, for specific protein-RNA binding the number and types of intermolecular interactions and preferred amino acids have been observed.^24,25^ It is crucial to understand the specific characteristics of protein-RNA complexes with respect to protein-DNA interactions, because RNA and DNA can bind the same protein, but on orthogonal binding sites.^26^ DNA and RNA double helices also differ in conformation: the former assumes a B-form and the latter an A-form. On the RNA side, the phosphate contributes less, and the sugar more, to the interaction than in protein-DNA complexes.^21,22^ Regarding the amino acid propensity to be at the protein-RNA interface, it has been observed that it is close to the one obtained for protein-DNA interfaces that are enriched in positively charged and polar residues.^21,27^ A crucial player in protein-RNA recognition is the 2*^′^*-OH of the ribose.^22^

Although the information above is undoubtedly valuable to understand many aspects of protein-RNA interfaces, it gives a rather static view of such complexes and at the same time does not take into account the relationship between experimental structural features and energetic costs (binding affinity). In fact, the anatomic description of protein-RNA interfaces and their physico-chemical properties generally relies on a static point of view. However, the dynamics of interfaces plays a major role in the mechanisms of complexation (involving both specific and non-specific complexes) and it can influence the resulting binding strength. Moreover, to date, in the large datasets of protein-RNA complexes available^23,28^ most complexes do not dispose of both the structure of the cognate unbound RNA or proteins and the related binding affinity. This probably explains why so little is known on the role of dynamics of protein-RNA interfaces and on the energetic aspects of such interactions. Only few studies have been conducted on specific systems using all-atom Molecular Dynamics (MD) simulations,^29,30^ for example short single-stranded RNAs bound to RNA recognition motifs (RRM) proteins (HuR-RRM3 and SRSF1-RRM2),^31^ the adaptive binding of spliceosomal U1A-RNA binding to a hairpin RNA and an internal loop RNA,^32^ a GGUG motif bound to Fused in sarcoma RRM protein,^33^ reduced and oxidized TFIIIA Zinc Fingers interacting with 5S RNA.^34^ Shaw and co-workers performed microsecond MD simulations to analyze the SARS-CoV-2 helicase replication–transcription complex, in particular the non-specific helicase 13, to highlight their mechanism and to characterize the interfaces.^35,36^ However, no systematic study has been conducted to assess if protein-RNA interfaces can undergo significant changes in simulations and if changes in the interfaces of native complexes can be observed as pointed out for protein-protein^37,38^ and protein-DNA interfaces also from some of us.^39–42^

In this context, water molecules also play a crucial role in the process of binding, in specific recognition, in interactions between protein-RNA as well as influence the structure and function of isolated biomolecules.^43^ In fact, they seem to be involved in mediating interactions,^44^ as well in the formation of H-bonding network to stabilize the interactions.^45^ At the interface, water molecules play a crucial role in determining the specificity and stability of the complex. However, with respect to protein(s) and DNA-protein interfaces,^46^ the water dynamics of RNAs and for protein-RNA complexation has been poorly investigated. Although several PDB structures have been analyzed, a static view has often been presented^44,47,48^ and only few studies have been conducted that take into account the interplay between water and protein/RNA dynamics and their interfaces.^49^

In this study, we aim to characterize the dynamics of protein-RNA complexes and their interfaces at molecular level by performing a more systematic and extensive analysis. To get insights on the dynamics of protein-RNA complexes and in the possibility of alternative interfaces, we investigated nine protein-RNA complexes for which at least the 3D structures of the complex and the unbound RNA molecule are available. We carried out standard all-atom MD simulations of complexes and unbound proteins/RNAs in explicit water with K^+^Cl*^−^* ions at physiological conditions. MD simulations were extensively characterized and several analyses were conducted allowing to propose a computational strategy that combines different tools. First, we studied how the binding could impact the stability, the conformation and the flexibility of each partner. A detailed analysis were conducted on the RNA structure, including the change of ribose puckering upon binding for its potential interest to better understand biochemical reactivity data as SHAPE. Then, we focused on the characterization of the stability of their interfaces and interfacial water molecules. To determine the presence or the absence of interface substates, in each system, we analyzed in detail the time evolution of the contacts between interface residues and we performed a hierarchical clustering on our trajectories based on the interface residue-residue contacts. Moreover, we characterized the interactions at the interface and the interface geometries, as well as the evolution of the contacts between interface residues and water molecules and the pairs of residues involved in mediated interactions via water molecules. Finally, in the case of alternative interfaces obtained with our approach, we performed a dynamic network analysis to assess how the change of contacts at the interface could impact the communication in the complex and to gather novel biological insights.

## Materials and Methods

### Data set

In order to analyze the protein-RNA interfaces, we took into account the datasets available in the literature for protein-RNA complexes.^19,50–53^ We performed a structural characterization of each candidate complex in order to avoid redundant structures and with the aim to assure a diversity in the size and type of the interfaces. To set-up the data set, we also took into account the variability in the RNA size and in the buried accessible surface area (ΔASA), the presence of different RNA motifs and different biological function. Another criterion was to choose RNA molecules whose size is compatible with SHAPE experiments. Based on that, we selected 9 complexes as reported in Table 1. Among them, the complex with PDB ID 3IEV was chosen for its RNA motif and small ΔASA, despite the potential difficulties to obtain reliable SHAPE experiments on it. The nine cases span various interface sizes (1341 - 3937 Å^2^) and RNA motifs, single strand (ss) or double strand (ds), at interface. The 3D structures of the 9 complexes under study are shown in Figure 1.

**Figure 1:**
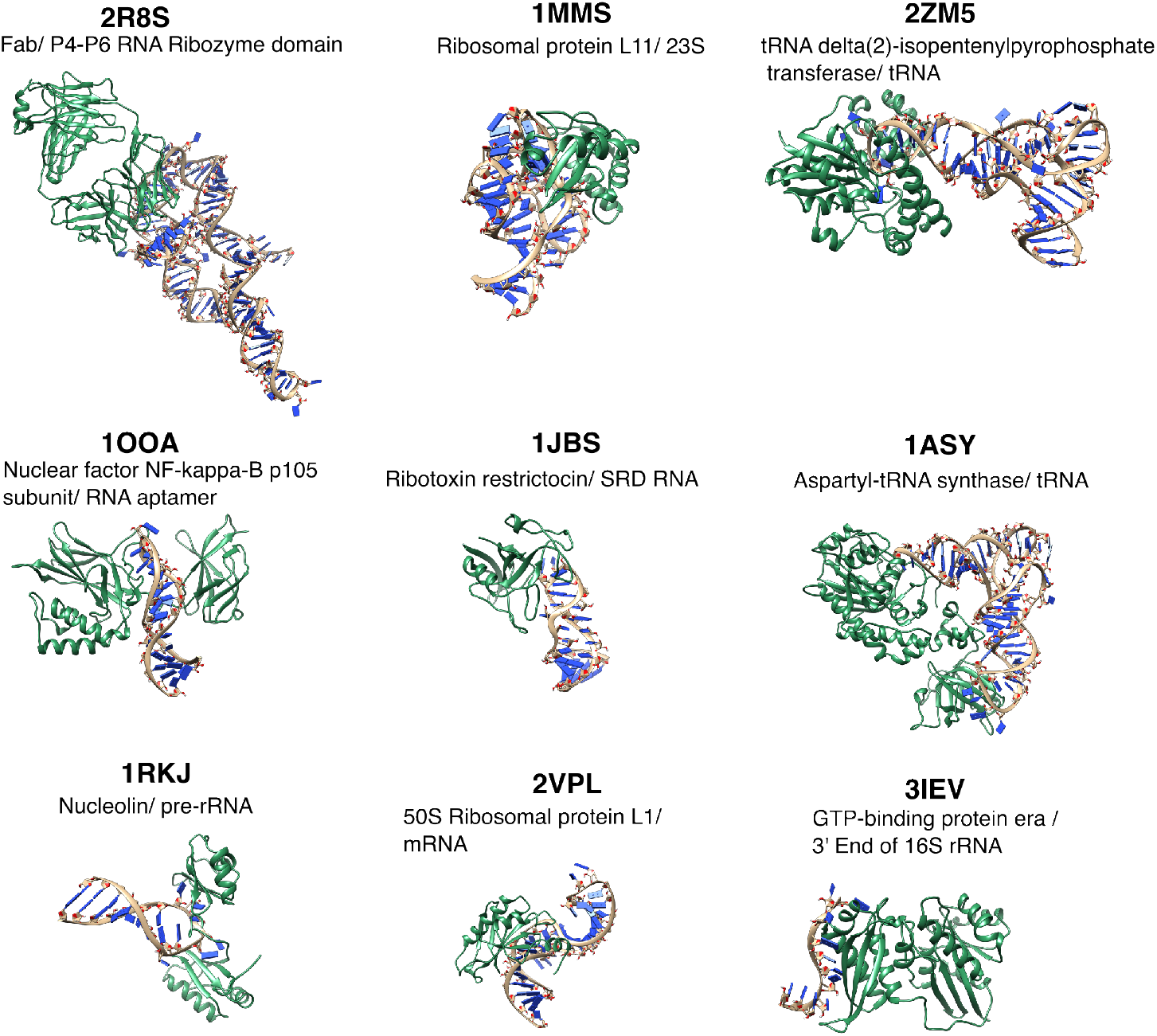
3D structures of the nine protein-RNA complexes studied in this study, with one partner colored in green (protein) and the other one in beige (RNA).

**Table 1:** Dataset of unbound-to-bound transition. b: bound; u: unbound; r: RNA; p: protein; RMSD*_exp_*: RMSD between the unbound and bound RNA structure; ss: single strand; ds: double strand. *: different NMR structures. *^w^*: crystallographic waters in the starting structure.

| PDB ID<sub>b</sub> | PDB ID<sub>u,r</sub> | PDB ID<sub>u,p</sub> | $RMSD_{exp}$ (Å) | Interface size (Å<sup>2</sup>) | RNA motifs |
| --- | --- | --- | --- | --- | --- |
| 1OOA(A:C) | 2JWV* | 3DO7 | 5.4 | 1910 | ss/ds |
| 1ASY(A:R) | 2TRA | 1EOV | 4.9 | 3585 | ss/ds |
| 2ZM5(A:C)<sup>w</sup> | 3L0U(A) |  | 4.2 | 3937 | ss/ds |
| 1MMS<sup>w</sup> | 3I9C(A) | 2K3F* | 1.6 | 2456 | ss/ds |
| 3IEV(A:D) <sup>w</sup> | 1SDR(B) |  | 7.6 | 2311 | ss |
| 2R8S(HL:R A)<sup>w</sup> | 1HR2 | 3IVK(AB) | 4.4 | 2416 | ss/ds |
| 2VPL(A:B) | 1U63 | 2OV7(A) | 3.7 | 2388 | ds |
| 1JBS(A:C) | 1AQZ | 1AQZ | 3.4 | 1341 | ss/ds |
| 1RKJ(A:B)* | 1QWA |  | 4.9 | 2218 | ss/ds |

To model our starting structures, several steps were necessary with the aim to use the same number of residues for the complex and the free protein and RNA molecule. Terminal residues present in the free protein/RNA but absent from the complex were chopped, and residues present in the complex but absent in the free protein/RNA were modeled based on the complex. In the absence of the PDB structure for the unbound protein, the bound form was taken into account. When one or more residues were absent, homology modeling was used to add the missing residue using either Modeller 10.2^54^ or ModeRNA 1.7.0^55^ for the protein or the RNA molecule, respectively. An exception was made for the unbound form of the 1RKJ complex protein, due to the absence of a correct homology model. For this reason, we used the structure extracted from the PDB of the complex. For the PDB structures obtained via NMR, we first carefully analyzed the overall quality of the produced models and then we used the software DSSR (Dissecting the Spatial Structure of RNA)^56^ to examine and annotate the RNA tertiary structures. As a result, we were able to determine both canonical and non-canonical base pairs and the number of hydrogen bonds in each structure. Then, in order to select the most representative model, we clustered the available models and constructed arc representations of the structures. To prepare the structures, we also used the crystallographic water molecules for the structures with divalent ions obtained by X-ray crystallography, the monovalent ions and Mg^2+^ ions. Finally, for the protein, we computed a pKa calculation on the complex structure using the software PDB2PQR^57^ and the algorithm PROPKA^58^ to define the protonation state of each residue. To deal with the same chemical species, we applied the same protonation state of the protein in the complex on the relative free protein(s).

### All-atom Molecular dynamics simulation

All-atom MD simulations were performed with the GROMACS 2018 package^59–62^ using the Amber ff14SB force field for proteins^63^ and the the Amber ff99bsc0 force field with OL3 modification (ff99bsc0*χ*_OL3_) for RNA molecules.^64^ Each RNA molecule was placed in a cubic box and solvated with TIP4P-EW to a depth of at least 14 Å.^65^ Each system was neutralized by adding potassium cations and then K^+^Cl*^−^* ion pairs to reach a physiological salt concentration of 0.15 M and by adding MgCl_2_ to reach a concentration of 0.02 M.^66^ Long-range electrostatic interactions were treated using the particle mesh Ewald method ^67,68^ with a real-space cutoff of 10 Å. The hydrogen bond lengths were restrained using P-LINCS,^69^ allowing a time step of 2 fs. Translational movement of the solute was removed every 1000 steps to avoid any kinetic energy build-up.^70^

Before the MD production, we carried out the energy minimization and equilibration. During equilibration (at least 10 ns) a Berendsen thermostat (*τ_T_* = 0.1 ps) and Berendsen pressure coupling (*τ_P_*= 0.5 ps)^71^ were used. The production part was carried out in an NTP ensemble at a temperature held at 300 K and a pressure held constant at 1 bar using the Bussi velocity-rescaling thermostat (*τ_T_*= 0.1 ps)^72^ and the Parrinello-Rahman barostat (*τ_P_* = 0.5 ps).^73^ During minimization and heating, RNA heavy atoms and protein backbone remained fixed by using positional restraints. During the equilibration, the restraints were gradually relaxed. Bond lengths were restrained using P-LINCS, allowing a time step of 2 fs. Initially, each simulation was 500 ns long. Due to the stability obtained during the simulations, most of them were extended, to a maximum of 1500 ns. All the simulation times are available in Table S1. A total simulation time of 21.75 *µ*s was performed.

### RMSD/RMSF analysis of complexes and single protein/RNA

We calculated RMSD time series on the unbound RNA, bound RNA, and complex using their starting structures as reference in order to determine system stability and binding effect. We computed RMSF for all atoms, averaged fluctuations for each residue, and superimposed the trajectory on the initial structure for unbound RNA and separated chains for complexes. Using this method, we were able to examine how each chain’s flexibility changed as it was bound. Finally, we performed a hierarchical cluster analysis using the software TTClust^74^ to determine the clusters for the unbound and bound systems. We chose this software since it performs a hierarchical analysis such as the algorithm implemented to clusterize the interfaces based on the contacts (see below) and it is capable of capturing global conformational changes that we are interested in studying rather than local ones.

### Determination of Free Energy Landscape (FEL)

To obtain a free energy landscape (FEL), first we carried out a principal component analysis (PCA) on the all-atom molecular dynamics simulations. To do, we considered only the heavy atoms and computed a least squares fit on the molecular system and the covariance matrix, and determined the eigenvalues and the eigenvectors as implemented in GROMACS 2018.^75,76^ Then, we computed the free energies using the command gmx sham. We performed the FEL on the unbound states, on the full complex, on the bound RNA and on the bound protein(s).

### RNA analysis

#### Puckering analysis

To take the ribose flexibility and its conformation, we computed the sugar pucker. It is important to highlight that the sugar ring is well described using pseudorotation parameters. Although there are four possible pseudorotation parameters for a five-membered ring (Frezza et al. ^77^), two in particular are useful to characterize the sugar conformation: the so-called phase (Pha) and amplitude (Amp). While the amplitude describes the degree of ring puckering, the phase describes which atoms are most out of the mean ring plane. We calculated these parameters using the expressions given below:

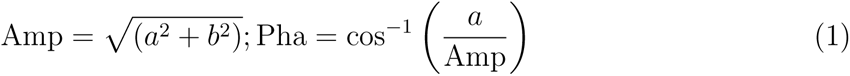

where 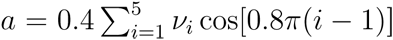 and 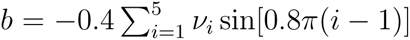 with *ν_i_* the ring dihedral angle *i*. This approach has the advantage of processing the ring dihedrals *ν*_1_ (C1’ -C2’ -C3’ -C4’) to *ν*_5_ (O4’ -C1’ -C2’ -C3’) in an equivalent manner. Conventionally, sugar ring puckers are divided into 10 families described by the atom which is most displaced from the mean ring plane (C1’, C2’, C3’, C4’ or O4’) and the direction of such displacement (*endo* for displacements on the side of the C5 atom and *exo* for displacements on the other side). In different sugar conformations, the distance between neighboring C2 atoms in the nucleobases and the orientation of P relative to the sugar/bases are different from each other. Based on previous works, these quantities are supposed to be related to the chemical reactivity of RNA with SHAPE reagents. In order to understand the interplay between the sugar conformation and protein-RNA binding, we grouped the sugar puckers into two large families. The sugar puckers C1’-*exo*, C2’-*endo*, C3’ -*exo*, C4’-*endo* belong to the B-like family, while C1’-*endo*,C2’-*exo*, C3’-*endo*, C4’-*exo* belong to the A-like family. For the single angular quantities, because of their periodicity, standard statistics cannot be used to compute the average and the fluctuations of angular values. For example, the arithmetic mean of 0° and 360° is 180°, which is misleading because for most purposes 360° is the same thing as 0°. Hence, we used the directional statistics instead, as explained in.^77^

#### Secondary Structure Analysis

In order to analyze the stability of unbound and bound RNA molecules and better characterize the secondary structure, we computed the hydrogen bonds and the stacking for each frame using the software MC-Annotate.^78–80^ The hydrogen bonds were classified using the Leontis/Westhof classification^81,82^ and the definition used in.^79^ In particular, we consider if the atoms involved in the HBs are the atom O2’, C8 or one oxygen of the phosphate group (OP) or belong to the Watson–Crick side (W), Hoogsten side (H), the sugar side (S), the B side (the position is between the W and H side or between the W and S side). For each type of HBs and stacking, we computed its average and the standard deviation (*sd*).

### Interface Analysis

The interface analysis is conducted on MD snapshots taken every 40 ps (i.e. 12500 to 25000 snapshots depending on the system). Part of this analysis was computed with in-house python scripts that are available at https://github.com/juliettemartin/DYN_INTERFACES.

#### Interface definition

Interface contacts are defined at the residue level using a 5 Å distance cutoff between heavy atoms. The solvent accessible surface area (ASA) was computed with the NACCESS software using the default radius of 1.4 Å to take into account the water molecule.^83^

Following the criteria introduced by Levi for the protein-protein interfaces, distinct regions were defined at the interface.^84^ Hence, residues with a change of ASA between the isolated chain and the complex are classified as: core if relative ASA in the isolated chain is greater than 25% and lower than 25% in the complex, rim if relative ASA in the complex is greater than 25%, support if the relative ASA in the isolated chain extracted from the complex is lower than 25%.

#### Interface properties

The accessible surface area buried by the interface is defined by:

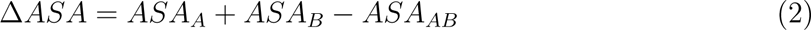

where *ASA*_A_ and *ASA*_B_ are the accessible surface areas of separate chains A and B respectively, and *ASA_AB_*represent the accessible surface area of the complex. The gap index evaluates the complementarity of the interacting surfaces in a complex and is defined as

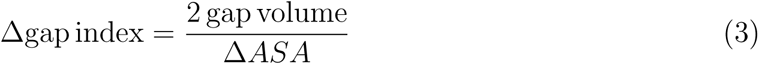

where the gap volume estimates the volume enclosed by any two molecules and is computed using the procedure developed by Laskowski and implemented in the SURFNET software.^85^

#### Inter hydrogen bonds and stacking

To characterize the hydrogen bonds between the protein and the RNA, at each frame of the trajectory, first we identified the HB based on cutoffs for the *Donor-H… Acceptor* distance and angle according to the Wernet-Nilsson criterion outlined in^86^ as implemented in mdtraj^87^ which considers a hydrogen bond formed if the distance between donor and acceptor heavy atoms is below a given distance cutoff dependent on the angle made by the hydrogen atom, donor, and acceptor atoms (“cone” criterion). Second, we computed the average and the standard deviation for the total number of HBs and for the HBs which involve the hydroxyl of the ribose (HB_2_*′_−OH_*).

To investigate the stacking interactions between the protein and the RNA at the interface, we considered the rings of PHE, TYR, TRP and HIS for the protein and the nucleobases for the RNA. First, we computed the center of mass for each sub-residue (**r***_COM,pr_* if it belongs to the protein and **r***_COM,r_* if it belongs to the RNA) and their distance (*d_stack_* = **r***_COM,pr_* − **r***_COM,r_*). Second, we computed the angle between the normal vectors to the sub-residues (*α_stack_*) defined as follows:

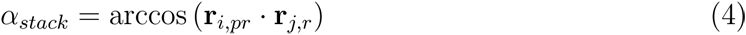

where **r***_i,pr_* = (**r**_1*,pr*_ − **r***_COM,pr_*) × (**r**_2*,pr*_ − **r***_COM,pr_*) are **r***_i,pr_* = (**r**_1*,r*_ − **r***_COM,r_*) × (**r**_2*,r*_ − **r***_COM,r_*) are the normal vectors to a ring of the protein and the nucleobase, respectively. The vector **r**_1_*_,pr_* represents the coordinates (*x, y, z*) of the atom CG and **r**_2_*_,pr_* represents the coordinates (*x, y, z*) of the atom CD1 for PHE and TRP, the atom ND1 for TYR and the atom NE1 for HIS. The vector **r**_1_*_,r_* represents the coordinates (*x, y, z*) of the atom N9 for G and A and the atom N1 for U and C and **r**_2_*_,pr_* represents the coordinates (*x, y, z*) of the atom C2 for all nucleobases. We considered that a stacking interaction is present if the distance *d_stack_* is below 5 Å and the absolute value of *α_stack_* is lower than 45*^◦^*. Finally, for each pair of residues of the protein and RNA involved in the stacking interactions, we computed their frequency along the full MD trajectory and for each interface cluster if present.

#### Interface clustering

To conduct a comparison between interfaces at various time points, we analyze the interface contacts of the respective snapshots, S1 and S2. The interface similarity is then measured by the Jaccard index *J* defined as:

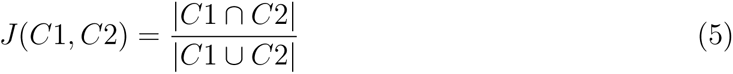

where C1 and C2 represents the two sets of contacts. The Jaccard index is equal to 1 if S1 and S2 have identical contacts. On the contrary it is equal to 0 if S1 and S2 have no common contacts.

First, the Jaccard indexes are converted into dissimilarity matrices by taking 1-*J*, followed by the computation of hierarchical clustering to identify clusters at the interfaces. To do so, the Ward.D2 method implemented in R was used.^88^ No generic method can be applied to choose the optimal number of clusters since the choice depends on the application. In this study, we aimed to have clusters that are different enough from each other, of reasonable size, and relatively stable in time. The choice of the optimal number of clusters was thus guided by the topology of the clustering dendrograms and the assessment of cluster size and stability in time when varying the number of clusters. We thus computed the size of the largest cluster, the size of the smallest cluster (number of snapshots in this cluster), and the number of cluster changes during the simulation time.

#### Choice of representative structures in each cluster

In each cluster, we defined the centroid as the snapshot with the highest average Jaccard similarity with respect to the other snapshots of the same cluster. To visualize the location of these centroids in each cluster, we applied principal component analysis on the Jaccard dissimilarity matrices and plotted the first two dimensions. In the SI, projections are presented.

#### Statistics of interface contacts in the different clusters

We computed the relative frequency of contacts in each cluster, defined by the proportion of snapshots exhibiting a contact in the cluster. Each contact is thus associated with *FN_c_*values of relative frequencies with *N_c_* the number of clusters. For each contact, we calculated the variance of these values among clusters. This quantity points out which contacts have similar frequencies (low variance) and which have variable frequencies across clusters (high variance). Variance values refer to contacts between residue pairs. To map these variance values at the level of residues, for each residue we selected the maximum variance, var(*X*), observed among contacts:

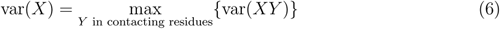

where var(*XY*) is the variance of the relative frequency of the *XY* contact among clusters. For example, a residue *X* seen in contact with residue *Y* with a variance of 0.2 and in with residue *Z* with a variance of 0.4 will receive a value of 0.4.

### Dynamical Network Analysis

To evaluate the impact of alternative interfaces on the allosteric communication in the complex, we performed a dynamical network analysis using the package dynetan.^89^ Each nucleotide was defined via two nodes. The first node is centered on either the phosphate or the O5’ for the nucleotide in 5’ and the second one on the atom N1 of the nucleobase. The protein is defined by only one node centered on C*α*. We computed the dynamical network analysis on each interface cluster by considering only one window, we compared the obtained communities between the clusters and for each complex we defined a specific path based on available pieces of information available in the literature. To visualize the different communities, optimal and suboptimal paths within the complex, we used VMD^90^ and Network View 2 GUI.^91^

### Water analysis: interfacial water molecules

To analyze the water at the interface, we used the same approach proposed by some of us in.^42^ Hence, first, we determined the water molecules whose distance, *d_w_*, from both protein and RNA is below 4 Å. To compute this distance, only the heavy atoms of the biomolecules and the oxygen of the water molecules were considered for determining the number of water molecules at the interface, also called interfacial water molecules. For the latter, to investigate the contacts mediated by water i.e., triplets constituted by a residue from one chain in contact with a water molecule which is in contact with a residue from the other chain, we also analyzed the contacts formed for each interfacial water molecule with the protein and RNA partners using the same distance cutoff *d_w_* as well their conservation along each MD simulations. Finally, we computed the variance of the contacts mediated by water when at least two interface clusters were determined. The data obtained from the MD simulations were analyzed using Python scripts, the packages mdtraj,^87^ NumPy^92^ and SciPy^93^ libraries.

### Statistical test

The correlation between variables was assessed using the Pearson correlation coefficient using the function coorcoeff in Matlab. The difference between two correlation coefficients was assessed using the Dunn and Clark test^94^ implemented in the R package cocor,^95^ that takes into account the intercorrelation between variables in the case of dependent groups.

## Results and Discussions

To study the dynamics of protein-RNA interfaces, first we investigated the impact of the formation of the complex at the level of the flexibility and conformational changes of RNA and proteins by computing RMSD, RMSF, clustering analysis and free-energy landscapes using PCA analysis. To evaluate the stability of RNA molecules, the time evolution of the RNA hydrogen bonds has also been taken into account. Second, we computed some geometric interface parameters (accessible surface area, gap volume and gap index) along each MD trajectory to better characterize the interface geometry for each complex and their stability, as well the hydrogen bonds and the stacking between the protein and the RNA molecules. Third, we clustered each trajectory based on the contacts at the interface in order to identify alternative interfaces. Each cluster was characterized based on several parameters including hydrogen bonds, stacking interactions, geometrical parameters. The clusters were also compared based on the number and type of contacts, by computing the variance. Moreover, based on dynamical network analysis we were able to access the impact of the different interfaces on the communication inside the complex. Since the interaction with a protein can impact the RNA flexibility and its reactivity, we also computed the puckering for each nucleotide and compared the results obtained for the free and bound RNA for each complex and within the interface clusters. To better characterize the interfaces, we also took into account the water molecules at the interfaces and we characterized the interactions mediated by them.

### Investigation of conformational changes and flexibility along molecular dynamics simulations and upon binding

As presented above in the Section Materials and Methods, to investigate the conformational changes along the MD simulations, we first performed a hierarchical clustering analysis using TTClust^74^ based on RMSD on the complexes and the single protein and RNA and we computed the RMSD time series. The RMSD time series, based on starting and clustering structures results, are summarized in Figures S1-S9; Table S2 reports the mean value. Second, we also computed the Free Energy Landscapes (FEL) for all the molecular species (unbound states, complex and bound RNA and protein) and compared them with the RMSD times series and the structural clusters (see Figures S13-21). Different landscapes are observed: with a single minimum, with two close minima, with broad minimum/minima, or rugged landscapes with multiple local minima and barriers. Based on these distinctions, the simulations conducted on the unbound proteins highlight varying degrees of conformational changes and number of conformational states: low (as observed in 3IEV and 2R8S where a single minimum is present), moderate, and high (e.g., 1RKJ where several distant minima are obtained). These changes involve conformational modifications confined to tails, specific loops, and relative movements between protein domains (e.g., in the case of 1ASY, where two minima are obtained and the movements involve the loop between the domains).

When unbound RNA is simulated, again low (e.g., 1RKJ where a single minimum is present and the structural clusters differ mostly only at the level of the loop), moderate and high (e.g. 2VPL where two distant minima are present) conformational changes are observed. These conformational changes include for example the tails (e.g., 2ZM5 where also three local minima are observed in the FEL), specific loops in the hairpin (e.g., 1JBS where the FEL also shows two distant minima) or backbone conformational changes which impact the whole structure and arise to relative movements between different parts of the RNA structure (e.g., 3IEV). It is interesting to point out that the presence of several structural clusters does not mean that several conformational states are obtained, like in the case of 2R8S where the FEL shows only a minimum and the structural clusters highlight the presence of backbone conformational changes which impact the whole structure. Despite these conformational changes, the RNA secondary structure for all the systems is stable as shown in Table S3-S5. In fact, the stacking interactions are stable (mostly upward stacking), as well as the canonical (cWW) and non-canonical HBs.

Regarding the simulations carried out on the complexes, we again obtained different scenarios. For example, for 2ZM5 the RMSD time series is stable and a single conformational state is present in the FEL (see Figure S20). Moreover, to understand the origin of the conformational changes and the presence of several states, we compared the FELs obtained for the complex and the separate partners. Based on this comparison, three main situations are observed: i) different local minima for one partner with no impact on the conformation of the complex (e.g. 2ZM5); ii) one conformational state for the partners and at least two conformational states for the complex suggesting a rigid-body movement between the protein and the RNA molecule (e.g. 1JBS); iii) several conformational states for the partners and the complex suggesting the presence of both local and rigid body movements (e. g. 1RKJ). The case ii) and iii) are shown in Figure 2. Based on structural clustering analysis, for the complexes 1ASY, 1OOA, and 1RKJ we can also observe that large pivotal movements of one biomolecule with respect to the other are obtained which are responsible for a high global change in the RMSD. Finally, by comparing the FELs obtained for the bound and unbound proteins and RNA molecules, we can assess if the complex allows the single partner to investigate more states or select a specific one. For example, for 2R8S the protein can investigate different conformational states in the complex and only one state in the unbound form. On the contrary, for 1JBS the protein investigates more states in the unbound form. Similar results are obtained for the RNA molecules. As for the unbound state, also for the bound RNA molecules under investigation in this work, the secondary structure for all the systems is stable and no dramatical changes are observed.

**Figure 2:**
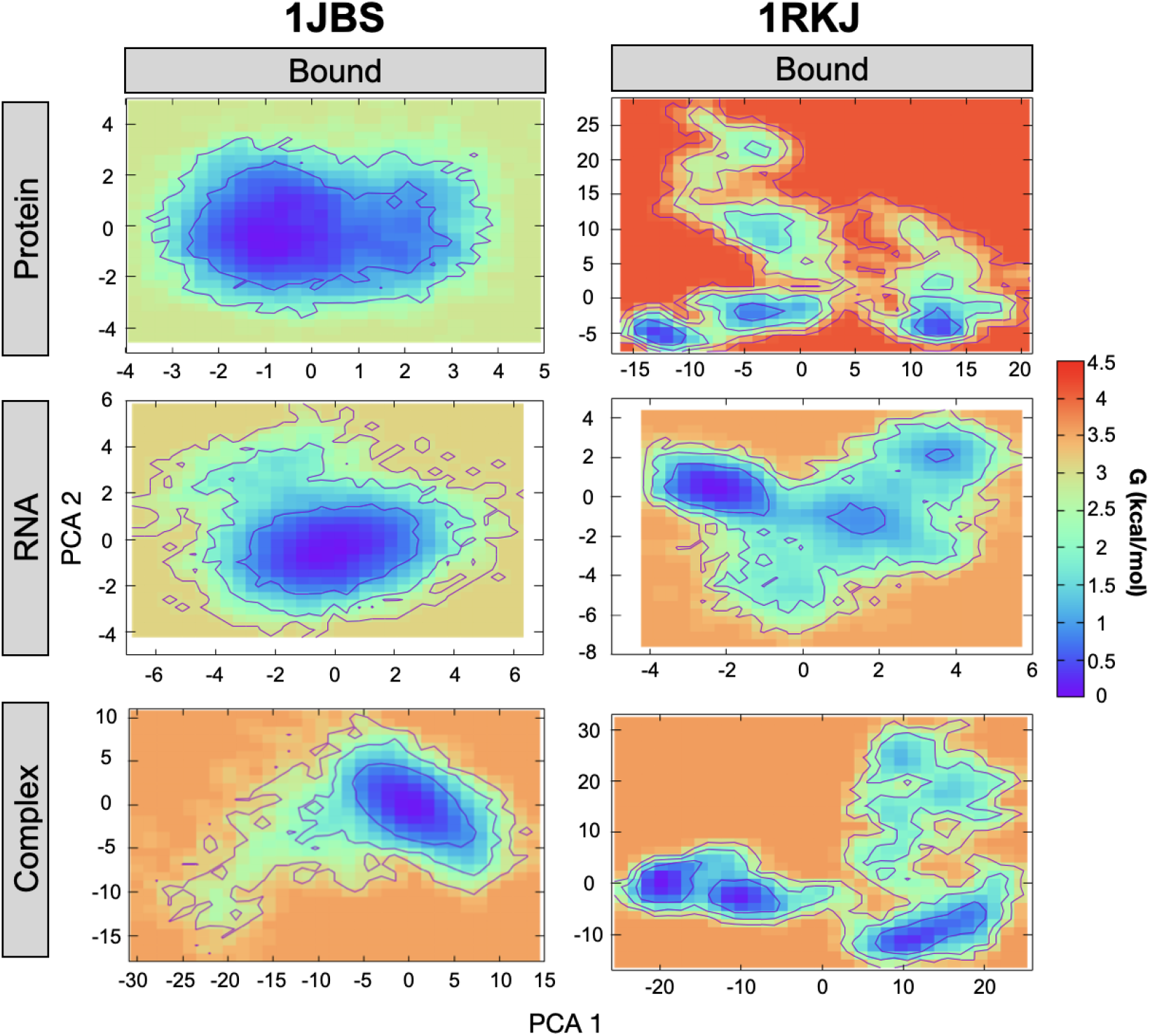
Free energy landscapes based on the first and second principal component (PCA 1 and PCA 2 respectively) for the bound simulations of 1JBS complex (left) and 1RKJ complex (right). FELs are computed separately for each chain.

The conformational changes observed along the trajectories and between the unbound and bound states can also impact the flexibility. To investigate the change of flexibility upon binding, we computed the RMSF for all the molecular species. RMSF profiles are shown in Figure S10-12, with 3D structures colored according to the change of flexibility between unbound states and complexes, defined as the difference between the RMSF computed in the bound state and the unbound ones. Based on these profiles, the flexibility of proteins and RNAs in complexes can be comparable to what is seen in the unbound forms, but also higher or lower than the one obtained in the free states. In fact, we can observe the flexibilization either a the level of the interface (e.g., 1JBS) or at sites distant from the interface (e.g., 1ASY), probably due to allosteric effects.

All these results highlight the dynamics of the protein and RNA that can impact at different level the complex and suggests that different interfaces may be present. However, no direct information on the interface and their dynamics is provided highlighting the need to more deeply investigate these aspects.

#### Puckering analysis: Comparison between unbound and bound forms

As observed from the FEL and the structural clusters, RNA molecules can undergo conformational changes in the unbound and bound state and change their flexibility upon binding as shown by the difference of RMSF. For RNA molecules, another key element is the ribose that can assume different conformations. To characterize the flexibility of the ribose and the impact of the interaction between protein and RNA on its conformation and flexibility, in particular for residues at the interface, we computed the sugar pucker as defined in the Materiel and Methods section and we compared the ribose puckering between unbound and bound forms.

All the results are reported in Figure 3 and in Figure S53-57. As expected, the ribose assumes mostly the C3’-*endo* conformation in both the unbound and bound form. However, in the bound form we can observe some changes in the ribose conformation. For example for 1RKJ complex, Figure 3 A highlight the influence of protein-RNA interaction on the sugar pucker. For the unbound RNA, C3’-*endo* form, and more widely A-like family, are the predominant ones except for residues 10 and 11. On the contrary, the bound form has a different layout, with an increase of C2’-*endo* form, and B-like family. This change in the conformation of the sugar occurs for residues that are highly involved in interactions with the protein, as illustrated by green bar and stars and in Figure 3 B. The same trend concerning interface residues as 1RKJ is found for the other complexes. As with 1ASY, we can also point out a change in the forms of the B-like family towards the A-like family between the unbound and bound forms for residues not present at the interface, such as for residues 15 to 20 and 50 to 60 of this complex. An exception is represented by 2VPL complex, which showed no major changes in ribose pucker in the absence or in the presence of the protein. Finally, for all systems, no puckering changes were observed in the presence or in the absence of hydrogen bonds involving the 2*^′^*-OH.

**Figure 3:**
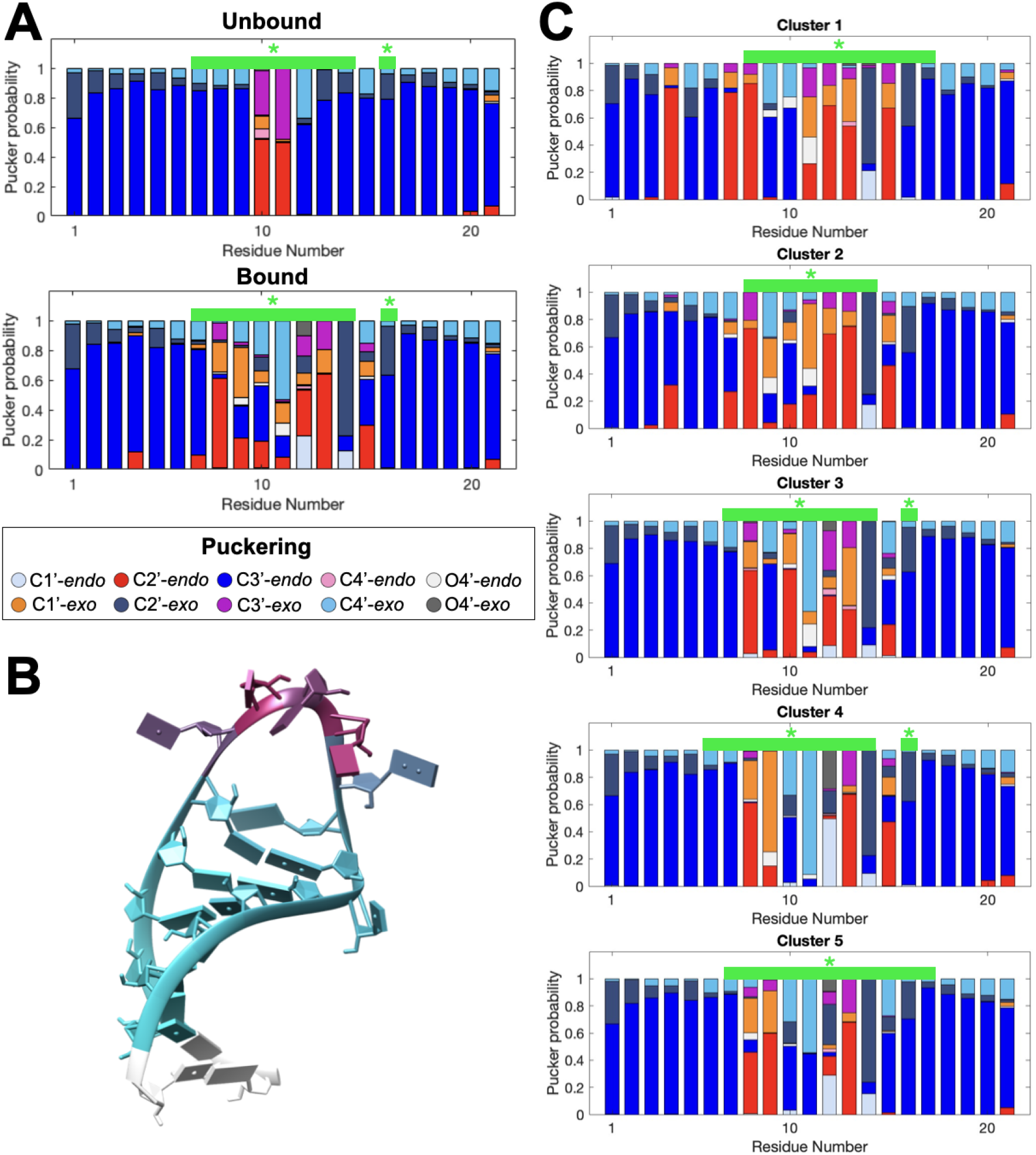
A. Ribose pucker probability of RNA unbound (top) and bound (bottom) for 1RKJ represented with stacked bar plot. B. RNA structure colored by interface contact for 1RKJ. White: not interface residues, cyan: residues at interface with a low frequency and pink: residues at interface with high frequency. C. Ribose pucker probability of RNA for each interface cluster of 1RKJ. For A and C, green bar and stars highlight the RNA residues involved in contacts with protein, with a frequency higher than or equal to 0.75. B-like family: Red: C2’-*endo*, Pink: C4’-*endo*, Orange red: C1’-*exo*, Magenta: C3’-*exo*. A-like family: Blue: C3’endo, light blue: C1’-*endo*, dark blue: C2’-*exo*, cyan: C4’-*exo*. Light grey: O4’-*endo*, dark grey: O4’-*exo*.

All these results highlight the impact of protein-RNA interaction on RNA sugar conformation and suggest a possible role in the stability of the interface. Moreover, it is known that C2’-*endo* and intermediate forms, enhanced in the bound state, should be involved in RNA reactivity and play a role in the interaction between proteins and RNA.^77,96,97^ The reactivity data as SHAPE have been largely increased in the last years with respect to available 3D structures and they are used to add soft constraints on the prediction of 2D structures for free and bound RNA.^98^ However, little is known on the relationship between the change in biochemical reactivity (like SHAPE) upon binding, the ribose pucker and the interface. Hence in this context, our findings represent an essential starting point to conduct this kind of analysis.

### Characterization of the interface

#### Geometric interface parameters along MD simulations

To get insights on the geometry of the interface, we computed the geometric interface parameters, the ΔASA, the gap volume and the gap index, along the MD trajectories and on the experimental starting structures, to investigate the impact of the dynamics on these parameters. Figure 4 and Figure S22-S27 show the distribution and Table S6 reports the relative values. All geometric interface parameters show variability during the simulation. While the initial values may deviate from the crystallographic ones, in the majority of cases, the simulations explore values in proximity to the crystallographic ones. By comparing these properties between the experimental structure, the starting structure (*t* = 0) and the average along the MD simulation, we highlighted different profiles. The 3IEV complex shows the greatest stability for the 3 parameters studied, while 2ZM5 and 1MMS, which are very stable for the gap index, show greater deviation for the other two. The greatest deviation of the gap index values, both experimental and baseline, is shown for the 1OOA and 1JBS complexes. However, this pattern is not found for ΔASA and gap volume, properties for which these complexes have an average deviation approximately equal to the other cases. These results show the considerable variability of the different complexes studied. Moreover, we observed on average a poorer packing of the complexes under investigation (as indicated by the large gap index as shown in Table S6 - average gap index equal to 3.91 +/- 2.26) than protein-ssDNA or protein-dsDNA as reported in previous works. ^27^ The only exception is provided by 3IEV with an average gap index equal to 1. This difference may result from the complex secondary structures that the other RNA molecules form. In fact, in the case of 3IEV the full RNA sequence is unpaired allowing the very close approach of the protein.

**Figure 4:**
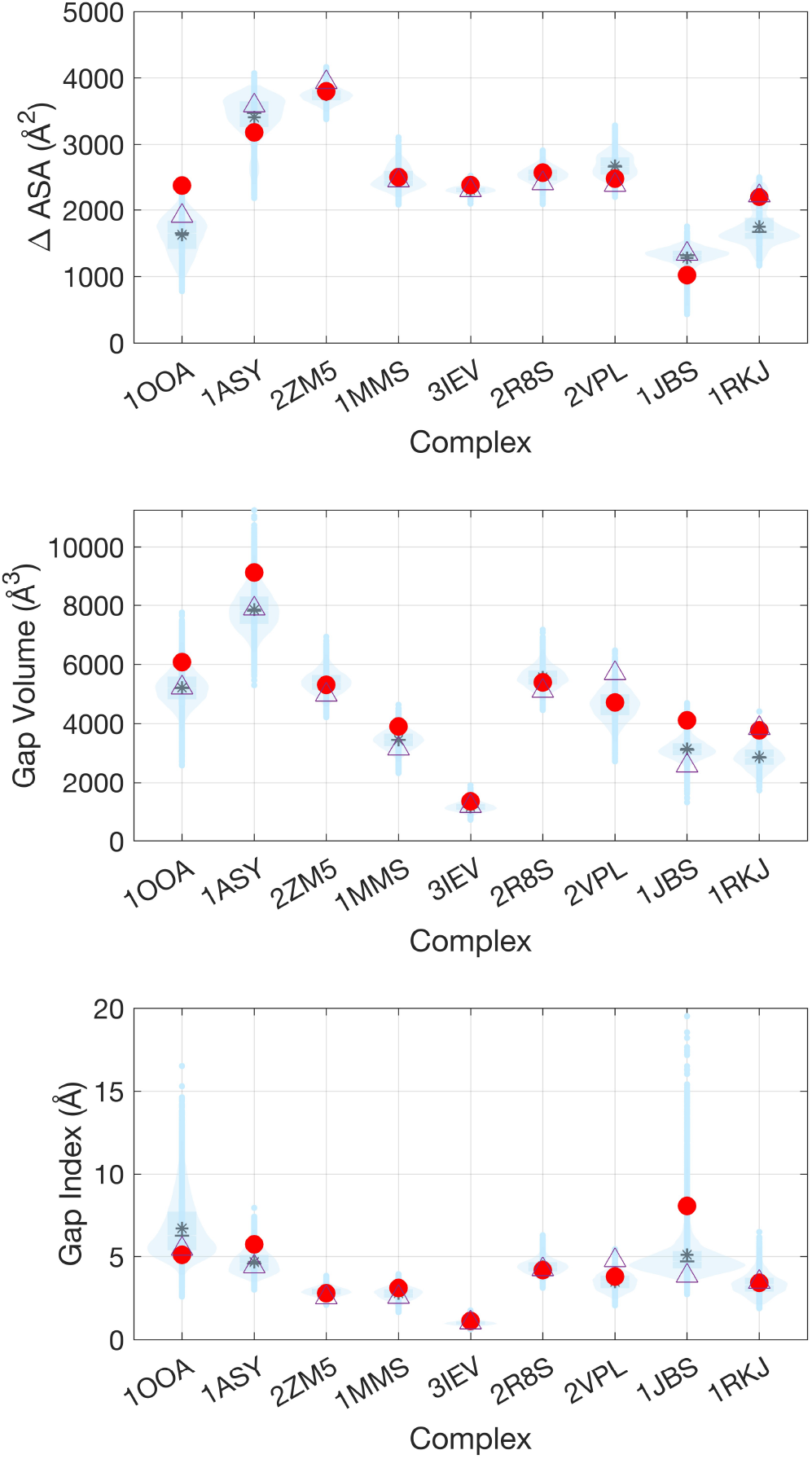
Distribution of geometric interface parameters for the nine complex represented in light blue as violin plots, with the average value indicated by an asterisk: Interface size measured by ΔASA (top), gap volume (middel) and gap index (bottom). Red filled circle: starting structure for MD production (*t* = 0). Purple triangle: experimental structure.

#### Non-bonded interactions via stacking and hydrogen bonds at the interface

To better characterize the interfaces, we also computed the inter HBs, as reported in Table S8 and S9. The total number of HBs (HB*_tot_*) depends on the system and its average varies between 6 for 1JBS and 27 for 2ZM5. To evaluate the impact of the interactions on the ribose, we also computed the number of HB that involves the 2’-OH of the ribose (HB_2_*′_−OH_*) as mentioned above and on average less than one interaction is found along the simulation.

No correlation is found between the average of HB*_tot_* and HB_2_*′_−OH_*. In fact for example for 2ZM5 only one HB that involves the 2’-OH of the ribose is observed on average out of 27. For 2R8S the HB_2_*′_−OH_* between Gly57 and G200 is present 60%. For 2VPL, the HB_2_*′_−OH_* between the Thr217 and G10 has a time occurrence equal to 96%. It is therefore possible to have HBs between the 2’-OH of the ribose and the protein residues, but these are only present occasionally in most complexes.

Finally, we also computed the total number of stacking interactions along the trajectory, its average and standard deviation (see Table S7). For 1JBS, 1MMS and 1OOA no stacking interactions between the protein and the RNA molecule are observed. One stacking interaction is observed 99.4 %, 99.8 % and 34.4 % of time along the trajectory for 1ASY, 2ZM5 and 2VPL, respectively. Two stacking interactions are observed more than 90% and 60% of time for 3IEV and 1RKJ. Finally, for 2R8S, U135 and U185 are always stacked during the simulation, oscillating between Tyr49, Tyr55 and Tyr322 and between Tyr246 and Tyr271, respectively. These results suggest that the nature of interactions at the interface could be different.

### Dynamics of interfaces

#### Conservation of initial interface contacts along MD simulations

Along the MD trajectories, we determined the conservation of initial interface contacts for each complex, as explained in the Material and Method. As shown in Figure S28, the preservation of initial contacts varies according to the protein-RNA complex under investigation. We highlighted two different patterns, a high very stable interface, with half of all complex maintaining at least 80% of initial contacts established (1MMS, 2R8S, 2VPL, 2ZM5 and 3IEV), and a major switch in contacts for the others complexes (1ASY, 1JBS, 1OOA and 1RKJ). The 3IEV complex, composed of *Escherichia coli* Ras-like protein bound to 3’end of 16S rRNA, has a medium size interface (ΔASA = 2,311 Å^2^) and presents the most conserved interface: about 90% of conserved contacts during the 500 ns of simulation. The MiaA protein complex with tRNA, 2ZM5, with the largest surface area (ΔASA = 3,937 Å^2^) also has a very stable interface (80%). Concerning 1JBS complex, the restrictocin and a 29-mer SRD RNA analog, with the smallest surface area (ΔASA = 1,341 Å^2^), the complex looses the majority of the initial contacts at the beginning of the MD simulation. The fraction of conserved contacts is decreasing to 25% at 15ns, then goes back to 75% during the rest of the simulation. On the contrary, 1OOA and 1RKJ retain less than 50% of the initial contacts. The highest change is noticed for 1RKJ complex, formed by the two N-terminal RNA-binding domains of nucleolin and a pre-rRNA target, (ΔASA = 2,218 Å^2^) with only 37% of preserved contacts. Regarding 1OOA complex, distinct conservation levels are observed suggesting different substates of protein-RNA interfaces, one at 50% and another one at 25%.

To better characterize the interface contacts and their interactions, we also investigated the conservation of the initial contacts and the total number of contacts along the trajectories by categorizing them based on the nature of the amino acids in interactions (i.e., polar or non-polar), see Figure S68. Furthermore, we investigated the evolution of stacking interactions between the RNA and the protein along MD simulations. As expected, there are more contacts formed by polar residues in all interfaces (around 70 to 80% of polar amino acid/RNA contacts). However, there are two exceptions: 1ASY and 3IEV. For 1ASY, there is a loss of polar contacts along the trajectory, while the number of apolar contacts remains constant. For 3IEV, there is also a loss of polar contacts and the majority of contacts are indeed contributed by apolar residues. These findings suggest that the observed interface rearrangements have the potential to alter the characteristics of the interfaces in a manner that depends on the specific case. Finally, the different physico-chemical natures of the interface could also have an impact on the mechanism of the recognition and the role of water molecules at the interface.

#### Analysis of the substates of protein-RNA interfaces

As explained in the Material and Methods section, the interface similarity was measured by the Jaccard index *J*, and then we performed hierarchical clustering based on interface contacts. This analysis unveiled the presence of distinct substates at the interfaces. Clustering interface results are reported in Figure 5 for 1RKJ complex and in the Figure S30-S37 for the other complexes. In the case of the two N-terminal RNA-binding domains of nucleolin (RBD12)/pre-rRNA (b2NRE) complex, 1RKJ, the Jaccard similarity matrix shows five purple squares along the diagonal with yellow off-diagonal rectangles, indicating five portions of the trajectory with high internal interface similarity and well distinct from each other (Figure 5A). Based on the dendrogram (Figure 5B), five main clusters corresponding to long-lived and well-populated substates (Figure 5C) are obtained. Except for cluster 1, the other clusters are evenly distributed along the simulation 5C.

**Figure 5:**
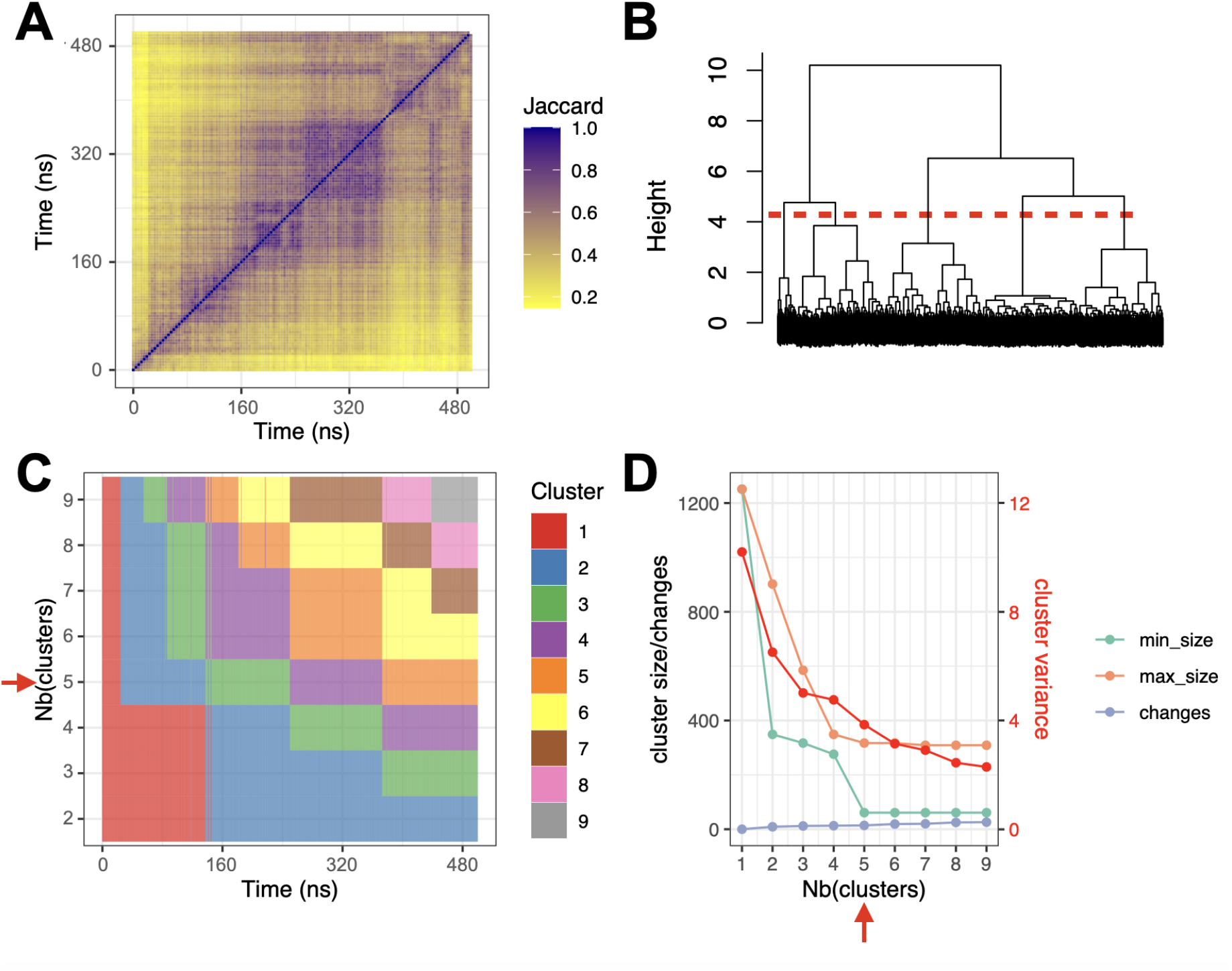
Clustering results for the 1RKJ complex. A. Jaccard similarity matrix. For each pair of snapshots, a yellow pixel indicates low interface similarity, and a purple pixel indicates high interface similarity. B. Clustering dendrogram. C. Cluster membership along the MD simulation, for different numbers of clusters. D. Cluster size, number of changes, and intra-cluster variance in red, for different numbers of clusters; min size: size of the smallest cluster, max size: size of the largest cluster, changes: number of cluster changes during the simulation. The red dashed line in panel B. and the red arrows in panels C.-D. indicate the optimal number of clusters.

In regards to the results, four of the nine complexes analyzed in this study exhibit different substates of their protein-RNA interaction interface. We obtained 3 clusters for 1ASY, 2 clusters for 1JBS and 4 clusters for 1OOA complex. Remarkably, interface clusters were found for the complexes with major switch in initial contacts, as described in the previous paragraph. That is confirmed by the conservation of initial interface contacts for the five other complexes (1MMS, 2R8S, 2VPL, 2ZM5 and 3IEV). Moreover, no relationship between ΔASA and the number of clusters is obtained, as previously also observed for protein-protein complexes.^42^

To explore the structural differences between substates, we computed the variance of each contact across the interface clusters. The matrix representation of the contact variance for 1RKJ is shown in Figure 6A. As highlighted in Figure 6B, at RNA level, all the residues initially in contact with the protein have high contact variance between clusters. In fact, both nucleotides involved in canonical and non-canonical pairing have a large contact variance. Same observation is made for the nucleolin RBD12 residues. Thus, a majority of contacts vary between the five clusters that are characterized by both conformational changes of the protein and the RNA and the rearrangement of the interface, as shown by the FEL in Figure S17. The clusters differ from one to another for the loop conformations and its relative orientation with respect to the two RBDs domains and the linker. In cluster 3 and 5 the bulged uridine base is involved at the interface. The presence of different alternative interfaces is compatible with the experimental observations where a lower stability with respect to the RBD12/nucleolin recognition element (NRE) complex is also observed.^99^ Based on that, we can speculate that our findings have a biological relevance since a low-level stability of the RBD12/b2NRE complex would allow for fast dissociation of the protein from the rRNA. Moreover, the identification of alternative interfaces, not accessible experimentally, could help to better understand the communication in the complex and the role of the different parts of the RNA structure in the recognition and the allostery in the complex.

**Figure 6:**
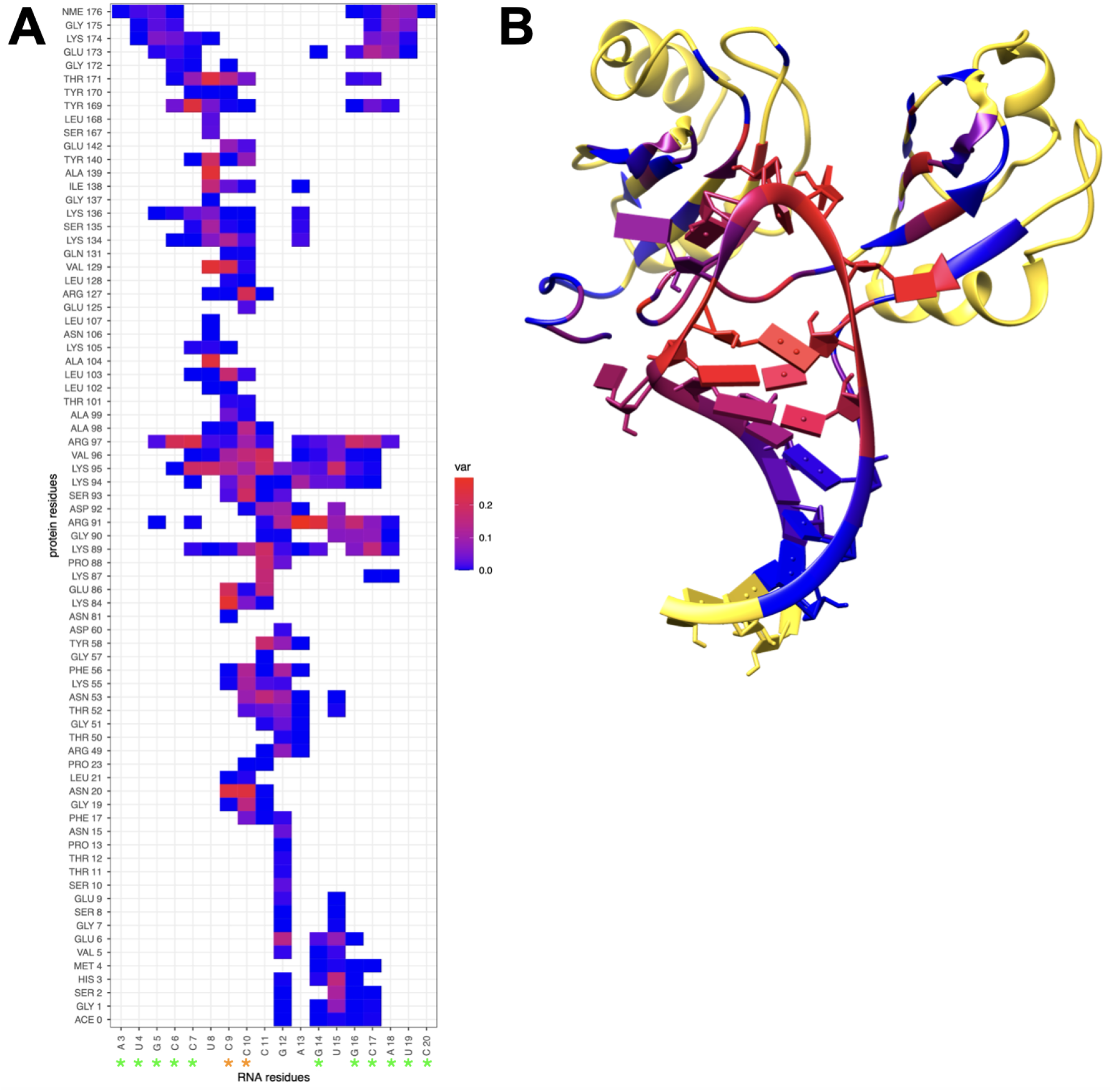
Contact variance in complex 1RKJ. A. Matrix representation of each contact variance across the five interface clusters. Blue: low variance; Red: high variance; Green stars: canonical pairing; Orange stars: non-canonical pairing. B. 3D structure representation of 1RKJ complex. Interface residues are colored by the maximum of contact variance (blue to red) and the other residues are colored in yellow.

Different characteristics are found for the other complexes. For 1ASY, 1JBS and 1OOA complexes, the matrices are available in Figure S38-S40. In contrast to 1RKJ, a majority of contacts, in the case of 1ASY, have stable frequencies across clusters. The results highlight one region at the protein-RNA interface which varies considerably for the three interface clusters, between C649 to C674 for RNA and Asp229 to Pro557 for protein. The changes at the interface are due to the conformational changes of protein and RNA, local rearrangements and relative orientation between the protein and RNA as shown in the FELs (see Figure S13). In this case, the alternative interfaces and the variation of contacts involve mostly the recognition sites: the single-stranded fragment at the beginning and the end of the anticodon stem and loop. Therefore the presence of interface substates with biological relevance allows one to better understand the recognition, as presented below.

In the case of 1JBS complex localized variations are also observed, but the origin is different. In fact, a rigid body movement occurs between the protein and the RNA as highlighted in the FEL (Figure S14). In fact, the two interfaces involve the nucleotides involved in the cleavage. In the cluster 2, the possibility of the cleavage is completely. Hence, we can hypothesize that the analysis of the two states and how to switch from one to another should allow us to get insight on the mechanisms behind and how it is activated. Finally, regarding the 1OOA complex, variations in different regions of the protein and RNA are observed. Based on FEL profiles reported in Figure S16 and the RMSD time series in Figure S4, these variations are mostly due to the dynamics of RNA which preserves the crucial intra HBs^100^ and interface rearrangements, although the protein can also visits different conformational states and its dynamics influences the interface. In other words, the stability of the complex is due to different patterns of contacts without strongly affecting the protein conformation. While the RNA aptamer is not natural, it binds the NF-*κ*B with high affinity and it has been suggested to be a promising therapeutics capable to inhibit the NF-*κ*B transcriptor factor.^100,101^ Hence, the identification of alternative interface substates allows us to determine the different contacts in terms of base pairing and stacking for example. These pieces of information are not accessible via x-ray crystallography, like the highly dynamics of the protein-RNA interface. These pieces of information are essential to better engineering an aptamer inhibitor of this transcriptor factor.

#### Characterization of interface substates

To characterize the different interface substates visited along MD simulations in the different systems, for each cluster we computed several interface properties (like the HBs and stacking at the interface) and the RNA puckering. Regarding the geometric interface parameters (ΔASA, gap volume and gap index) already computed for the entire trajectory (see above), their average computed for each interface cluster are reported in Table S6 and distributions are shown in Figure S22-S27 (complexes with only one cluster are also integrated for the sake of completeness). For all the complexes studied, the results indicate a clear difference between the clusters for the following two parameters: ΔASA and the gap index. Notably, the gap volume exhibited a a noticeable difference solely within the 1RKJ complex.

When examining the distribution of these parameters within each cluster, both the gap index and gap volume are distributed across the same region for all clusters for all complexes. However, the ΔASA property exhibits distinct patterns depending on the cluster being analyzed. For example for 1JBS the two clusters sample different ΔASA ranges, indicating interfaces of different sizes. The last observation also applies to complexes 1ASY, 1OOA and 1RKJ, where the different clusters correspond to interfaces of different size.

We also computed the total number of HB between the protein and the RNA along each cluster. The total number of HBs can change radically depending on the complex observed, with, for example, an average of 13 for the second cluster of 1ASY versus 20 for the third cluster. The same applies to 1JBS, which loses half its HBs between cluster 1 and 2 (from 6 to 3). On the contrary, for 1OOA and 1RKJ the differences are less noticeable between clusters.

Since the 2*^′^*-hydroxyl group of the ribose both can be involved in the interactions with the protein via hydrogen bonds and is related to the reactivity/stability of the RNA molecules, for each cluster we computed the HBs between this chemical group and the protein (HB_2_*′_−OH_*, see Table S9) and we analyzed if there was or not a relationship with the change of the puckering upon binding. For 1RKJ, one HB_2_*′_−OH_* between Tyr140 and U8 was only observed in cluster 1 and 2 with a time occurrence of 87% and 78%, respectively. The nucleotide involved in the HB changes its puckering conformation upon binding (see Figure 3C). The other three clusters present different HB_2_*′_−OH_* with an occurrence below 30%. For the clusters of the other complexes, the HB_2_*′_−OH_*obtained have an occurrence below 50%. Hence, although other nucleotides are at the interface (indicated by the green bar and stars in Figure 3C and Figure S53 and S54) and present different distributions of sugar pucker conformations with respect to the unbound form, they do not interact with the protein via HB_2_*′_−OH_*. It is important to point out that each cluster presents a different distribution of sugar pucker conformations and the nucleotides at the interfaces present a puckering mostly belonging to the B-like family. These results suggest that the reactivity of the RNA should also change based on the interface substate and depending on the chemical probe these changes could be captured or not.

Finally, we compared the clusters by taking into account the stacking interactions at the interface (see Table S7). For 1ASY, one stacking interaction between Phe127 and U635 is observed 99% of the time along each cluster. On the contrary, for 1OOA and 1JBS no stacking interactions with a time occurance above 50% are obtained. For 1OOA, a stacking interaction between His63 and U14 is present 15.5% of time along the MD simulation, and in particular it is observed 35.8% of the time in cluster 2. For 1RKJ, we obtained two prevalent stacking interactions (Phe56-C11 and Tyr140-C9) with an occurrence above 60% along the MD trajectory, but they are differently distributed in the clusters. For example, the stacking interaction between Phe17 and C11 is present 3.8% of the time during the simulation but is observed 73.7% of the time in cluster 1 and it is absent in the other clusters. Another example is given by the stacking between Phe17 and C10 that is present 78.9% of the time during the simulation but absent in cluster 1. All these results again highlighted that protein-RNA interface dynamics vary significantly depending on the complex studied and the interface cluster.

### Dynamical Network Analysis

To investigate the impact of the alternative interfaces obtained by our interface clustering analysis, we also performed a dynamical network analysis (DNA) on each cluster as explained above. For each complex, we compared the number of communities predicted, their location, the edges with the highest betweenness and the path obtained. In dynamical networks, a community is identified when the network’s nodes can be organized into clusters where nodes within each cluster exhibit strong internal connections.

For the 1RKJ complex, Figure 7 summarizes the results obtained (see also Figure S49-S52). In each cluster nine communities are at least identified. The different clusters differ from one another for the communities at the interface (see Figure 7B). The residues at the interface belong at least to four different communities. Here, the communities can include only residues of the protein, or only residues of the RNA, otherwise some residues of both partners for example at the interface. Hence, the differences at the level of the contacts and communities bring to different dynamical networks based on the interface substate. For example, within the clusters the edges with the highest betweenness changes, as well as the communication between the two RBDs change. In cluster 1 and 2, their communication is mediated by the RNA’s loop. On the other three clusters, the upper part of the stem is involved in the communication between the RBDs, but at the same time the two RBDs can interact and communicate without involving the RNA. Therefore, the paths between the two RDBs (nodes considered: Val5 and Asp132), between the RNA and RBD1 (nodes considered:G2-P and Val5) and between the RNA and RDB2 (node involved: G2-P and Asp132) and from are different. In fact, the optimal path between the termini of the protein involves the RNA only for cluster 1 and 2 in slightly different matter. The path between the nodes G2-P and Asp132 seems to be slightly affected by the changes of the contacts at the interface, as also suggested by the presence of several edges with high betweenness in all clusters. The path between the RNA and RBD1 is the most affected by the different interfaces. In fact, in cluster 1, 3 and 5 the loop is involved. Interestingly, for cluster 5, both strands of the RNA molecule are involved in the communication. These findings allow us to hypothesize that the communication between the RGB2 and the RNA is less influenced by the alternative interface than the other possible paths. Therefore, based on this analysis, the interaction of the upper part of the stem and the protein influences the allosteric communication as well as mostly the communication within the protein and the RNA and the RGB1.

**Figure 7:**
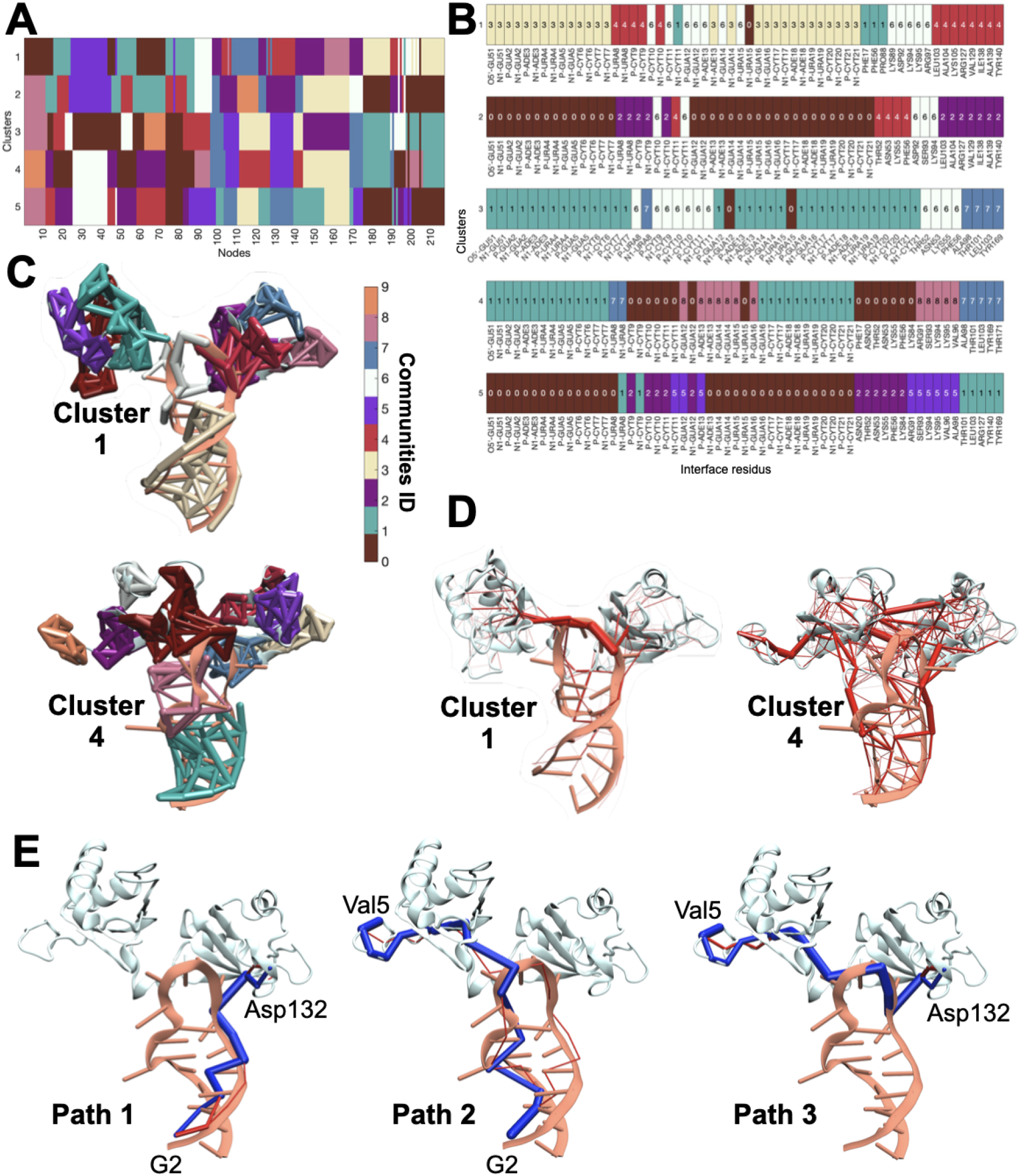
A. Community of the nodes for the different interface clusters of the 1RKJ complex. B. Community of the nodes for the different interface clusters of the 1RKJ complex for the residues at the interface between the protein and the RNA. C. Communities representation in the complex structure for the interface cluster 1 and 4. D. Edge betweenness for the interface cluster 1 and 4. Thickness depends on the betweenneess value. E. Optimal path (blue) and sub-optimal path (red) from G2 to Asp132 (left), G2 to Val5 (middle) and from Val5 to Asp132 (right).

For 1OOA, in each cluster eleven communities are identified (see Figure S46-S47). The residues at the interface belong at least to two different communities. In cluster 1, two separate communities are present for the protein and the RNA. On the contrary in the other clusters, the communities involved both residues of the protein and the RNA. The predicted dynamical networks are different and the edges with the highest betweenness vary for each cluster (see Figure S46-S47). Therefore, the communication within the complex changes based on the interface substate, as well as the path connecting the termini of the protein and the RNA and the end of the two protein domains (see Figure S48). In particular, we observed that the allosteric communication in the protein is mostly affected by the presence of alternative interfaces pointing out that the protein allostery and the communication within the protein are modulated by the protein-RNA interfaces and in particular by the conformational changes of the aptamer. These findings support the fact that this aptamer plays a crucial role in the regulation of the NF-*κ*B transcriptor factor.

For 1JBS, at least nine communities are obtained (9 for cluster 1 and 11 for cluster 2, see Figure S44). Also for these alternative substates the communities at the interface are different, but the same edge with the highest betweenness is obtained between LYS111 and G10 suggesting that this interaction is key to anchor the RNA on the protein before the cleavage since in cluster 2 several key contacts essential for this biological function are lost. Moreover, the communication between the identity residue and the cleavage sites are not affected suggesting that the RNA conformation is not affected by the alternative interfaces while the allosteric communication within the complex changes between the inactive and active state (see Figure S45).

For 1ASY, in each cluster twelve communities are at least identified (12 for cluster 1, 14 for cluster 2 and 13 for cluster 3, see Figure S42). Although the communities involved at the interface are different, the global dynamical network are similar, as well as the edges with the highest betweenness (see Figure S43). We also computed the optimal and sub-optimal paths between the node C602-P and the node U632-P for their biological relevance and no relevant differences are observed as function of the interface cluster (see Figure S43). Hence, in other words, the presence of different interface contacts does not strongly impact the communication at the interface and their allostery.

### Water analysis

To get more insights into the protein-RNA interfaces and in particular into the role of the water molecules at the interface and their interactions, we also evaluated the evolution of interfacial water molecules (water molecules whose distance, *d_w_*, from both protein and RNA is below 4 Å) along the MD simulations, as well as the contacts between interface residues and interface water molecules.

#### Evolution of interfacial waters

We monitored the number of interfacial water molecules along time. The average number of interfacial water molecules and their standard deviation (*sd*) are shown in Table S3. Figure 8 displays the result for complex 1RKJ. The number of interfacial waters rapidly drops from around 100 to 75 during the first tens of nano-seconds of simulations and stays around this value until 250 ns, during which two sub-states are visited. Then we see two augmentations, corresponding each to a new interface substate. It appears that the interface substates correlate with the number of interfacial waters, contrary to what was seen with proteins.^42^ The results for the eight others complexes are available in the Figure S59. Similar to 1RKJ complex, a switch in the number of water molecules in contact with the residues at the interface was also observed for clusters of complexes 1ASY and 1JBS. However, the 1OOA complex, which has 4 clusters at the interface, has a quite variable number of water molecules (*sd* = 20), but the link with the cluster seems less obvious. Finally, complexes with a single cluster (1MMS, 2R8S, 2VPL, 2ZM5 and 3IEV) generally show a low variability in their number of interfacial waters.

**Figure 8:**
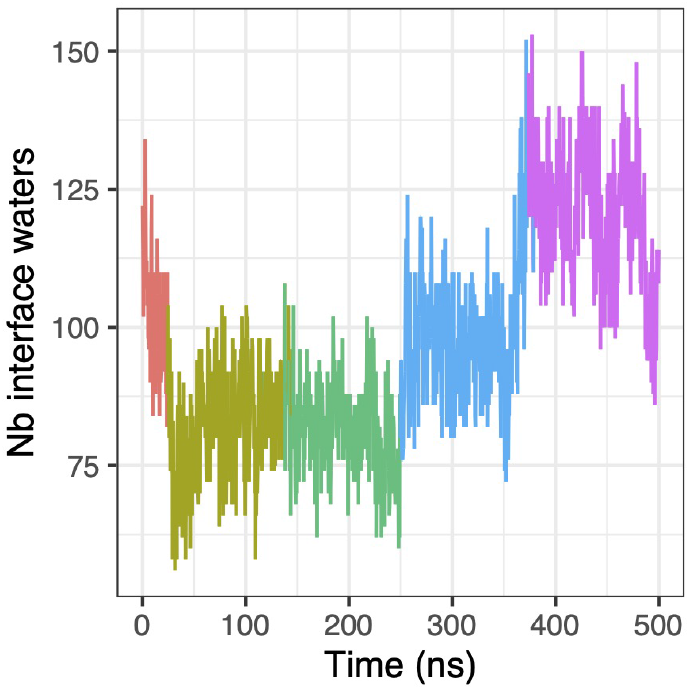
Number of interfacial water molecules in contact with interface residues, colored according to MD clusters for 1RKJ protein-RNA complex.

The stability of the total amount of interface water molecules hides the structural re-arrangement of water molecules at the interface. Moreover, the location of these water molecules at the interface differ between clusters even when the number of water molecules at the interface does not significantly change.

To rationalize these findings, we computed the Pearson correlation, as well a linear regression fit, between the average number of interfacial water molecules (*n_iwm_*) and geometrical properties obtained for each cluster (in the case of one cluster the complete trajectory was taken into account), as shown in Figure 9. A significant correlation was obtained between both ΔASA and the number of interfacial water molecules (*ρ* = 0.8345) and Gap Volume and the number of interfacial water molecules (*ρ* = 0.9044), but no significant difference between these two dependent correlations was observed (*p*-value equal to 0.2837). Surprisingly, a non-linear trend was obtained for the number of water molecules as a function of the gap index. In fact, for two clusters (the cluster 2 of 1JBS and the cluster 3 of 1OOA) a large value of gap index is associated to a small number of interfacial water molecules.

**Figure 9:**
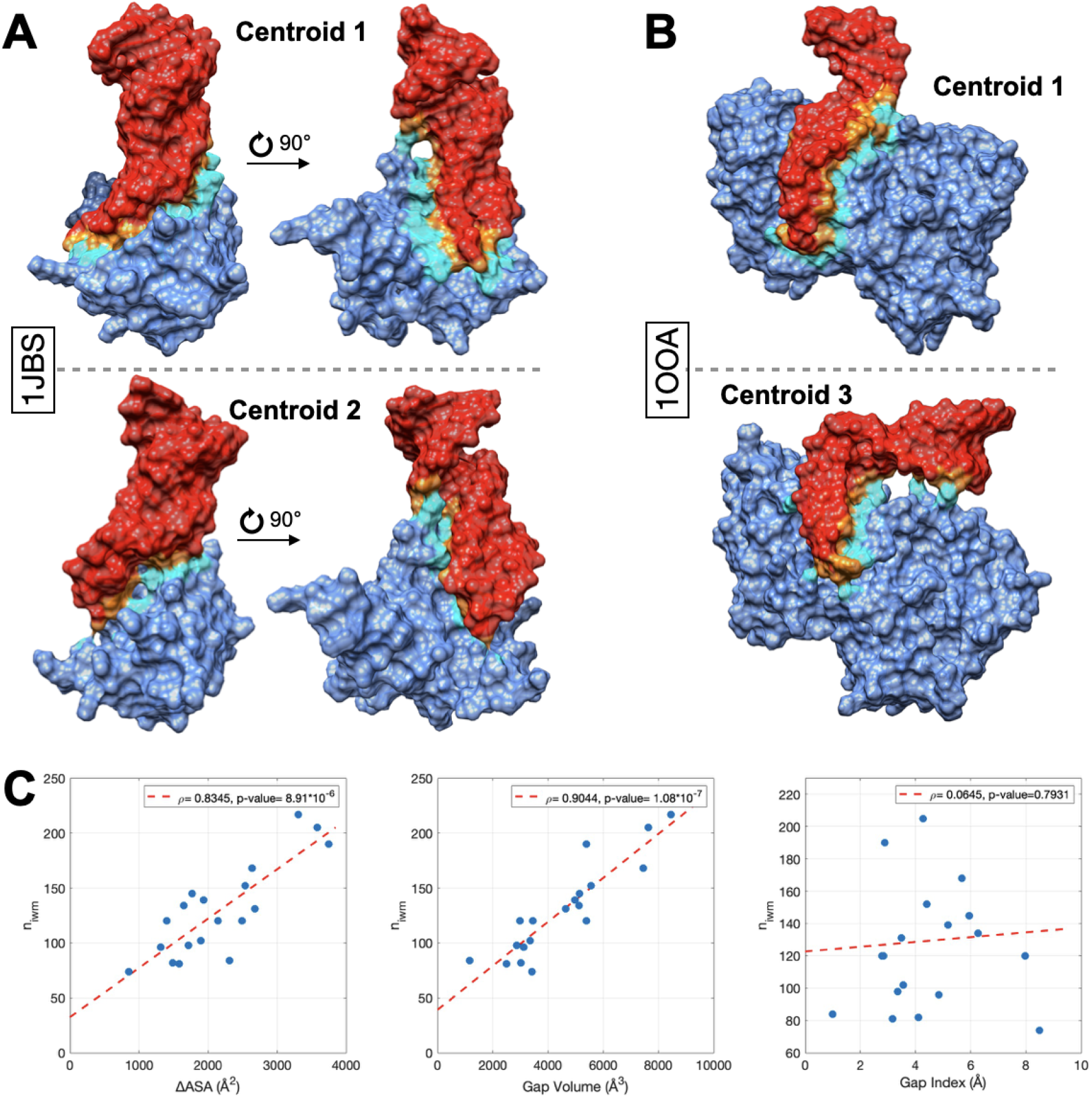
A. Centroid of cluster 1 and 2 obtained for 1JBS. B. Centroid of cluster 1 and 3 obtained for 1OOA. For A and B, RNA = red; protein = blue; Interface residue of RNA = orange and protein = cyan. C. Scatter plot of the number of interfacial water molecules (*n_iwm_*) as a function of the interface size (ΔASA, left), the gap volume (middle) and the gap index (right). In each panel, the red dash line is the linear regression fit and the Pearson *ρ* coefficient and associated *p*-values are annotated in box.

Based on these results, we can hypothesize that the packing between the protein and the RNA molecule at the interface may have a crucial role on the number of water molecules at the interface and their dynamics. For low values of gap index (dense packing as observed in protein-protein complexes) although the location of water molecules at the interface differ between clusters, the layer of water molecules is thin and can easily and fast adapt when a new interface is formed. In an intermediate regime, the water molecules at the interface can fill a larger volume (correlation between the number of interfacial molecules and the gap volume) and possible confinement effects on the water dynamics can be hypothesized since some molecules could be trapped. Finally when the gap volume is greater than 6-7 Å, the packing of the interface is loose and an opening of the interface is observed in our systems (see Figure 9 for the superimposition of the centroids for the cluster 1 and the cluster 2 of 1JBS and for the cluster 1 and cluster 3 of 1OOA). In this case, the water molecules can easily escape and despite the large gap volume, the change of accessible surface area has lower value and several molecules are not more involved in the interaction with both partners, hence their dynamics should be similar to the bulk water. We can finally speculate that the difference in structural water at the interface could also have a biological relevance since it is known that water molecules play a crucial role in the recognition and in our study the alternative interface not only have a different number of water molecules but also the location of these water molecules at the interface differ between clusters.

#### Contacts between interface residues and interface water molecules

We examined the interactions between interface residues and interface water molecules and compared these findings with the interface clustering analysis. To do so, we computed the number of interfacial water molecules in contact with residues at the interface along the MD trajectory. Figure 10 presents the distribution of the number of interface water molecules, in each cluster of 1RKJ complex, for all the residues with a variable number of contacts interfacial waters (here, with a standard deviation over the trajectory greater than 2). This figure allows to clearly highlight the residues with different solvation states between clusters. Some residues display a variable number of contacts with interface water molecules, with no evident correlation with interface clusters, like residues Asn106 and Asn53, and RNA nucleobases G12 and G16. On the contrary, some residues have specific profiles depending on the cluster. For examples, residues Gly175, Gly7, Lys174 and Met4, and RNA nucleobases C20, C21, C7 have a few number of contacts with interfaces water molecules in clusters 1-4, but high number of contacts in cluster 5. Thus, some interface residues have a variable number of contacts with interfacial waters throughout the simulations, while other residues exhibit specific signatures according to the substates. Same observation was made for the complexes with different substates (1ASY, 1JBS and 1OOA), as shown in the Figures S60-S67 (for the sake of completeness, single cluster complexes are also included). This analysis provides another perspective on interfaces, portraying them as dynamic entities capable of rearranging contacts between residues from interacting chains and water molecules.

**Figure 10:**
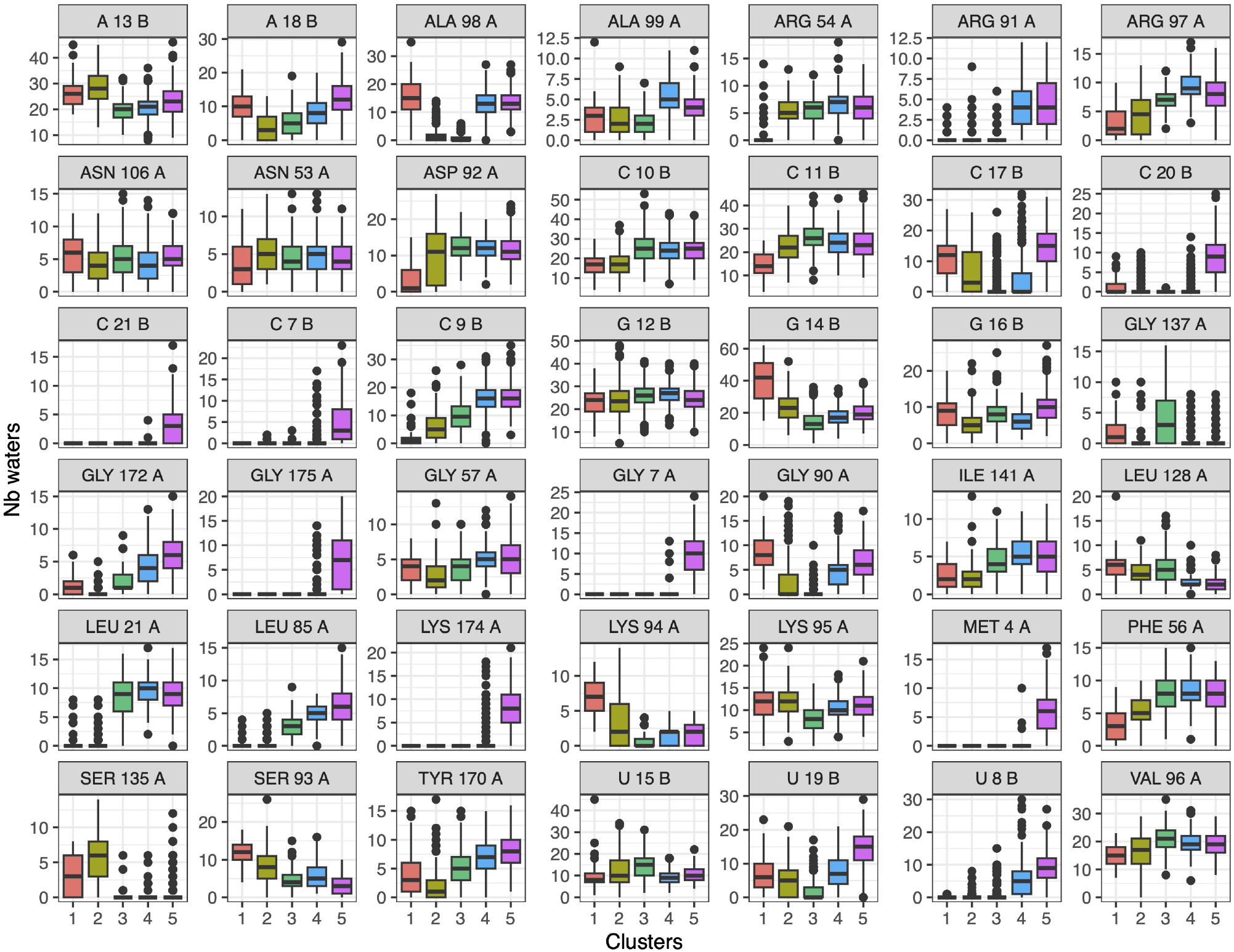
Number of interfacial water molecules in contact with interface residues, in each MD clusters for 1RKJ complex. Distributions are shown as boxplots in each interface cluster.

#### Contacts mediated by water molecules at the interface

To enhance our understanding of water-related contacts and their impact on interactions, we also investigated contacts mediated by water molecules at the interface. These contacts involve triplets consisting of a residue from one chain in contact with a water molecule, which at its turn is in contact with a residue from the other chain. A water-mediated contact between two residues is identified by the presence of such triplets in the snapshots. It is important to note that the water molecule connecting the residues may vary between snapshots if its dynamics is faster than the time between each snapshot. Since this study does not delve into water dynamics, our focus remains on assessing the stability of the residue pairs involved in water-mediated contacts and their evolution.

First, we extracted the water-mediated contacts and evaluated their preservation along the MD simulations and compared these results with the direct residue-residue preserved contacts (see Figure S68 A and B). This comparison allowed us to get insights into the potential relationship between the loss of initial contacts and water molecules. The decrease in initial contacts seems to be only partially related to the loss of contacts mediated by water molecules. We also investigated the evolution of the number of water-mediated contacts along simulations and the number of direct residue-residue contacts (see Figure S68 C and D). For example, in the case of 1ASY, the number of contacts decreases and then increases again, while the number of preserved contacts does not increase, meaning that the complex forms new contacts. For 1OOA, the complex moves away from its native contacts throughout the simulation and moves towards a configuration with fewer contacts since there is an opening of the interface that also induces the loss of water molecules at the interface. Although the number of water-mediated contacts along simulations usually follows the same evolution as the number of direct residue-residue contacts, there are some exceptions. In the case of 2VPL and 2RS8, along the trajectory on the one hand a decrease of the number of contacts can be observed and on the other an increasing number of contacts mediated by water can be obtained. These different trends are due to two possible cases obtained by our MD simulations: remodelling of the interface (in this case there is a correlation between the number of contacts and the number of contacts mediated by the water, which are at the periphery), and opening of the interface (in this case there is an anti-correlation between number of contacts between residues and number of contacts mediated by water, because water molecules can more easily introduce themselves and have ample space to occupy between the two partners). In one case, the contacts mediated by water follow the contacts between residues, in the second scenario they replace them.

To better characterize the details of the contacts mediated by water molecules, we computed their frequency for each complex and their interface substates if present (see Figure S69-S75). First, for all complexes, multiple water-mediated contacts are observed at their interface. These contacts can involve residues that are less exposed in the experimental structures. Furthermore, we note that certain contacts remain relatively stable throughout the simulation period, appearing 75% of the time or more. This underscores the structural significance of interface water molecules. Moreover, as expected, numerous contacts involve positively charged protein residues. However, occasionally these contacts also involve negatively charged residues from protein (Asp and Glu), illustrating the screening effect of water molecules that enables their presence at the protein-RNA interface and stabilizes their interactions with the RNA molecules. Lastly, sometimes non-polar protein residues are also involved in these contacts.

To compare the interfaces substates, we also computed the variance of the contacts mediated by water molecules (see Figure S69-S75). This analysis allows to highlight the contacts mediated by water conserved in the different alternative interfaces for which we can hypothesize that the water molecules could be trapped and slow down their dynamics, as well as the differences between substates in terms of solvation since the loss of a water mediated contacts means either the formation of a direct contact between the pair of residues or the fact that one residue or both are no longer part of the interface. For example, for 1RKJ, Lys94 and Arg97 are involved 95% and 49% of the time at least in one water-mediated interaction along the MD trajectory, respectively. The variance for Lys94 and Arg97 is respectively below 0.17 and 0.24. Our results confirm the hypothesis that these residues are involved in water-mediated interactions with the RNA.^99^ Moreover, our analysis allows to identify other residues involved in these interactions that are not accessible experimentally, like Phe56 (frequency equal to 0.65 and variance below 0.07) which is known to be involved at the interface. Interestingly, in the case of 1JBS, a conserved contact mediated by water is obtained between Glu95 and A17, which is placed in the cleavage site. For 1OOA, conserved contacts mediated by water are obtained between Glu60 and Gly66, a region of the complex that is hypothesized to be critical for its stability. Our results strengthen this hypothesis and propose to explain the origin of the stability. For 1ASY, the conserved contacts mediated by water are at the end of the anticodon stem and loop (G630-G639), a crucial region for the recognition. Both positively and negatively charged residues are conserved. For example, the contact between Asp176 and G634 has a frequency of 0.34 with a variance equal to 0.02. Our analysis suggests that the conserved contacts mediated by water are biologically relevant allowing to determine other key residues involved in the stability of the interface in combination with the pieces of information obtained by the analysis of direct contacts. Moreover, we can speculate that the comparison between different clusters at the level of the residues with high variance should again allow to better understand the role of water molecules in the dynamics of the interface and how the water molecules influence the stability, the possible allosteric mechanisms and pathways.

## Conclusions

In this work, we aimed at setting up a computational strategy that combines different metrics and tools capable of investigating and characterizing the dynamics of protein-RNA interfaces. To do so, we defined a benchmark of nine complexes with different RNA motifs, interface sizes and geometries. We performed all-atom molecular simulations on the complexes, as well as on their bound and unbound forms. In this context, first we investigated the conformational changes and the change of flexibility along molecular dynamics simulations and upon binding on the one hand by computing RMSD time series, RMSF profiles, and free energy landscapes, and on the other by analyzing the preservation of the RNA structure at the level of base-pairing, stacking and puckering conformations. Second, we characterized the interfaces by computing some geometrical parameters such as the buried surface area, the gap volume and the gap index, the conservation of the initial contacts, that were also categorized based on their physicochemical properties, the inter hydrogen-bonds, the stacking interactions at the interface. In this context, we also determined the presence or the absence of interface substates using a hierarchical clustering approach. Each interface cluster was characterized and compared with the other ones. To get insights on the allosteric communication within the complex and how it is impacted by the alternative interfaces, we performed a dynamical network analysis. Finally, we evaluated the role of water at the interface by computing the number and the evolution of water molecules at the interfaces, as well as the contacts mediated by them.

In the complexes under investigation, both proteins and RNA molecules can visit different conformational states in the bound and unbound form. Moreover, the dynamics of the complex and its conformational changes can be due to either the dynamics of both partners that also impacts the interface or the relative rigid movements between them. For the RNA molecules under investigation, the 2D structure is preserved as shown by the stability of base pairing and stacking along MD simulations. A change of flexibility upon binding is also observed involving either only one or both biomolecules. A rigidification or a flexibilization of the biomolecule(s) can be induced for the formation of the complex. Moreover, the distribution of the ribose puckering is affected by protein-RNA interactions since more reactive conformations are observed for the residue s at the interface. Surprisingly, no correlation has been obtained between the change of ribose pucker and the presence of hydrogen bonds involved the hydroxyl group of the ribose and the protein at the interface. As expected a poor packing of the complexes under investigation (as indicated by the large gap index expect for one complex) was observed. At the same time, a variability of the geometrical parameters and of their distributions was also detected.

Our hierarchical clustering based on the contacts at the interface highlighted that at the scale of our simulations (up to 1 *µ*s) almost half of RNA-protein complexes under investigation visit distinct interface substates. Interface substates exhibit altered interface contacts, either through direct interaction or mediated by water molecules. The geometrical properties between substates are notably influenced in a manner dependent on the specific case, as well as the number of hydrogen bonds and the stacking interactions at the interface. Regarding the RNA, the ribose pucker is also distributed differently between clusters, always with a majority for C2*^′^*-*endo* and associated B-like family forms. The adjustments of interface contacts are accompanied by shifts in the arrangement of interface waters, which at their turn influence the hydration levels of residues. It is crucial to emphasize that these alterations are not limited to the edges of the interfaces; they also impact the residues at the core of the interface. This variation could lead to distinct complex stability and contribute to the reversibility of the interaction. Moreover, the impact of these interfaces on the allostery of the complex can also be analyzed via a dynamical network analysis that allowed us to identify different paths in the complexes.

Our work highlights that we cannot any longer consider protein-RNA interfaces as static objects, but we have to take into account their dynamics. Therefore, a computational strategy which includes several descriptors and techniques as proposed here is necessary to properly characterize protein-RNA complexes considering the different main actors in their stability and dynamics. Regarding the RNA molecules, our analysis conducted on the complete MD simulations and on the interface clusters to investigate the ribose puckering, known to play a role in their reactivity and flexibility, suggests that a better characterization of the relationship between the dynamics of the interface and RNA molecules, the ribose puckering and chemical reactivity should improve our capability to predict protein-RNA complexes and distinguish the origin of the change of chemical reactivity due to the interface and the RNA conformational changes. By combining the analysis of FELs and the characterization of interface clusters, we were able to better understand the origin of the variability of the contacts at the interface. Moreover, we showed that the alternative interfaces predicted in our study are biologically relevant and their characterization allows to identify key direct and water-mediated contacts at the interface. In fact, our capability to identify alternative interfaces is also crucial to understand and unveil how the allosteric communication within the complex and the bound biomolecules is influenced and to study different biological pathways as well as to determine key residues in the biomolecule and at the interface providing new pieces of information on the complexes under investigation. In this context, it is important to point out that the interface clustering method presented here cannot only be applied to standard MD simulations, but it can also be extended to characterize different repeats or enhanced sampled MD simulations, allowing to identify very distant interface substates and determine their pathways. Our analysis conducted on the number of interfacial water and the water mediated-contacts also suggests that the packing between the protein and the RNA molecule at the interface may have a crucial role on the water at the interface and its dynamics. These results seem to indicate that in protein-RNA docking the presence of water molecules at the interface may open a new route to predict them. Finally, our results also suggest that our approach could be exploited to design new aptamer or more in general RNA molecules as specific therapeutics.

## Author Contributions

The study was designed by EF. AS, CH and EF performed all-atom MD simulations. AS, CH, JM and EF analyzed all-atom MD simulations. EF and JM supervised AS. EF supervised CH. All authors contributed to write the manuscript.

## Acknowledgement

The authors thank all members of the “Structure et traduction des ARN viraux” team at CiTCoM for useful comments and suggestions all along the process. EF and CH also thank Dr. M.C.R. Melo for his help with the dynamical network analysis.

## Supporting Information Available

The Supporting Information is available free of charge on the ACS Publications website at DOI:

Table of simulation time, average RMSD value, stacking/pairing RNA, interface parameters values and interaction type appearance; RMSD/RMSF/FEL profiles; figures of geometric interface parameters distributions; initial contact time series; principal component analysis projection; interface clustering results; contact variance matrix; network analysis; puckering probability; interface water molecules and water mediated contact.

## Data and Software Availability

The MD simulations obtained for the unbound and bound structures without water are available in the repository Zenodo. Part of the interface analysis is conducted with in-house python scripts that are available at https://github.com/juliettemartin/DYN_INTERFACES.

