## Supplementary Information for "The dynamics of protein-RNA interfaces using all-atom molecular dynamics simulations"

§*Laboratory of Biology and Modeling of the Cell, Université de Lyon, ENS de Lyon, Université Claude Bernard, CNRS UMR 5239, Inserm, U1293, Lyon, France.*

### Contents


**S1** **Tables** **S4**

**S2** **RMSD profiles** **S13**

**S3** **RMSF profiles** **S22**

**S4** **Free Energy Landscape** **S25**

**S5** **Burried Assessible Surface Area (ΔASA)** **S34**

**S6** **Gap Volume** **S36**

**S7** **Gap Index** **S38**

**S8** **Initial contacts** **S40**

**S9** **Principal component analysis** **S41**

**S10** **Interface clustering** **S42**

**S11** **Variance of contact frequencies across clusters** **S50**

**S12** **Network** **S53**

**S13** **Puckering** **S65**

**S14** **Puckering clusters** **S70**

**S15** **Interfacial water molecules evolution** **S71**

**S16** **Interfacial water molecules number** **S72**

**S17** **Contact type** **S80**

**S18 Contact water mediated** **S81**

### S1 Tables

Table S1: Simulation times of the nine protein-RNA complexes. b-$_b$: bound; $_u$: unbound.

| PDB ID$_b$ | Protein/RNA$_b$ (ns) | Protein$_u$ (ns) | RNA$_u$ (ns) |
|---|---|---|---|
| 1OOA | 1500 | 1000 | 1000 |
| 1ASY | 1000 | 1000 | 1000 |
| 2ZM5 | 1000 | 500 | 1000 |
| 1MMS | 1000 | 1000 | 1000 |
| 3IEV | 500 | 500 | 1000 |
| 2R8S | 500 | 500 | 500 |
| 2VPL | 1000 | 500 | 1000 |
| 1JBS | 1000 | 500 | 750 |
| 1RKJ | 500 | 500 | 500 |

Table S2: Mean RMSD value of the nine protein-RNA complexes with standard deviation inside brackets and with the starting structure or the main cluster structure as reference. Ref.: Reference structure; $_b$: bound; $_u$: unbound; $_s$: structure.

| PDB ID$_b$ | Ref. | Protein/RNA$_b$ (Å) | Protein$_b$ (Å) | RNA$_b$ (Å) | Protein$_u$ (Å) | RNA$_u$ (Å) |
|---|---|---|---|---|---|---|
| 1ASY | Starting$_s$ | 6.2 (1.1) | 6.3 (1.1) | 4.0 (0.9) | 4.5 (0.8) | 3.3 (0.4) |
| | Cluster$_s$ | 3.9 (1.3) | 3.7 (1.2) | 2.6 (1.0) | 3.5 (0.8) | 2.3 (0.6) |
| 1JBS | Starting$_s$ | 2.7 (0.8) | 2.0 (0.2) | 1.9 (0.5) | 2.7 (0.3) | 4.8 (0.9) |
| | Cluster$_s$ | 2.5 (0.9) | 1.7 (0.2) | 1.7 (0.4) | 2.0 (0.3) | 3.4 (1.3) |
| 1MMS | Starting$_s$ | 2.1 (0.3) | 2.0 (0.3) | 1.8 (0.2) | 6.6 (0.7) | 2.7 (0.2) |
| | Cluster$_s$ | 1.8 (0.3) | 1.8 (0.4) | 1.4 (0.2) | 2.5 (0.8) | 1.5 (0.3) |
| 1OOA | Starting$_s$ | 9.6 (1.4) | 9.4 (1.2) | 4.2 (1.6) | 7.0 (1.1) | 3.3 (0.9) |
| | Cluster$_s$ | 4.5 (1.5) | 2.9 (1.5) | 4.8 (1.3) | 3.3 (1.1) | 3.2 (1.0) |
| 1RKJ | Starting$_s$ | 12.0 (1.6) | 11.0 (1.4) | 5.6 (0.6) | 9.1 (0.9) | 2.0 (0.5) |
| | Cluster$_s$ | 5.6 (2.1) | 4.9 (2.0) | 2.7 (0.6) | 6.0 (2.6) | 2.1 (0.5) |
| 2R8S | Starting$_s$ | 4.0 (0.6) | 2.2 (0.2) | 4.3 (0.6) | 2.6 (0.3) | 4.0 (0.6) |
| | Cluster$_s$ | 3.0 (0.6) | 2.0 (0.3) | 3.0 (0.8) | 2.1 (0.3) | 2.4 (0.5) |
| 2VPL | Starting$_s$ | 2.8 (0.4) | 2.0 (0.2) | 3.1 (0.5) | 2.9 (0.3) | 9.9 (3.6) |
| | Cluster$_s$ | 2.6 (0.4) | 1.9 (0.2) | 2.8 (0.6) | 2.5 (0.6) | 6.2 (3.4) |
| 2ZM5 | Starting$_s$ | 2.7 (0.4) | 2.1 (0.2) | 2.7 (0.3) | 6.5 (1.5) | 3.7 (0.7) |
| | Cluster$_s$ | 2.3 (0.5) | 1.7 (0.2) | 2.0 (0.5) | 4.2 (1.7) | 2.7 (0.8) |
| 3IEV | Starting$_s$ | 2.1 (0.2) | 1.8 (0.1) | 2.5 (0.3) | 1.9 (0.1) | 7.5 (1.5) |
| | Cluster$_s$ | 2.1 (0.2) | 1.7 (0.1) | 3.0 (0.7) | 1.7 (0.1) | 4.4 (2.7) |

Table S3: Average of the different stacking types for the bound or unbound RNA of the nine protein-RNA complexes during the MD trajectory, with standard deviation inside brackets. $_b$: bound; $_u$: unbound.

| PDB ID | States | Upward | Inward | Outward | Downward |
|---|---|---|---|---|---|
| 1ASY | $RNA_b$ | 40.3 (2.1) | 3.6 (0.6) | 10.5 (1.3) | 0 (0) |
| | $RNA_u$ | 41.2 (2.6) | 3.6 (1.1) | 9.4 (1.4) | 0.1 (0.2) |
| 1JBS | $RNA_b$ | 18.0 (1.4) | 2.0 (0.1) | 3.9 (0.7) | 0.1 (0.3) |
| | $RNA_u$ | 15.3 (1.7) | 0.3 (0.5) | 2.6 (0.9) | 0 (0.1) |
| 1MMS | $RNA_b$ | 36.1 (1.9) | 1.5 (0.6) | 5.6 (1.1) | 2.0 (0.1) |
| | $RNA_u$ | 35.8 (1.9) | 1.8 (0.7) | 5.3 (0.9) | 2.6 (0.5) |
| 1OOA | $RNA_b$ | 16.1 (2.1) | 1.0 (0.2) | 1.3 (0.7) | 0 (0.2) |
| | $RNA_u$ | 16.7 (1.6) | 0.9 (0.3) | 1.7 (0.9) | 0.6 (0.5) |
| 1RKJ | $RNA_b$ | 11.2 (1.2) | 0.2 (0.4) | 1.1 (0.4) | 0 (0) |
| | $RNA_u$ | 14.0 (1.3) | 0 (0) | 1.4 (0.6) | 0 (0) |
| 2R8S | $RNA_b$ | 101.3 (3.0) | 5.8 (0.6) | 18.0 (1.6) | 2.9 (0.2) |
| | $RNA_u$ | 100.5 (3.1) | 5.3 (0.8) | 16.2 (1.6) | 3.1 (0.4) |
| 2VPL | $RNA_b$ | 32.3 (2.1) | 1.0 (0.3) | 5.3 (1.0) | 0 (0.2) |
| | $RNA_u$ | 31.9 (2.4) | 0.5 (0.6) | 4.2 (1.2) | 0.1 (0.3) |
| 2ZM5 | $RNA_b$ | 42.1 (2.0) | 3.4 (0.7) | 9.3 (1.3) | 0 (0) |
| | $RNA_u$ | 44.4 (2.2) | 3.1 (0.8) | 8.1 (1.3) | 0 (0) |
| 3IEV | $RNA_b$ | 2.7 (0.6) | 0.5 (0.5) | 0 (0) | 0 (0) |
| | $RNA_u$ | 3.3 (1.9) | 0.8 (0.7) | 0.2 (0.4) | 0.8 (0.8) |

Table S4: Average of the different pairing types for the bound or unbound RNA of the nine protein-RNA complexes during the MD trajectory, with standard deviation inside brackets. t: trans; c: cis; W: Watson–Crick side; H: Hoogsten side; S: sugar side; B: B side; $_b$: bound; $_u$: unbound. (1)

| Type | States | 1ASY | 1JBS | 1MMS | 1OOA | 1RKJ | 2R8S | 2VPL | 2ZM5 | 3IEV |
|---|---|---|---|---|---|---|---|---|---|---|
| tWW | $RNA_b$ | 2.0 (0.4) | 0 (0) | 1.1 (0.6) | 0 (0.2) | 0 (0) | 1.3 (0.5) | 0 (0.2) | 2.1 (0.4) | 0 (0) |
| | $RNA_u$ | 0.6 (0.6) | 0 (0.1) | 0 (0.2) | 0 (0.1) | 0.9 (0.3) | 1.3 (0.6) | 0 (0.2) | 1.1 (0.5) | 0.4 (0.6) |
| cWW | $RNA_b$ | 21.8 (1.1) | 7.3 (0.6) | 16.5 (0.7) | 5.3 (1.9) | 7.0 (0.1) | 47.8 (0.9) | 17.4 (0.6) | 22.8 (0.8) | 0 (0) |
| | $RNA_u$ | 20.9 (1.0) | 7.2 (0.5) | 16.6 (0.7) | 5.9 (1.3) | 7.9 (0.5) | 47.6 (1.2) | 17.2 (1.0) | 21.7 (0.7) | 0.1 (0.2) |
| tWH | $RNA_b$ | 1.7 (0.5) | 1.0 (0.2) | 0.2 (0.4) | 0 (0.2) | 0 (0.1) | 3.4 (0.6) | 0 (0.1) | 3.0 (0.3) | 0 (0) |
| | $RNA_u$ | 2.2 (0.5) | 0.4 (0.5) | 3.0 (1.0) | 0.1 (0.2) | 0 (0.1) | 4.1 (0.7) | 0 (0.2) | 2.5 (0.6) | 0.1 (0.3) |
| cWH | $RNA_b$ | 0.1 (0.4) | 0 (0.1) | 2.4 (0.6) | 0.1 (0.3) | 0 (0.1) | 1.6 (0.6) | 0 (0.1) | 0.4 (0.6) | 0 (0) |
| | $RNA_u$ | 0.3 (0.5) | 0 (0.1) | 3.2 (0.7) | 0.1 (0.3) | 0 (0.2) | 1.1 (0.3) | 0 (0.1) | 1.0 (0.5) | 0 (0.2) |
| tWB | $RNA_b$ | 1.2 (0.7) | 0 (0.1) | 0 (0.1) | 0 (0) | 0 (0) | 0.1 (0.2) | 0 (0) | 0.9 (0.3) | 0 (0) |
| | $RNA_u$ | 0.3 (0.5) | 0 (0) | 0 (0.2) | 0 (0) | 0 (0.1) | 0.1 (0.4) | 0 (0) | 0.8 (0.4) | 0 (0.1) |
| cWB | $RNA_b$ | 0.7 (0.8) | 0 (0.1) | 0.2 (0.4) | 0.5 (0.6) | 0 (0.1) | 0.4 (0.6) | 0 (0.1) | 0 (0.2) | 0 (0) |
| | $RNA_u$ | 0.1 (0.3) | 0 (0.2) | 0.2 (0.5) | 0.1 (0.2) | 0 (0.1) | 0.8 (0.7) | 0.1 (0.2) | 0.1 (0.3) | 0 (0) |
| tWS | $RNA_b$ | 0 (0.1) | 0 (0.1) | 0 (0.1) | 0 (0) | 0 (0) | 1.3 (0.7) | 0 (0.1) | 0 (0.1) | 0 (0) |
| | $RNA_u$ | 0.2 (0.4) | 0.1 (0.3) | 0 (0.1) | 0 (0.1) | 0 (0) | 1.1 (0.7) | 0 (0.1) | 0.2 (0.4) | 0.1 (0.3) |
| cWS | $RNA_b$ | 0 (0.2) | 0 (0.1) | 0 (0) | 0.9 (1.1) | 0 (0) | 0.7 (0.6) | 0 (0.2) | 0.7 (0.5) | 0 (0) |
| | $RNA_u$ | 0.2 (0.5) | 0.3 (0.6) | 0 (0.1) | 0 (0.1) | 0 (0) | 0.7 (0.7) | 0.1 (0.4) | 0.1 (0.2) | 0 (0.1) |
| tHH | $RNA_b$ | 0.2 (0.4) | 1.0 (0.1) | 0 (0) | 0 (0) | 0 (0) | 0 (0.1) | 0 (0) | 0.8 (0.4) | 0 (0) |
| | $RNA_u$ | 0.2 (0.4) | 0 (0.1) | 0 (0) | 0 (0) | 0 (0) | 0 (0) | 0.1 (0.3) | 0.1 (0.3) | 0 (0) |
| cHH | $RNA_b$ | 0.1 (0.2) | 0 (0.1) | 0 (0) | 0 (0) | 0 (0) | 0 (0) | 0 (0) | 0 (0.1) | 0 (0) |
| | $RNA_u$ | 0.1 (0.3) | 0 (0) | 0 (0) | 0 (0) | 0 (0) | 0 (0.1) | 0 (0.1) | 0 (0) | 0 (0) |
| tHB | $RNA_b$ | 0 (0.2) | 0 (0.2) | 0 (0.1) | 0.1 (0.3) | 0 (0) | 0.1 (0.3) | 0 (0.1) | 0 (0.1) | 0 (0) |
| | $RNA_u$ | 0 (0.2) | 0 (0.2) | 0 (0) | 0.2 (0.4) | 0 (0) | 0.1 (0.3) | 0 (0.1) | 0.2 (0.4) | 0 (0) |
| cHB | $RNA_b$ | 0.1 (0.3) | 0 (0.1) | 0 (0.1) | 0 (0) | 0 (0) | 0 (0.1) | 0 (0.1) | 0.1 (0.3) | 0 (0.1) |
| | $RNA_u$ | 0.4 (0.5) | 0 (0.1) | 0 (0.1) | 0 (0.1) | 0 (0.1) | 0 (0.1) | 0 (0) | 0 (0) | 0 (0) |
| tHS | $RNA_b$ | 0 (0.1) | 1.8 (0.4) | 1.7 (0.5) | 1.8 (0.8) | 0 (0) | 6.6 (0.9) | 1.2 (0.5) | 0.2 (0.4) | 0 (0) |
| | $RNA_u$ | 1.2 (0.6) | 1.3 (0.5) | 1.7 (0.5) | 2.1 (1.0) | 0 (0.2) | 4.9 (0.8) | 1.1 (0.5) | 0.1 (0.4) | 0 (0.1) |
| cHS | $RNA_b$ | 0.4 (0.6) | 0.9 (0.3) | 0.9 (0.2) | 0.2 (0.4) | 0 (0) | 0.9 (0.6) | 0.2 (0.4) | 0.1 (0.3) | 0.1 (0.3) |
| | $RNA_u$ | 0.7 (0.6) | 0.8 (0.4) | 0.9 (0.3) | 0.4 (0.6) | 0.2 (0.4) | 1.7 (0.9) | 0.2 (0.4) | 0.1 (0.3) | 0 (0) |
| tBB | $RNA_b$ | 0 (0) | 0 (0) | 0 (0) | 0 (0) | 0 (0) | 0 (0) | 0 (0) | 0 (0) | 0 (0) |
| | $RNA_u$ | 0 (0) | 0 (0) | 0 (0) | 0 (0) | 0 (0) | 0 (0) | 0 (0) | 0 (0) | 0 (0) |
| cBB | $RNA_b$ | 0 (0) | 0 (0) | 0 (0) | 0 (0) | 0 (0) | 0 (0) | 0 (0) | 0 (0) | 0 (0) |
| | $RNA_u$ | 0 (0) | 0 (0) | 0 (0) | 0 (0) | 0 (0) | 0 (0) | 0 (0) | 0 (0) | 0 (0) |
| tBS | $RNA_b$ | 0 (0) | 0 (0.1) | 0 (0.1) | 0 (0.1) | 0 (0) | 0.3 (0.5) | 0.3 (0.5) | 0 (0.2) | 0 (0) |
| | $RNA_u$ | 0 (0.2) | 0 (0) | 0 (0.2) | 0.1 (0.3) | 0 (0.1) | 0.1 (0.3) | 0 (0.2) | 0.1 (0.3) | 0 (0) |
| cBS | $RNA_b$ | 0 (0) | 0 (0.1) | 0 (0.2) | 0 (0.1) | 0 (0) | 0.1 (0.3) | 0 (0) | 0 (0) | 0 (0) |
| | $RNA_u$ | 0 (0) | 0 (0.1) | 0 (0.1) | 0 (0) | 0 (0) | 0.2 (0.5) | 0 (0) | 0 (0) | 0 (0) |
| tSS | $RNA_b$ | 0 (0) | 0 (0) | 1.0 (0.5) | 0 (0) | 0 (0) | 2.1 (0.7) | 1.0 (0.5) | 0 (0) | 0 (0) |
| | $RNA_u$ | 0 (0) | 0 (0) | 0.1 (0.3) | 0 (0) | 0 (0) | 2.0 (0.8) | 0 (0.2) | 0 (0) | 0 (0) |
| cSS | $RNA_b$ | 0 (0) | 0 (0) | 0 (0) | 0 (0) | 0 (0.2) | 0 (0) | 0 (0) | 0 (0) | 0 (0) |
| | $RNA_u$ | 0 (0) | 0 (0) | 0 (0.1) | 0 (0) | 0 (0) | 0.3 (0.5) | 0 (0) | 0 (0) | 0 (0) |

Table S5: Average of the different pairing types for the bound or unbound RNA of the nine protein-RNA complexes during the MD trajectory, with standard deviation inside brackets. t: trans; c: cis; W: Watson–Crick side; H: Hoogsten side; S: sugar side; B: B side; O2': O2' atom; C8: C8 atom; OP2: oxygen of the phosphate group; $_b$: bound; $_u$: unbound. (2)

| Type | States | 1ASY | 1JBS | 1MMS | 1OOA | 1RKJ | 2R8S | 2VPL | 2ZM5 | 3IEV |
|---|---|---|---|---|---|---|---|---|---|---|
| O2'W | $RNA_b$ | 0.8 (0.8) | 0.2 (0.4) | 2.9 (0.6) | 0.2 (0.4) | 0 (0.2) | 2.6 (1.3) | 0.9 (0.7) | 1.8 (0.9) | 0 (0) |
| | $RNA_u$ | 2.2 (1.1) | 0.3 (0.5) | 0.7 (0.6) | 0.9 (0.7) | 0.1 (0.2) | 3.1 (1.3) | 0.3 (0.5) | 2.0 (1.0) | 0.3 (0.5) |
| O2'H | $RNA_b$ | 1.8 (0.9) | 0 (0.1) | 3.3 (1.1) | 0.6 (0.6) | 0.1 (0.3) | 2.1 (1.1) | 0.3 (0.5) | 1.9 (0.6) | 0 (0.1) |
| | $RNA_u$ | 1.0 (0.7) | 0.5 (0.7) | 1.3 (1.0) | 0.6 (0.5) | 0.1 (0.4) | 2.0 (1.1) | 0.3 (0.6) | 2.8 (0.8) | 0.2 (0.4) |
| O2'B | $RNA_b$ | 1.3 (1.0) | 0.2 (0.4) | 2.5 (1.0) | 0.8 (1.0) | 0.2 (0.5) | 4.3 (1.5) | 0.2 (0.4) | 2.1 (0.8) | 0 (0) |
| | $RNA_u$ | 1.0 (0.8) | 0.7 (0.9) | 1.0 (0.9) | 1.1 (1.0) | 0 (0.1) | 3.8 (1.6) | 0.7 (0.9) | 2.4 (1.1) | 0.1 (0.2) |
| O2'S | $RNA_b$ | 0.4 (0.5) | 0 (0) | 1.1 (0.7) | 0.1 (0.3) | 0 (0.1) | 3.5 (1.0) | 0.1 (0.3) | 0.1 (0.3) | 0 (0) |
| | $RNA_u$ | 0.2 (0.4) | 0.1 (0.3) | 0.6 (0.7) | 0 (0.1) | 0 (0) | 3.2 (1.0) | 0.2 (0.5) | 0 (0.1) | 0.1 (0.3) |
| O2PW | $RNA_b$ | 0.8 (0.6) | 1.3 (0.6) | 3.1 (0.9) | 0 (0.1) | 0 (0.1) | 3.1 (1.1) | 0.6 (0.5) | 1.1 (0.4) | 0 (0) |
| | $RNA_u$ | 0.2 (0.5) | 1.4 (0.7) | 1.2 (0.8) | 0.6 (0.6) | 0 (0) | 1.3 (0.7) | 0.3 (0.5) | 3.2 (0.9) | 0 (0.2) |
| O2PH | $RNA_b$ | 0 (0.2) | 0 (0) | 0.2 (0.4) | 0 (0.1) | 0 (0) | 0.1 (0.3) | 0.1 (0.3) | 0 (0.1) | 0 (0) |
| | $RNA_u$ | 0 (0.1) | 0.2 (0.4) | 0 (0.2) | 0.1 (0.3) | 0 (0) | 0.6 (0.5) | 0 (0.1) | 0.2 (0.4) | 0 (0) |
| O2PB | $RNA_b$ | 2.8 (0.9) | 0.3 (0.5) | 1.7 (1.0) | 0.8 (0.8) | 1.1 (0.8) | 4.3 (1.4) | 0.9 (0.9) | 2.6 (0.7) | 0 (0) |
| | $RNA_u$ | 2.5 (1.0) | 1.7 (1.0) | 1.8 (1.2) | 0.2 (0.4) | 1.0 (0.4) | 4.0 (1.3) | 0.8 (0.8) | 1.4 (1.0) | 0.5 (0.8) |
| O2PS | $RNA_b$ | 0 (0) | 0 (0) | 0 (0.1) | 0 (0) | 0 (0) | 0 (0.1) | 0 (0) | 0 (0) | 0 (0) |
| | $RNA_u$ | 0 (0) | 0 (0.1) | 0 (0) | 0 (0) | 0 (0) | 0 (0.1) | 0 (0) | 0 (0) | 0 (0) |
| tWC8 | $RNA_b$ | 0.1 (0.3) | 0 (0) | 0 (0) | 0 (0) | 0 (0) | 0 (0) | 0 (0) | 0 (0) | 0 (0) |
| | $RNA_u$ | 0 (0.1) | 0 (0) | 0 (0) | 0 (0) | 0 (0) | 0 (0) | 0 (0) | 0 (0) | 0 (0) |
| cWC8 | $RNA_b$ | 0 (0) | 0 (0) | 0 (0) | 0 (0) | 0 (0) | 0 (0) | 0 (0) | 0 (0) | 0 (0) |
| | $RNA_u$ | 0 (0) | 0 (0) | 0 (0) | 0 (0) | 0 (0) | 0 (0) | 0 (0) | 0 (0) | 0 (0) |
| tHC8 | $RNA_b$ | 0 (0.1) | 0 (0) | 0 (0) | 0 (0) | 0 (0) | 0 (0) | 0 (0) | 0 (0) | 0 (0) |
| | $RNA_u$ | 0 (0.1) | 0 (0) | 0 (0) | 0 (0) | 0 (0) | 0 (0) | 0 (0) | 0 (0) | 0 (0) |
| cHC8 | $RNA_b$ | 0.1 (0.2) | 0 (0) | 0 (0) | 0 (0) | 0 (0) | 0 (0) | 0 (0) | 0 (0) | 0 (0) |
| | $RNA_u$ | 0.1 (0.2) | 0 (0) | 0 (0) | 0 (0) | 0 (0) | 0 (0) | 0 (0) | 0 (0) | 0 (0) |
| tBC8 | $RNA_b$ | 0 (0) | 0 (0) | 0 (0) | 0 (0) | 0 (0) | 0 (0) | 0 (0) | 0 (0) | 0 (0) |
| | $RNA_u$ | 0 (0) | 0 (0) | 0 (0) | 0 (0) | 0 (0) | 0 (0) | 0 (0) | 0 (0) | 0 (0) |
| cBC8 | $RNA_b$ | 0 (0) | 0 (0) | 0 (0) | 0 (0) | 0 (0) | 0 (0) | 0 (0) | 0 (0) | 0 (0) |
| | $RNA_u$ | 0 (0) | 0 (0) | 0 (0) | 0 (0) | 0 (0) | 0 (0) | 0 (0) | 0 (0) | 0 (0) |
| tSC8 | $RNA_b$ | 0 (0) | 0 (0) | 0 (0.2) | 0 (0) | 0 (0) | 0 (0) | 0 (0) | 0 (0) | 0 (0) |
| | $RNA_u$ | 0 (0) | 0 (0.1) | 0 (0) | 0 (0) | 0 (0) | 0 (0.1) | 0 (0) | 0 (0) | 0 (0) |
| cSC8 | $RNA_b$ | 0 (0) | 0 (0) | 0 (0) | 0 (0) | 0 (0) | 0 (0.1) | 0 (0) | 0 (0) | 0 (0) |
| | $RNA_u$ | 0 (0) | 0 (0.1) | 0 (0) | 0 (0) | 0 (0) | 0 (0.1) | 0 (0) | 0 (0) | 0 (0) |
| O2'C8 | $RNA_b$ | 0 (0.2) | 0 (0.1) | 0.1 (0.3) | 0.1 (0.3) | 0.1 (0.2) | 0.2 (0.4) | 0 (0.2) | 0 (0.1) | 0.1 (0.3) |
| | $RNA_u$ | 0 (0.1) | 0.1 (0.3) | 0.2 (0.4) | 0 (0.2) | 0 (0) | 0.2 (0.5) | 0 (0.3) | 0 (0.2) | 0 (0.2) |
| O2PC8 | $RNA_b$ | 0 (0) | 0 (0) | 0 (0) | 0 (0) | 0 (0) | 0 (0.1) | 0 (0) | 0 (0) | 0 (0) |
| | $RNA_u$ | 0 (0) | 0 (0.2) | 0 (0) | 0 (0.1) | 0 (0) | 0 (0.1) | 0 (0.1) | 0 (0) | 0 (0) |

Table S6: Interface parameters values for the different structures and mean values along MD simulation and for interface cluster of the nine complexes with standard deviation inside brackets. $Structure_e$= experimental structure; $Structure_s$= starting structure; $MD_t$= Molecular dynamics trajectory; $_{nb}$= number

| PDB ID | States | Delta ASA (Å$^2$) | Gap Volume (Å$^3$) | Gap Index (Å) | Interface water$_{nb}$ |
|---|---|---|---|---|---|
| 1ASY | $Structure_e$ | 3584 | 7886 | 4.40 | - |
| | $Structure_s$ | 3182 | 9135 | 5.74 | 240 |
| | $MD_t$ | 3406 (336) | 7855 (691) | 4.67 (0.69) | 205 (21) |
| | Cluster 1 | 3310 (175) | 8455 (590) | 5.13 (0.49) | 217 (17) |
| | Cluster 2 | 2637 (172) | 7445 (545) | 5.68 (0.63) | 168 (18) |
| | Cluster 3 | 3581 (182) | 7632 (564) | 4.28 (0.44) | 205 (15) |
| 1JBS | $Structure_e$ | 1341 | 2579 | 3.84 | - |
| | $Structure_s$ | 1022 | 4113 | 8.05 | 88 |
| | $MD_t$ | 1280 (183) | 3143 (333) | 5.11 (1.53) | 94 (13) |
| | Cluster 1 | 1313 (131) | 3122 (291) | 4.84 (1.01) | 96 (10) |
| | Cluster 2 | 856 (219) | 3414 (600) | 8.49 (2.60) | 74 (19) |
| 1MMS | $Structure_e$ | 2456 | 3148 | 2.56 | - |
| | $Structure_s$ | 2508 | 3901 | 3.11 | 118 |
| | $MD_t$ | 2492 (158) | 3450 (314) | 2.79 (0.36) | 120 (13) |
| 1OOA | $Structure_e$ | 1910 | 5216 | 5.46 | - |
| | $Structure_s$ | 2383 | 6078 | 5.10 | 162 |
| | $MD_t$ | 1627 (292) | 5219 (612) | 6.70 (1.78) | 132 (20) |
| | Cluster 1 | 1938 (210) | 4983 (746) | 5.17 (0.75) | 139 (18) |
| | Cluster 2 | 1769 (210) | 5149 (476) | 5.95 (1.19) | 145 (16) |
| | Cluster 3 | 1398 (229) | 5387 (615) | 7.97 (1.81) | 120 (17) |
| | Cluster 4 | 1650 (147) | 5128 (622) | 6.26 (0.96) | 134 (14) |
| 1RKJ | $Structure_e$ | 2219 | 3846 | 3.47 | - |
| | $Structure_s$ | 2204 | 3770 | 3.42 | 122 |
| | $MD_t$ | 1750 (290) | 2872 (391) | 3.36 (0.70) | 97 (19) |
| | Cluster 1 | 1894 (156) | 3345 (346) | 3.55 (0.38) | 102 (11) |
| | Cluster 2 | 1486 (148) | 3011 (324) | 4.11 (0.72) | 82 (10) |
| | Cluster 3 | 1581 (74) | 2495 (287) | 3.17 (0.43) | 81 (8) |
| | Cluster 4 | 1718 (97) | 2872 (352) | 3.35 (0.40) | 98 (11) |
| | Cluster 5 | 2141 (219) | 2981 (311) | 2.83 (0.55) | 120 (12) |
| 2R8S | $Structure_e$ | 2416 | 5103 | 4.22 | - |
| | $Structure_s$ | 2575 | 5390 | 4.19 | 152 |
| | $MD_t$ | 2536 (124) | 5566 (362) | 4.41 (0.42) | 152 (11) |
| 2VPL | $Structure_e$ | 2388 | 5704 | 4.78 | - |
| | $Structure_s$ | 2486 | 4713 | 3.79 | 124 |
| | $MD_t$ | 2677 (168) | 4637 (522) | 3.49 (0.51) | 131 (16) |
| 2ZM5 | $Structure_e$ | 3937 | 4977 | 2.53 | - |
| | $Structure_s$ | 3800 | 5314 | 2.79 | 174 |
| | $MD_t$ | 3748 (121) | 5401 (374) | 2.89 (0.26) | 190 (11) |
| 3IEV | $Structure_e$ | 2311 | 1180 | 1.02 | - |
| | $Structure_s$ | 2389 | 1347 | 1.12 | 90 |
| | $MD_t$ | 2314 (54) | 1161 (157) | 1.00 (0.14) | 84 (8) |

Table S7: Interface stacking between protein and RNA along MD simulation and for interface cluster of the nine complexes. $Pro_{resID}$: Protein residue ID; $RNA_{resID}$: RNA residue ID; $Freq_{tot}$: Total frequency along MD trajectory; $Cluster_{nb}$: interface cluster number.

| PDB ID | $Pro_{resID}$ | $RNA_{resID}$ | $Freq_{tot}$ (%) | $Cluster_{nb}$ (Freq (%)) |
|---|---|---|---|---|
| 1ASY | Phe127 | G634 | 4.5 | 1 (3.0) |
| | | | | 3 (5.4) |
| | Phe127 | U635 | 99.4 | 1 (99.5) |
| | | | | 2 (99.7) |
| | | | | 3 (99.5) |
| 1JBS | His49 | A17 | 1.7 | 2 (21.5) |
| 1MMS | | | | |
| 1OOA | Tyr56 | A9 | 0.8 | 1 (2.6) |
| | | | | 2 (2.0) |
| | His63 | U14 | 15.5 | 1 (8.8) |
| | | | | 2 (35.8) |
| | | | | 3 (3.0) |
| | | | | 4 (0.9) |
| 1RKJ | His3 | U15 | 10.8 | 4 (0.3) |
| | | | | 5 (42.3) |
| | Phe17 | C11 | 3.8 | 1 (73.7) |
| | | | | 2 (1.0) |
| | Phe56 | C10 | 2.5 | 1 (8.2) |
| | | | | 3 (3.6) |
| | | | | 5 (6.6) |
| | | C11 | 78.9 | 2 (79.9) |
| | | | | 3 (99.6) |
| | | | | 4 (99.4) |
| | | | | 5 (53.6) |
| | Tyr58 | G12 | 0.2 | 1 (4.9) |
| | Tyr140 | U8 | 20.5 | 1 (80.3) |
| | | | | 2 (72.2) |
| | | C9 | 61.4 | 1 (96.7) |
| | | | | 2 (98.6) |
| | | | | 3 (14.1) |
| | | | | 4 (40.1) |
| | | | | 5 (82.7) |
| | Tyr169 | U8 | 20.5 | 2 (0.7) |
| | | | | 3 (60.1) |
| | | | | 4 (95.5) |
| | | | | 5 (93.4) |
| 2R8S | Tyr49 | U135 | 61.3 | |
| | Tyr55 | U135 | 1.0 | |
| | Tyr92 | A179 | 37.5 | |
| | Tyr246 | U185 | 54.3 | |
| | Tyr271 | U185 | 14.2 | |
| | Tyr322 | U135 | 37.5 | |
| 2VPL | His172 | A8 | 34.2 | |
| 2ZM5 | Trp285 | U32 | 99.8 | |
| 3IEV | Tyr276 | U1537 | 98.4 | |
| | | C1538 | 6.0 | |
| | Trp280 | A1534 | 91.0 | |

Table S8: Interface mean total HB between protein and RNA and involving the 2'-OH of the ribose, along MD simulation and for interface cluster of the nine complexes with standard deviation inside brackets. $MD_t$= Molecular dynamics trajectory; $HB_{tot}$: HB total; $HB_{2'-OH}$: HB involving ribose 2'-OH.

| PDB ID | States | $HB_{tot}$ | $HB_{2'-OH}$ |
|---|---|---|---|
| 1ASY | $MD_t$ | 19.6 (3.6) | 0.3 (0.5) |
| | Cluster 1 | 19.3 (2.9) | 0.2 (0.5) |
| | Cluster 2 | 13.2 (2.2) | 0.1 (0.4) |
| | Cluster 3 | 20.8 (3.0) | 0.3 (0.5) |
| 1JBS | $MD_t$ | 6.1 (2.1) | 0.3 (0.5) |
| | Cluster 1 | 6.0 (1.9) | 0.3 (0.5) |
| | Cluster 2 | 3.0 (1.7) | 0.3 (0.5) |
| 1MMS | $MD_t$ | 14.8 (2.3) | 0.7 (0.7) |
| 1OOA | $MD_t$ | 9.0 (2.8) | 0.5 (0.7) |
| | Cluster 1 | 10.9 (3.2) | 0.7 (0.8) |
| | Cluster 2 | 9.1 (2.6) | 0.5 (0.7) |
| | Cluster 3 | 7.8 (2.1) | 0.2 (0.4) |
| | Cluster 4 | 10.4 (2.7) | 1.2 (0.9) |
| 1RKJ | $MD_t$ | 8.0 (2.1) | 0.7 (0.7) |
| | Cluster 1 | 8.2 (1.8) | 1.0 (0.5) |
| | Cluster 2 | 7.6 (1.9) | 1.1 (0.6) |
| | Cluster 3 | 6.8 (1.8) | 0.8 (0.7) |
| | Cluster 4 | 8.4 (1.6) | 0.1 (0.3) |
| | Cluster 5 | 9.1 (2.3) | 0.6 (0.7) |
| 2R8S | $MD_t$ | 12.5 (2.2) | 1.1 (0.8) |
| 2VPL | $MD_t$ | 14.4 (2.4) | 1.8 (0.7) |
| 2ZM5 | $MD_t$ | 27.1 (2.7) | 1.0 (0.8) |
| 3IEV | $MD_t$ | 13.7 (2.0) | 0.3 (0.5) |

Table S9: Percentage of interface HB between protein and RNA involving the 2'-OH of the ribose, along MD simulation and for interface cluster of the nine complexes. $Pro_{resID}$: Protein residue ID; $RNA_{resID}$: RNA residue ID; $Freq_{tot}$: Total frequency along MD trajectory; $Cluster_{nb}$: interface cluster number; $MD_t$= Molecular dynamics trajectory.

| PDB ID | $Pro_{resID}$ | $RNA_{resID}$ | Freq (%) | $MD_t$/$Cluster_{nb}$ |
|---|---|---|---|---|
| 1ASY | Arg556 | U666 | 11.4 | 3 |
| 1JBS | Lys111 | A9 | 19.8 | 2 |
| 1MMS | Lys133 | C1079 | 28.7 | $MD_t$ |
| | Ser90 | C1064 | 13.7 | $MD_t$ |
| | Gln12 | U1061 | 18.5 | $MD_t$ |
| 1OOA | Asn244 | U7 | 27.3 | 1 |
| | Lys145 | A9 | 25.3 | 1 |
| | Lys77 | A16 | 10.0 | 1 |
| | Lys145 | C24 | 19.7 | 2 |
| | Lys241 | G22 | 10.8 | 2 |
| | Lys74 | A17 | 18.3 | 4 |
| | Asn136 | A16 | 32.2 | 4 |
| | Thr143 | U7 | 12.2 | 4 |
| | Lys145 | U6 | 47.0 | 4 |
| 1RKJ | Tyr140 | U8 | 22.6 | $MD_t$ |
| | | | 87.1 | 1 |
| | | | 78.5 | 2 |
| | Lys95 | C11 | 13.1 | 2 |
| | Arg127 | C9 | 26.0 | 3 |
| | Lys95 | C7 | 23.8 | 3 |
| | Arg97 | C17 | 13.5 | 5 |
| | Arg91 | G12 | 10.7 | 5 |
| 2R8S | Thr321 | C170 | 23 | $MD_t$ |
| | Gly57 | G 200 | 60.8 | $MD_t$ |
| | Ser93 | G 180 | 15.7 | $MD_t$ |
| 2VPL | Lys167 | U6 | 16.4 | $MD_t$ |
| | | G7 | 47.6 | $MD_t$ |
| | Thr217 | G10 | 95.7 | $MD_t$ |
| 2ZM5 | Ala125 | G34 | 29.5 | $MD_t$ |
| | Arg270 | C25 | 43.2 | $MD_t$ |
| 3IEV | Arg255 | A1534 | 21.3 | $MD_t$ |

#### S2 RMSD profiles

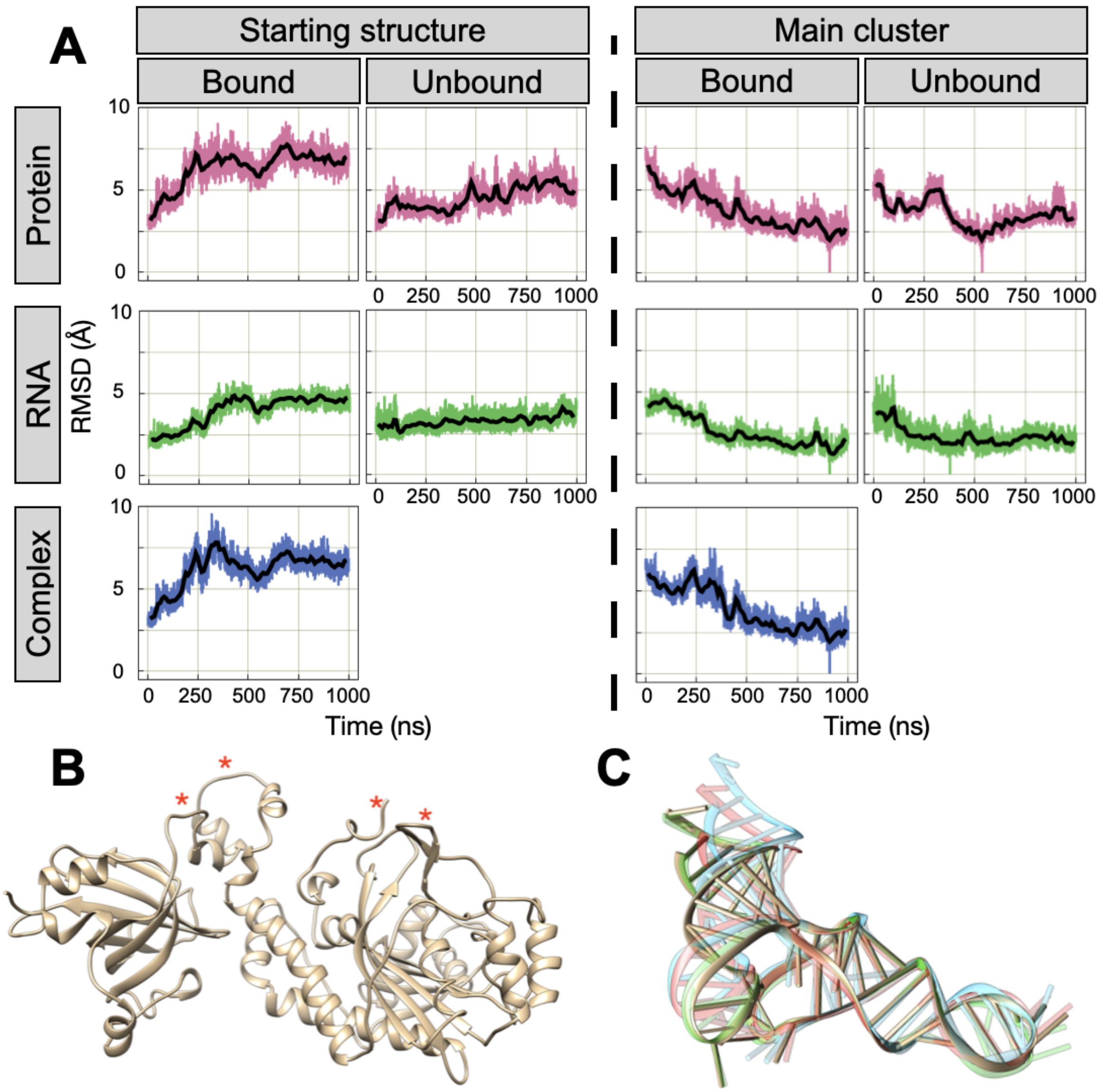


Figure S1: A. RMSD time serie for the bound and unbound simulation of 1ASY complex. RMSDs are computed separately for each chain, with the starting structure (left) or the main cluster structure (right) as reference. Protein is pink, RNA is green and the whole structure is blue. B. Protein structure representation. Red stars highlight the most flexible regions. C. RNA structure representation of the different cluster structure obtained during the simulation. Cyan: Cluster 1; Red: Cluster 2; Green: Cluster 3; Sand: Cluster 4 (main).

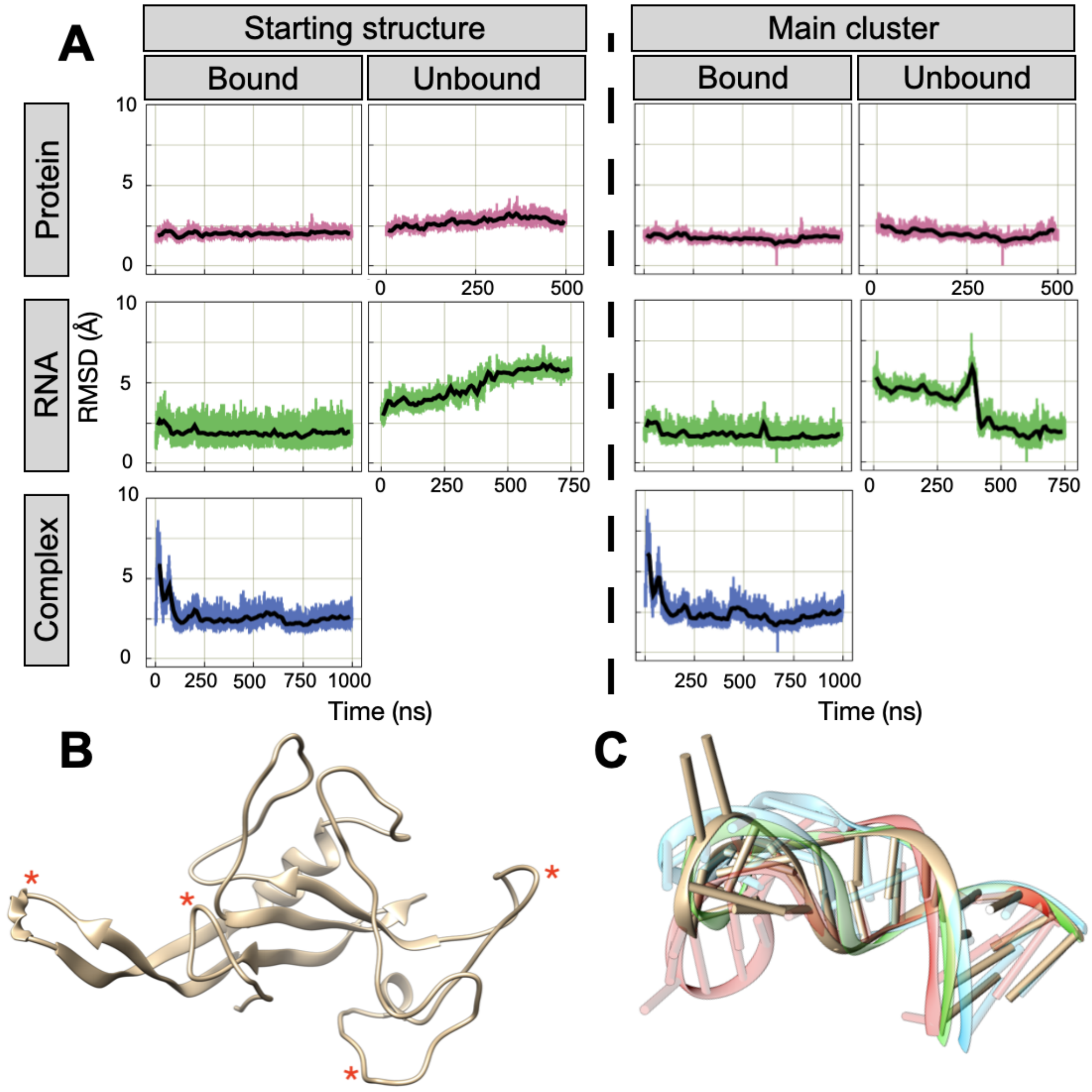


Figure S2: A. RMSD time serie for the bound and unbound simulation of 1JBS complex. RMSDs are computed separately for each chain, with the starting structure (left) or the main cluster structure (right) as reference. Protein is pink, RNA is green and the whole structure is blue. B. Protein structure representation. Red stars highlight the most flexible regions. C. RNA structure representation of the different cluster structure obtained during the simulation. Cyan: Cluster 1; Red: Cluster 2; Green: Cluster 3; Sand: Cluster 4 (main).

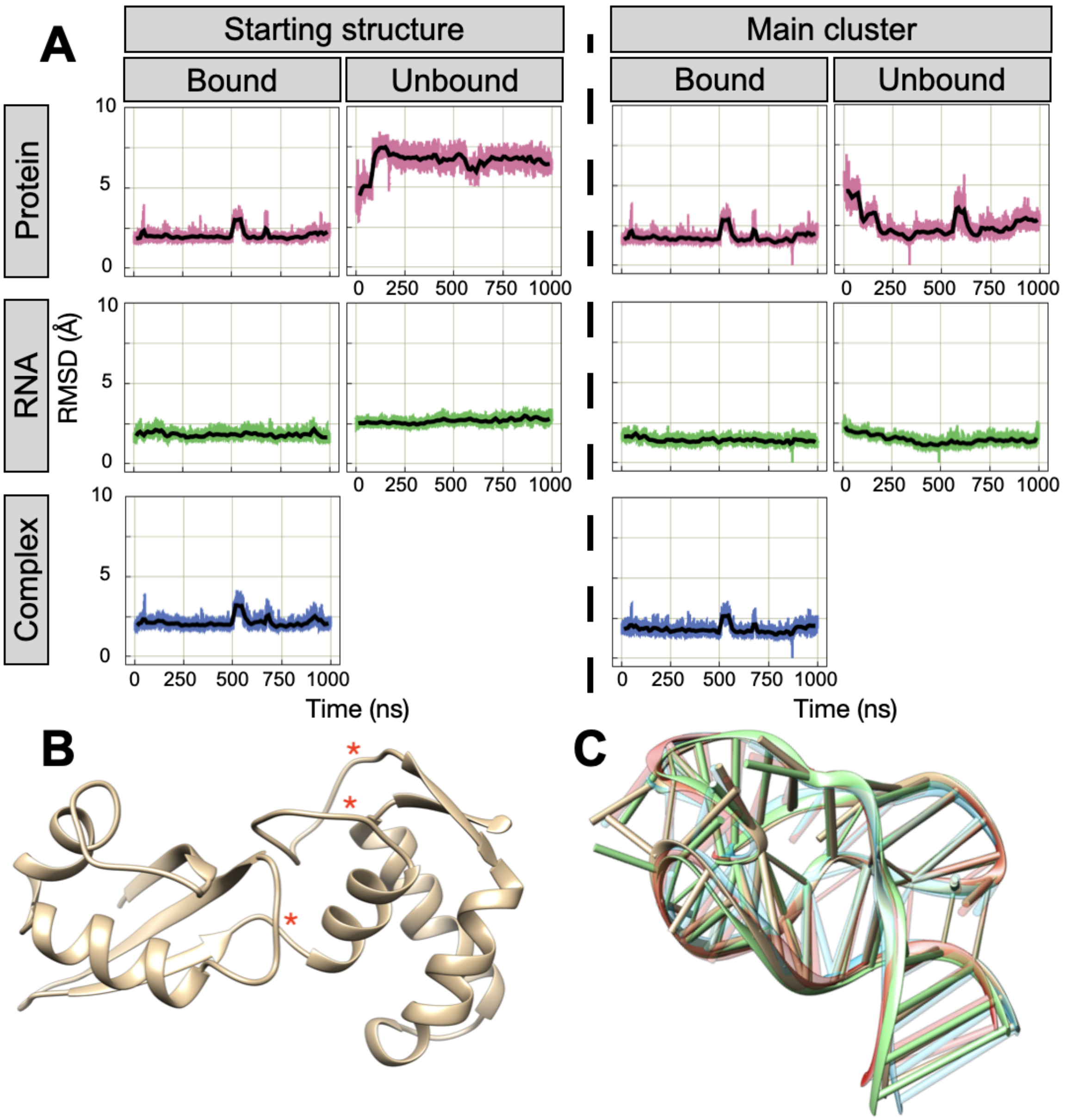


Figure S3: A. RMSD time serie for the bound and unbound simulation of 1MMS complex. RMSDs are computed separately for each chain, with the starting structure (left) or the main cluster structure (right) as reference. Protein is pink, RNA is green and the whole structure is blue. B. Protein structure representation. Red stars highlight the most flexible regions. C. RNA structure representation of the different cluster structure obtained during the simulation. Cyan: Cluster 1; Red: Cluster 2; Sand: Cluster 3 (main); Green: Cluster 4.

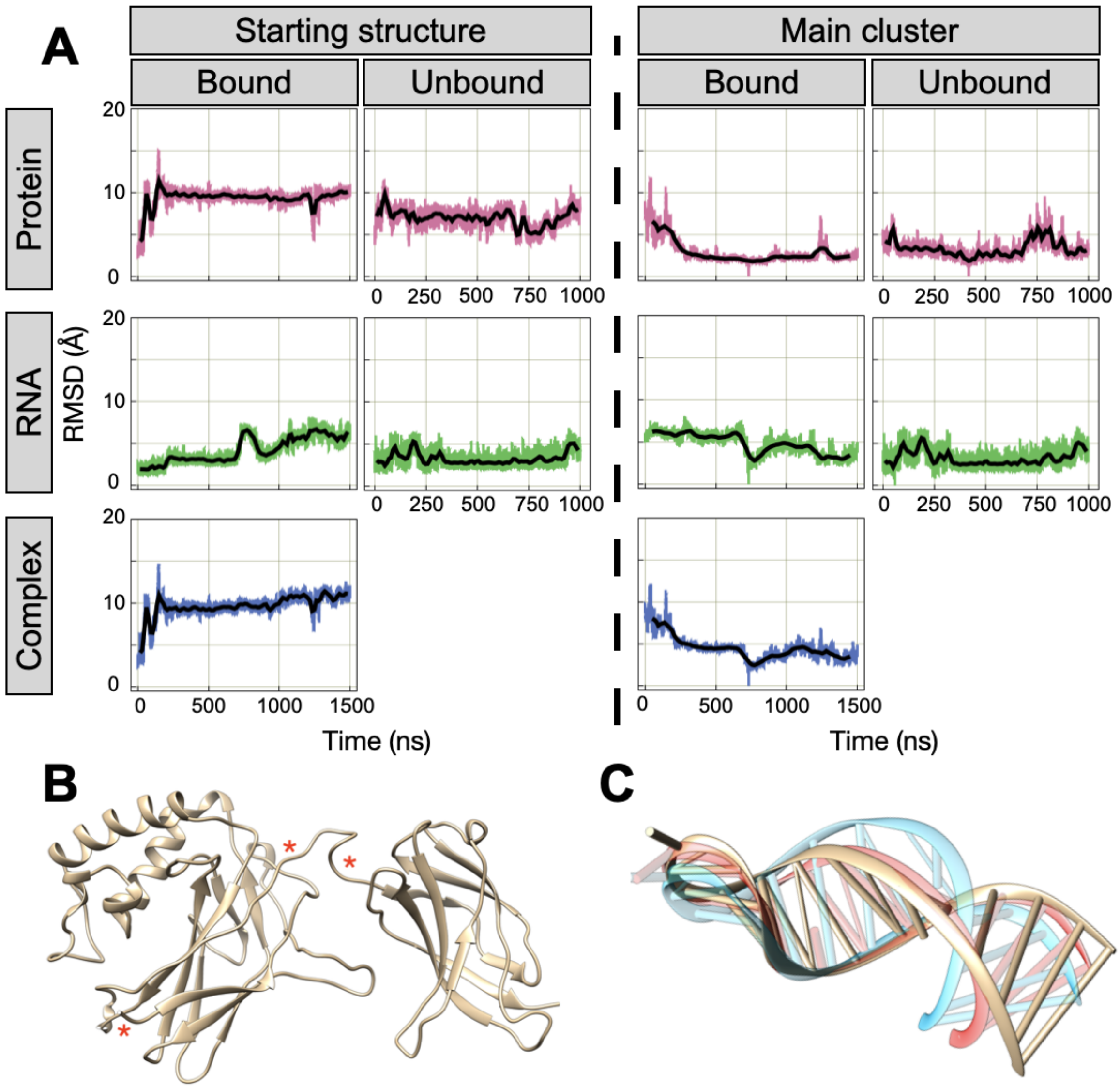


Figure S4: A. RMSD time serie for the bound and unbound simulation of 1OOA complex. RMSDs are computed separately for each chain, with the starting structure (left) or the main cluster structure (right) as reference. Protein is pink, RNA is green and the whole structure is blue. B. Protein structure representation. Red stars highlight the most flexible regions. C. RNA structure representation of the different cluster structure obtained during the simulation. Sand: Cluster 1 (main); Cyan: Cluster 2; Red: Cluster 3.

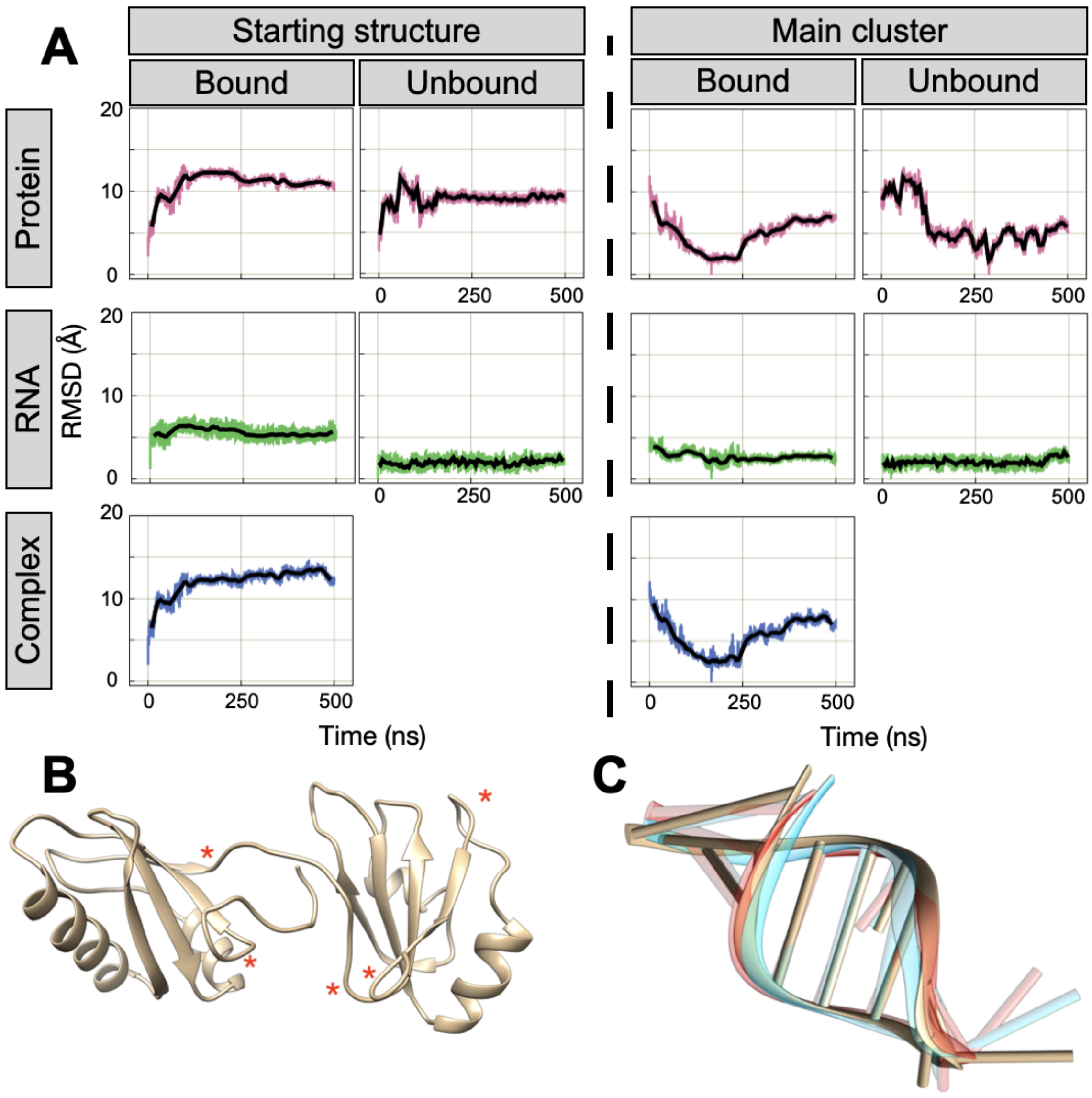


Figure S5: A. RMSD time serie for the bound and unbound simulation of 1RKJ complex. RMSDs are computed separately for each chain, with the starting structure (left) or the main cluster structure (right) as reference. Protein is pink, RNA is green and the whole structure is blue. B. Protein structure representation. Red stars highlight the most flexible regions. C. RNA structure representation of the different cluster structure obtained during the simulation.

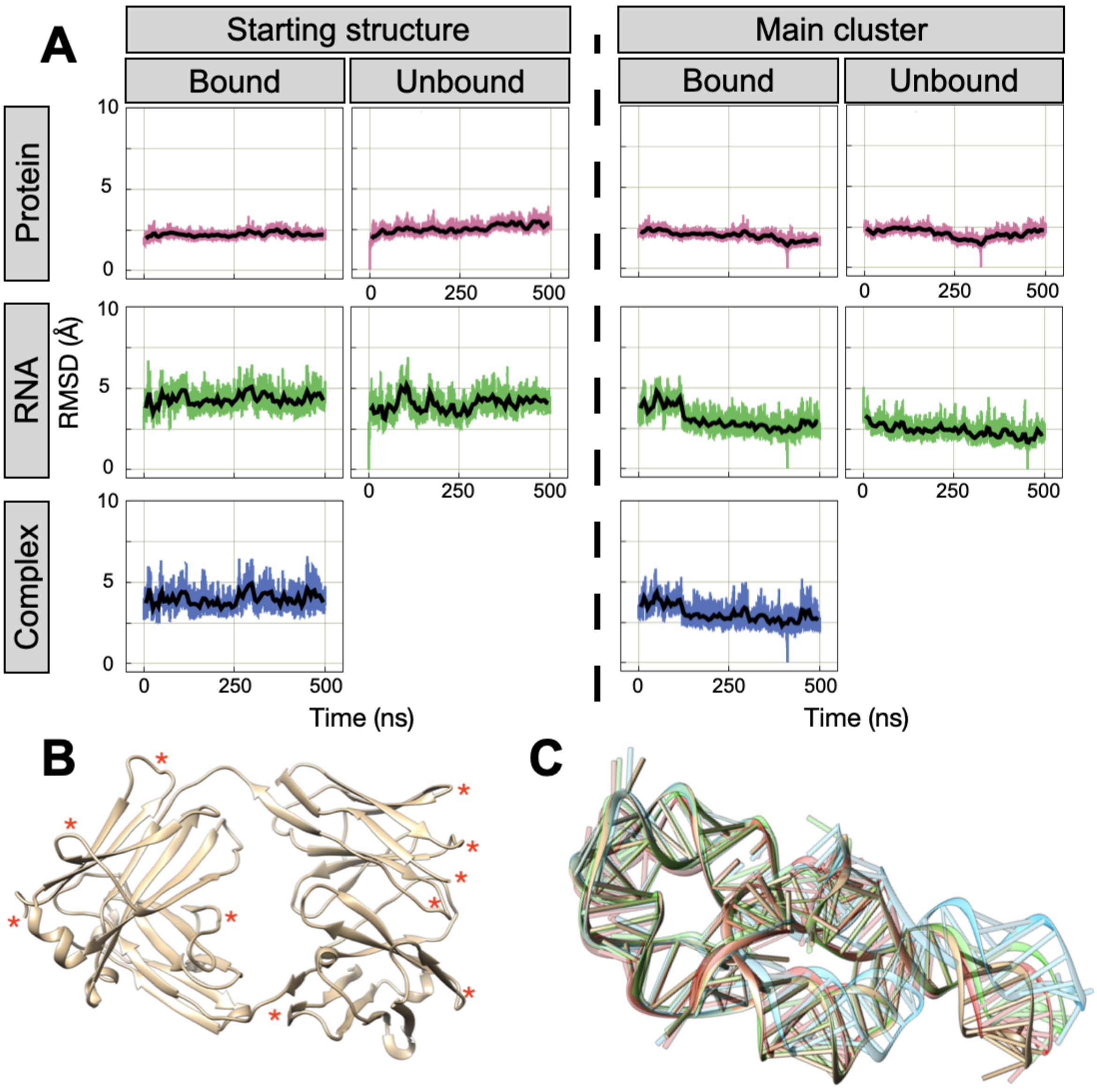


Figure S6: A. RMSD time serie for the bound and unbound simulation of 2R8S complex. RMSDs are computed separately for each chain, with the starting structure (left) or the main cluster structure (right) as reference. Protein is pink, RNA is green and the whole structure is blue. B. Protein structure representation. Red stars highlight the most flexible regions. C. RNA structure representation of the different cluster structure obtained during the simulation. Cyan: Cluster 1; Red: Cluster 2; Sand: Cluster 3 (main); Green: Cluster 4.

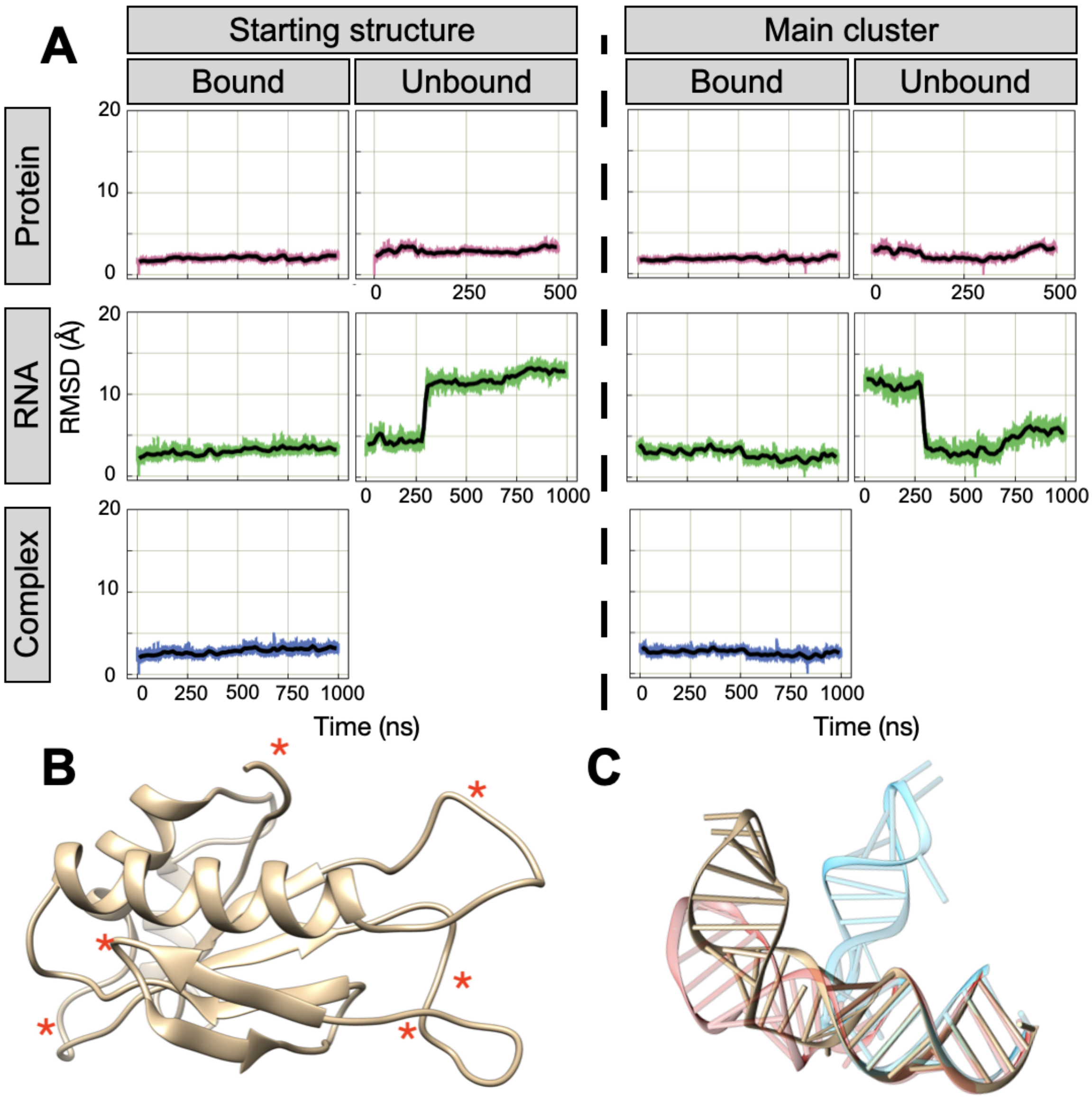


Figure S7: A. RMSD time serie for the bound and unbound simulation of 2VPL complex. RMSDs are computed separately for each chain, with the starting structure (left) or the main cluster structure (right) as reference. Protein is pink, RNA is green and the whole structure is blue. B. Protein structure representation. Red stars highlight the most flexible regions. C. RNA structure representation of the different cluster structure obtained during the simulation. Cyan: Cluster 1; Sand: Cluster 2 (main); Red: Cluster 3.

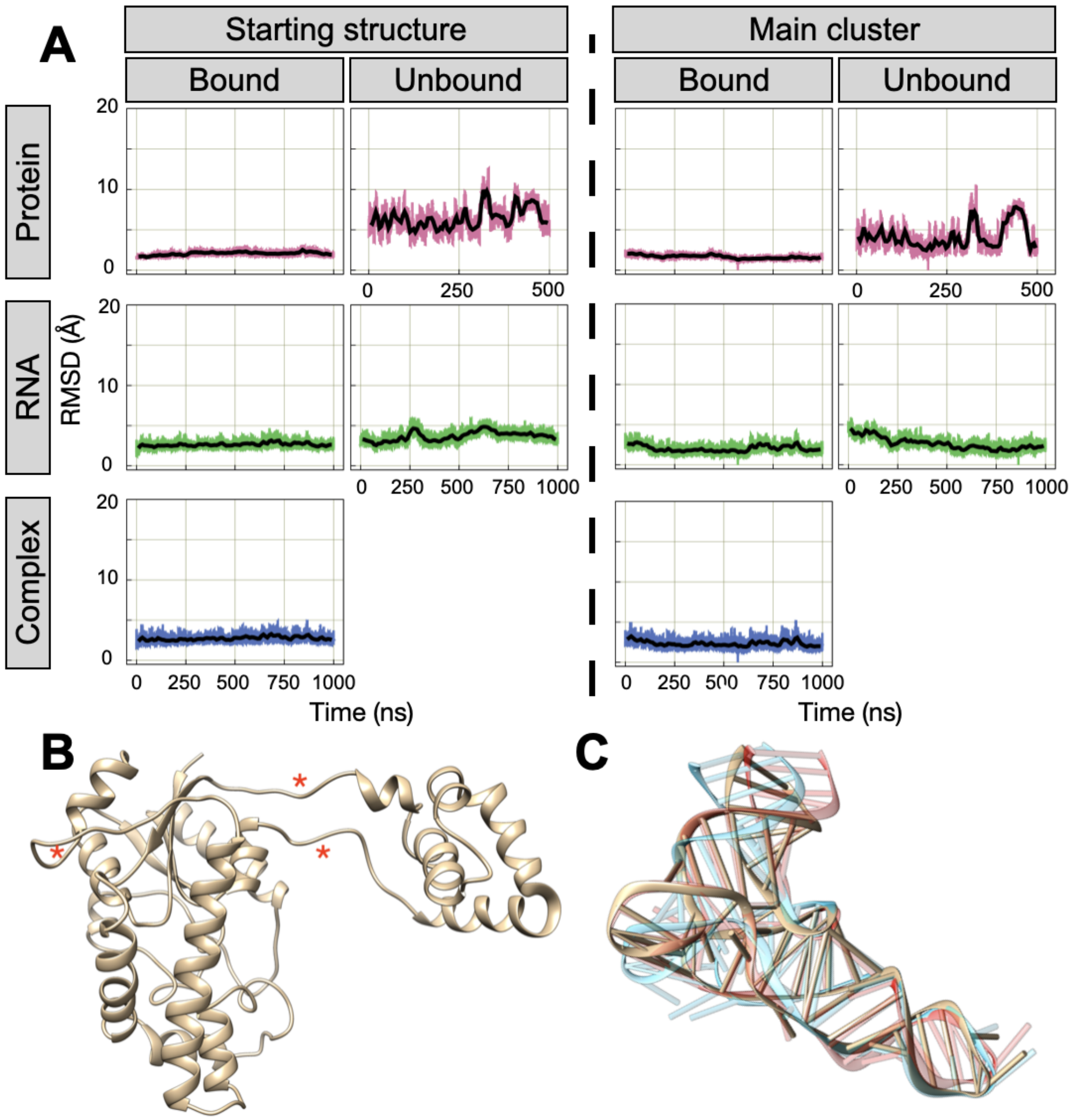


Figure S8: A. RMSD time serie for the bound and unbound simulation of 2ZM5 complex. RMSDs are computed separately for each chain, with the starting structure (left) or the main cluster structure (right) as reference. Protein is pink, RNA is green and the whole structure is blue. B. Protein structure representation. Red stars highlight the most flexible regions. C. RNA structure representation of the different cluster structure obtained during the simulation. Cyan: Cluster 1; Red: Cluster 2; Sand: Cluster 3 (main).

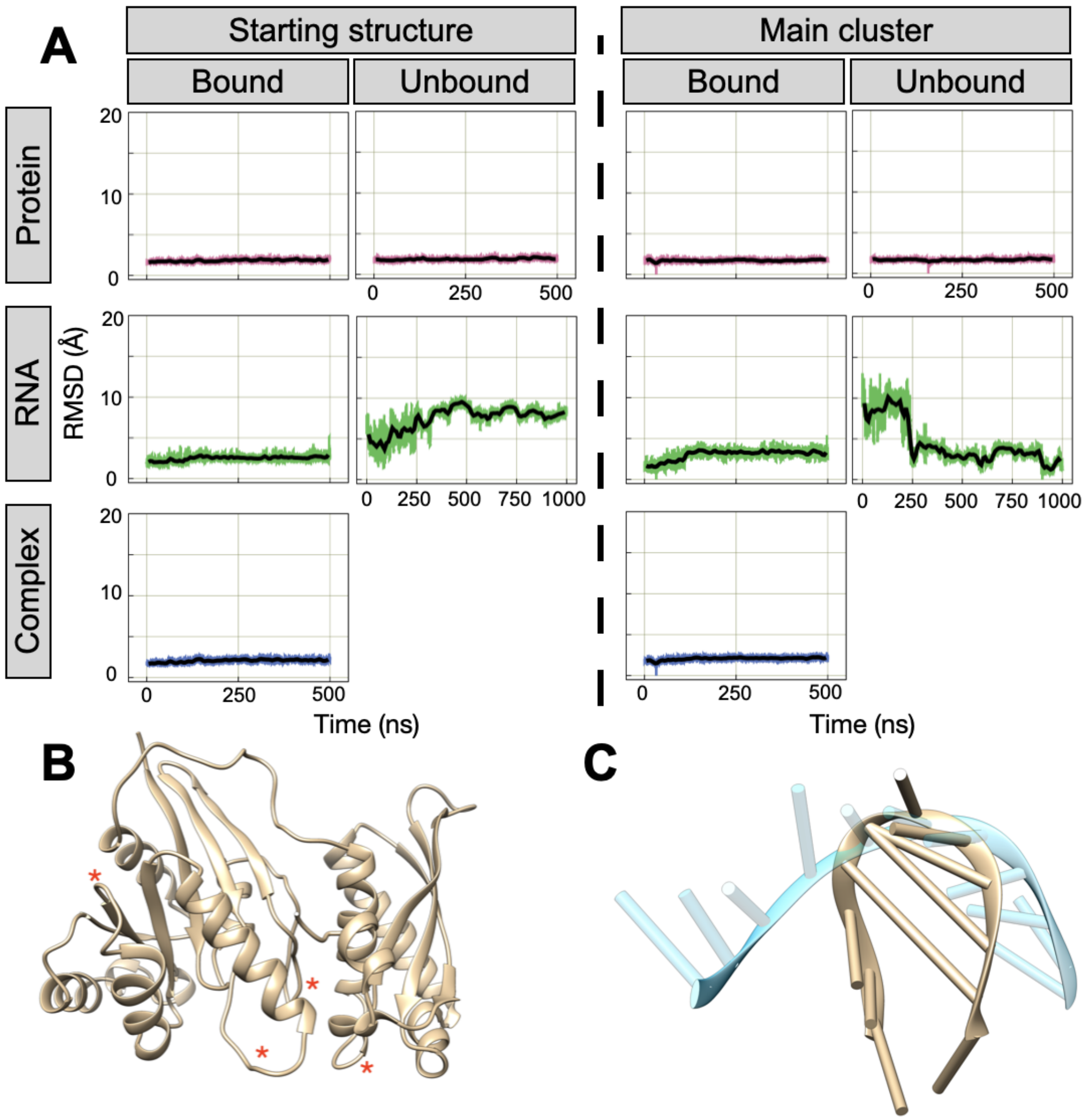


Figure S9: A. RMSD time serie for the bound and unbound simulation of 3IEV complex. RMSDs are computed separately for each chain, with the starting structure (left) or the main cluster structure (right) as reference. Protein is pink, RNA is green and the whole structure is blue. B. Protein structure representation. Red stars highlight the most flexible regions. C. RNA structure representation of the different cluster structure obtained during the simulation. Cyan: Cluster 1; Sand: Cluster 2 (main).

#### S3 RMSF profiles

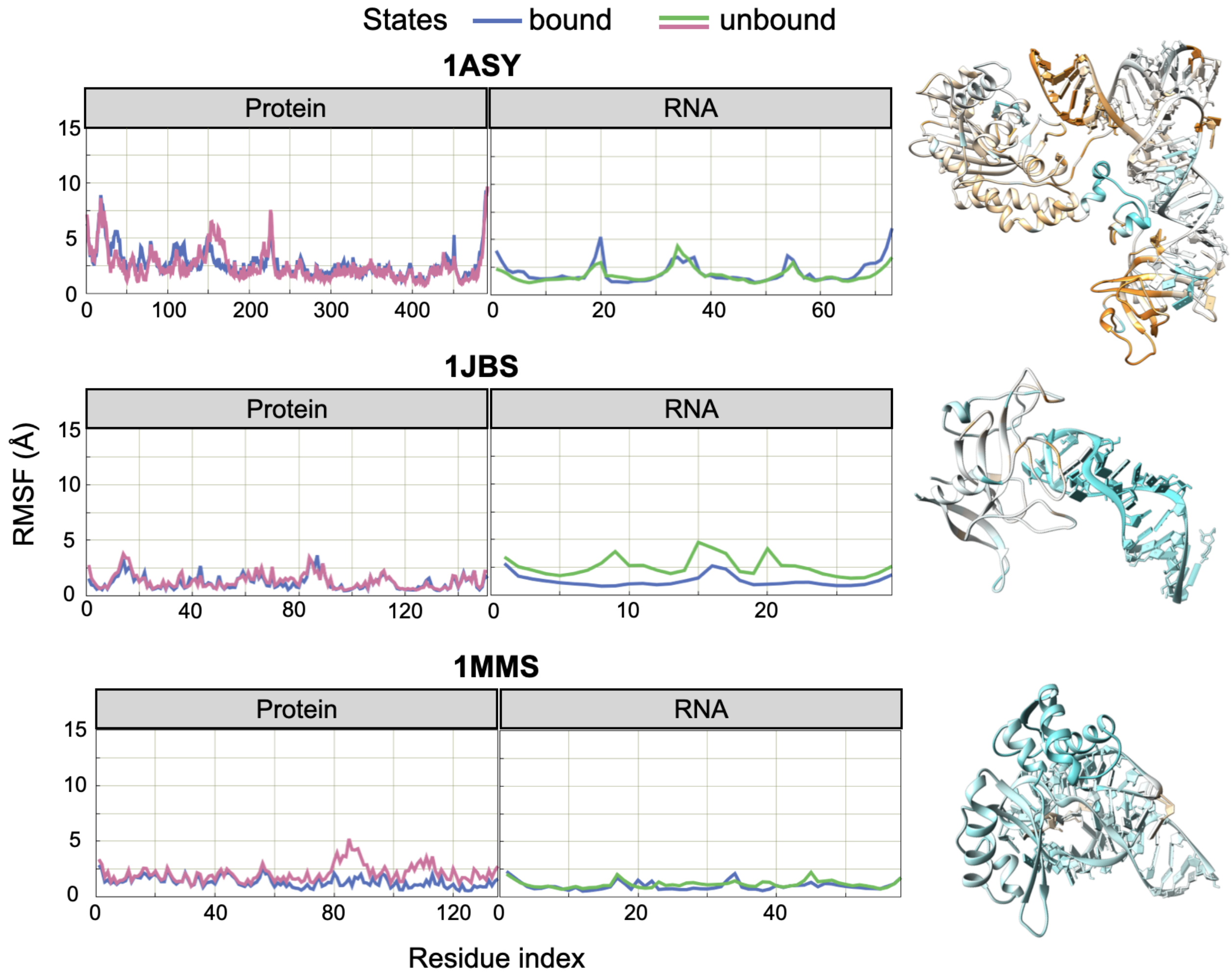


Figure S10: RMSF profiles computed from simulations for proteins or RNA in complexes (blue) and unbound states (proteins in pink and RNA in green) for 1ASY, 1JBS and 1MMS complexes. 3D structures are colored according to the difference in RMSF, between bound and unbound structure, on a scale going from -1.2 Å (cyan) to +1.2 Å (orange).

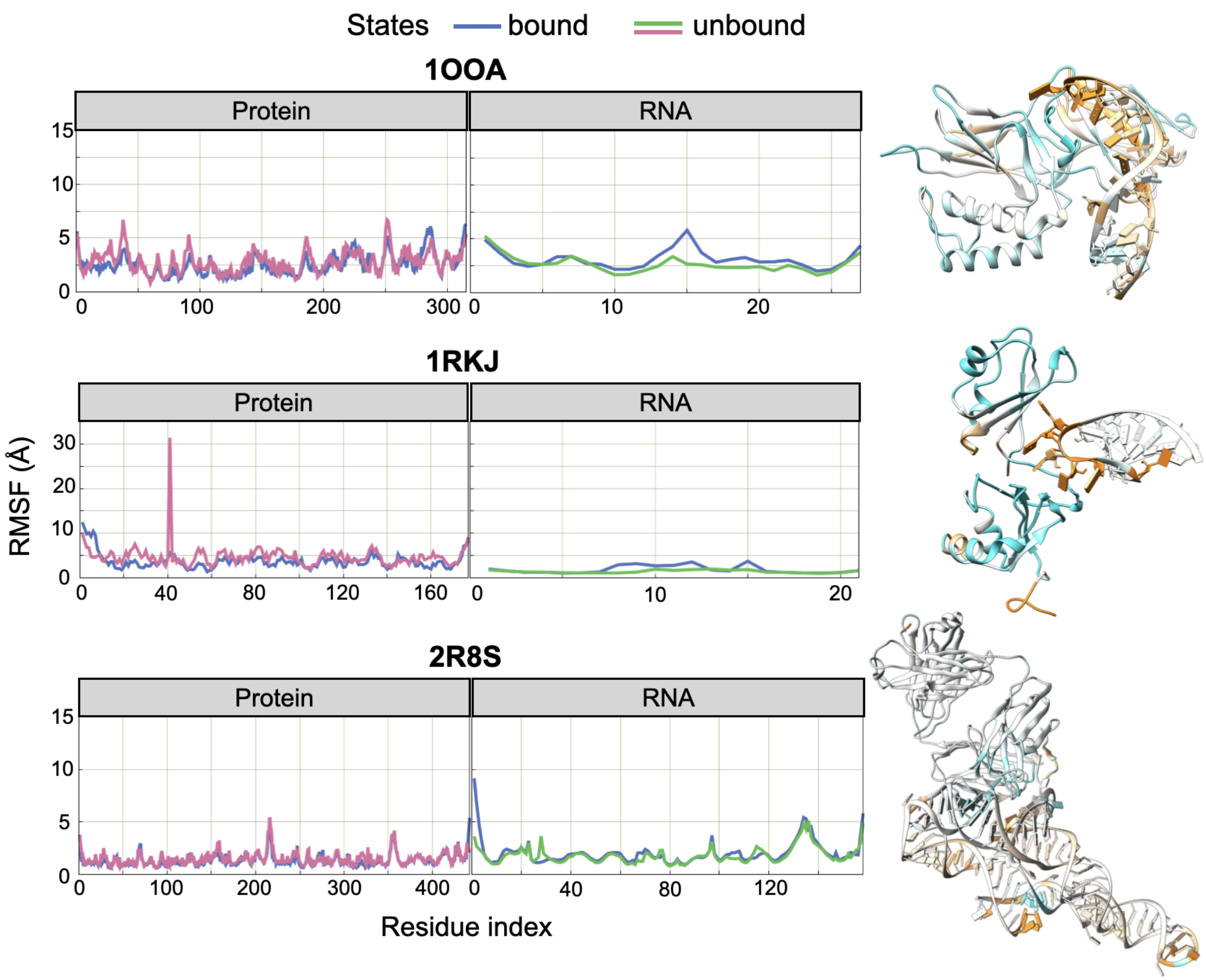


Figure S11: RMSF profiles computed from simulations for proteins or RNA in complexes (blue) and unbound states (proteins in pink and RNA in green) for 1OOA, 1RKJ and 2R8S complexes. 3D structures are colored according to the difference in RMSF, between bound and unbound structure, on a scale going from -1.2 Å (cyan) to +1.2 Å (orange).

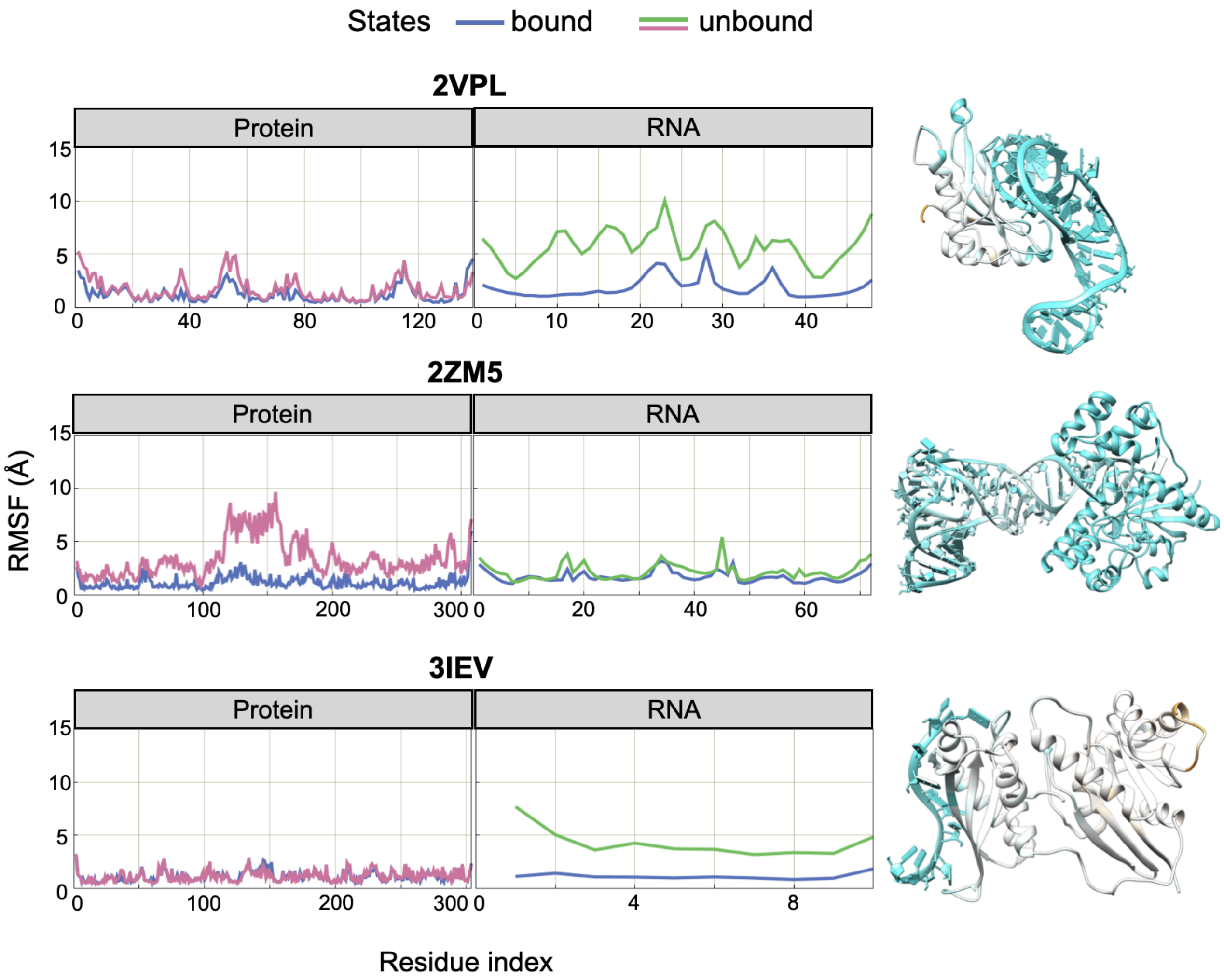


Figure S12: RMSF profiles computed from simulations for proteins or RNA in complexes (blue) and unbound states (proteins in pink and RNA in green) for 2VPL, 2ZM5 and 3IEV complexes. 3D structures are colored according to the difference in RMSF, between bound and unbound structure, on a scale going from -1.2 Å (cyan) to +1.2 Å (orange).

### S4 Free Energy Landscape

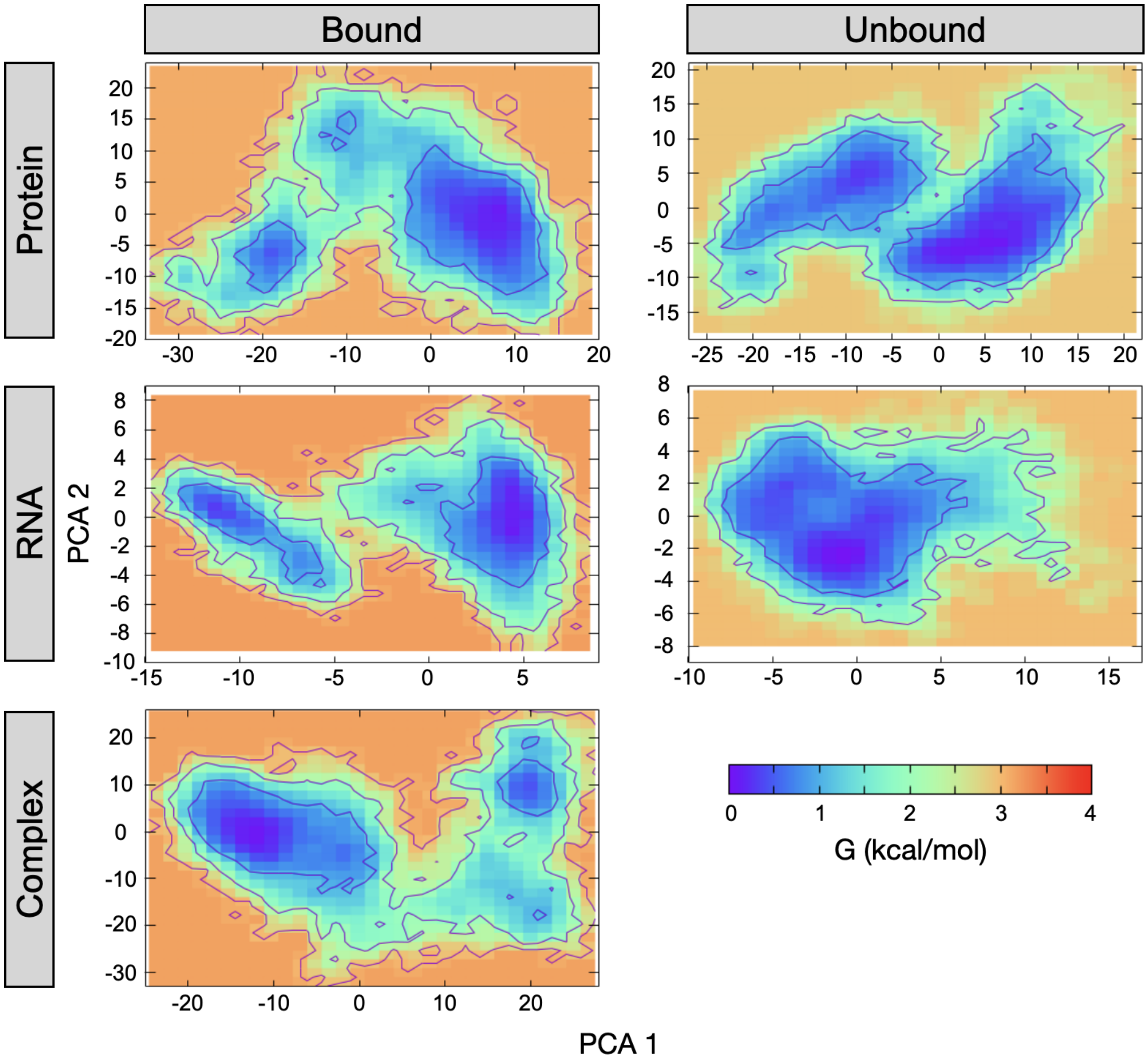


Figure S13: Free energy landscape for the bound and unbound simulation of 1ASY complex. FELs are computed separately for each chain.

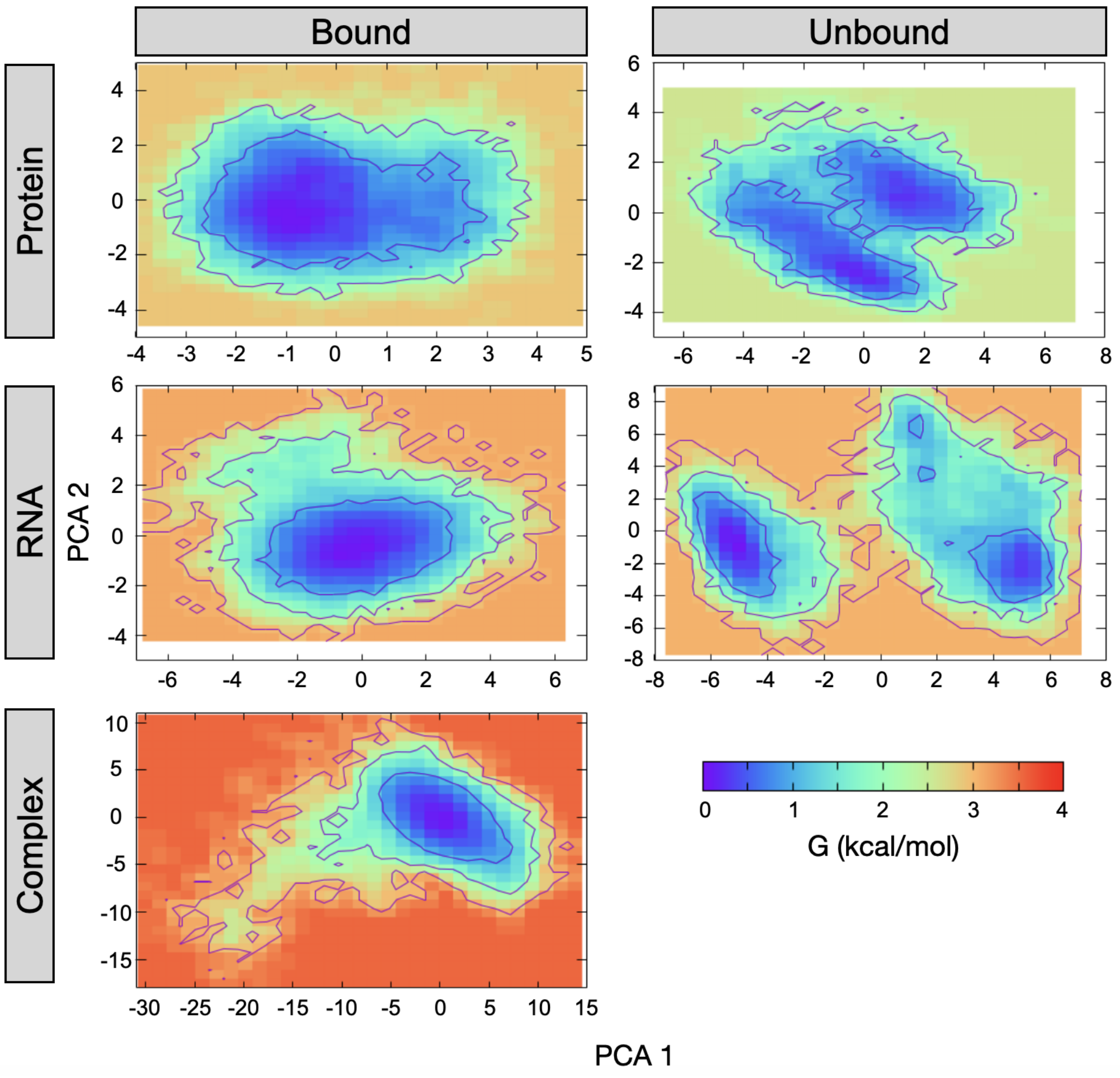


Figure S14: Free energy landscape for the bound and unbound simulation of 1JBS complex. FELs are computed separately for each chain.

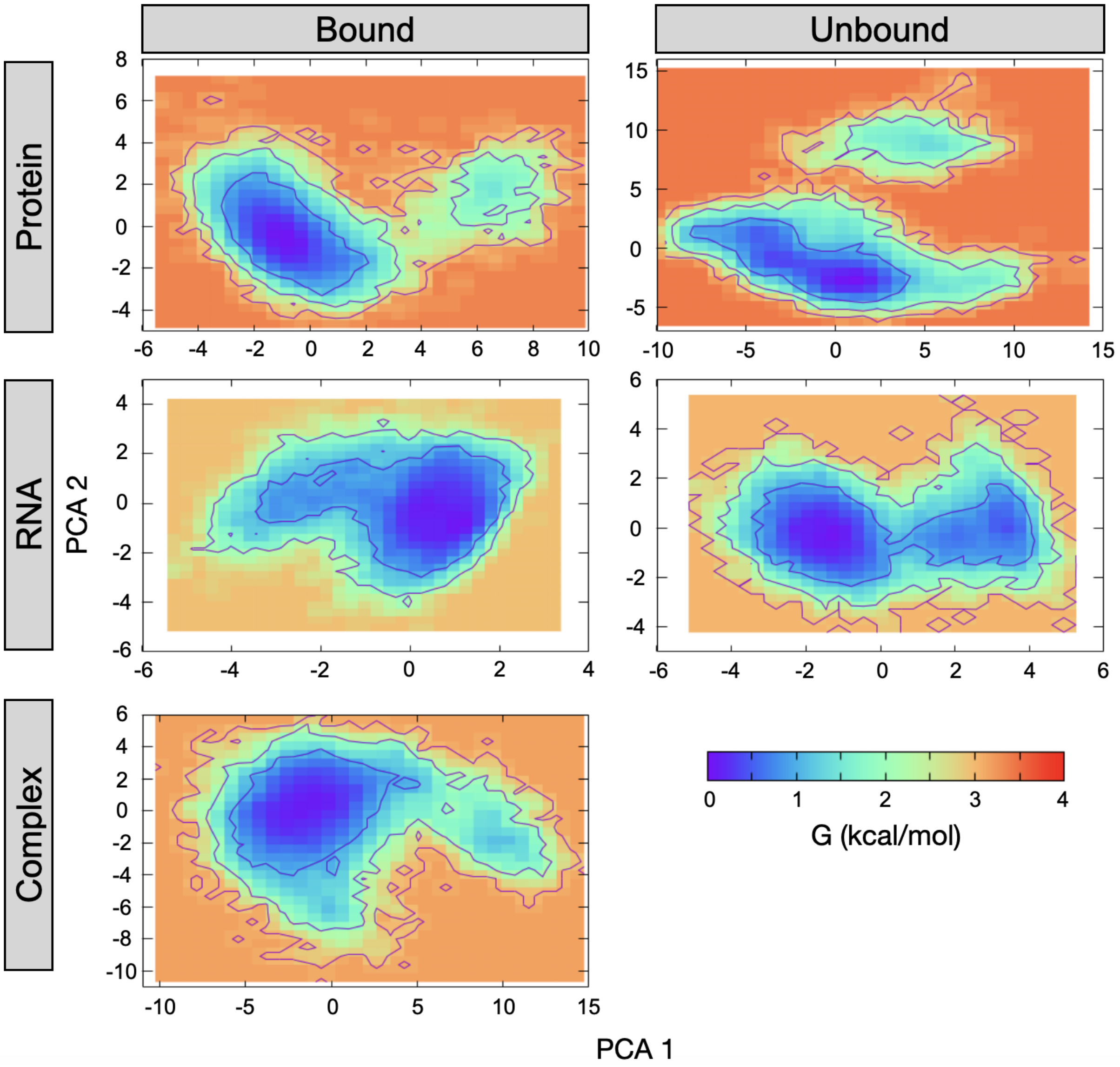


Figure S15: Free energy landscape for the bound and unbound simulation of 1MMS complex. FELs are computed separately for each chain.

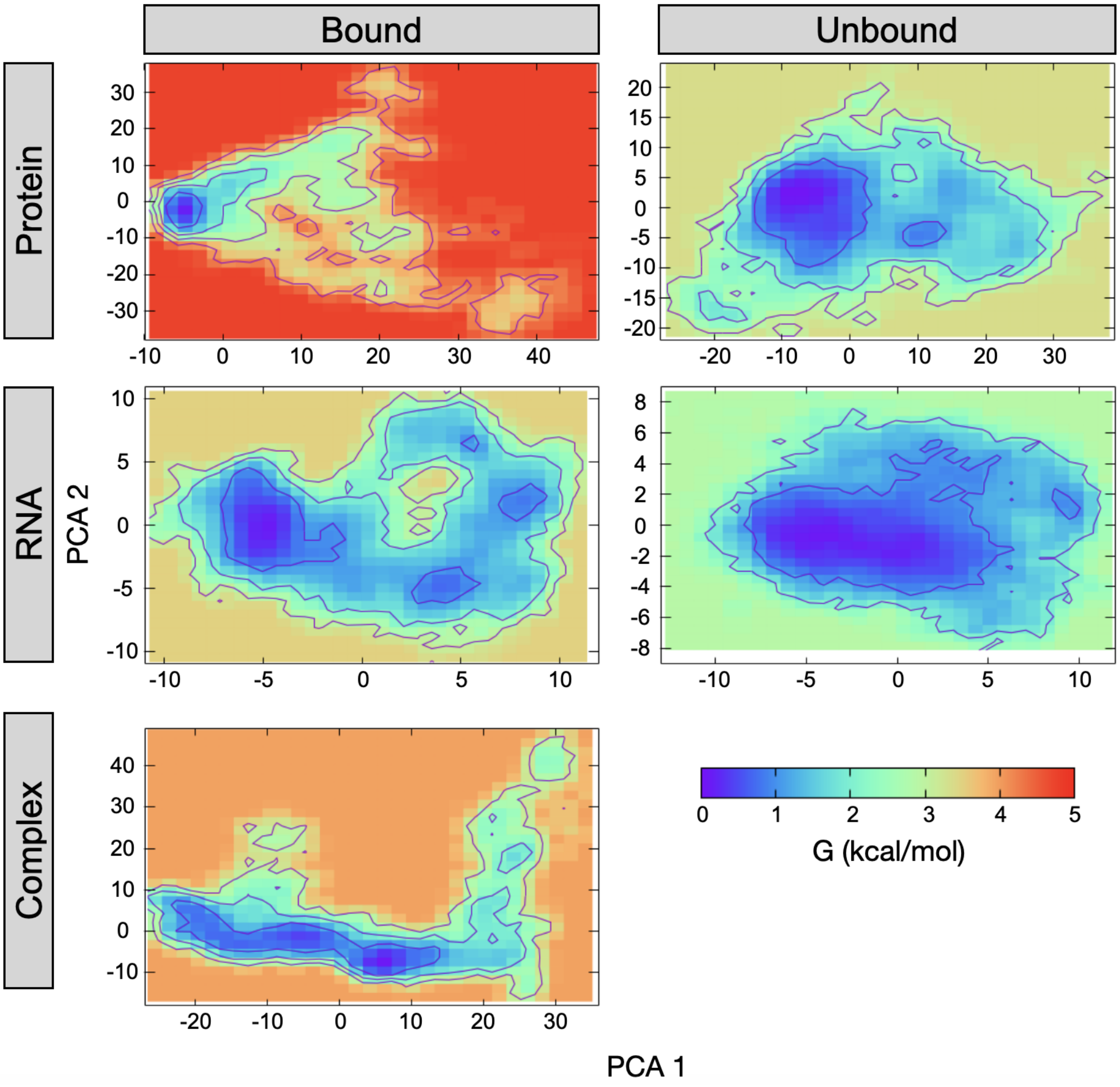


Figure S16: Free energy landscape for the bound and unbound simulation of 1OOA complex. FELs are computed separately for each chain.

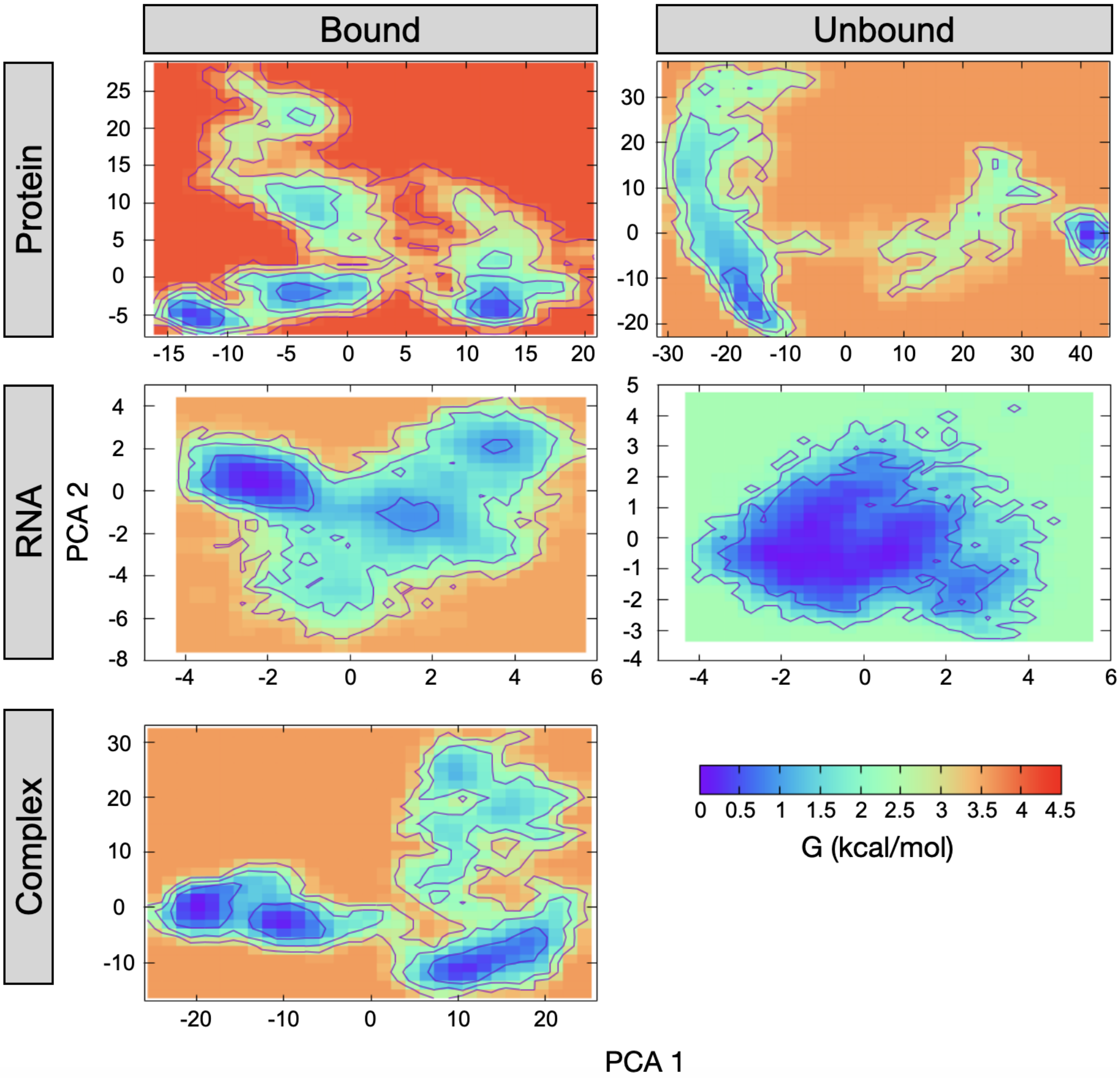


Figure S17: Free energy landscape for the bound and unbound simulation of 1RKJ complex. FELs are computed separately for each chain.

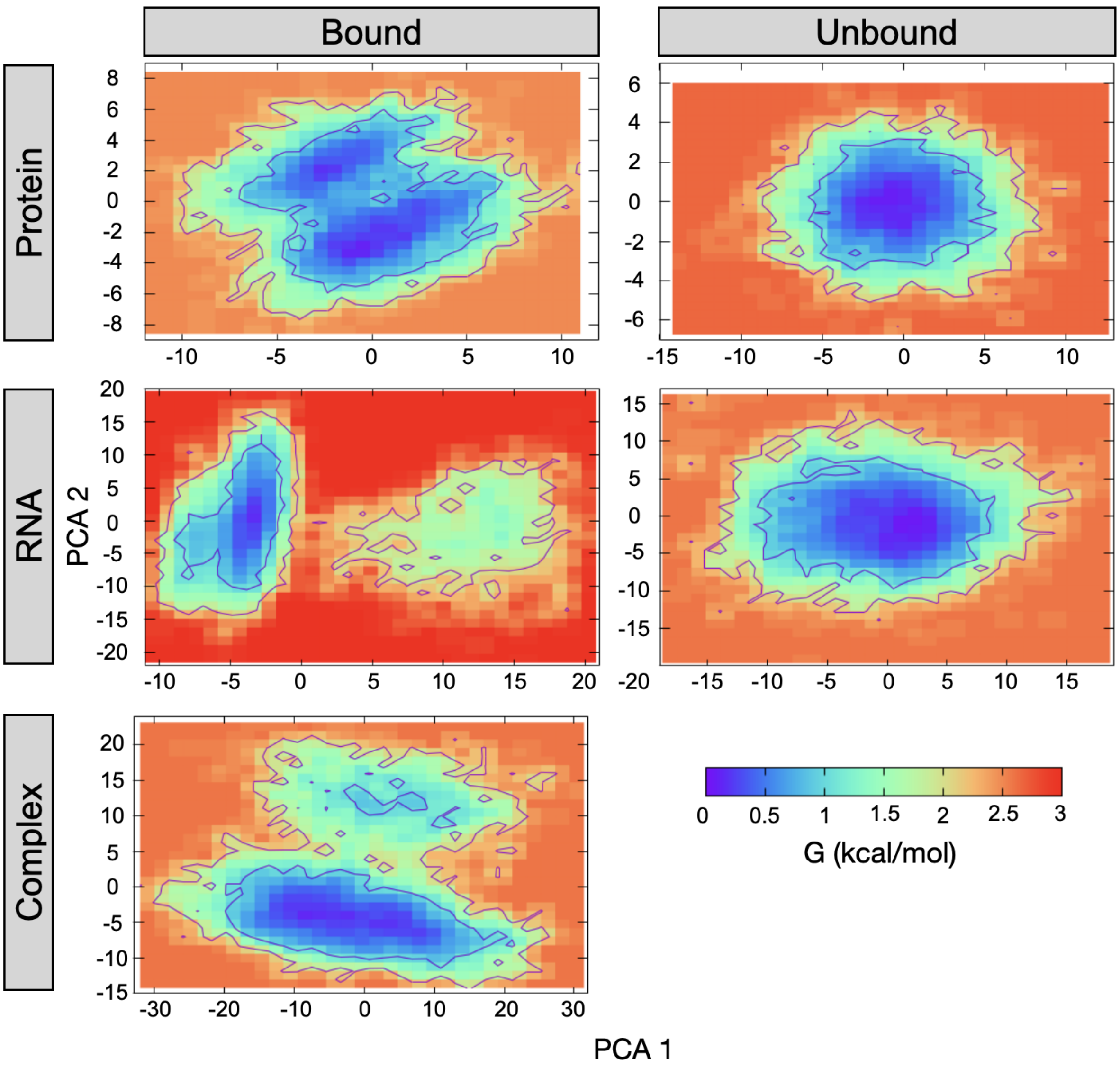


Figure S18: Free energy landscape for the bound and unbound simulation of 2R8S complex. FELs are computed separately for each chain.

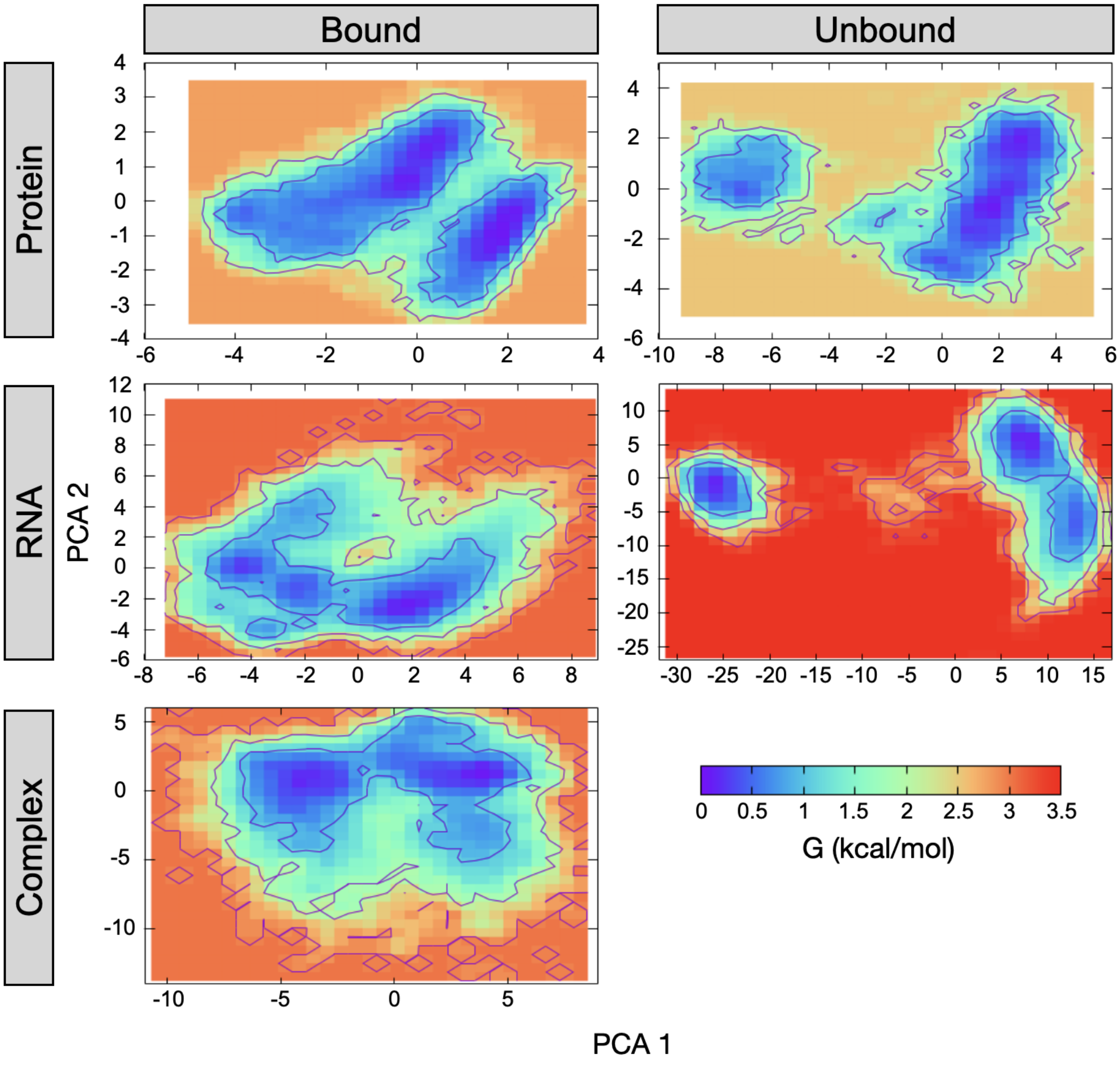


Figure S19: Free energy landscape for the bound and unbound simulation of 2VPL complex. FELs are computed separately for each chain.

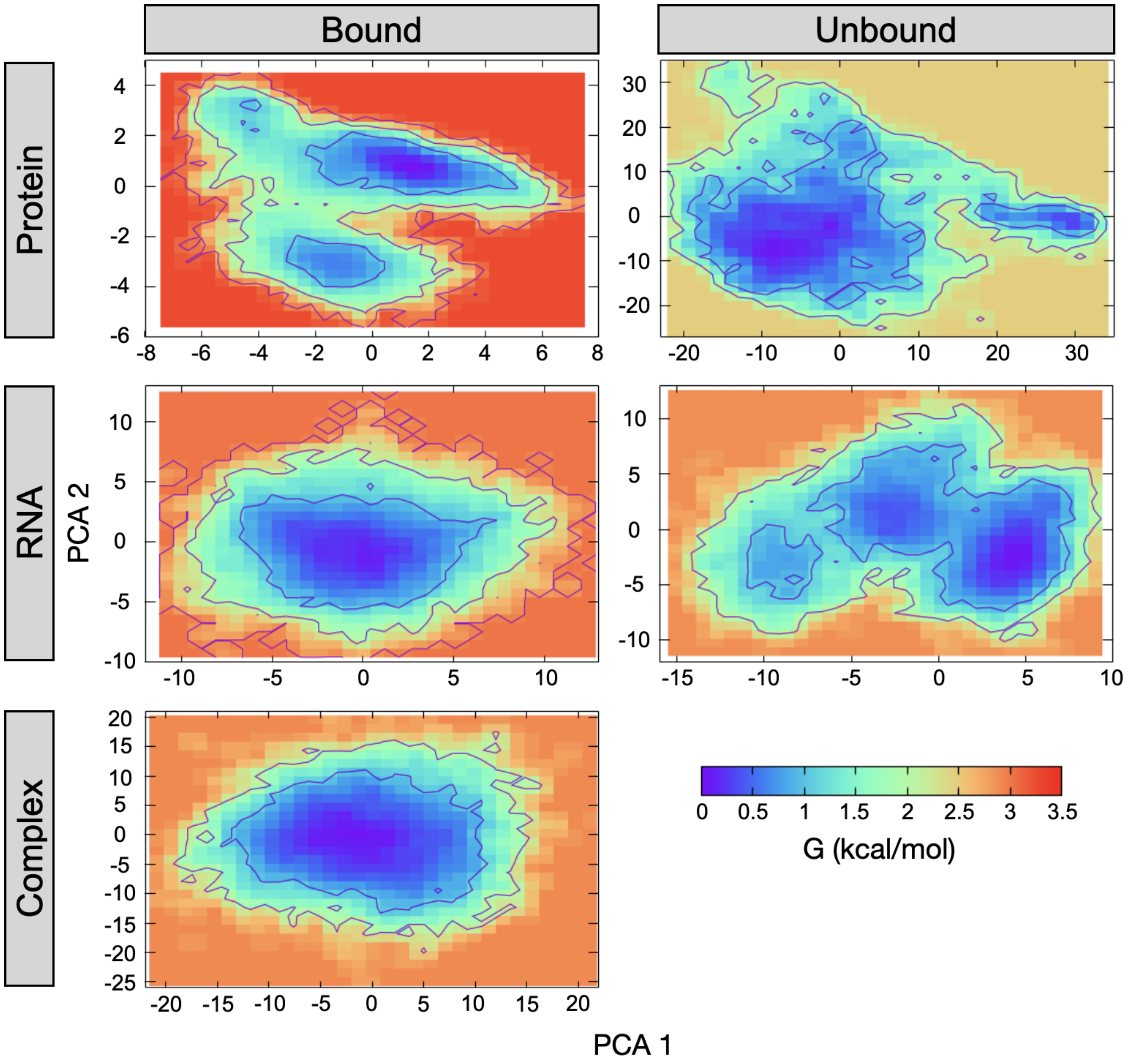


Figure S20: Free energy landscape for the bound and unbound simulation of 2ZM5 complex. FELs are computed separately for each chain.

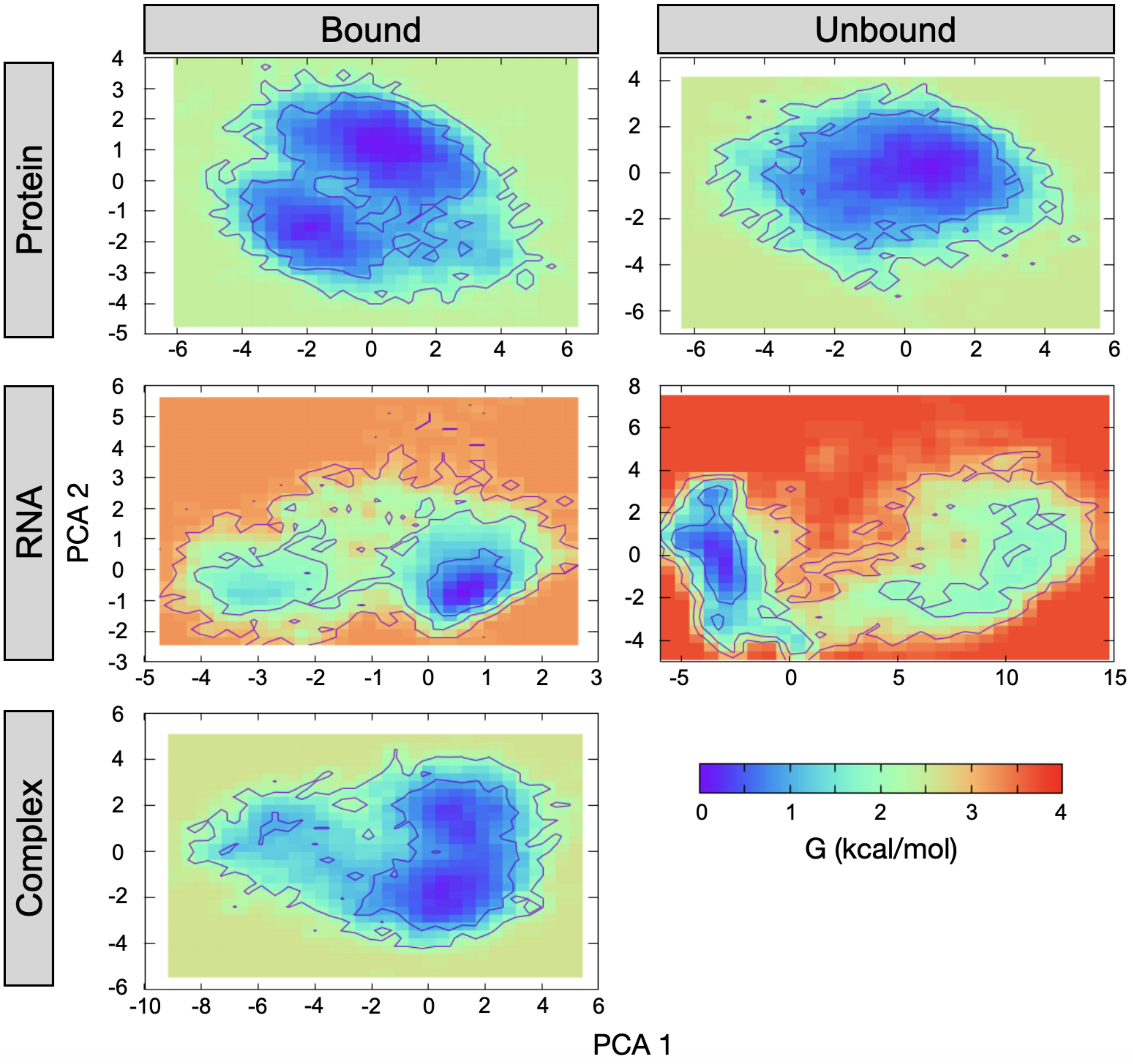


Figure S21: Free energy landscape for the bound and unbound simulation of 3IEV complex. FELs are computed separately for each chain.

### S5 Burried Assessible Surface Area (ΔASA)

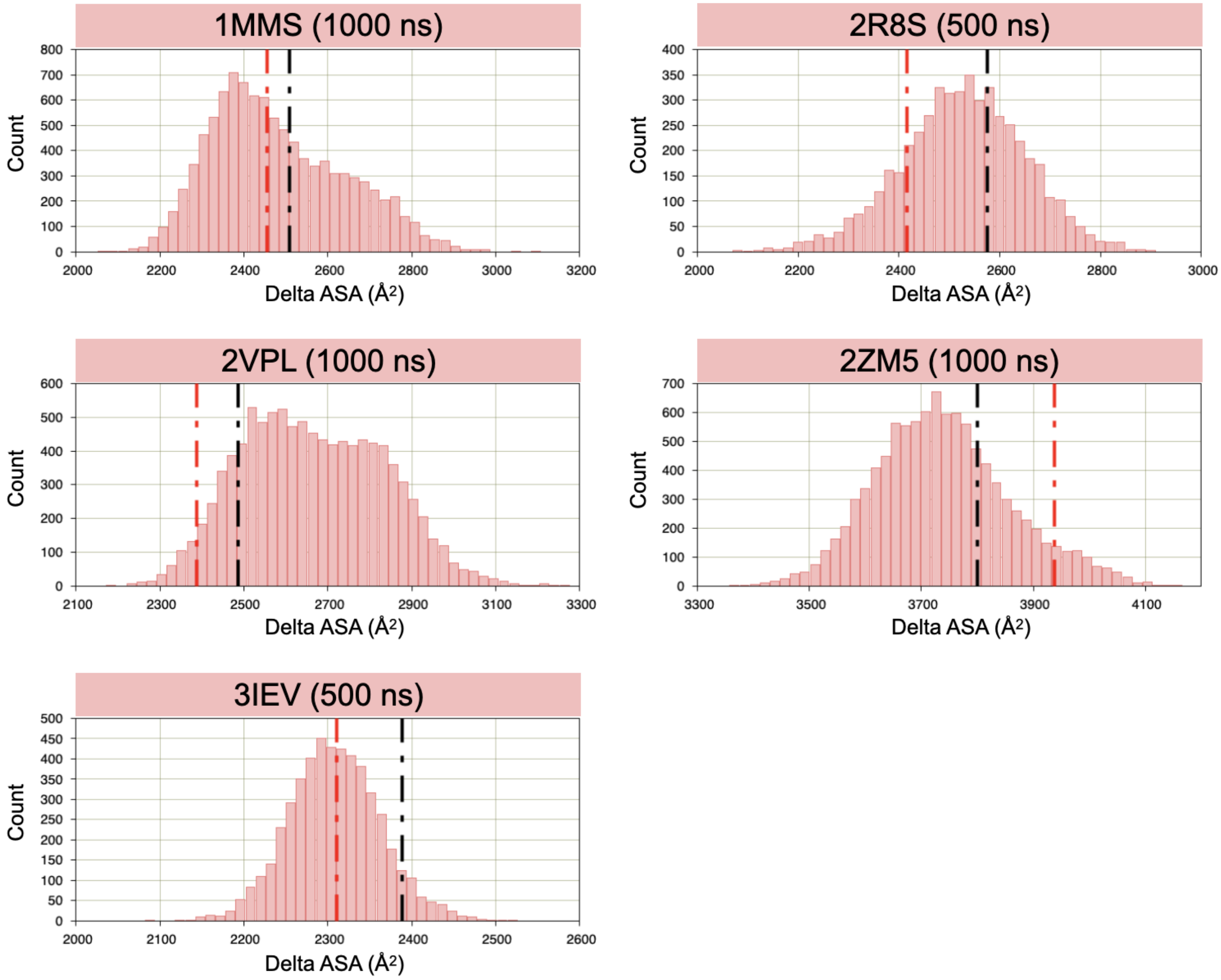


Figure S22: Distribution of interface ΔASA (Å$^2$) for complex with one cluster. The colored ribbon at the top of each plot represents the time series of visited clusters during the simulation. Black dotted lines indicate initial values and red dotted lines indicate values in the crystal structures.

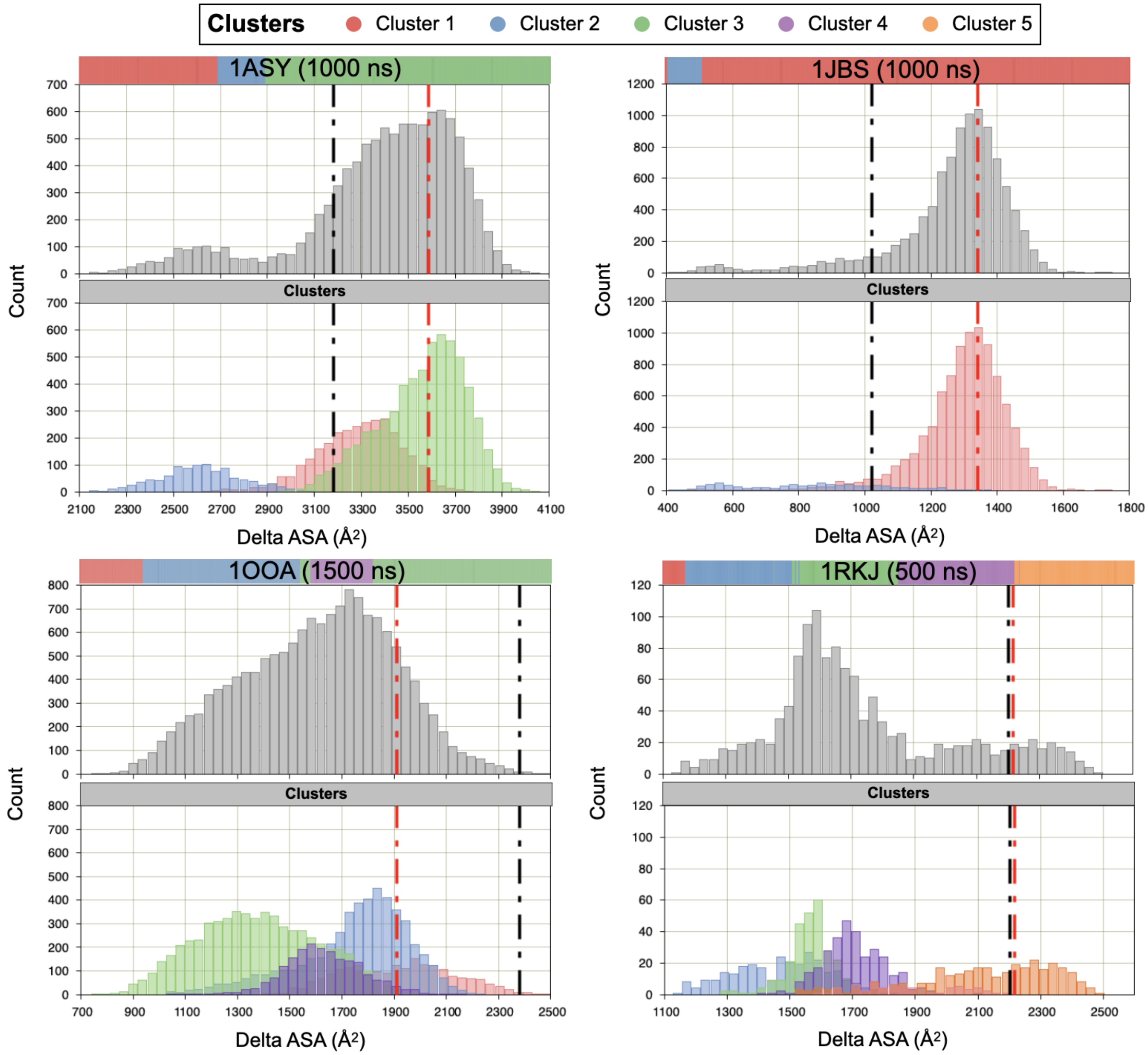


Figure S23: Distribution of interface ΔASA (Å$^2$) for complex with more than one cluster. For each complex, the global distribution is plotted in gray and the distributions in each cluster in colors. The colored ribbon at the top of each plot represents the time series of visited clusters during the simulation. Black dotted lines indicate initial values and red dotted lines indicate values in the crystal structures.

#### S6 Gap Volume

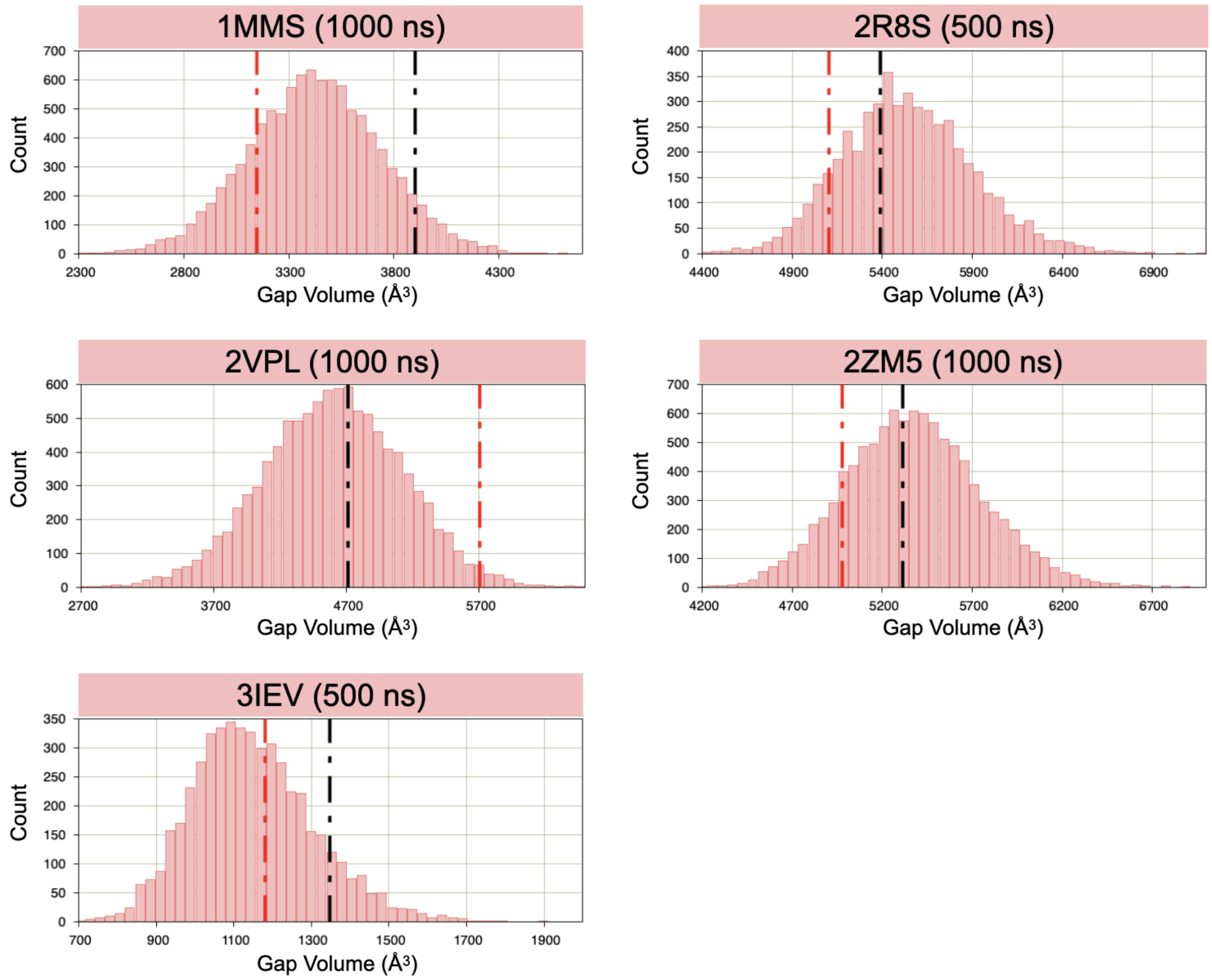


Figure S24: Distribution of gap volume (Å$^3$) for complex with one cluster. The colored ribbon at the top of each plot represents the time series of visited clusters during the simulation. Black dotted lines indicate initial values and red dotted lines indicate values in the crystal structures.

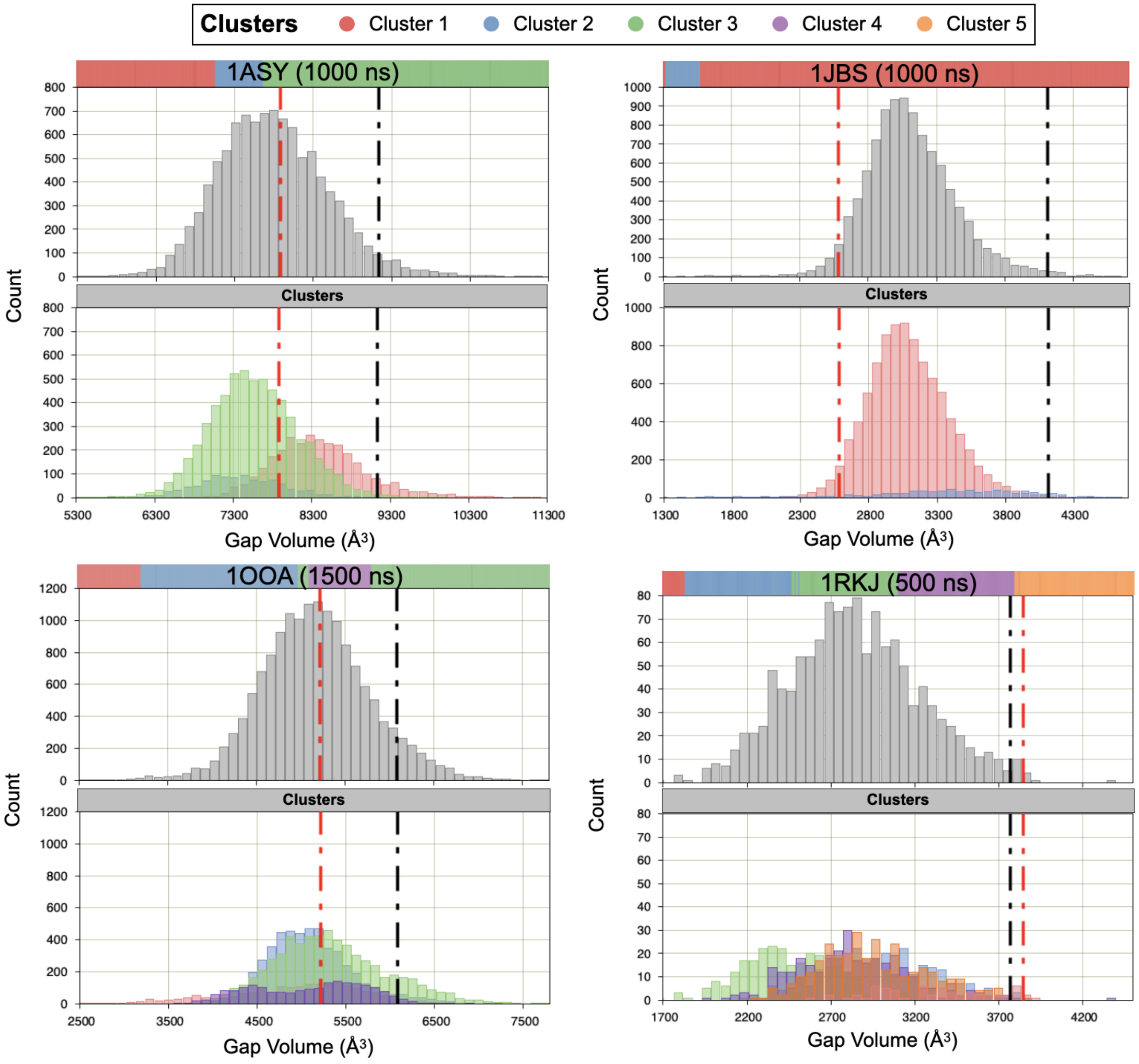


Figure S25: Distribution of gap volume (Å$^3$) for complex with more than one cluster. For each complex, the global distribution is plotted in gray and the distributions in each cluster in colors. The colored ribbon at the top of each plot represents the time series of visited clusters during the simulation. Black dotted lines indicate initial values and red dotted lines indicate values in the crystal structures.

### S7 Gap Index

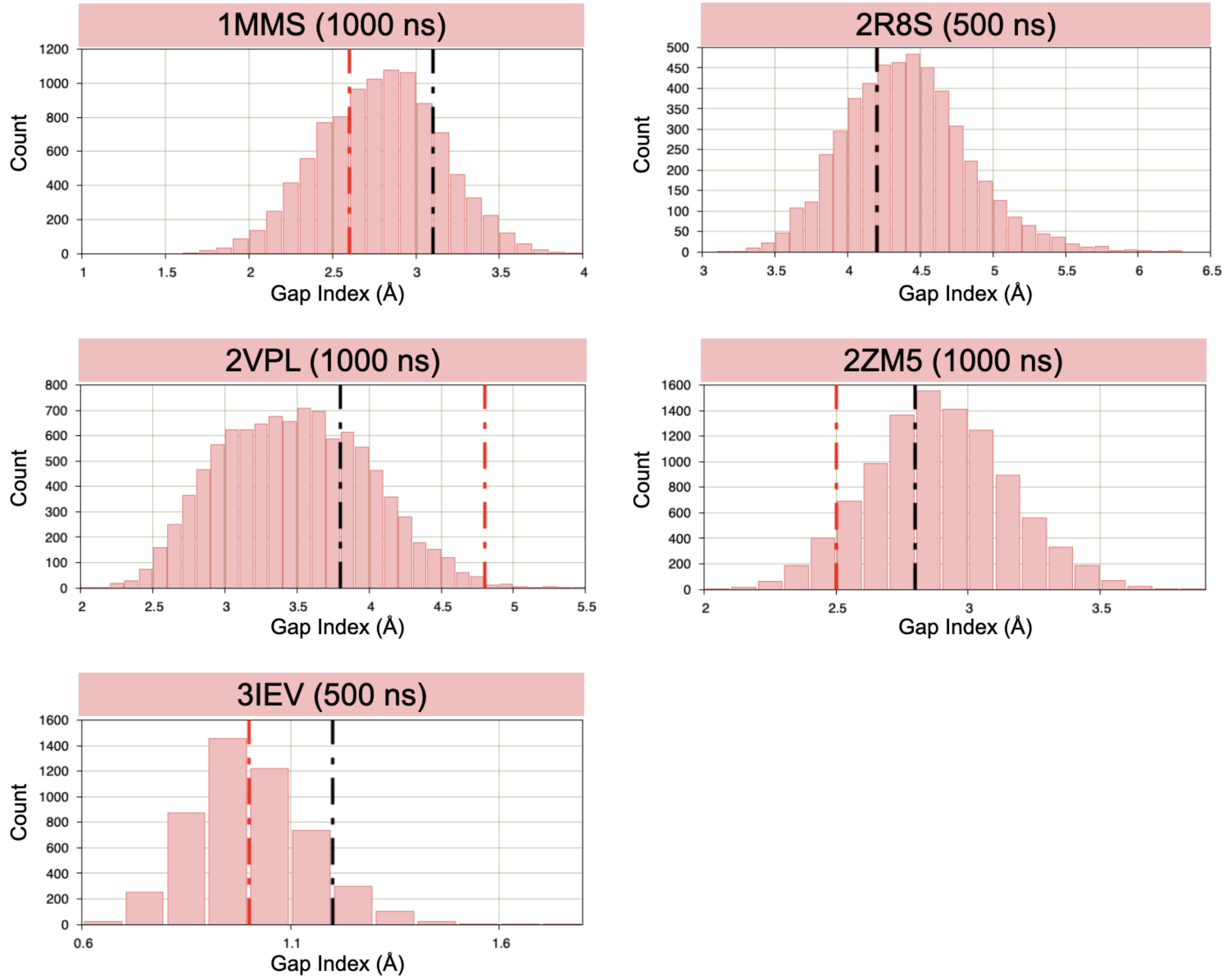


Figure S26: Distribution of gap index (Å) for complex with one cluster. The colored ribbon at the top of each plot represents the time series of visited clusters during the simulation. Black dotted lines indicate initial values and red dotted lines indicate values in the crystal structures.

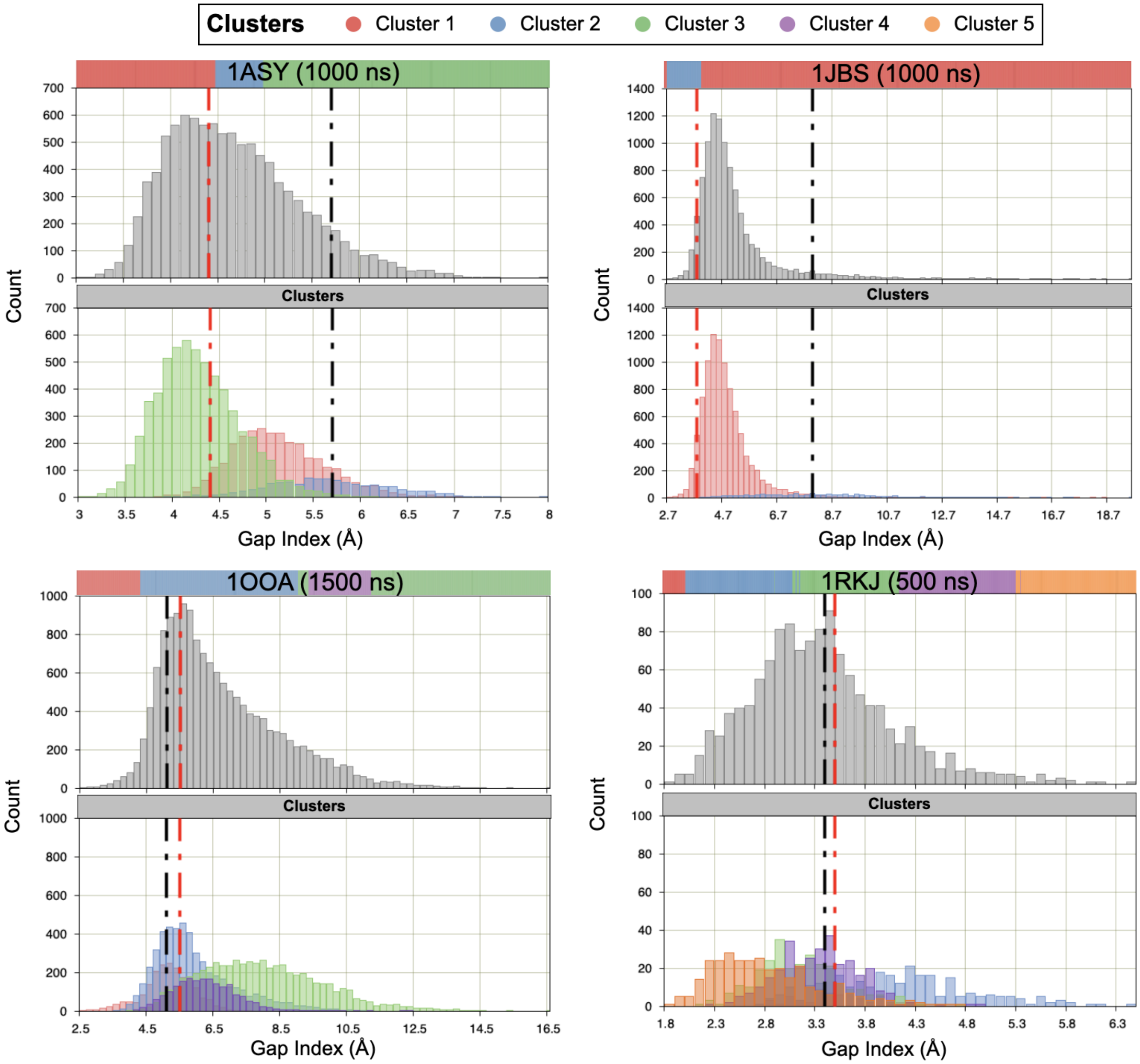


Figure S27: Distribution of gap index (Å) for complex with more than one cluster. For each complex, the global distribution is plotted in gray and the distributions in each cluster in colors. The colored ribbon at the top of each plot represents the time series of visited clusters during the simulation. Black dotted lines indicate initial values and red dotted lines indicate values in the crystal structures.

#### S8 Initial contacts

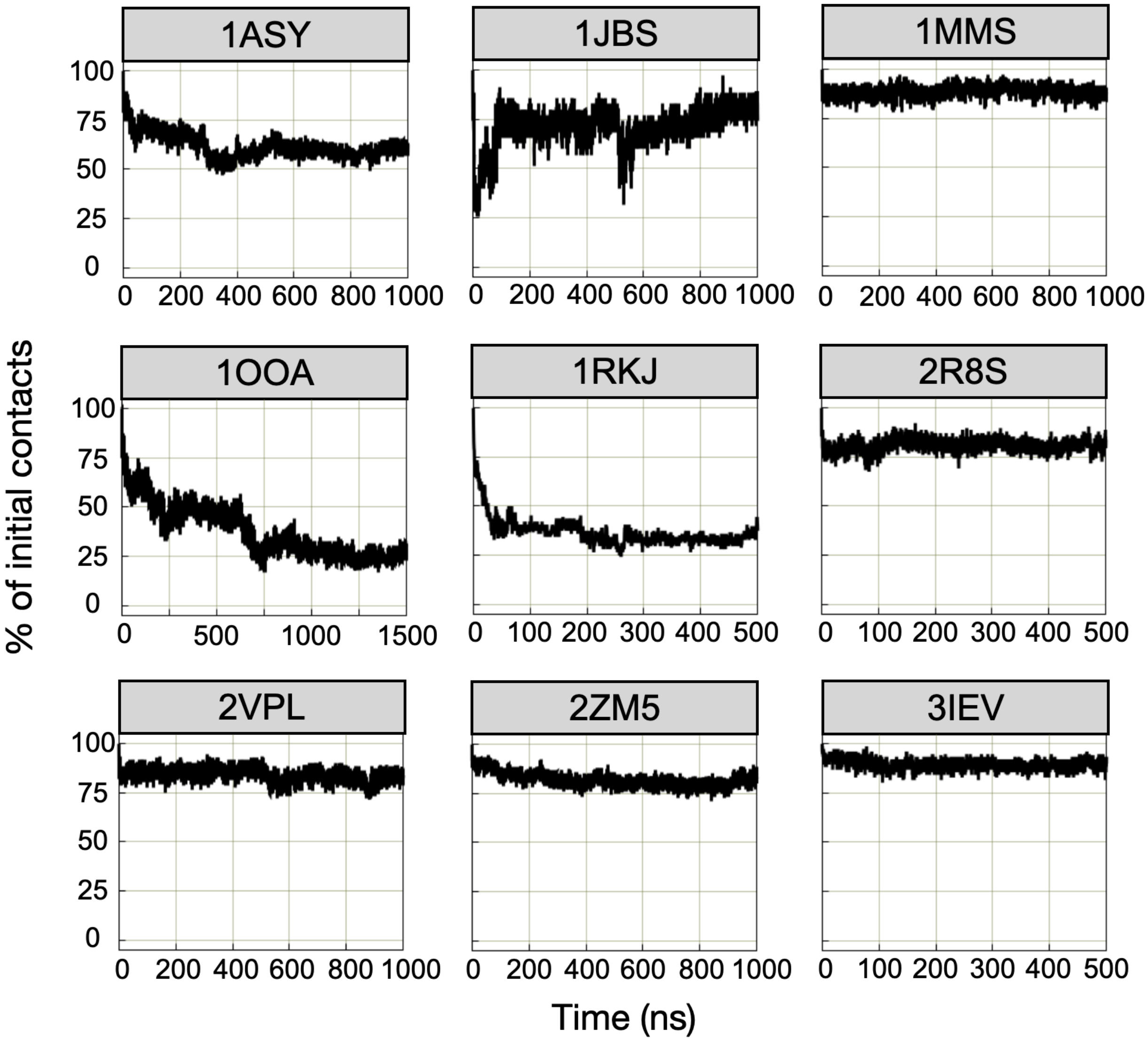


Figure S28: Time series of the preservation of initial contacts for the nine complexes.

#### S9 Principal component analysis

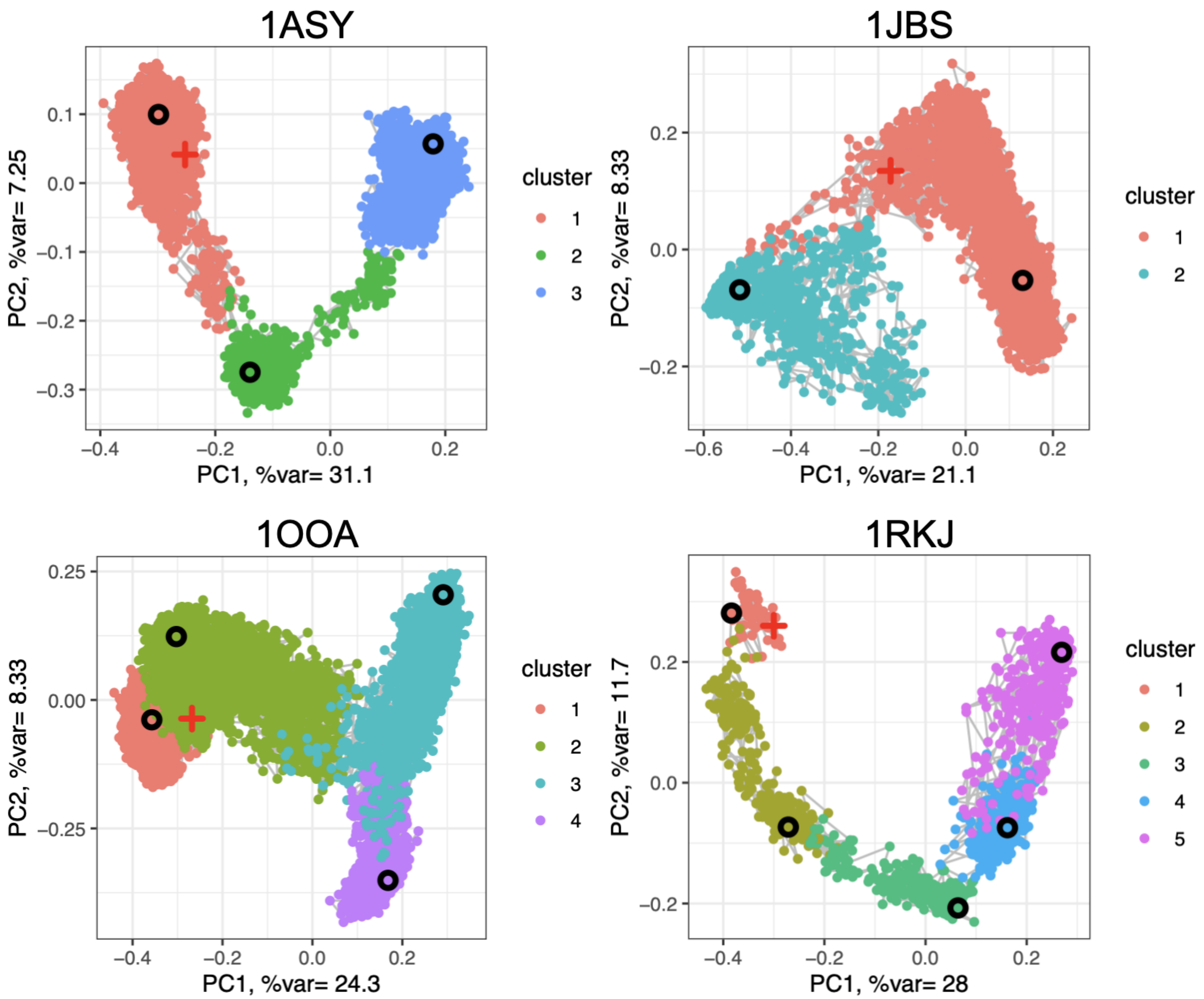


Figure S29: Projection of snapshots and cluster centroids on the first two dimensions of the principal component analysis. Points are colored according to the cluster, connected in their order in time. Red crosses indicate the starting point of the simulations; black circles indicate cluster centroids. The percentage of variance explained by each principal component is indicated on each axis.

### S10 Interface clustering

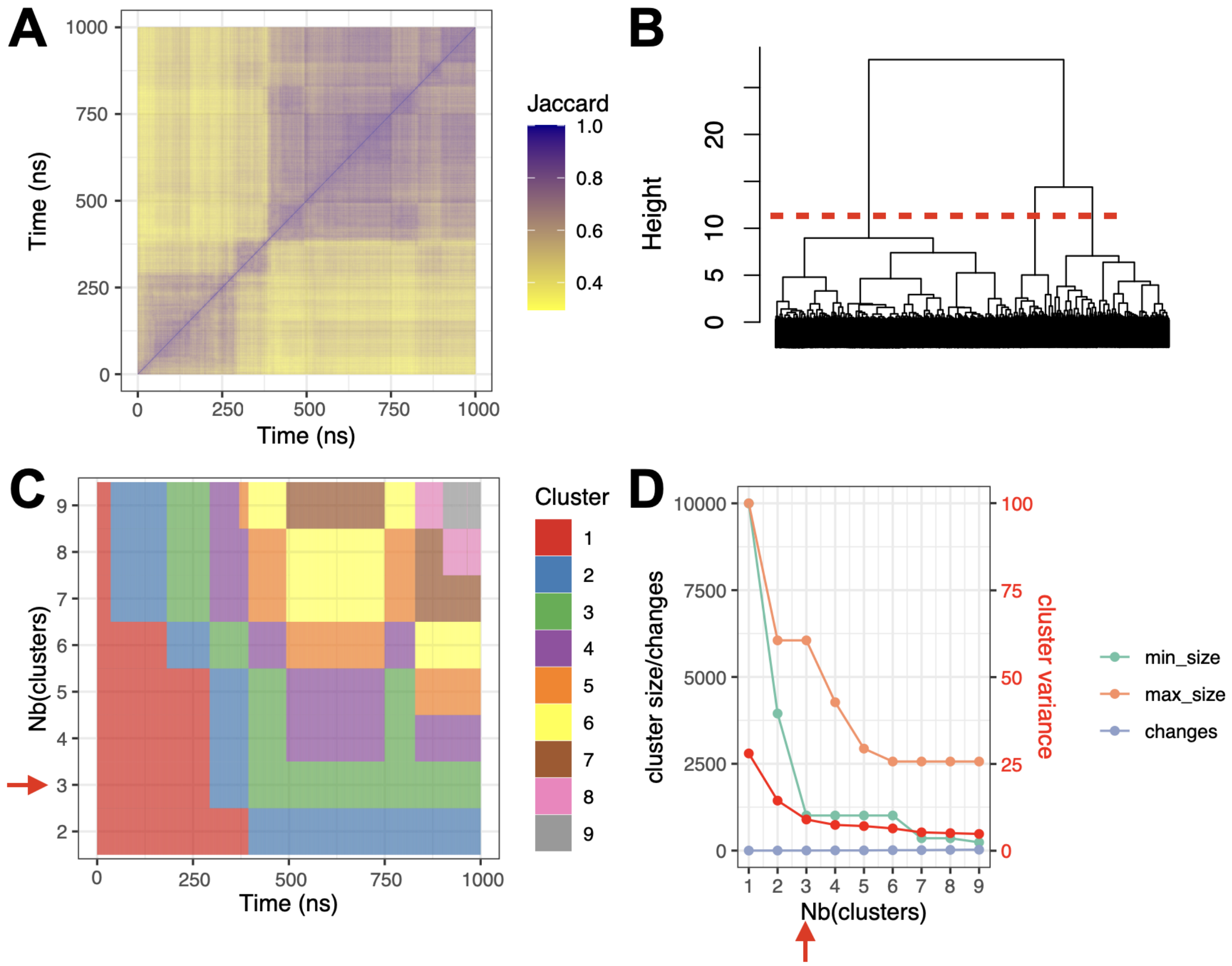


Figure S30: Interface clustering results for complex 1ASY (3 clusters). A. Jaccard similarity matrix. For each pair of snapshots, a yellow pixel indicates low interface similarity and a purple pixel indicates high interface similarity. B. Clustering dendrogram. C. Cluster membership along simulation time, for different numbers of clusters. D. Cluster size, number of changes, and intra-cluster variance (red), for different numbers of clusters. Min size: size of the smallest cluster, max size: size of the largest cluster, changes: number of cluster changes during the simulation. Red dashed line (B) and red arrow (C, D) highlight the optimal number of clusters.

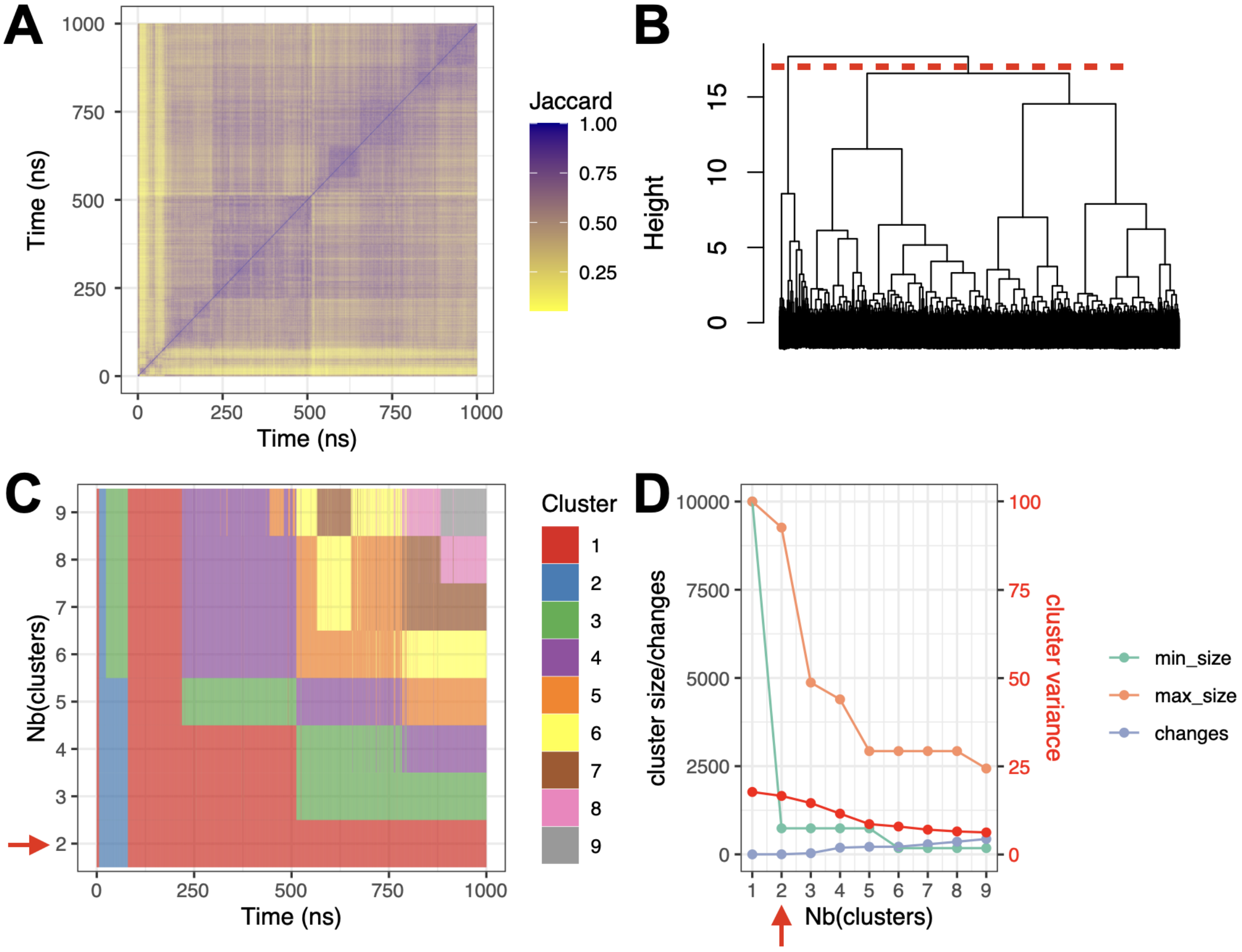


Figure S31: Interface clustering results for complex 1JBS (2 clusters). A. Jaccard similarity matrix. For each pair of snapshots, a yellow pixel indicates low interface similarity and a purple pixel indicates high interface similarity. B. Clustering dendrogram. C. Cluster membership along simulation time, for different numbers of clusters. D. Cluster size, number of changes, and intra-cluster variance (red), for different numbers of clusters. Min size: size of the smallest cluster, max size: size of the largest cluster, changes: number of cluster changes during the simulation. Red dashed line (B) and red arrow (C, D) highlight the optimal number of clusters.

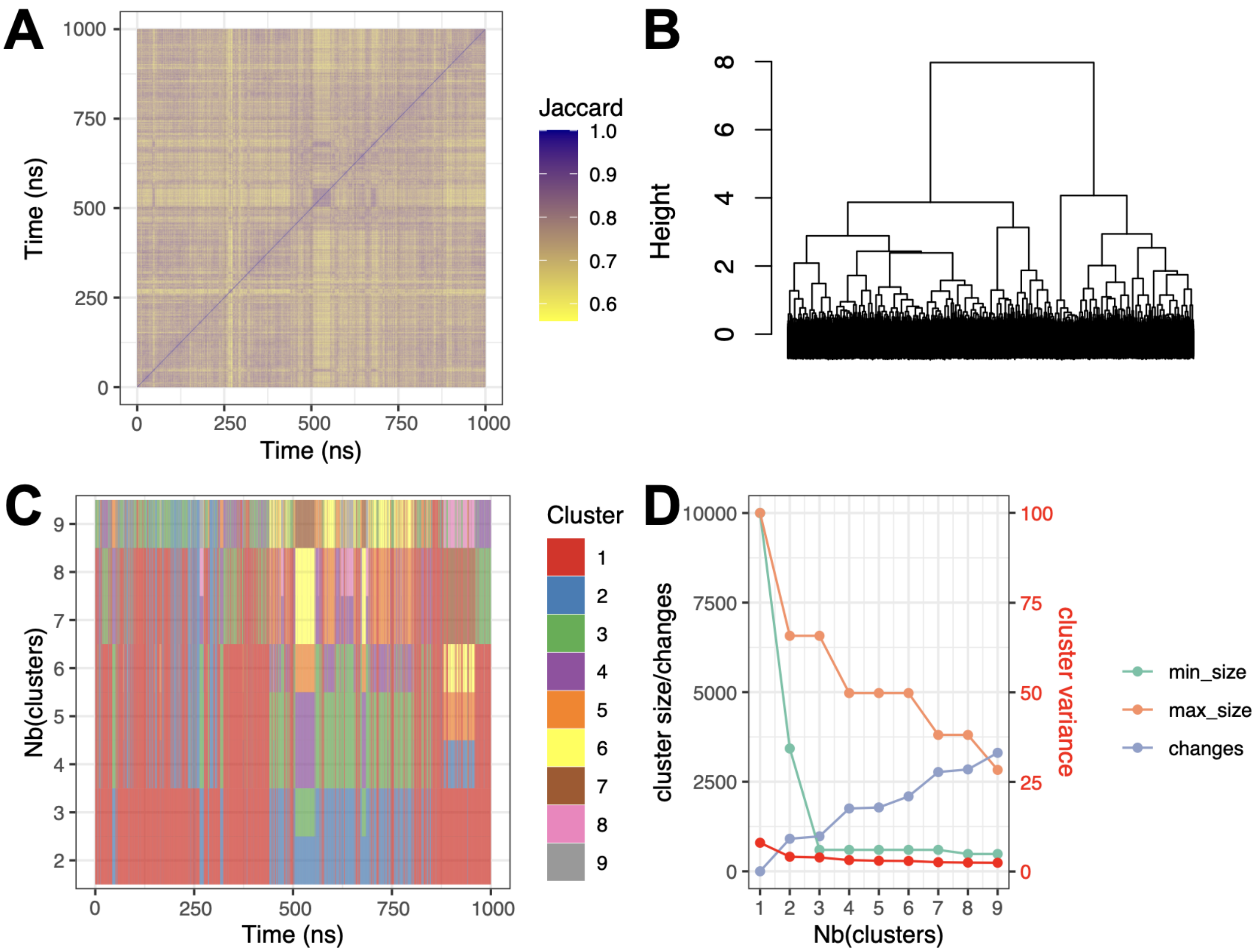


Figure S32: Interface clustering results for complex 1MMS (1 cluster). A. Jaccard similarity matrix. For each pair of snapshots, a yellow pixel indicates low interface similarity and a purple pixel indicates high interface similarity. B. Clustering dendrogram. C. Cluster membership along simulation time, for different numbers of clusters. D. Cluster size, number of changes, and intra-cluster variance (red), for different numbers of clusters. Min size: size of the smallest cluster, max size: size of the largest cluster, changes: number of cluster changes during the simulation.

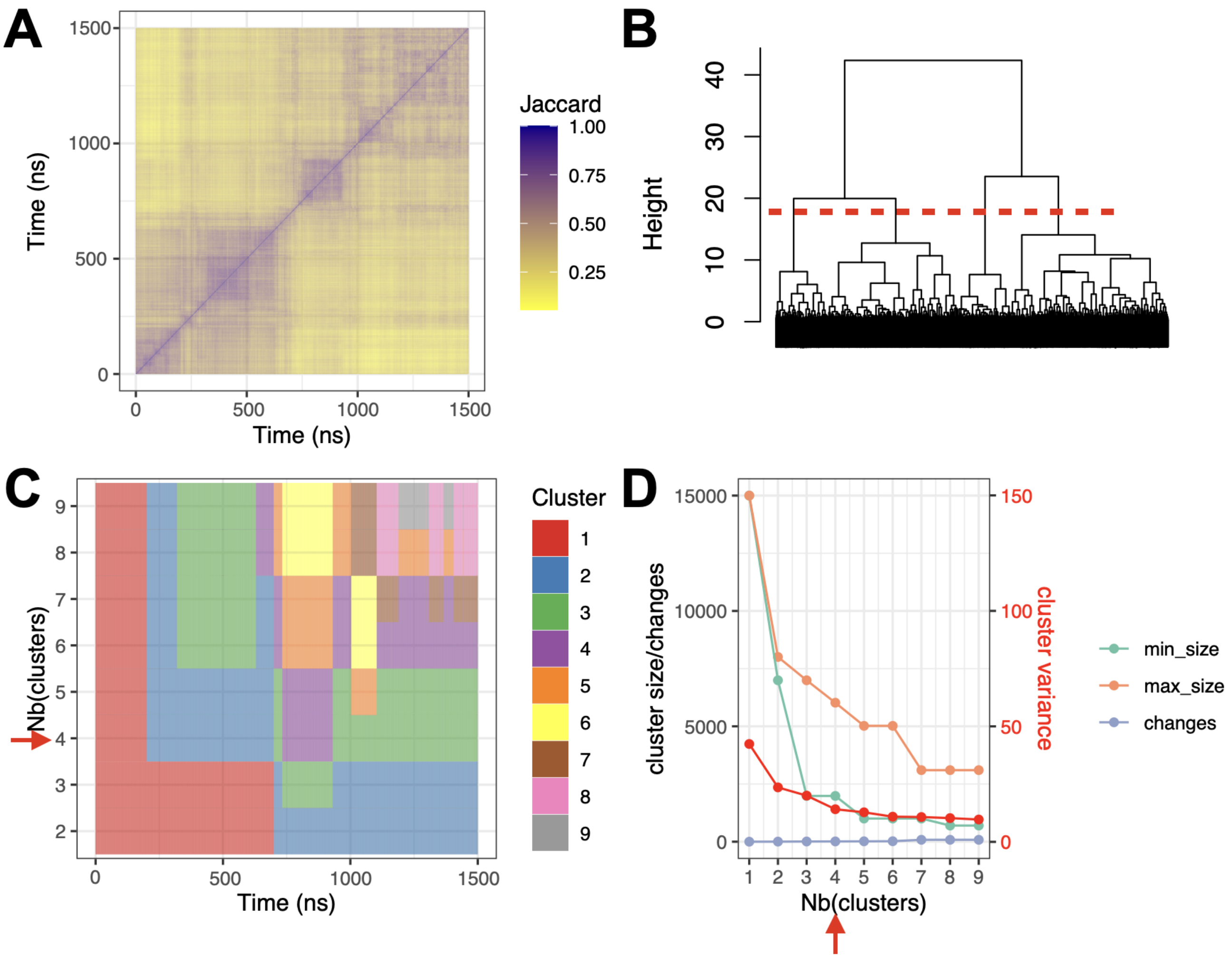


Figure S33: Interface clustering results for complex 1OOA (4 clusters). A. Jaccard similarity matrix. For each pair of snapshots, a yellow pixel indicates low interface similarity and a purple pixel indicates high interface similarity. B. Clustering dendrogram. C. Cluster membership along simulation time, for different numbers of clusters. D. Cluster size, number of changes, and intra-cluster variance (red), for different numbers of clusters. Min size: size of the smallest cluster, max size: size of the largest cluster, changes: number of cluster changes during the simulation. Red dashed line (B) and red arrow (C, D) highlight the optimal number of clusters.

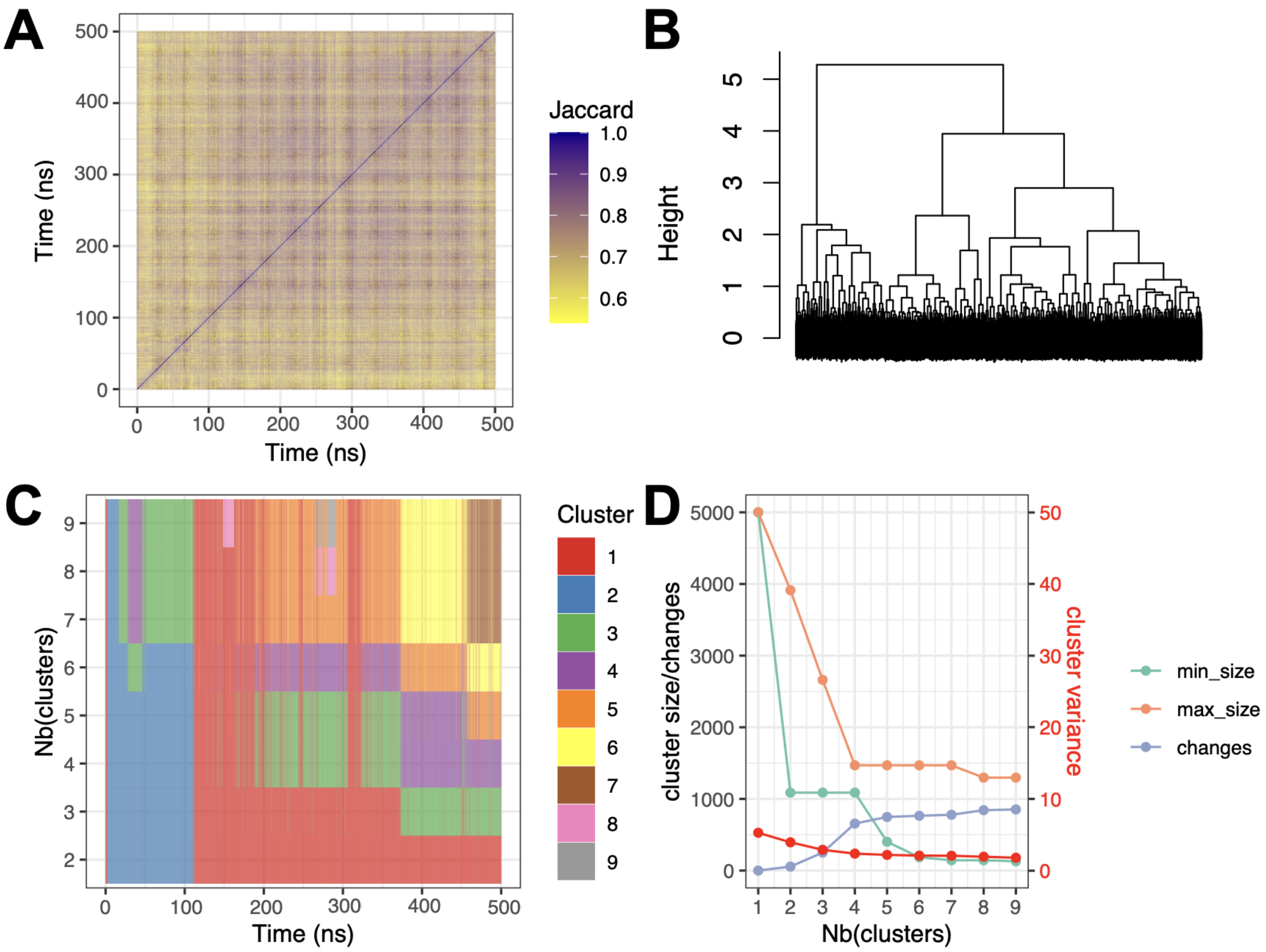


Figure S34: Interface clustering results for complex 2R8S (1 cluster). A. Jaccard similarity matrix. For each pair of snapshots, a yellow pixel indicates low interface similarity and a purple pixel indicates high interface similarity. B. Clustering dendrogram. C. Cluster membership along simulation time, for different numbers of clusters. D. Cluster size, number of changes, and intra-cluster variance (red), for different numbers of clusters. Min size: size of the smallest cluster, max size: size of the largest cluster, changes: number of cluster changes during the simulation.

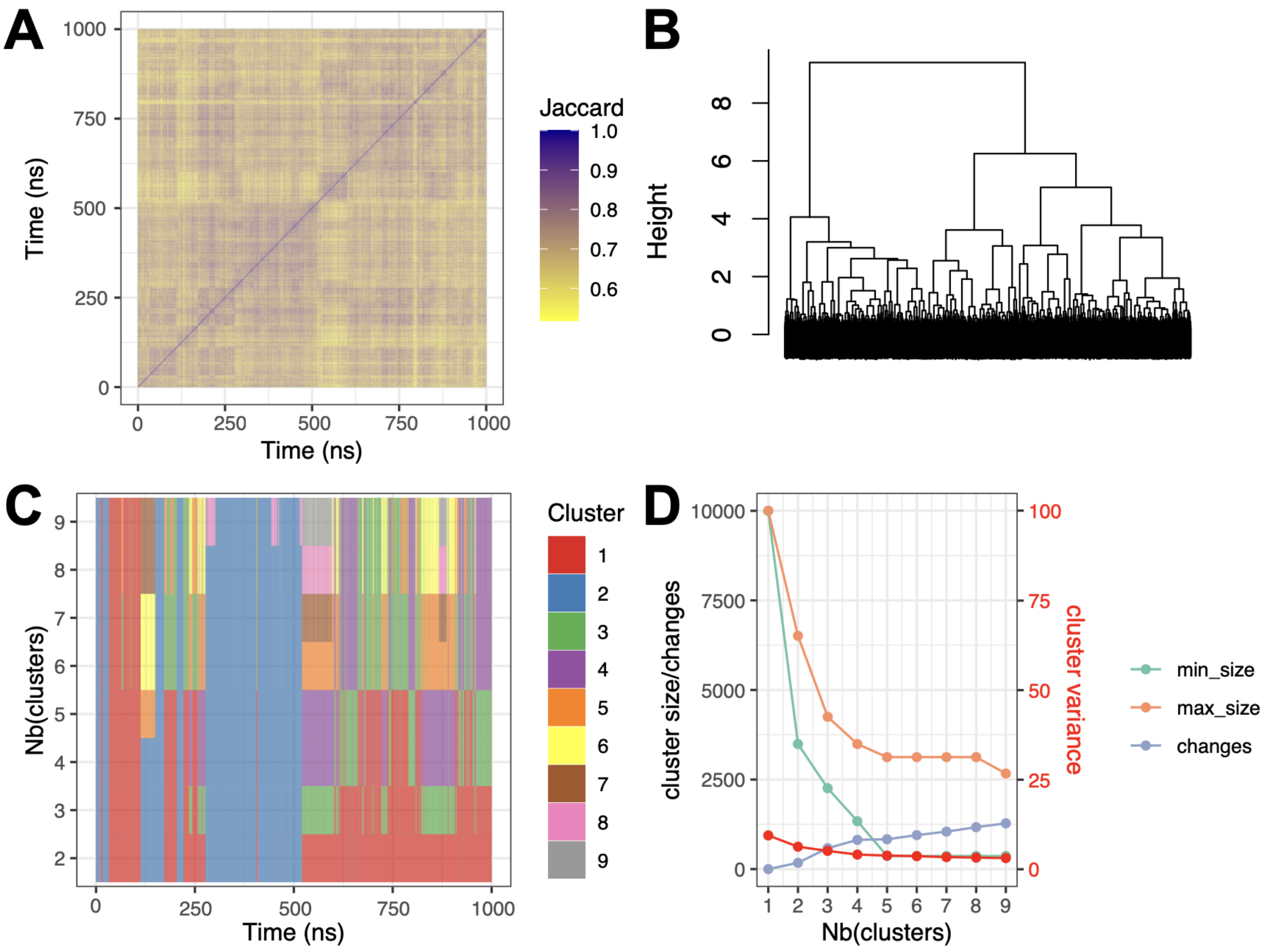


Figure S35: Interface clustering results for complex 2VPL (1 cluster). A. Jaccard similarity matrix. For each pair of snapshots, a yellow pixel indicates low interface similarity and a purple pixel indicates high interface similarity. B. Clustering dendrogram. C. Cluster membership along simulation time, for different numbers of clusters. D. Cluster size, number of changes, and intra-cluster variance (red), for different numbers of clusters. Min size: size of the smallest cluster, max size: size of the largest cluster, changes: number of cluster changes during the simulation.

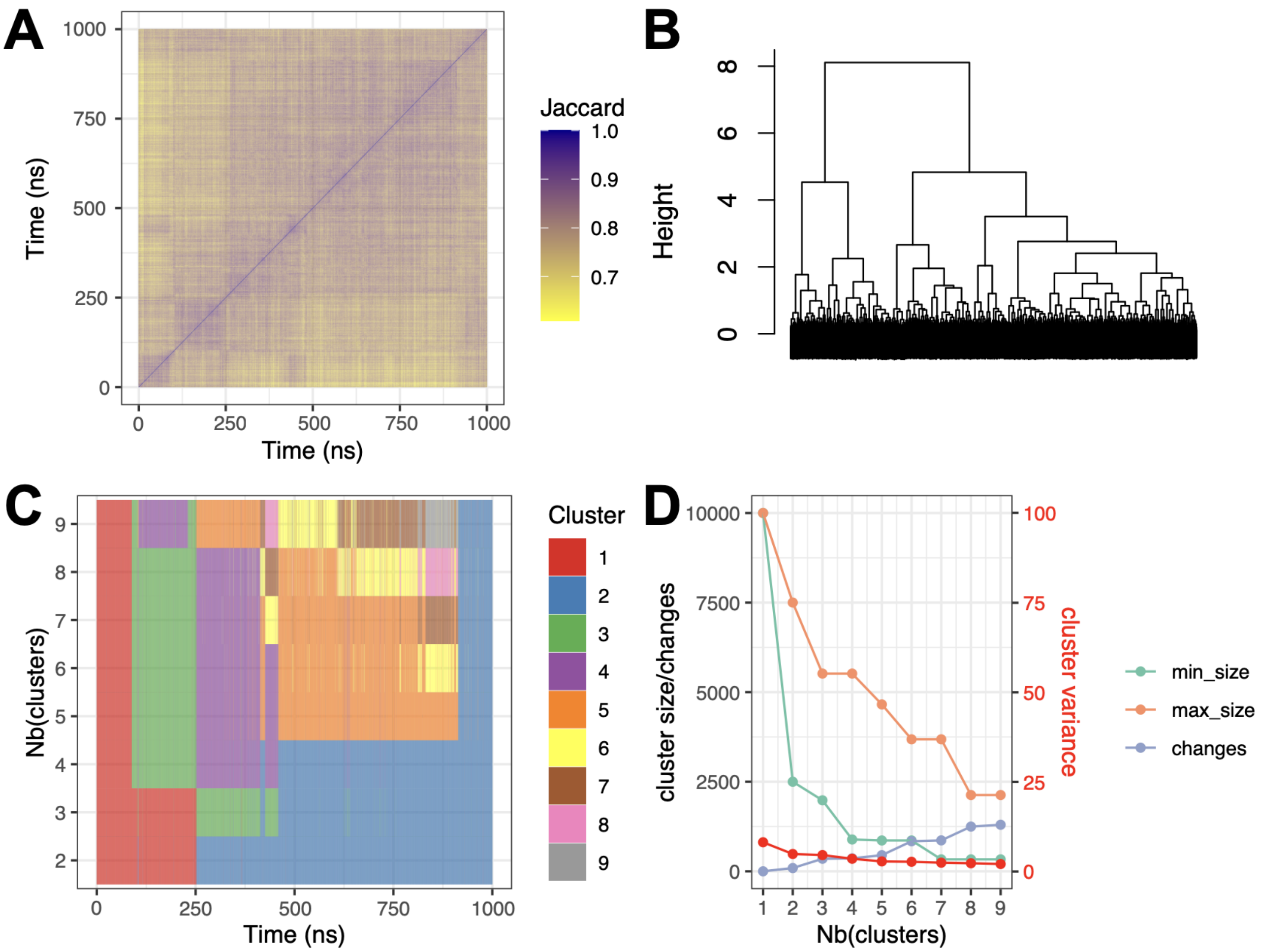


Figure S36: Interface clustering results for complex 2ZM5 (1 cluster). A. Jaccard similarity matrix. For each pair of snapshots, a yellow pixel indicates low interface similarity and a purple pixel indicates high interface similarity. B. Clustering dendrogram. C. Cluster membership along simulation time, for different numbers of clusters. D. Cluster size, number of changes, and intra-cluster variance (red), for different numbers of clusters. Min size: size of the smallest cluster, max size: size of the largest cluster, changes: number of cluster changes during the simulation.

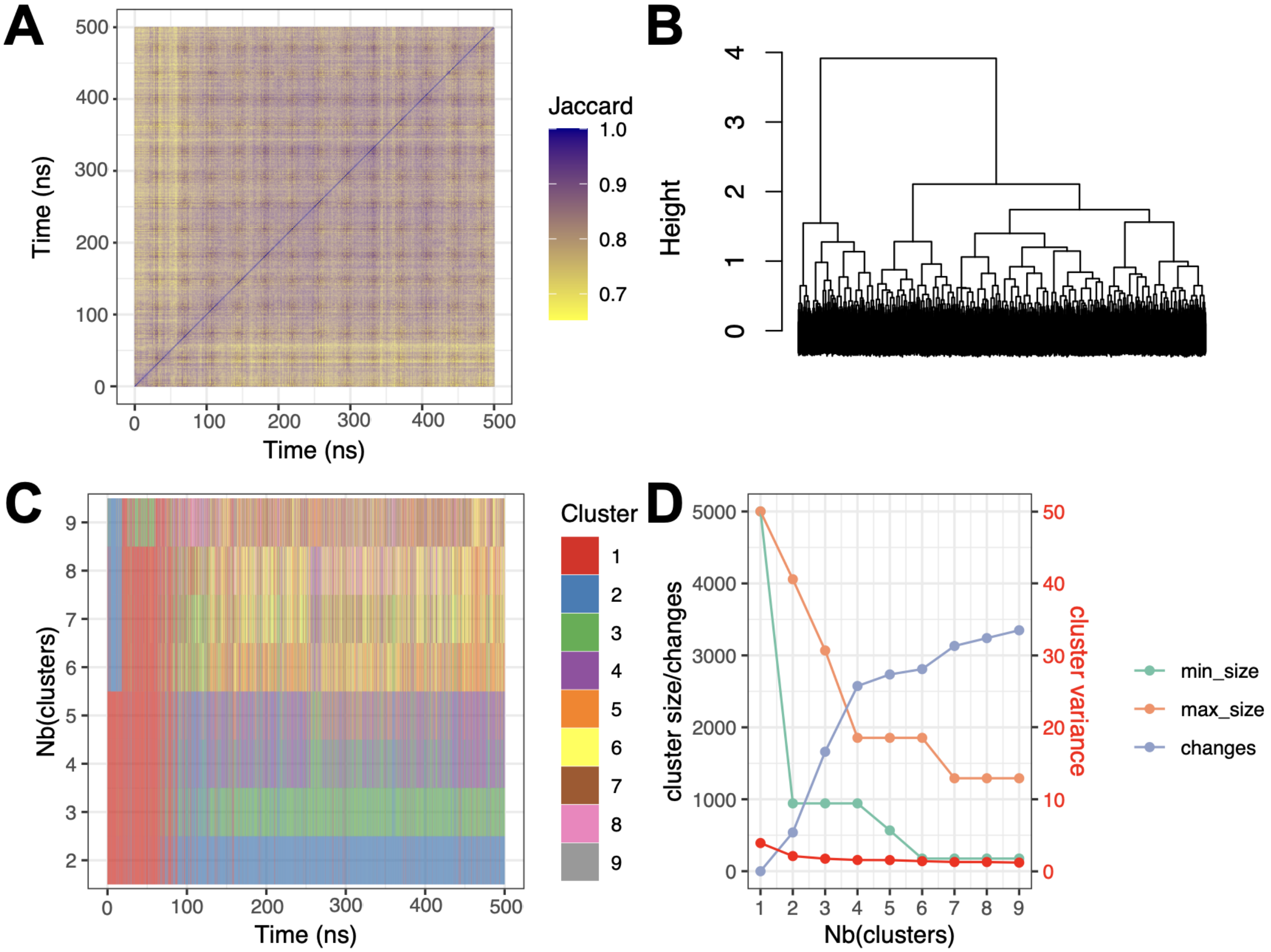


Figure S37: Interface clustering results for complex 3IEV (1 cluster). A. Jaccard similarity matrix. For each pair of snapshots, a yellow pixel indicates low interface similarity and a purple pixel indicates high interface similarity. B. Clustering dendrogram. C. Cluster membership along simulation time, for different numbers of clusters. D. Cluster size, number of changes, and intra-cluster variance (red), for different numbers of clusters. Min size: size of the smallest cluster, max size: size of the largest cluster, changes: number of cluster changes during the simulation.

### S11 Variance of contact frequencies across clusters

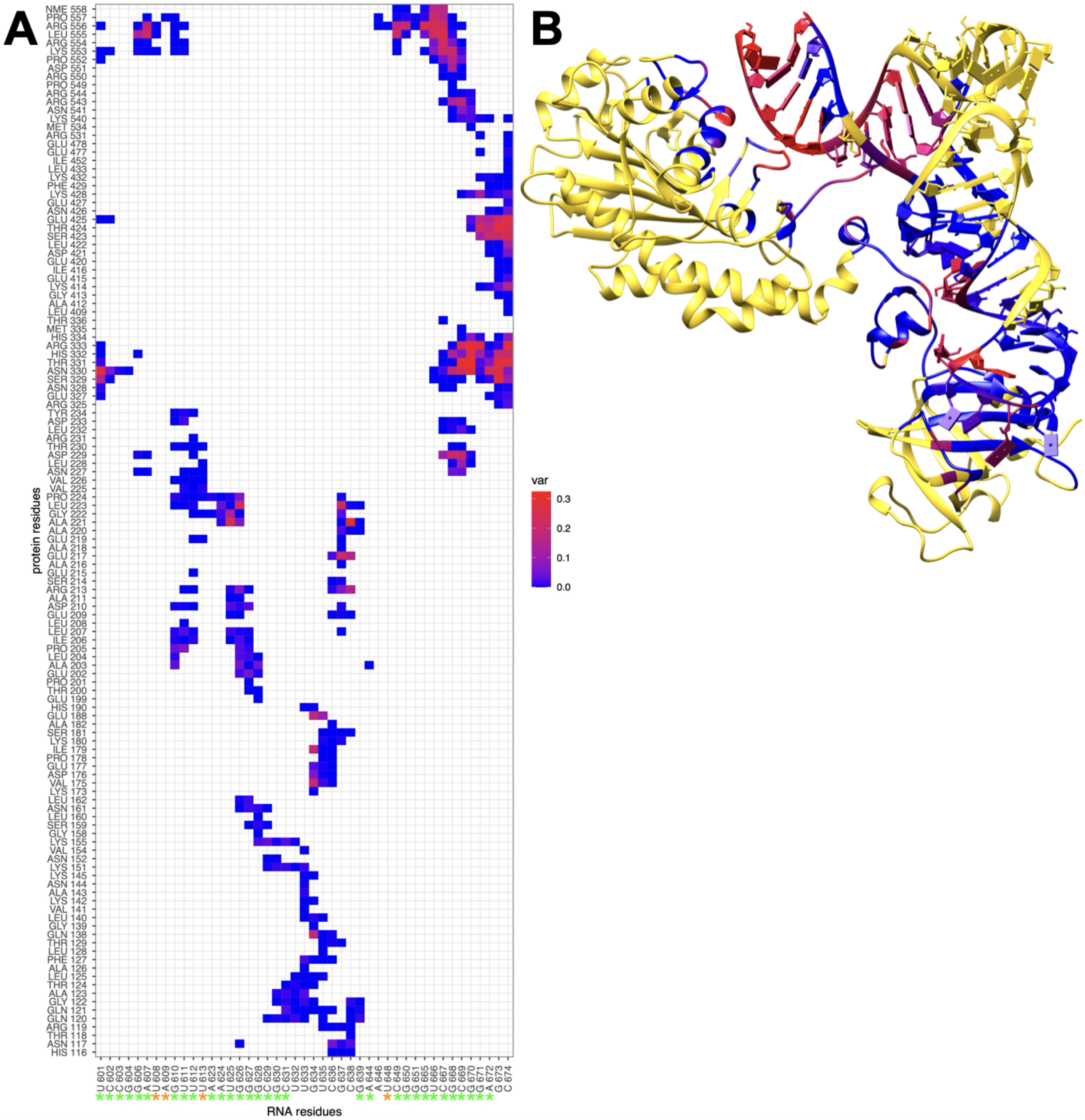

Figure S38: Contact variance in complex 1ASY. A. Matrix representation of each contact variance across the three interface clusters. Blue: low variance; Red: high variance; Green stars: canonical pairing; Orange stars: non-canonical pairing. B. 3D structure representation of 1ASY complex. Interface residues are colored by the maximum of contact variance (blue to red) and the other residues are colored in yellow.

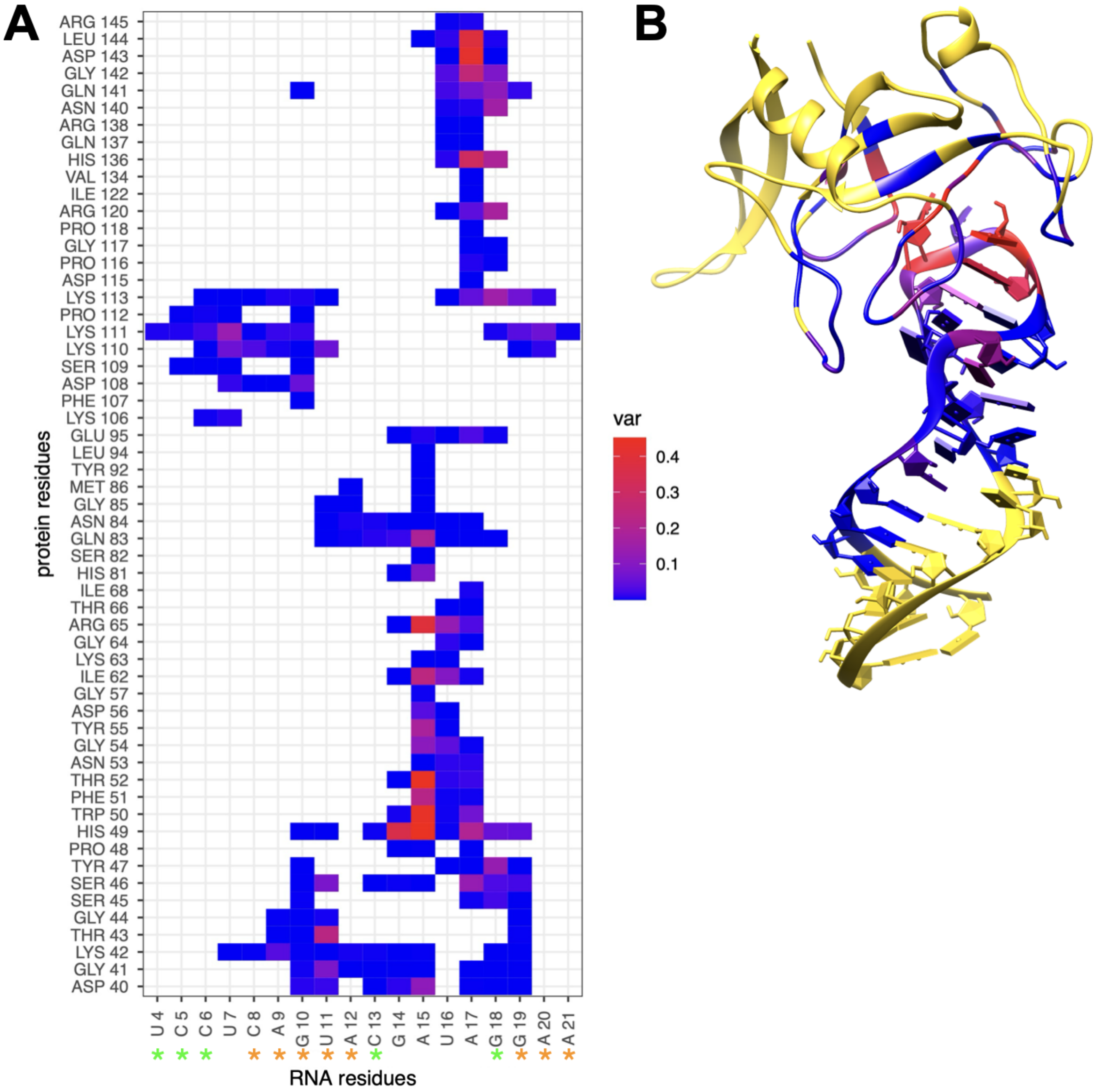


Figure S39: Contact variance in complex 1JBS. A. Matrix representation of each contact variance across the two interface clusters. Blue: low variance; Red: high variance; Green stars: canonical pairing; Orange stars: non-canonical pairing. B. 3D structure representation of 1JBS complex. Interface residues are colored by the maximum of contact variance (blue to red) and the other residues are colored in yellow.

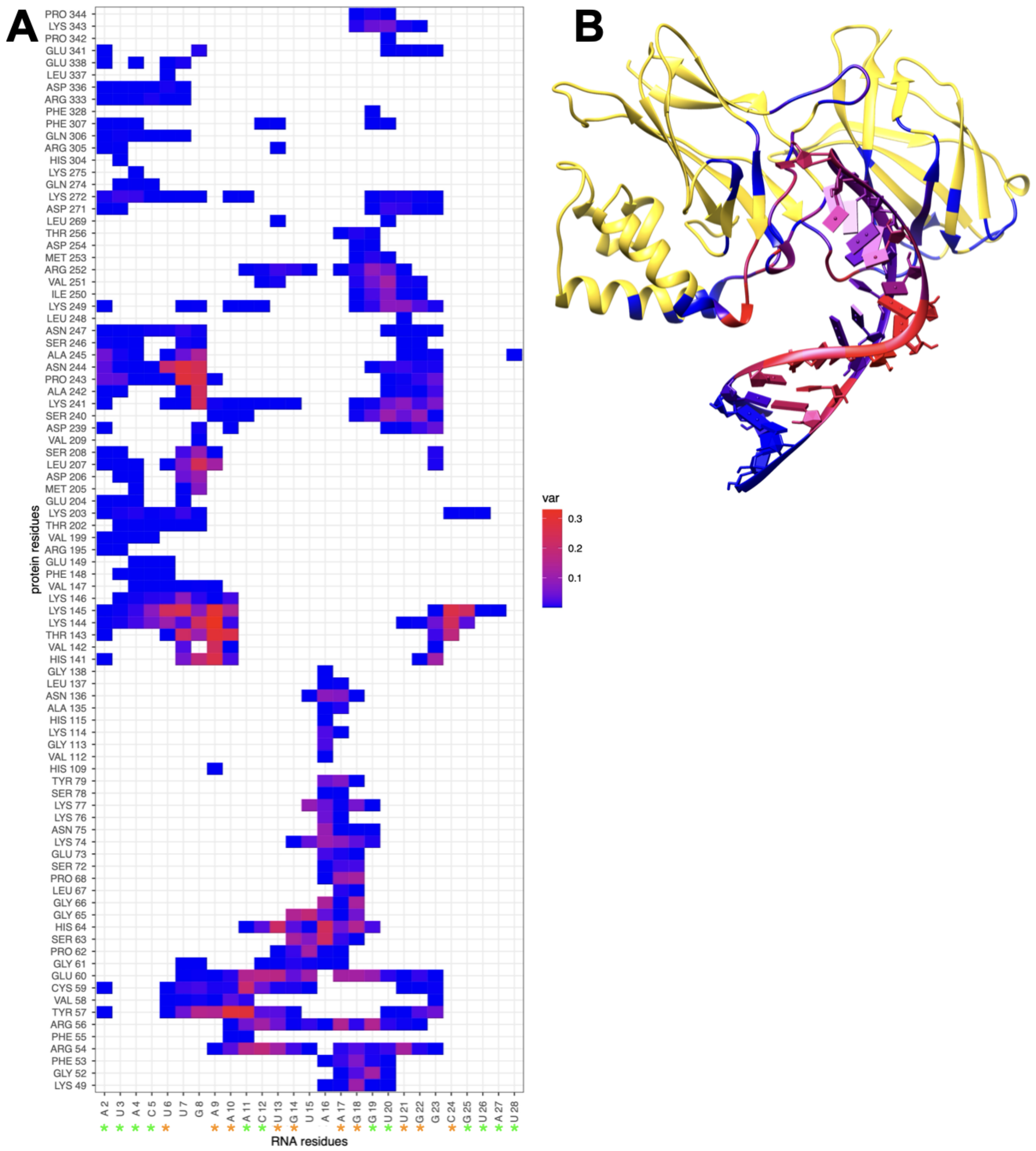


Figure S40: Contact variance in complex 1OOA. A. Matrix representation of each contact variance across the four interface clusters. Blue: low variance; Red: high variance; Green stars: canonical pairing; Orange stars: non-canonical pairing. B. 3D structure representation of 1OOA complex. Interface residues are colored by the maximum of contact variance (blue to red) and the other residues are colored in yellow.

### S12 Network

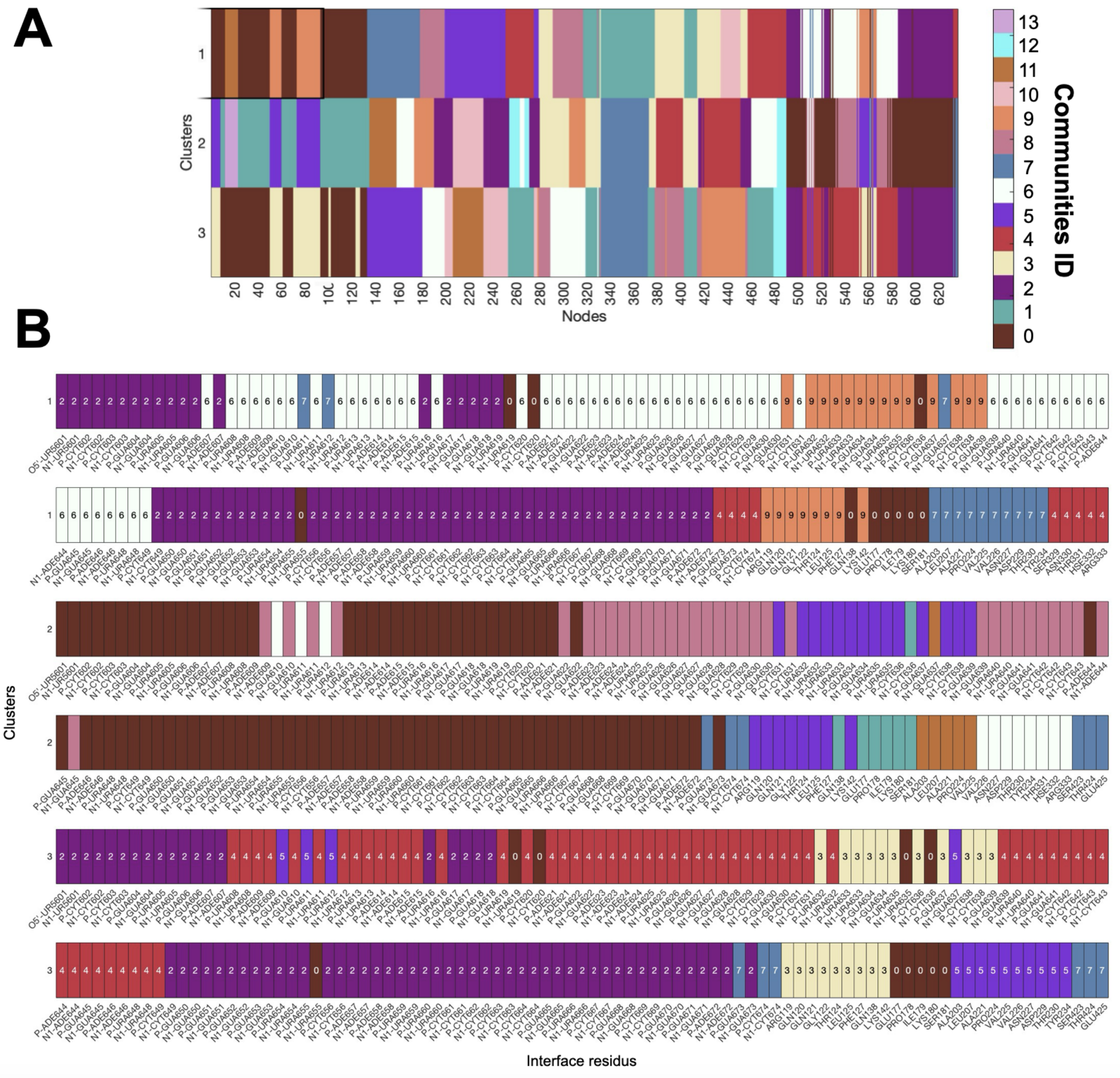


Figure S41: A. Community of the nodes for the different interface clusters of 1ASY complex. B. Community of the nodes for the different interface clusters of 1ASY complex for the residues at the interface between the protein and the RNA.

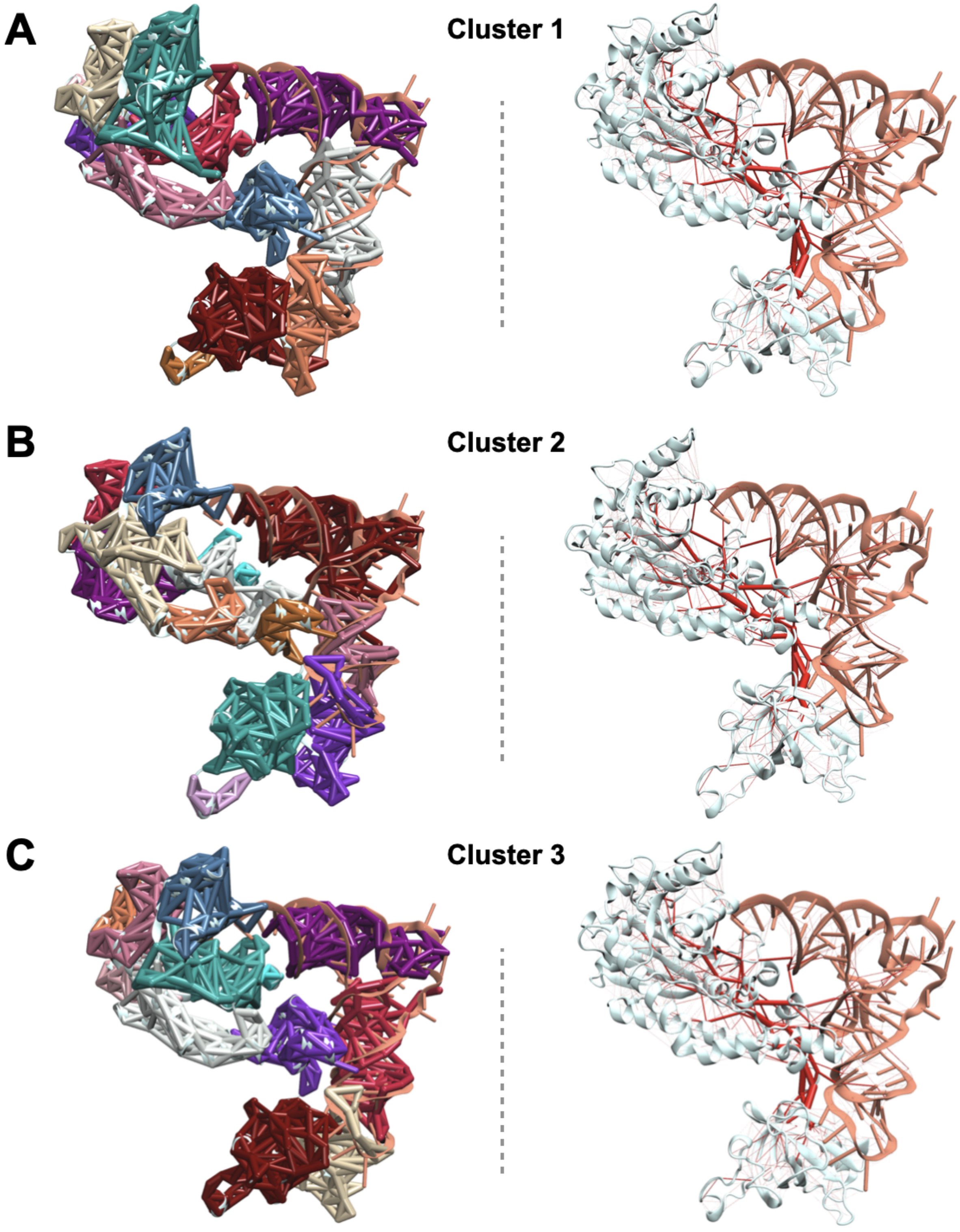


Figure S42: Communities representation in the complex structure (left) and edge betweenness (right) for the interface cluster 1 (A), 2 (B) and 3 (C) of 1ASY complex. Thickness depends on the betweenneess value.

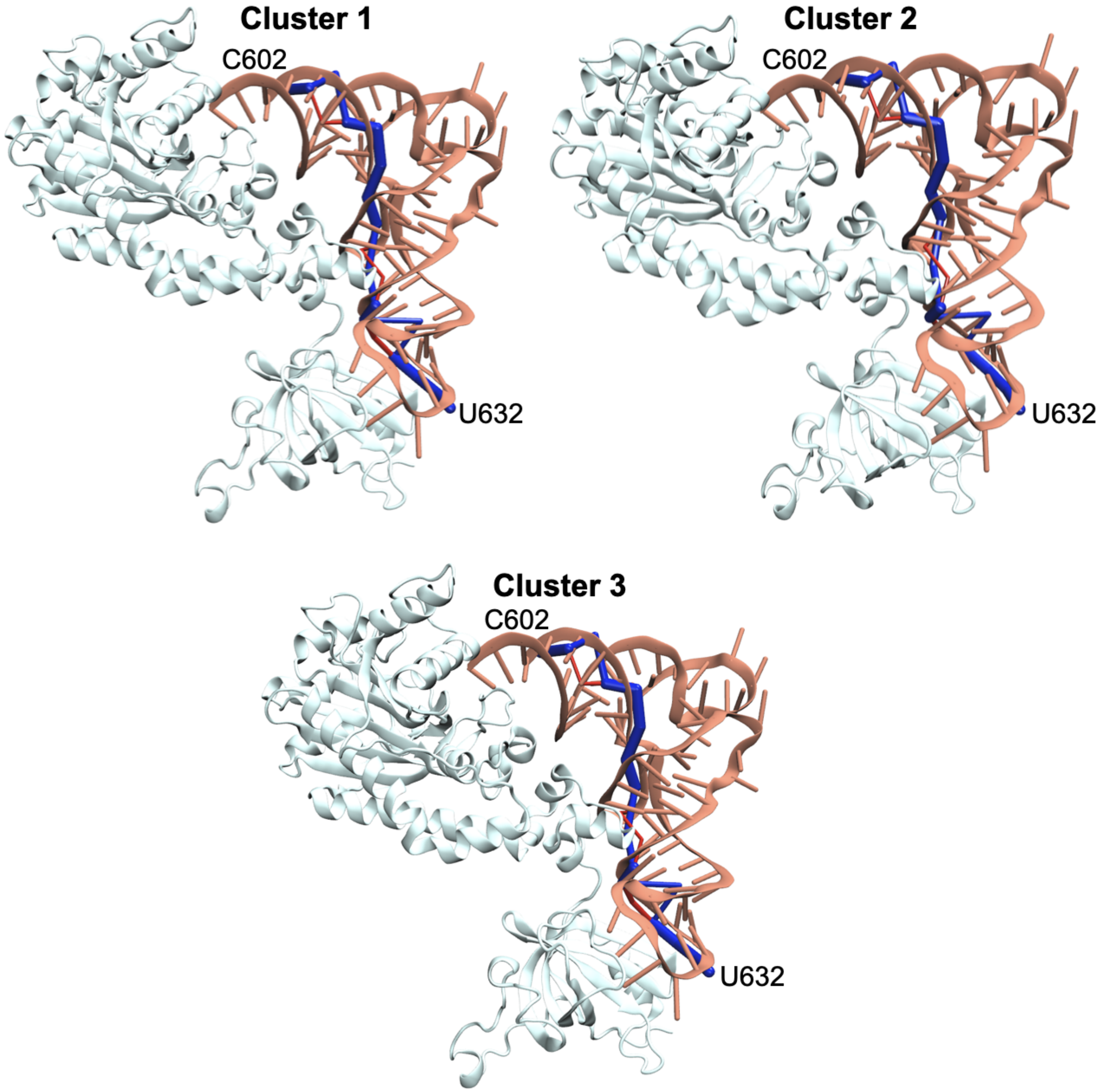


Figure S43: Optimal path (blue) and sub-optimal path (red) from U632 to C602 for cluster 1, 2 and 3 of 1ASY complex.

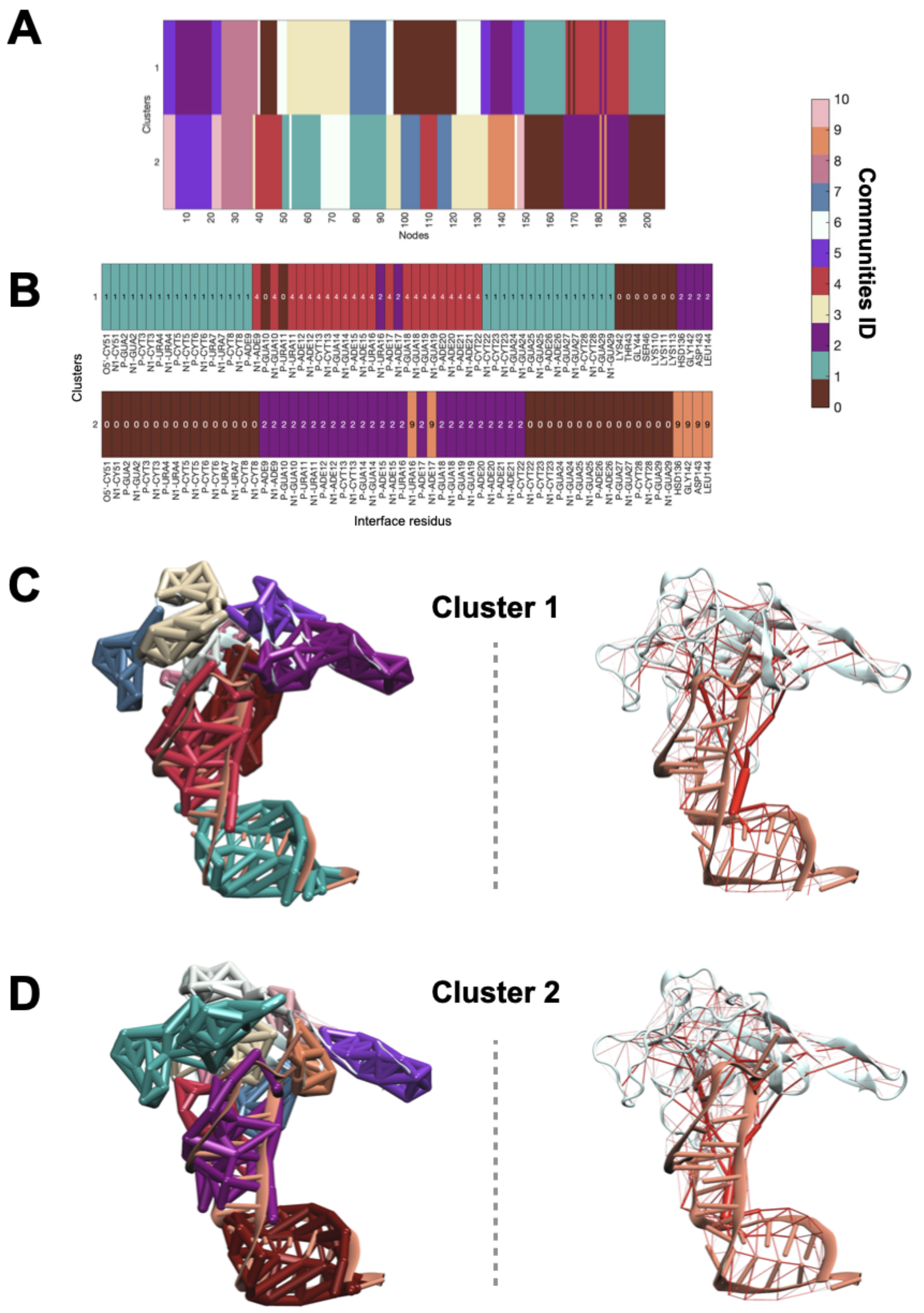

Figure S44: A. Community of the nodes for the different interface clusters of 1JBS complex. B. Community of the nodes for the different interface clusters of 1JBS complex for the residues at the interface between the protein and the RNA. Communities representation in the complex structure (left) and edge betweenness (right) for the interface cluster 1 (C) and 2 (D) of 1JBS complex. Thickness depends on the betweenneess value.

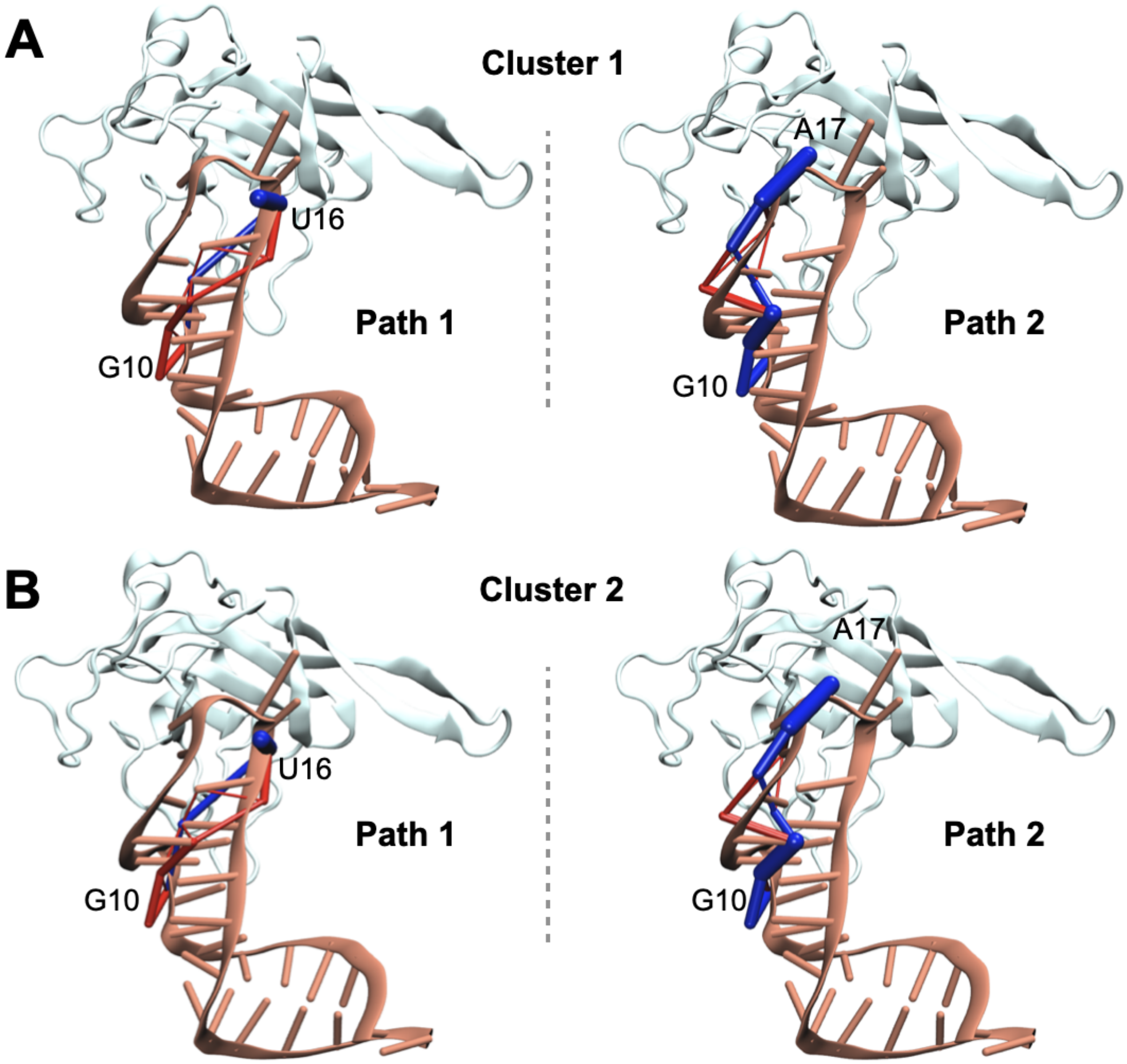


Figure S45: Optimal path (blue) and sub-optimal path (red) from G10 to U16 (left) and from G10 to A17 (right) for cluster 1 (A) and 2 (B) of 1JBS complex.

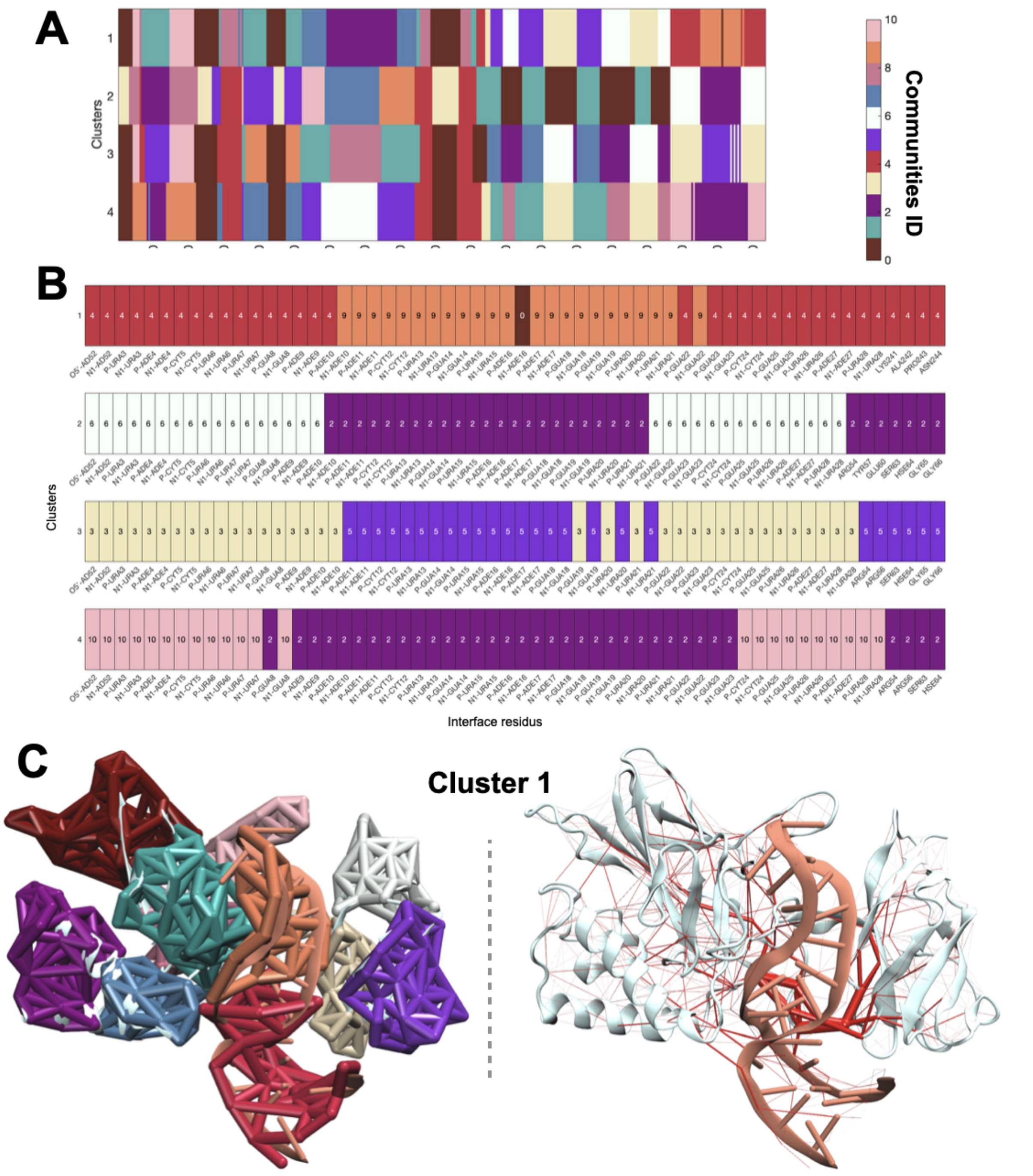


Figure S46: A. Community of the nodes for the different interface clusters of 1OOA complex. B. Community of the nodes for the different interface clusters of 1OOA complex for the residues at the interface between the protein and the RNA. Communities representation in the complex structure (left) and edge betweenness (right) for the interface cluster 1 (C) of 1OOA complex. Thickness depends on the betweeneess value.

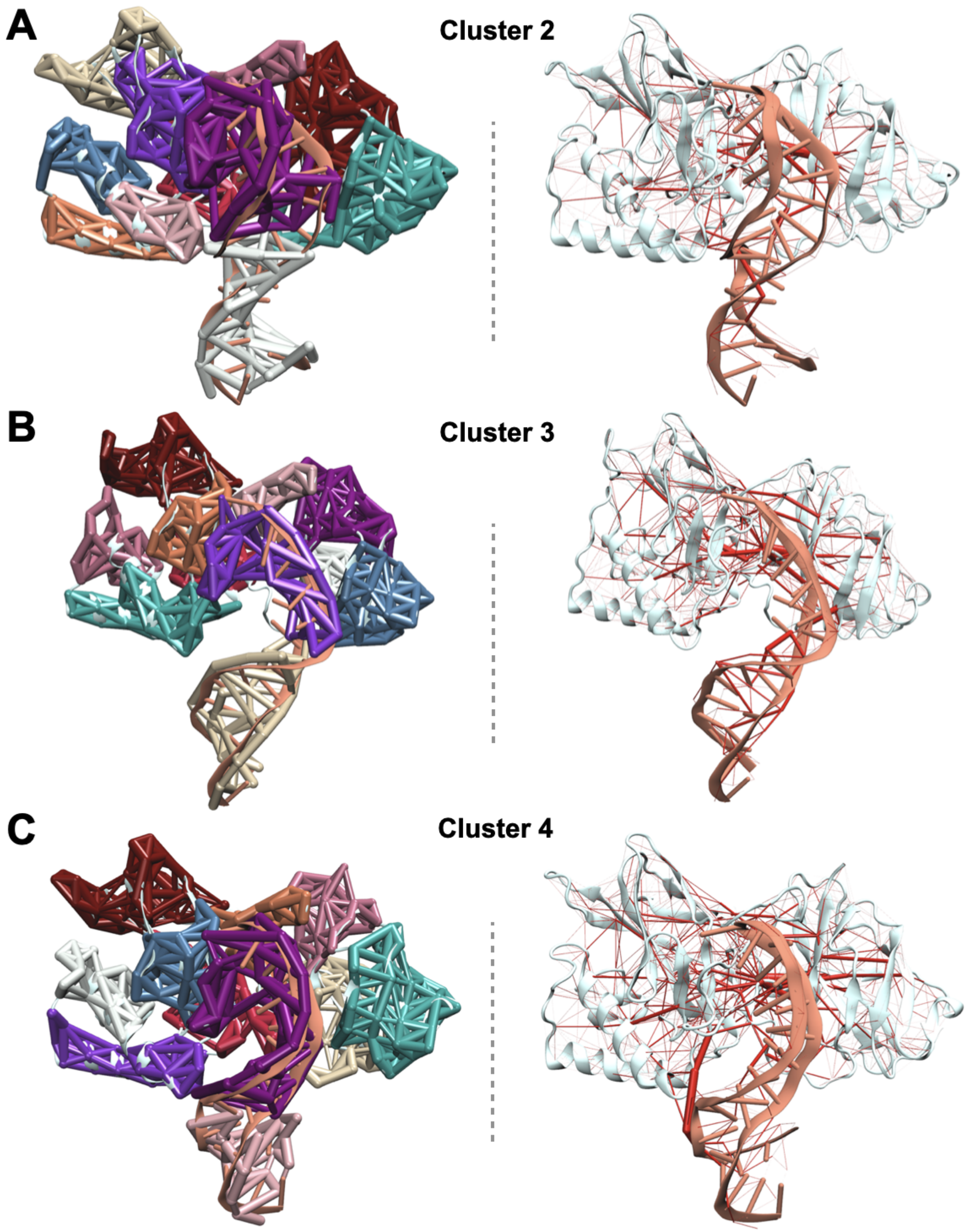


Figure S47: Communities representation in the complex structure (left) and edge betweenness (right) for the interface cluster 2 (A), 3 (B) and 4 (C) of 1OOA complex. Thickness depends on the betweenneess value.

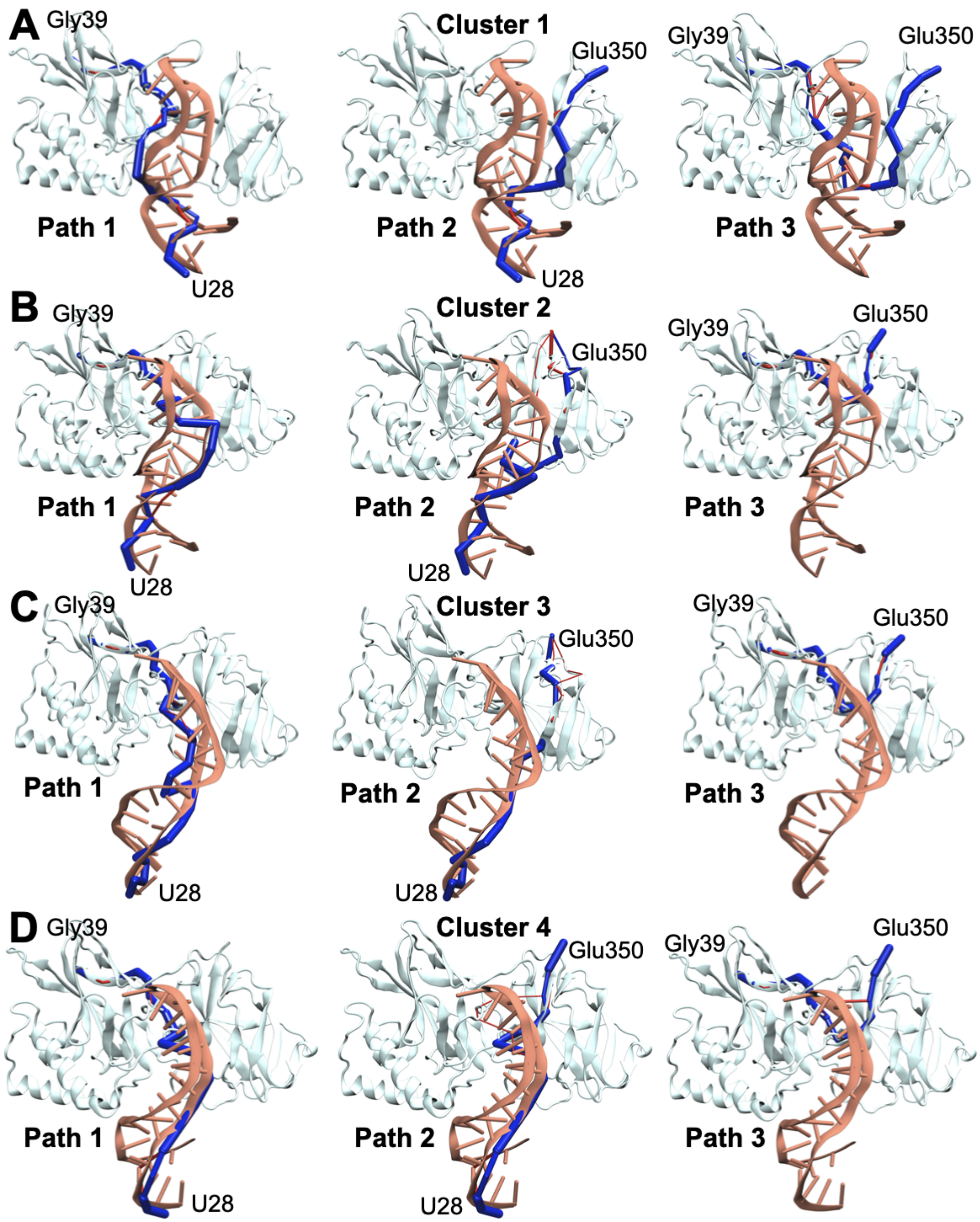


Figure S48: Optimal path (blue) and sub-optimal path (red) from U28 to Gly39 (left), U28 to Glu350 (middle) and from G10 to A17 (right) for cluster 1 (A), 2 (B), 3 (C) and 4 (D) of 1OOA complex.

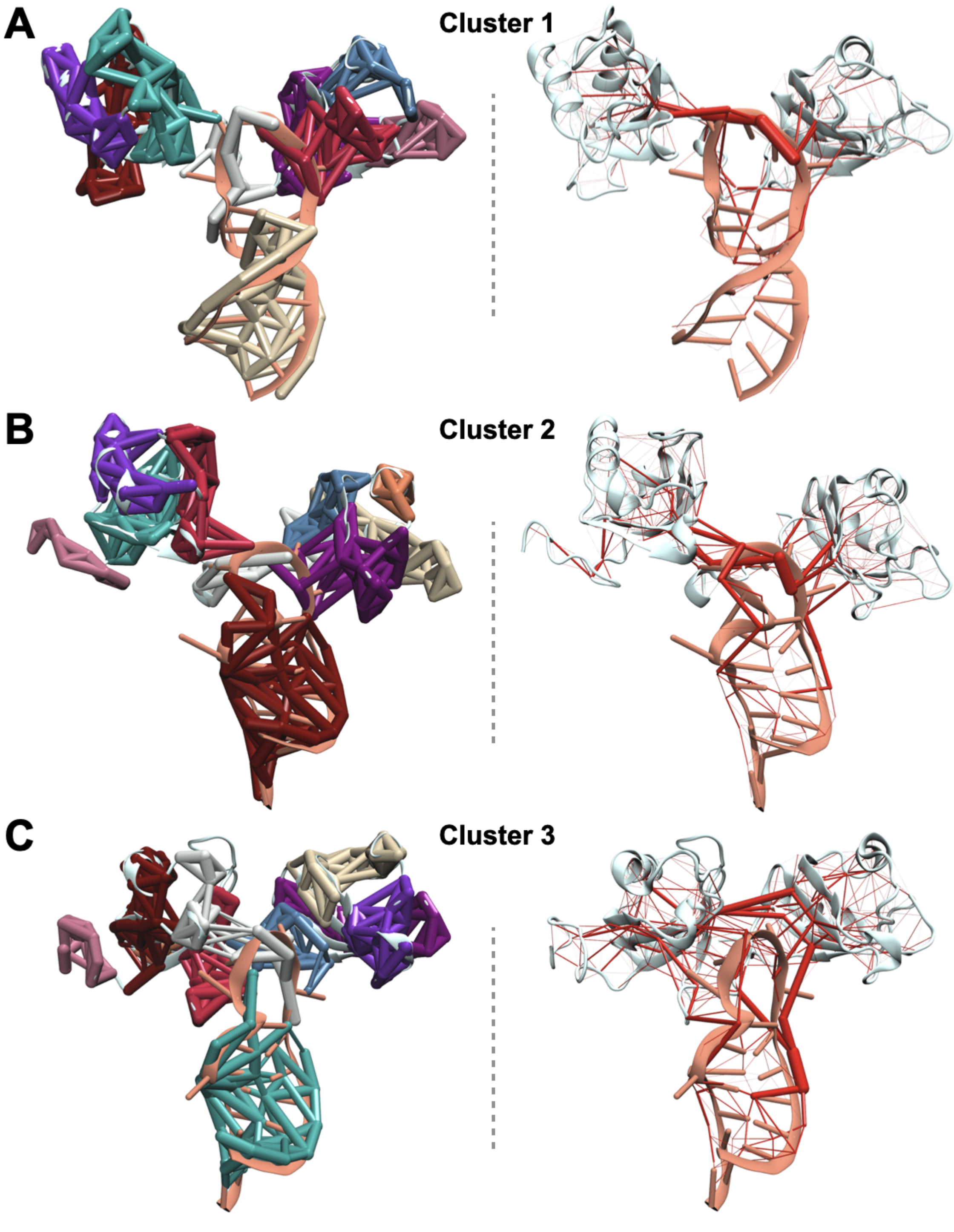


Figure S49: Communities representation in the complex structure (left) and edge betweenness (right) for the interface cluster 1 (A), 2 (B) and 3 (C) of 1RKJ complex. Thickness depends on the betweenness value.

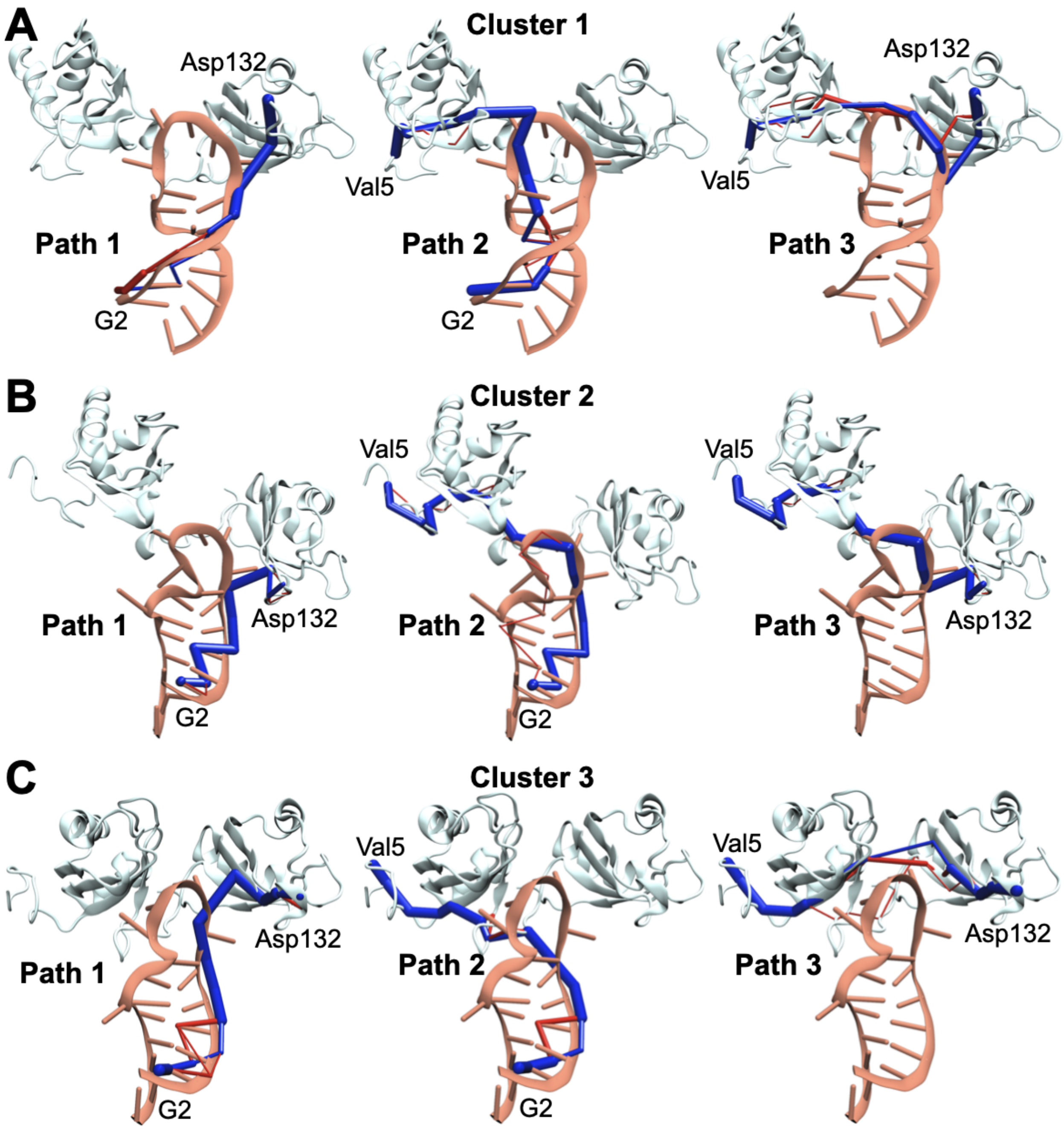


Figure S50: Optimal path (blue) and sub-optimal path (red) from G2 to Asp132 (left), G2 to Val5 (middle) and from Val5 to Asp132 (right) for cluster 1 (A), 2 (B) and 3 (C) of 1RKJ complex.

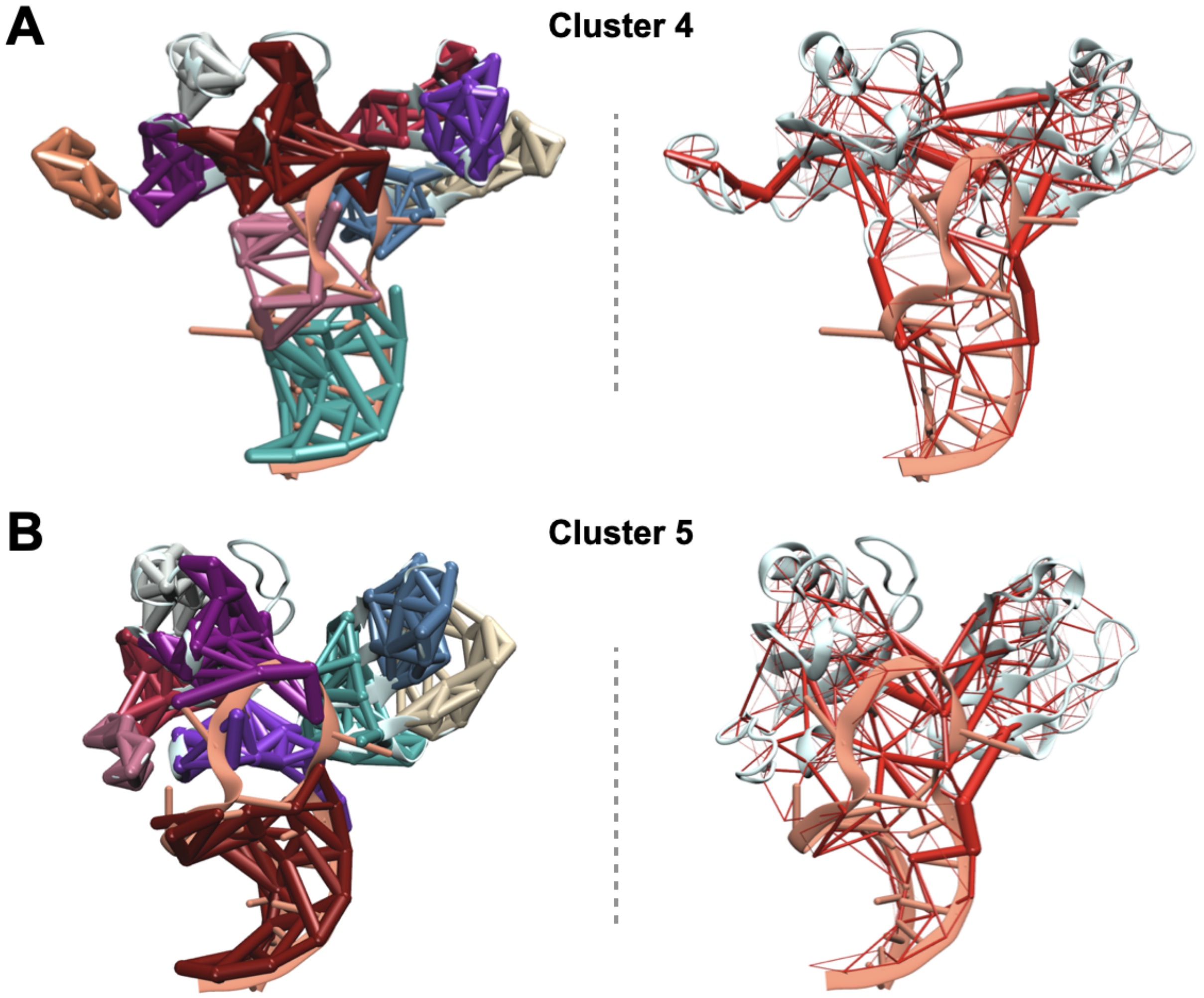


Figure S51: Communities representation in the complex structure (left) and edge betweenness (right) for the interface cluster 4 (A) and 5 (B) of 1RKJ complex. Thickness depends on the betweenneess value.

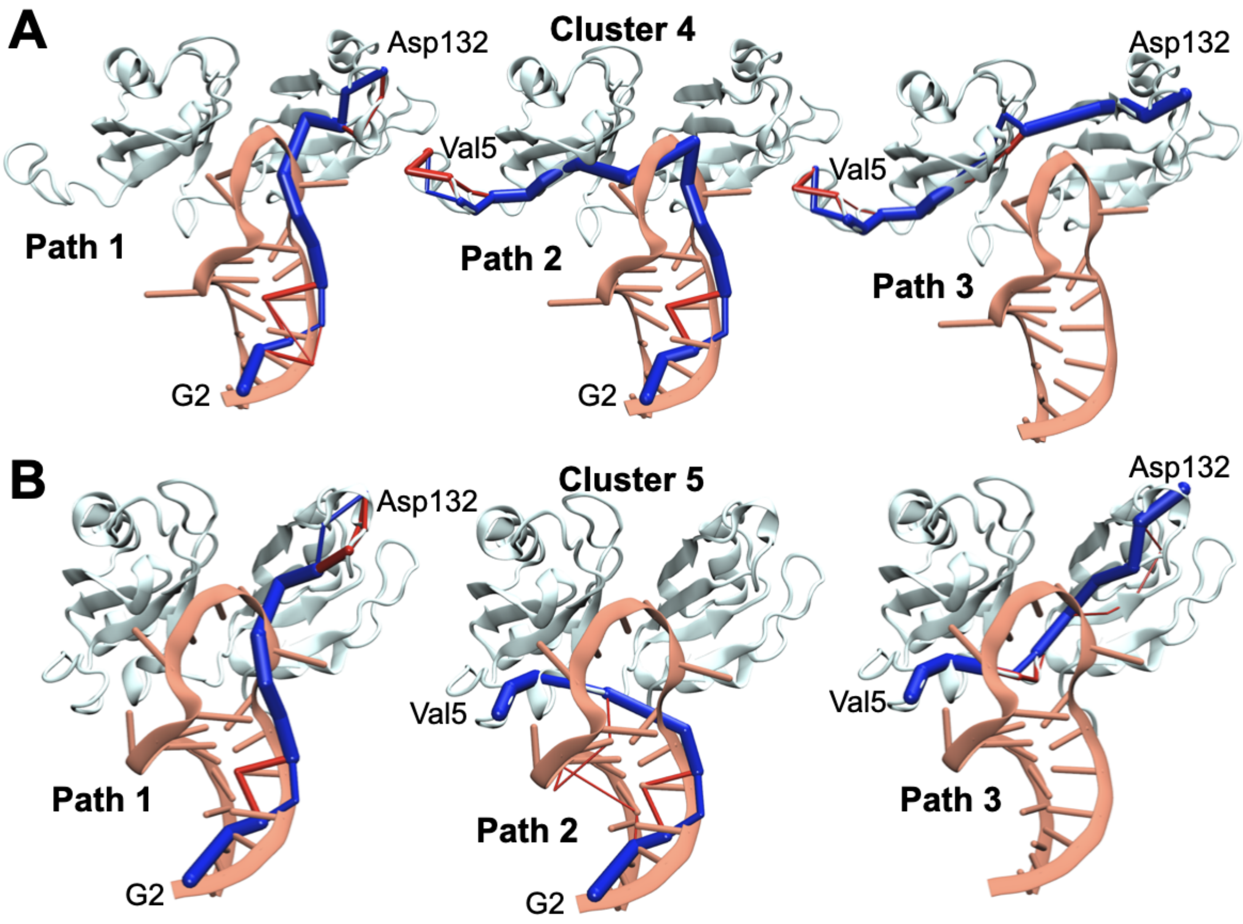


Figure S52: Optimal path (blue) and sub-optimal path (red) from G2 to Asp132 (left), G2 to Val5 (middle) and from Val5 to Asp132 (right) for cluster 4 (A) and 5 (B) of 1RKJ complex.

#### S13 Puckering

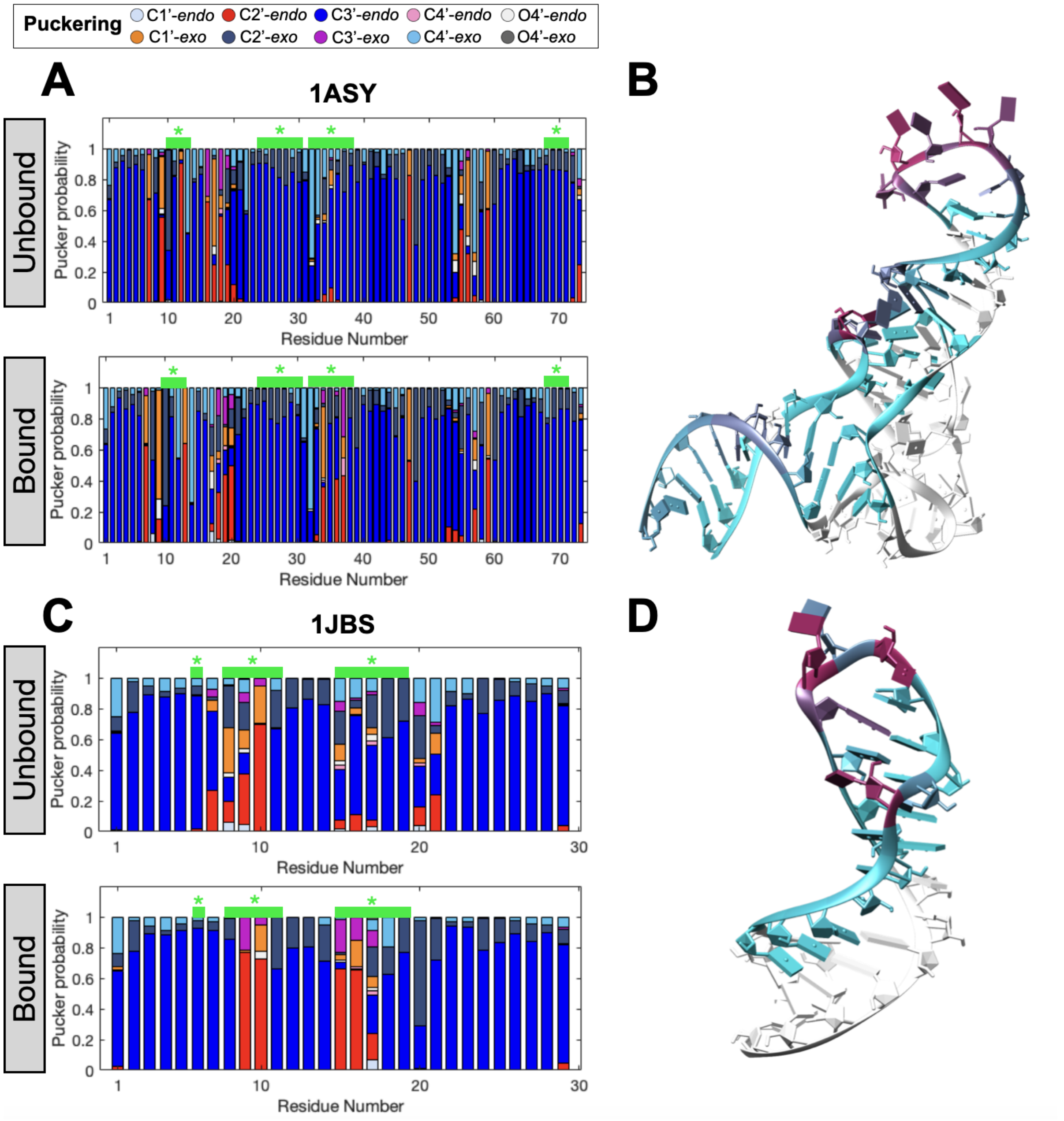


Figure S53: A-C. Pucker probability of RNA unbound (top) and bound (bottom) for 1ASY (A) and 1JBS (C) represented with stacked bar plot. Green bar and stars highlight the RNA residues involved in contacts with protein, with a frequency higher than or equal to 0.75. B-like family: Red: C2'-*endo*, Pink: C4'-*endo*, Orange red: C1'-*exo*, Magenta: C3'-*exo*. A-like family: Blue: C3'- endo, light blue: C1'-*endo*, dark blue: C2'-*exo*, cyan: C4'-*exo*. Light grey: O4'-*endo*, dark grey: O4'-*exo*. B-D. RNA structure colored by interface contact for 1ASY (B) and 1JBS (D). White: not interface residues, cyan: residues at interface with a low frequency and pink: residues at interface with high frequency.

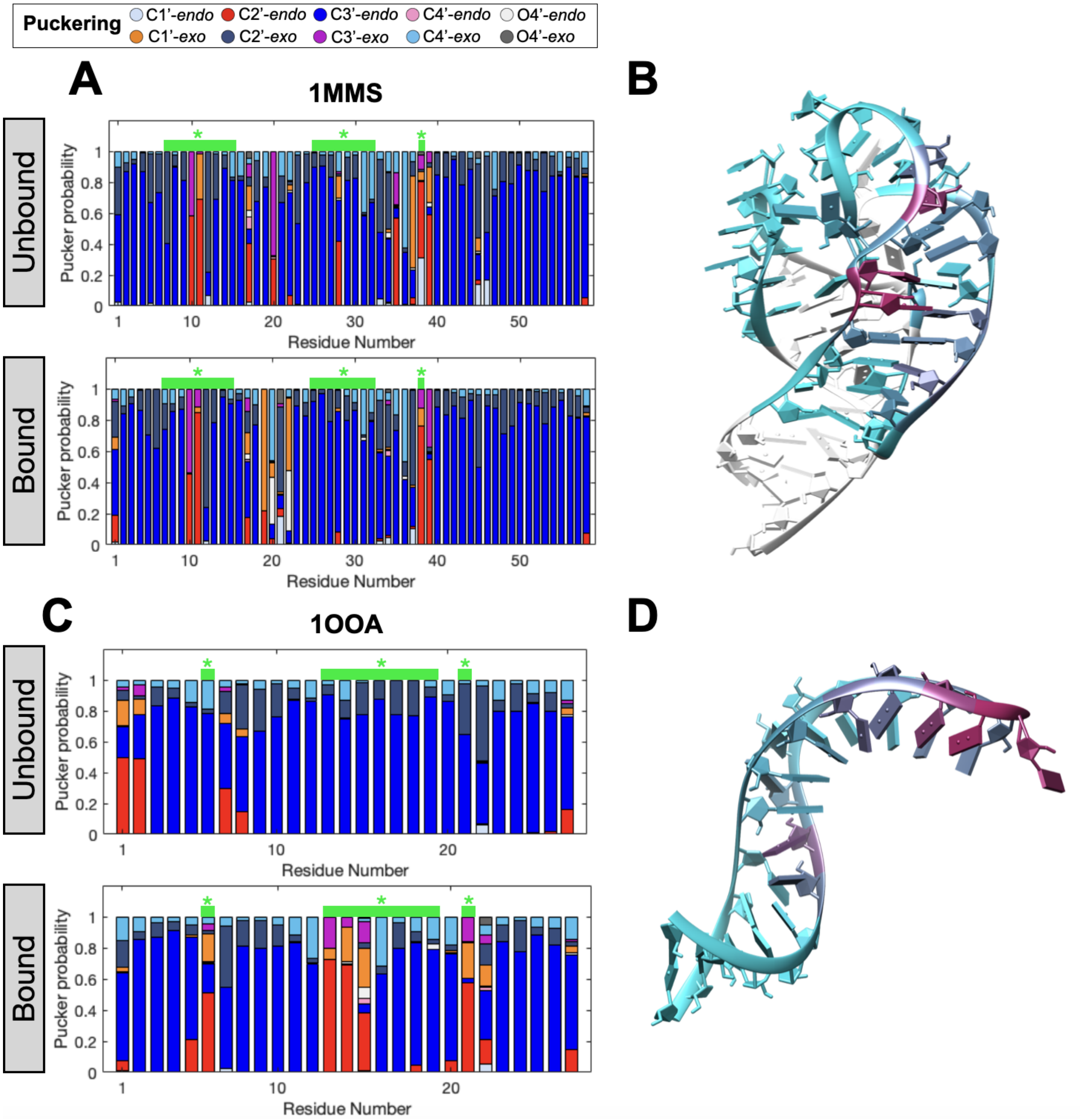


Figure S54: A-C. Pucker probability of RNA unbound (top) and bound (bottom) for 1MMS (A) and 1OOA (C) represented with stacked bar plot. Green bar and stars highlight the RNA residues involved in contacts with protein, with a frequency higher than or equal to 0.75. B-like family: Red: C2'-*endo*, Pink: C4'-*endo*, Orange red: C1'-*exo*, Magenta: C3'-*exo*. A-like family: Blue: C3'- endo, light blue: C1'-*endo*, dark blue: C2'-*exo*, cyan: C4'-*exo*. Light grey: O4'-*endo*, dark grey: O4'-*exo*. B-D. RNA structure colored by interface contact for 1MMS (B) and 1OOA (D). White: not interface residues, cyan: residues at interface with a low frequency and pink: residues at interface with high frequency.

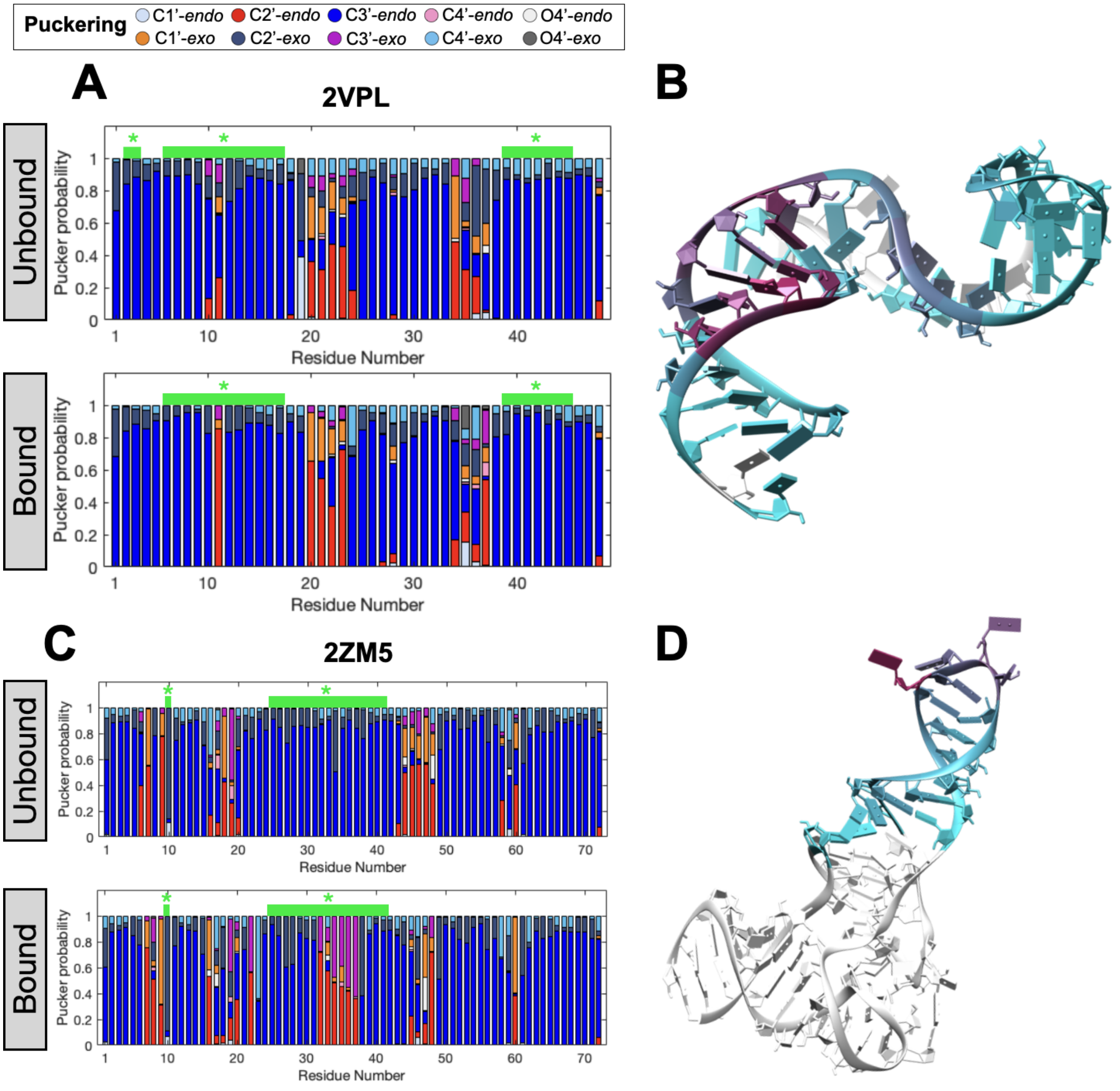


Figure S55: A-C. Pucker probability of RNA unbound (top) and bound (bottom) for 2VPL (A) and 2ZM5 (C) represented with stacked bar plot. Green bar and stars highlight the RNA residues involved in contacts with protein, with a frequency higher than or equal to 0.75. B-like family: Red: C2'-*endo*, Pink: C4'-*endo*, Orange red: C1'-*exo*, Magenta: C3'-*exo*. A-like family: Blue: C3'- endo, light blue: C1'-*endo*, dark blue: C2'-*exo*, cyan: C4'-*exo*. Light grey: O4'-*endo*, dark grey: O4'-*exo*. B-D. RNA structure colored by interface contact for 2VPL (B) and 2ZM5 (D). White: not interface residues, cyan: residues at interface with a low frequency and pink: residues at interface with high frequency.

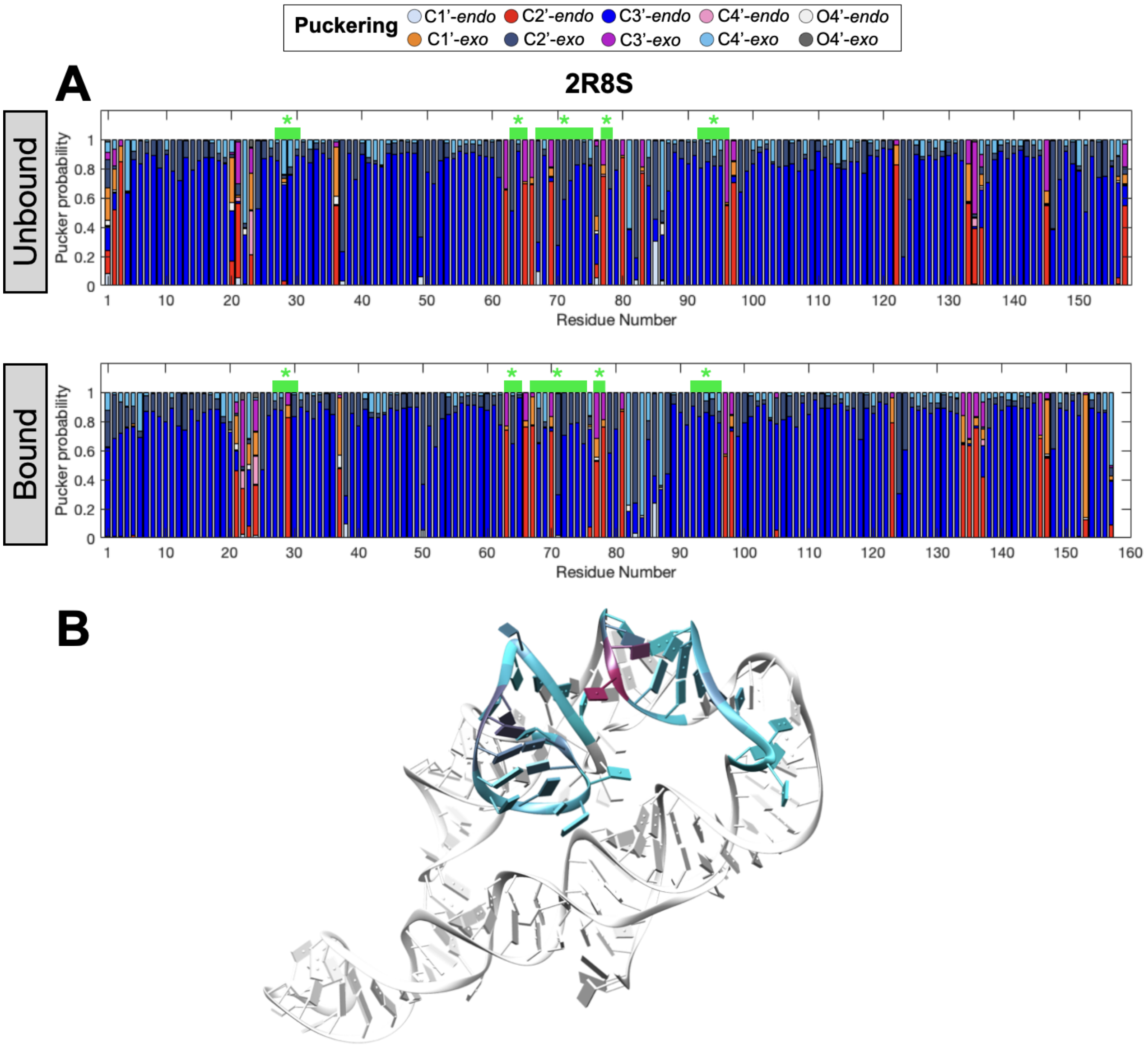


Figure S56: A. Pucker probability of RNA unbound (top) and bound (bottom) for 2R8S represented with stacked bar plot. Green bar and stars highlight the RNA residues involved in contacts with protein, with a frequency higher than or equal to 0.75. B-like family: Red: C2'-*endo*, Pink: C4'-*endo*, Orange red: C1'-*exo*, Magenta: C3'-*exo*. A-like family: Blue: C3'- endo, light blue: C1'-*endo*, dark blue: C2'-*exo*, cyan: C4'-*exo*. Light grey: O4'-*endo*, dark grey: O4'-*exo*. B. RNA structure colored by interface contact for 2R8S. White: not interface residues, cyan: residues at interface with a low frequency and pink: residues at interface with high frequency.

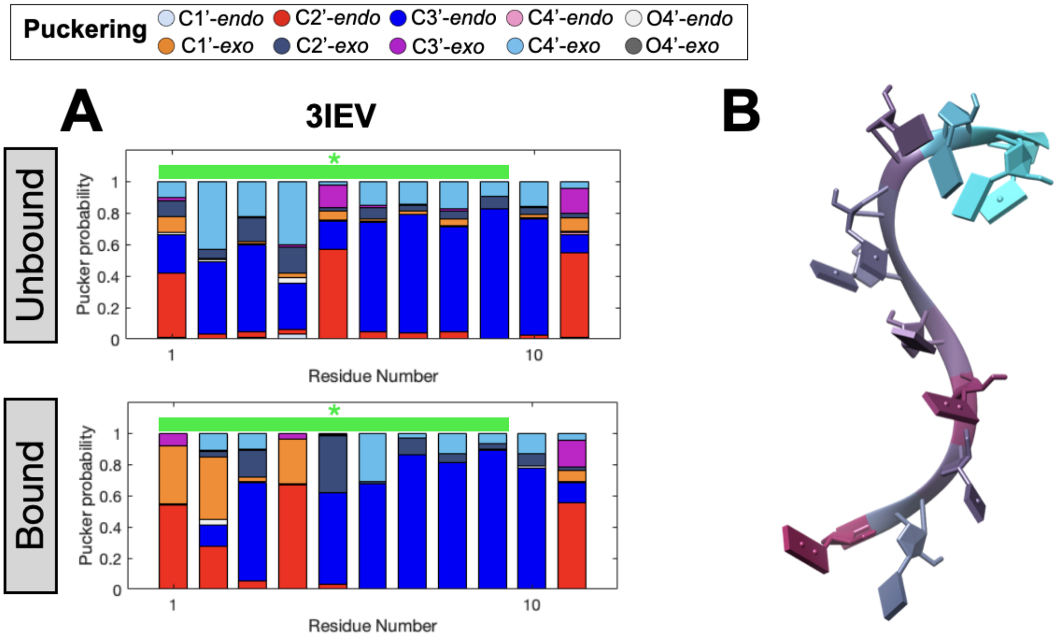


Figure S57: A. Pucker probability of RNA unbound (top) and bound (bottom) for 3IEV represented with stacked bar plot. Green bar and stars highlight the RNA residues involved in contacts with protein, with a frequency higher than or equal to 0.75. B-like family: Red: C2'-*endo*, Pink: C4'-*endo*, Orange red: C1'-*exo*, Magenta: C3'-*exo*. A-like family: Blue: C3'- endo, light blue: C1'-*endo*, dark blue: C2'-*exo*, cyan: C4'-*exo*. Light grey: O4'-*endo*, dark grey: O4'-*exo*. B. RNA structure colored by interface contact for 3IEV. White: not interface residues, cyan: residues at interface with a low frequency and pink: residues at interface with high frequency.

#### S14 Puckering clusters

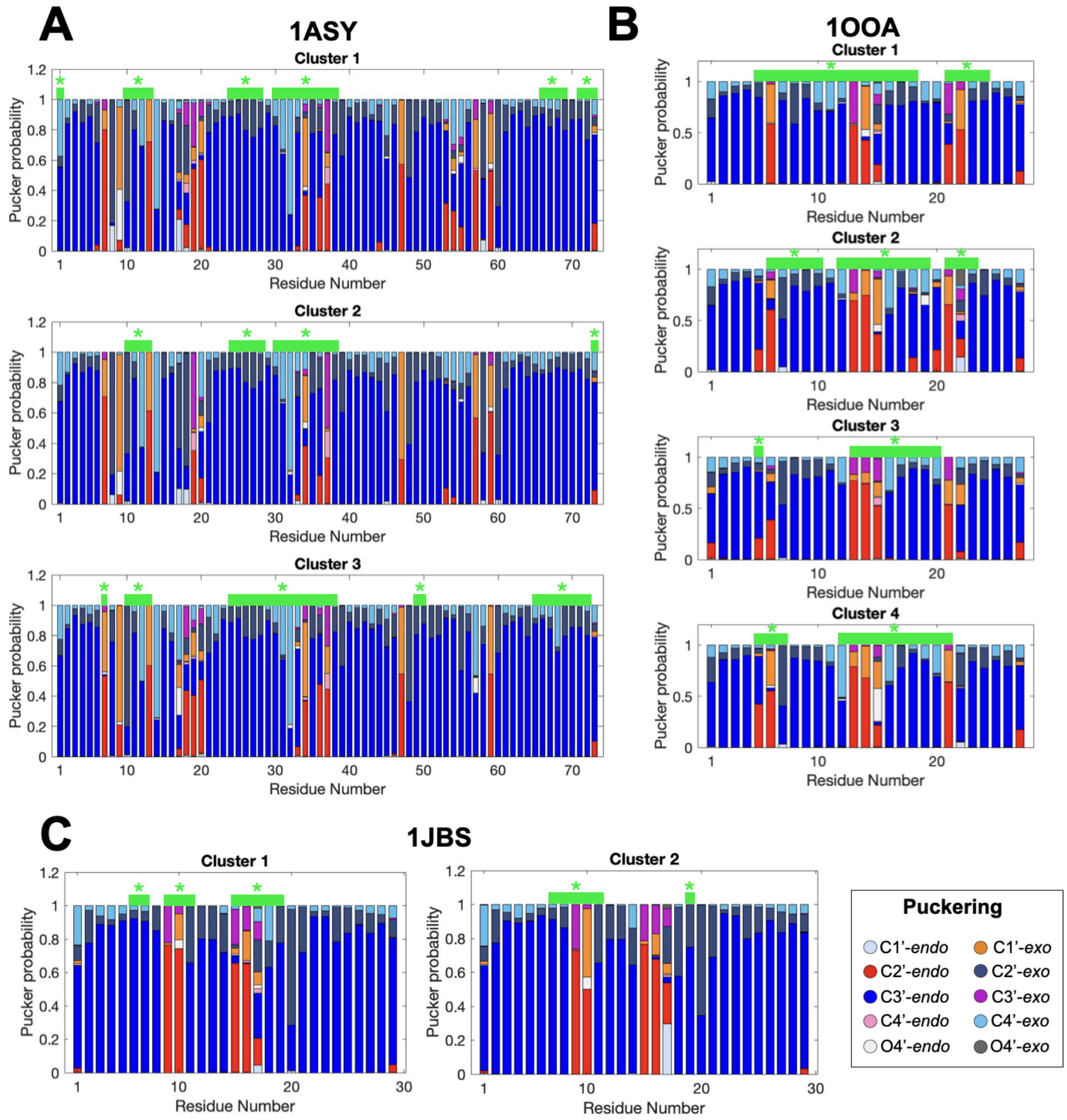


Figure S58: Pucker probability of RNA for each interface cluster of 1ASY (A), 1OOA (B) and 1JBS (C) complex represented with stacked bar plot. Green bar and stars highlight the RNA residues involved in contacts with protein, with a frequency higher than or equal to 0.75. B-like family: Red: C2'-*endo*, Pink: C4'-*endo*, Orange red: C1'-*exo*, Magenta: C3'-*exo*. A-like family: Blue: C3'- endo, light blue: C1'-*endo*, dark blue: C2'-*exo*, cyan: C4'-*exo*. Light grey: O4'-*endo*, dark grey: O4'-*exo*.

### S15 Interfacial water molecules evolution

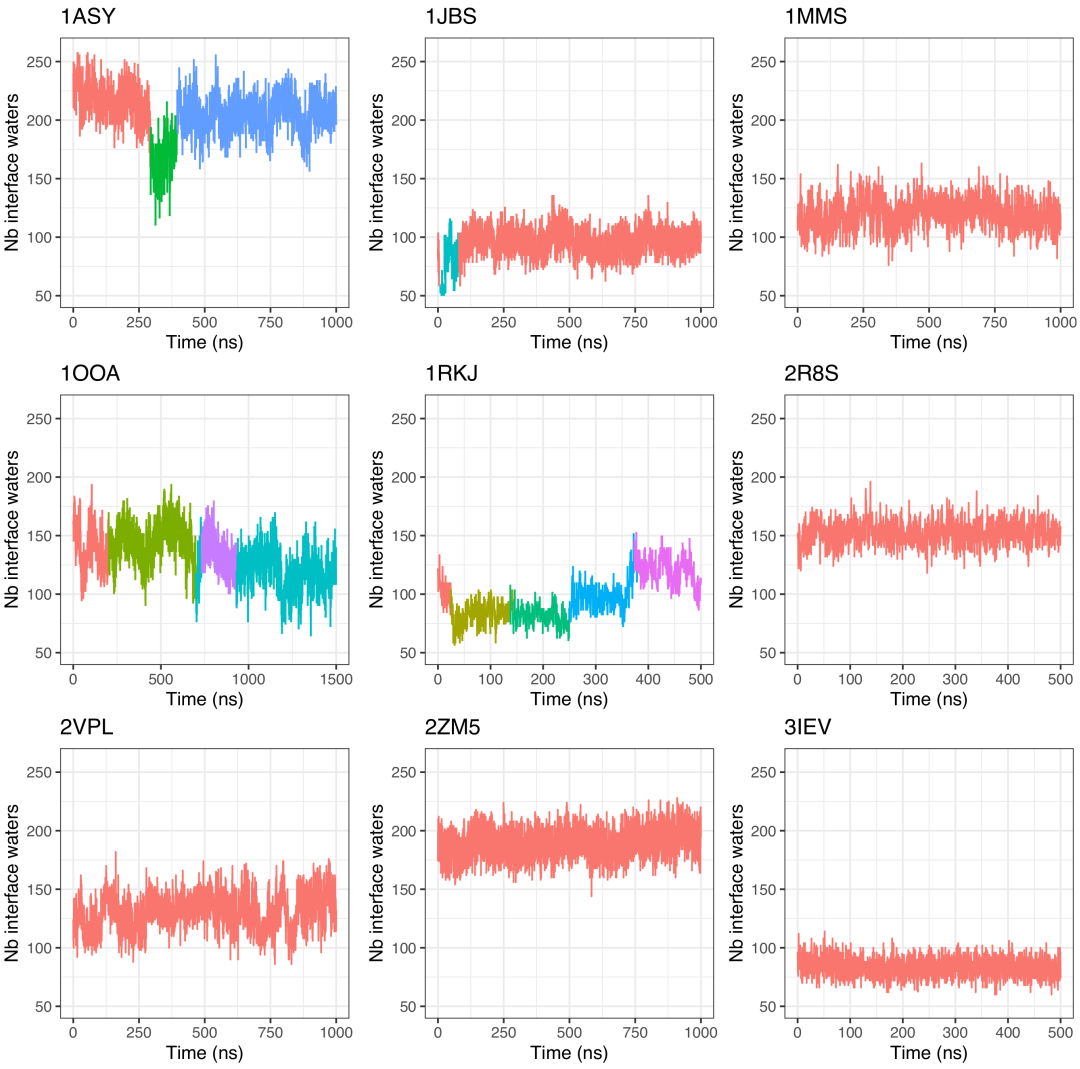


Figure S59: Number of interfacial water molecules in contact with interface residues, in each protein-RNA complexes.

### S16 Interfacial water molecules number

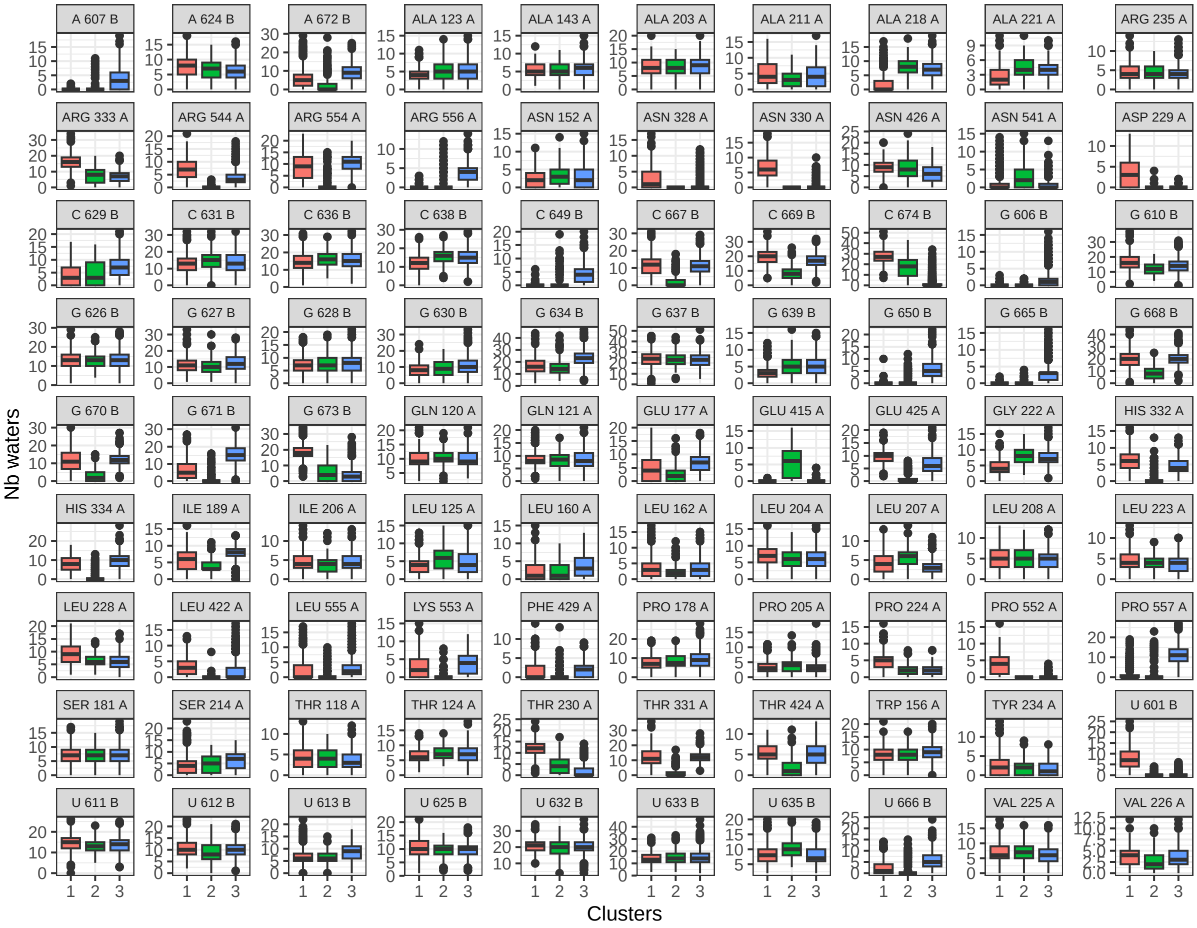


Figure S60: Number of interfacial water molecules in contact with interface residues, in each MD clusters for 1ASY complex.

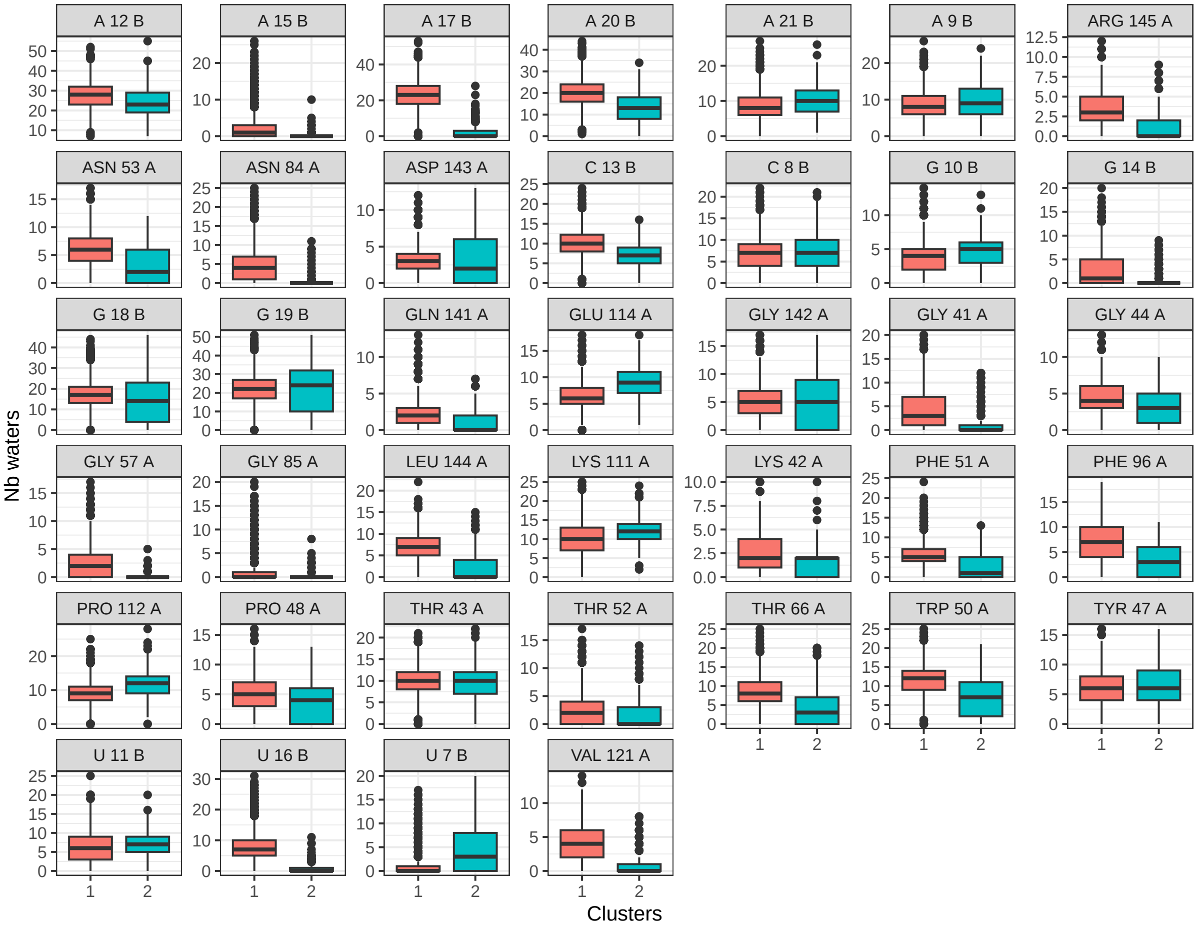


Figure S61: Number of interfacial water molecules in contact with interface residues, in each MD clusters for 1JBS complex.

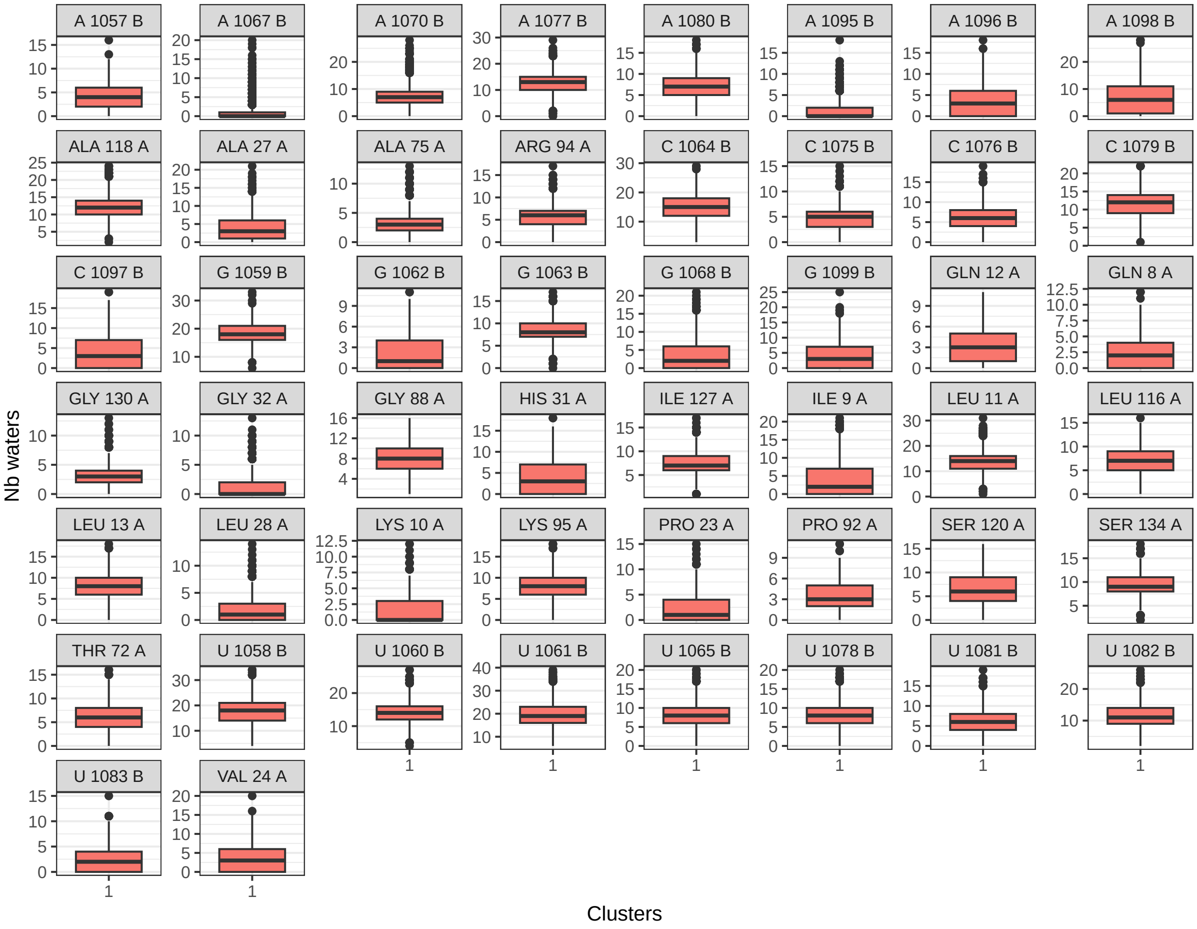


Figure S62: Number of interfacial water molecules in contact with interface residues, in each MD clusters for 1MMS complex.

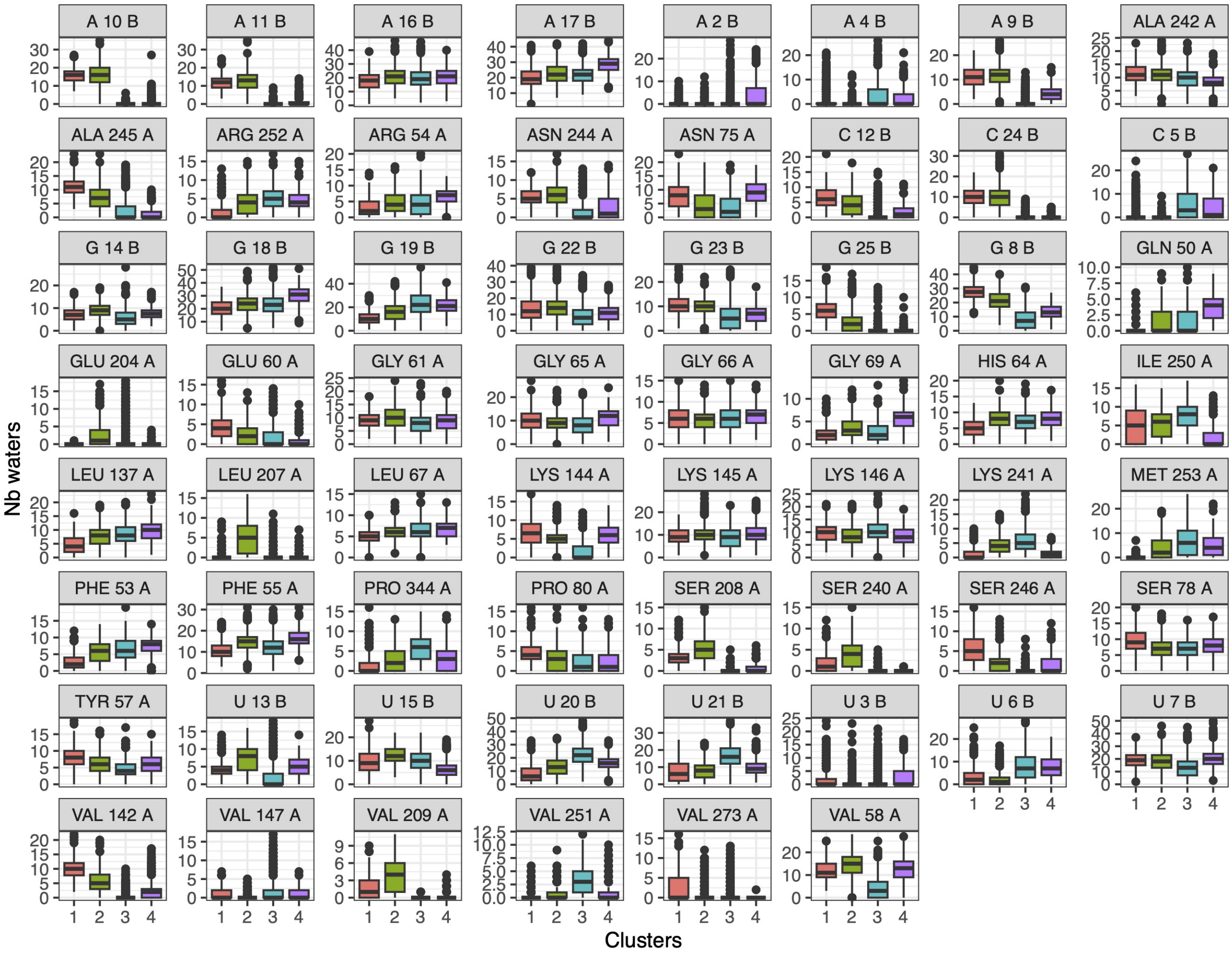


Figure S63: Number of interfacial water molecules in contact with interface residues, in each MD clusters for 1OOA complex.

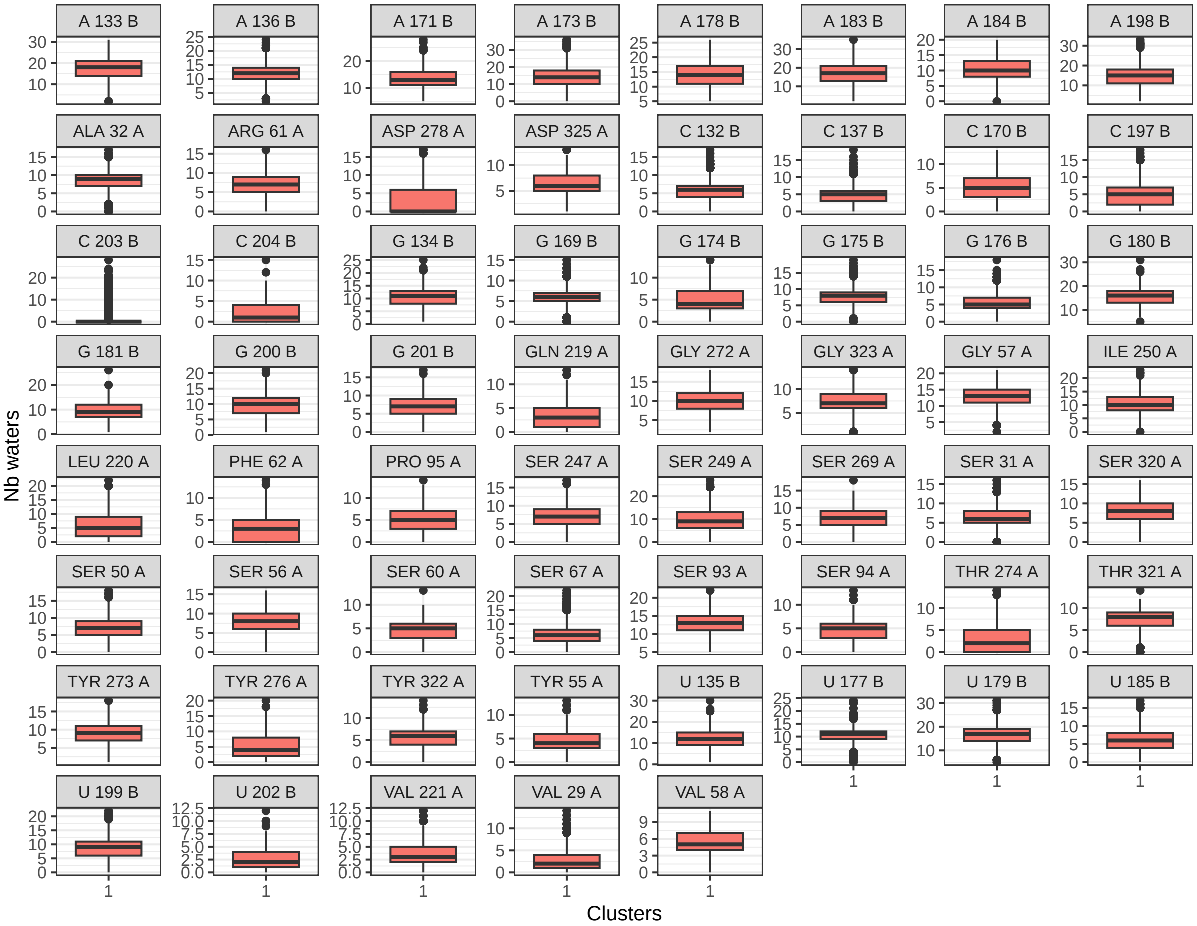


Figure S64: Number of interfacial water molecules in contact with interface residues, in each MD clusters for 2R8S complex.

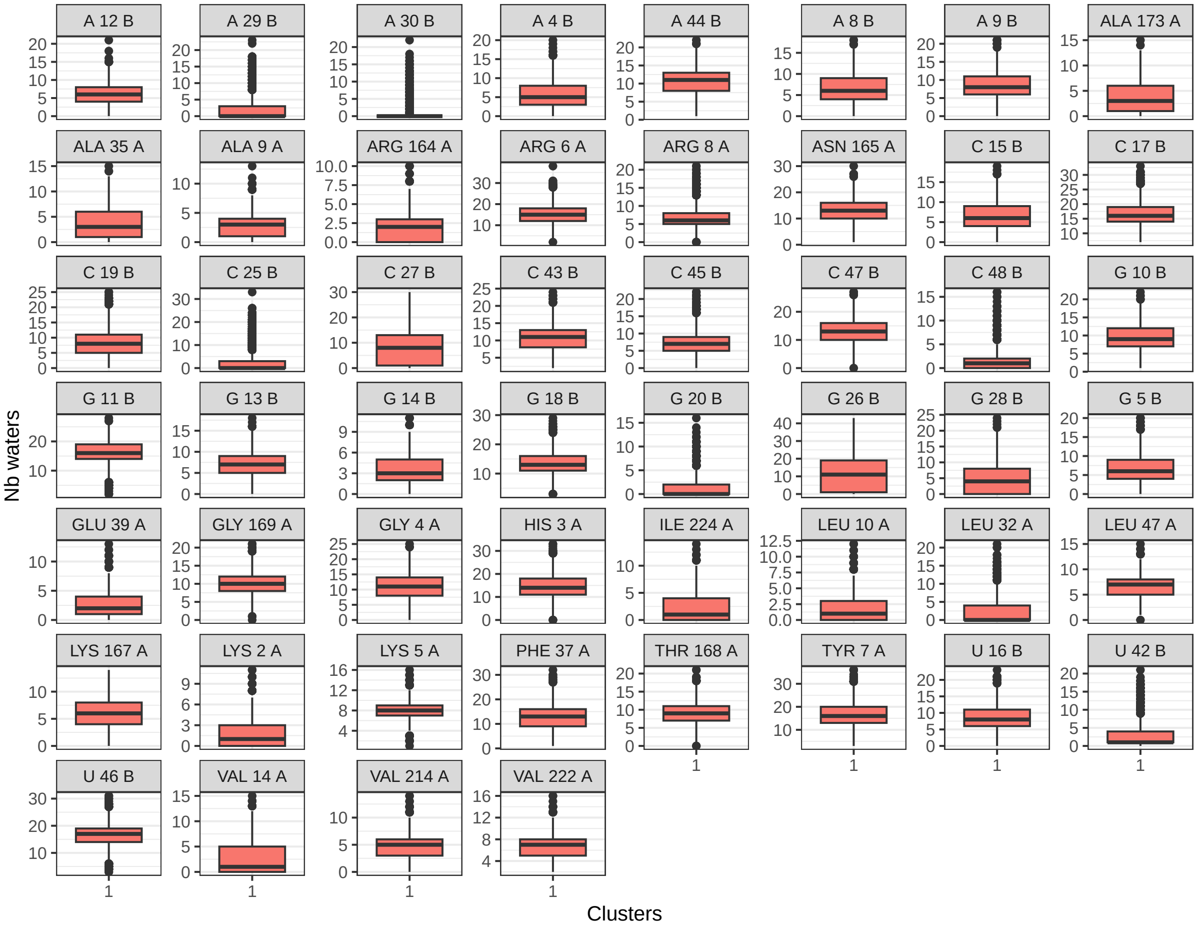


Figure S65: Number of interfacial water molecules in contact with interface residues, in each MD clusters for 2VPL complex.

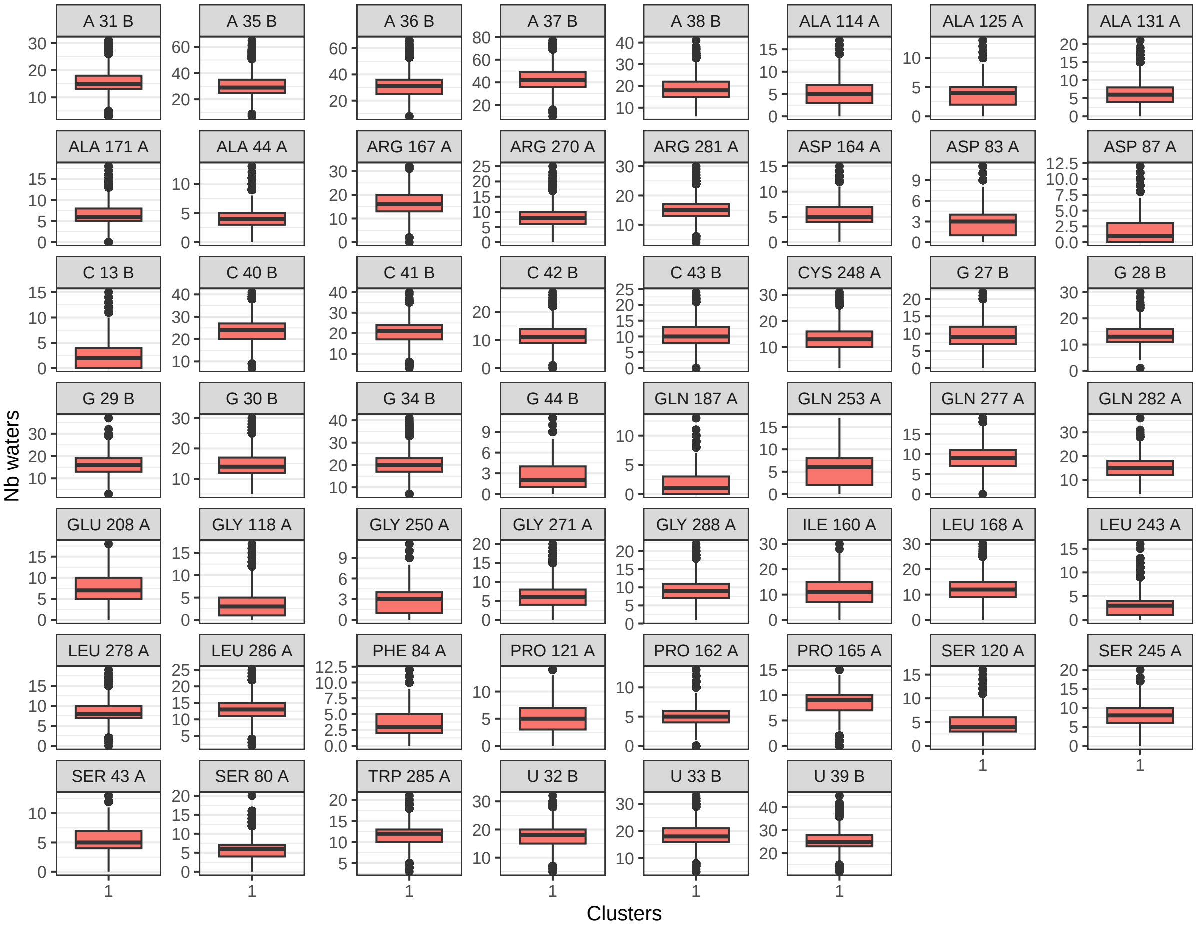


Figure S66: Number of interfacial water molecules in contact with interface residues, in each MD clusters for 2ZM5 complex.

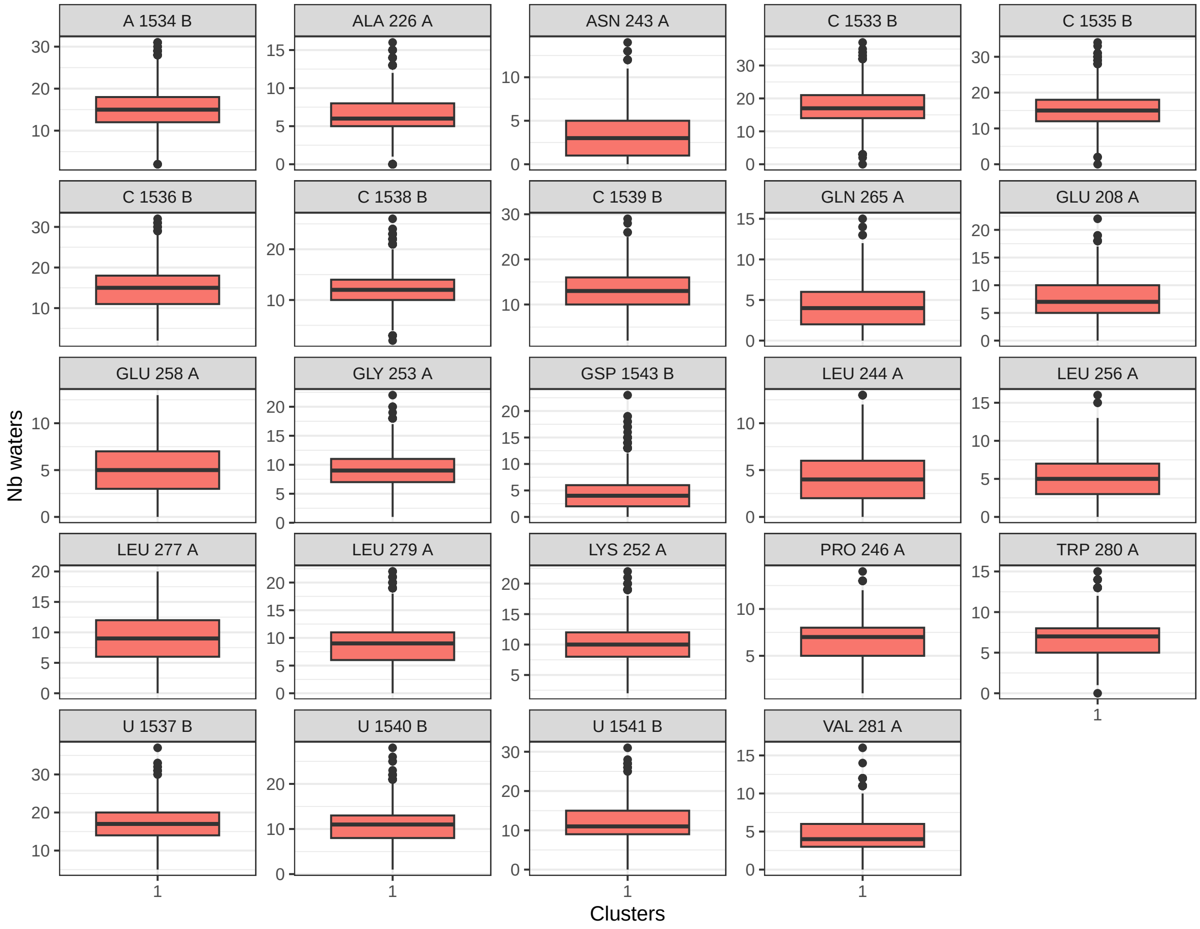


Figure S67: Number of interfacial water molecules in contact with interface residues, in each MD clusters for 3IEV complex.

### S17 Contact type

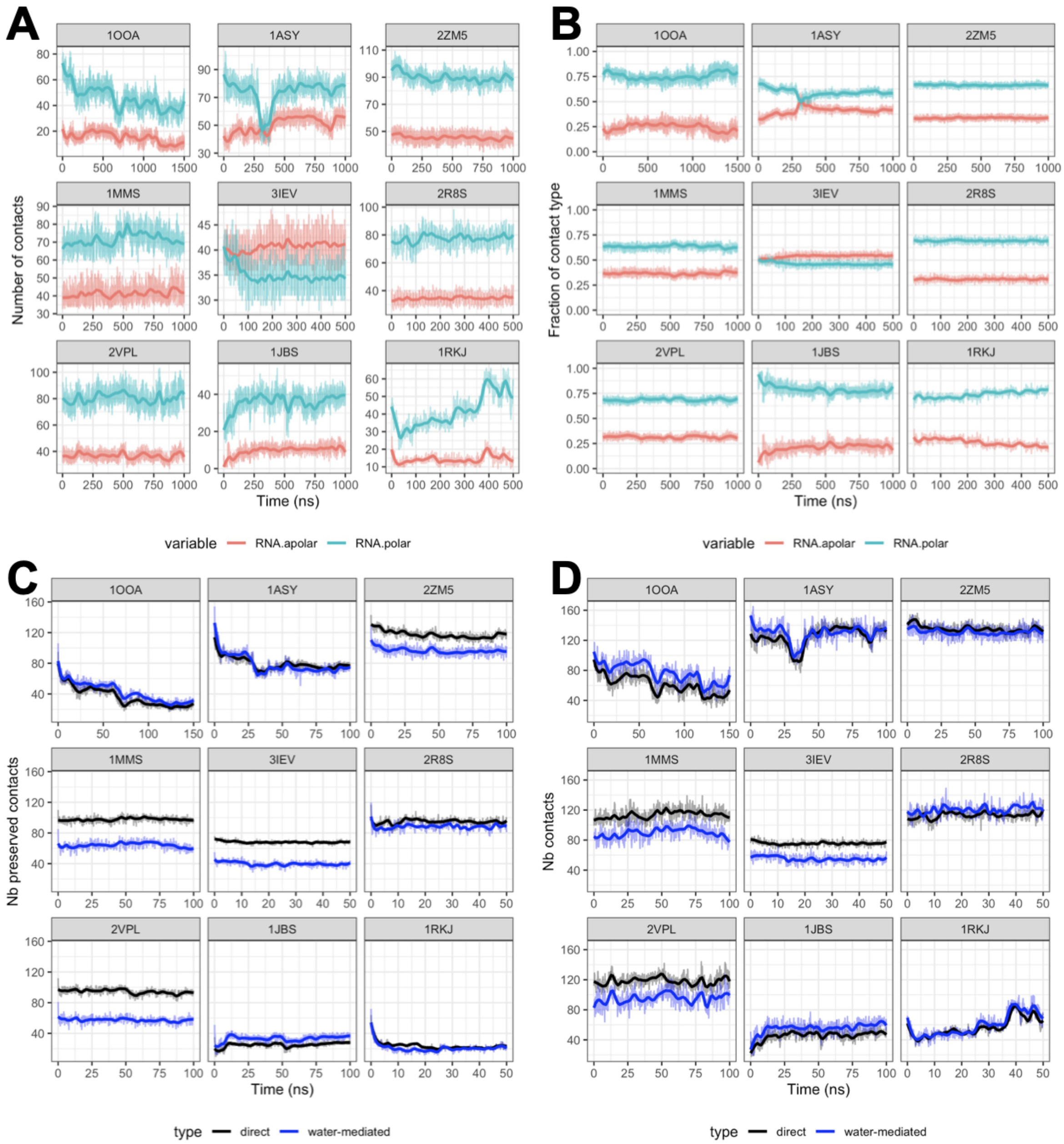


Figure S68: A: Time series of the total number of contact for each contact type at the interface. B: Time series of the fraction of each contact type at the interface. C: time series of the number of initial contacts that are preserved, direct (black) or mediated by water molecules (blue). D: Time series of the number of interface contacts, direct (black) or mediated by water molecules (blue). The amino-acids are classified as follows: polar =ASP, GLU, ASN, GLN, SER, THR, TYR, LYS, ARG, HIS; apolar=ALA, VAL, LEU, ILE, PRO, PHE, MET, TRP, GLY, CYS.

### S18 Contact water mediated

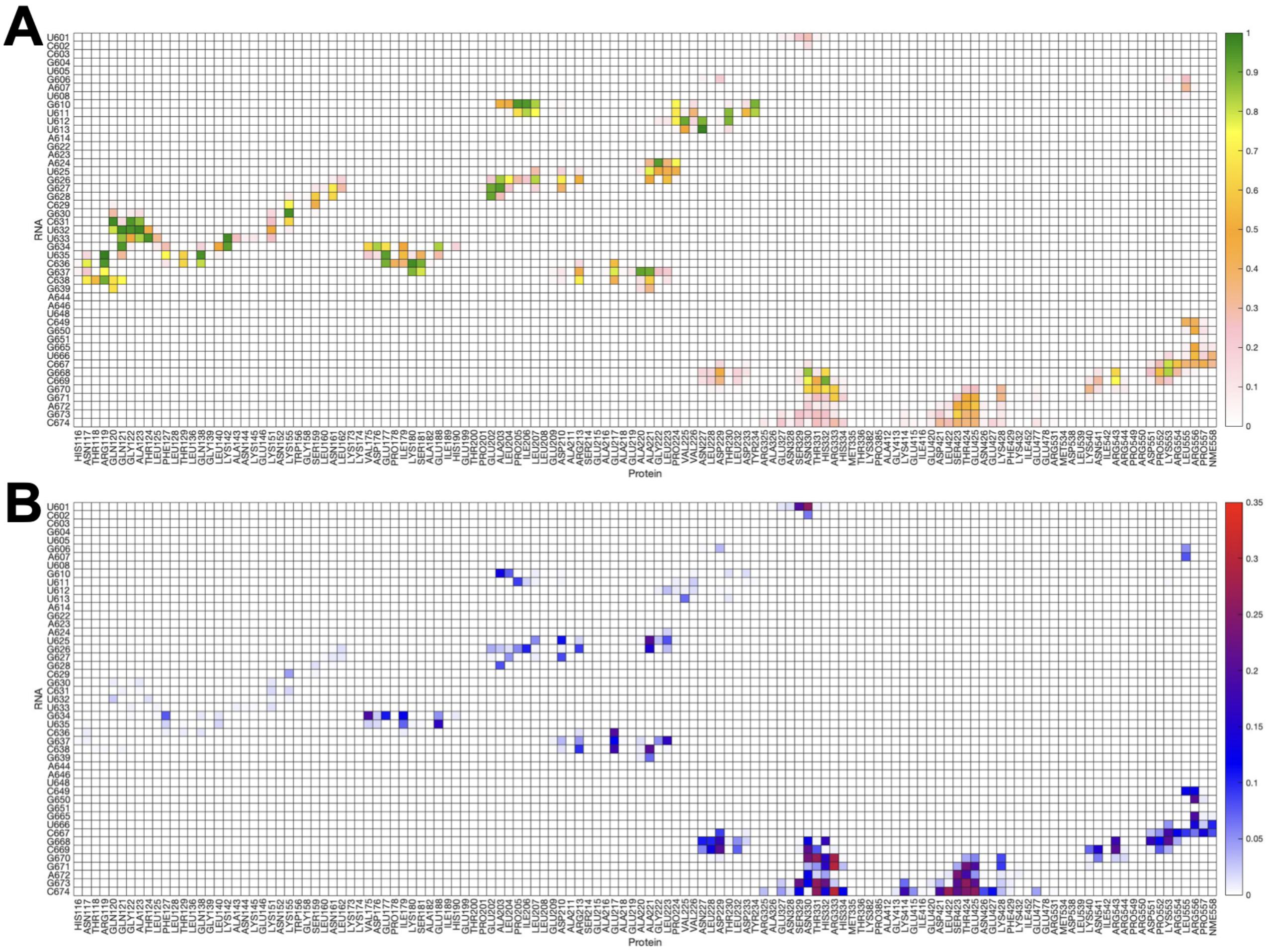


Figure S69: Relative frequency of water-mediated contacts (A) and matrix representation of each water-mediated contact variance across the three interface clusters (B) of 1ASY. Blue: low variance; Red: high variance.

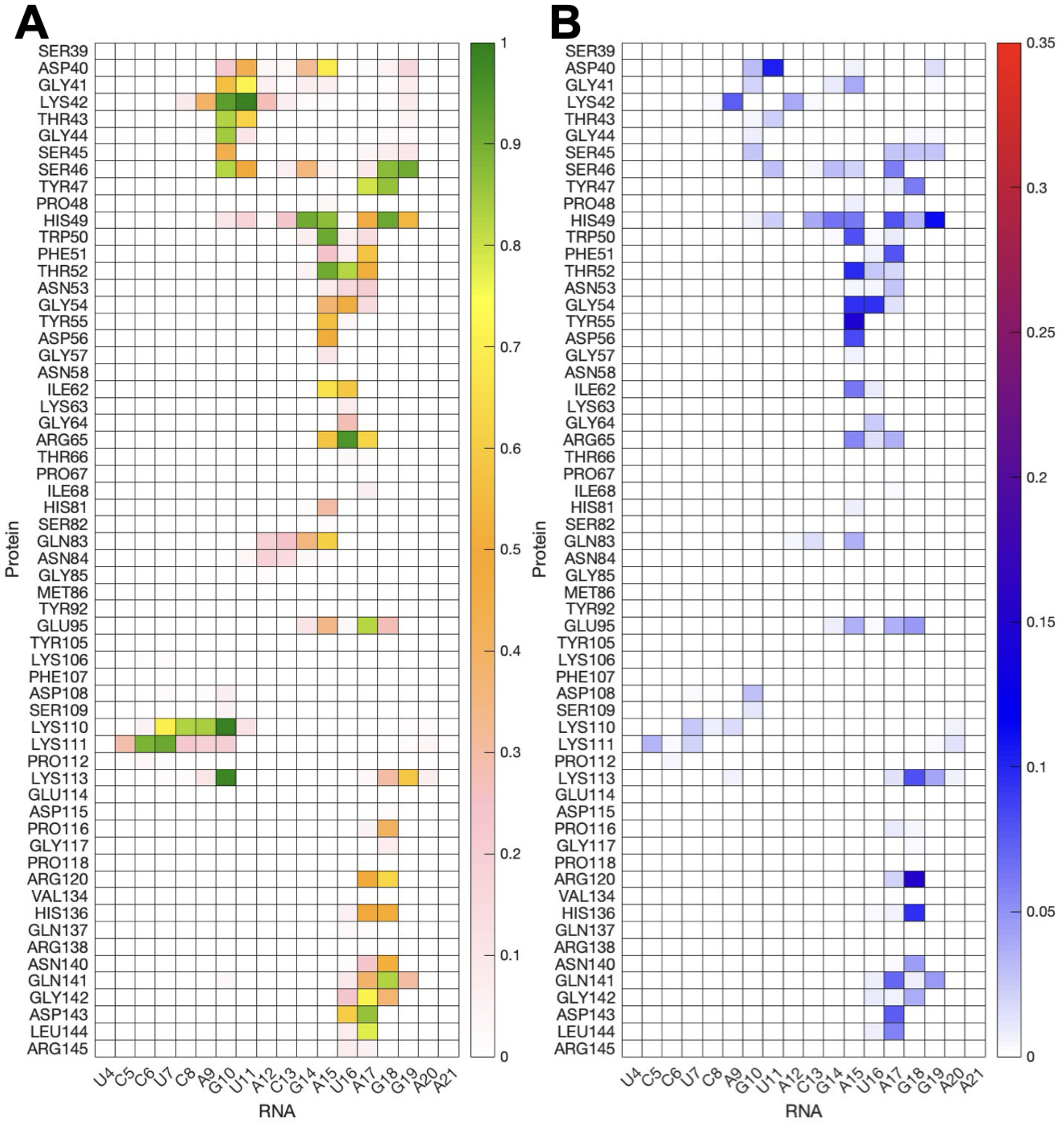


Figure S70: Relative frequency of water-mediated contacts (A) and matrix representation of each water-mediated contact variance across the two interface clusters (B) of 1JBS Blue: low variance; Red: high variance.

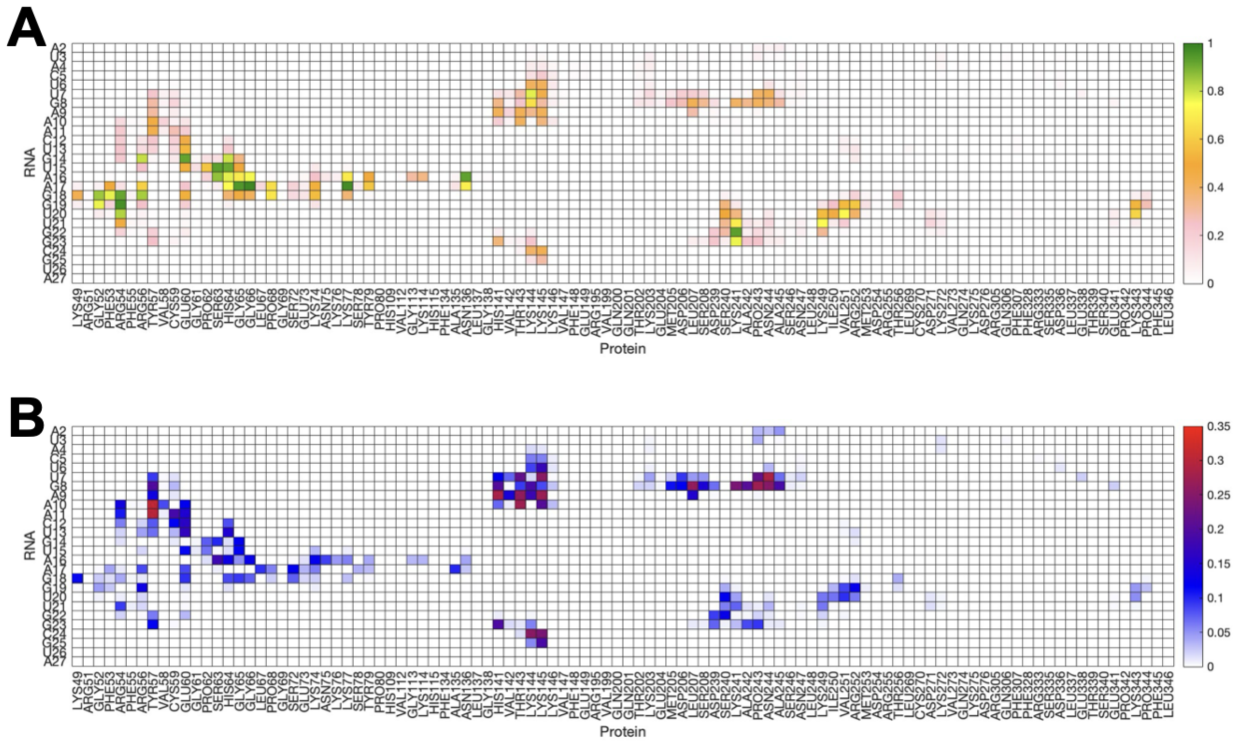


Figure S71: Relative frequency of water-mediated contacts (A) and matrix representation of each water-mediated contact variance across the four interface clusters (B) of 1OOA. Blue: low variance; Red: high variance.

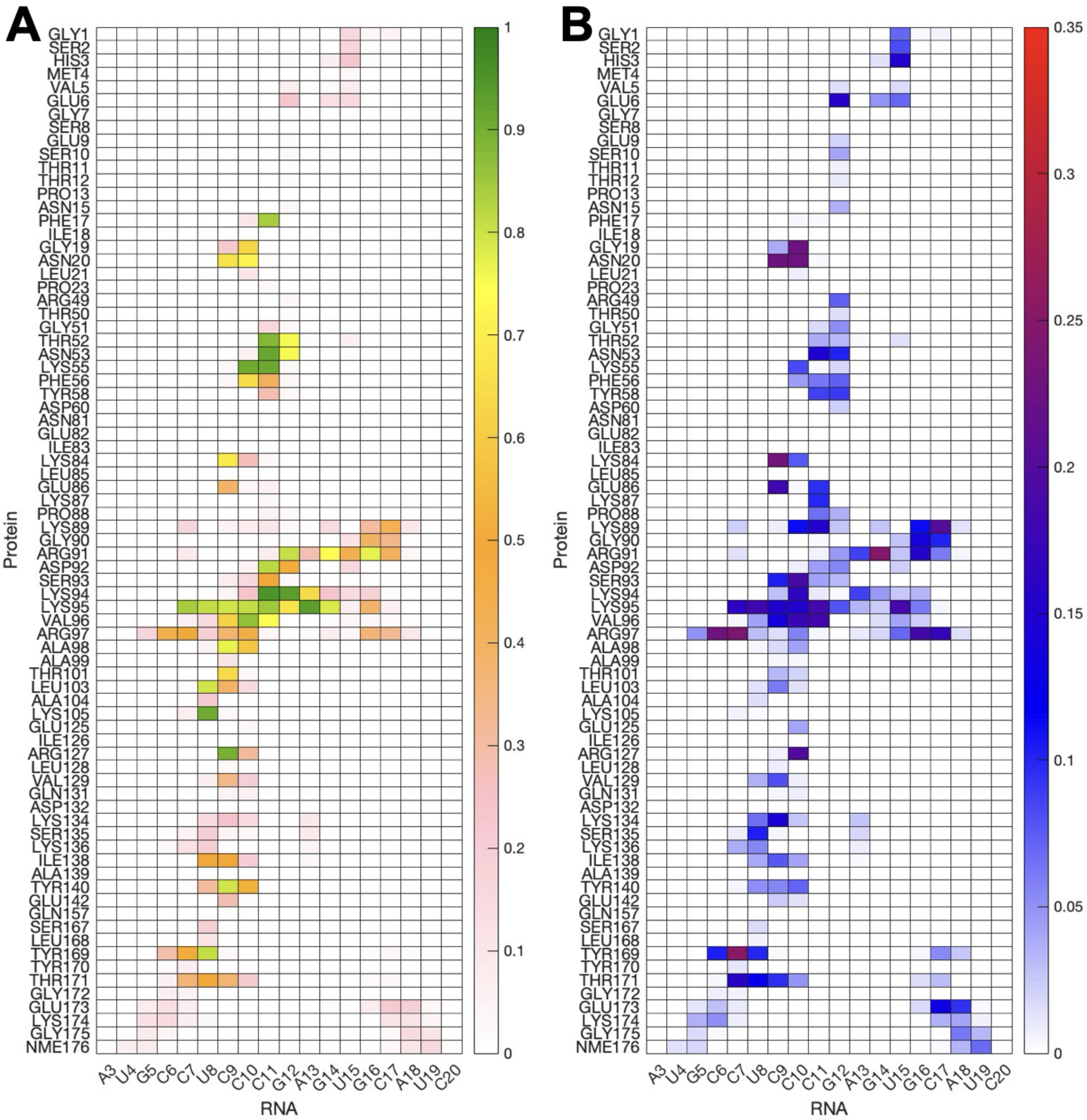


Figure S72: Relative frequency of water-mediated contacts (A) and matrix representation of each water-mediated contact variance across the five interface clusters (B) of 1RKJ. Blue: low variance; Red: high variance.

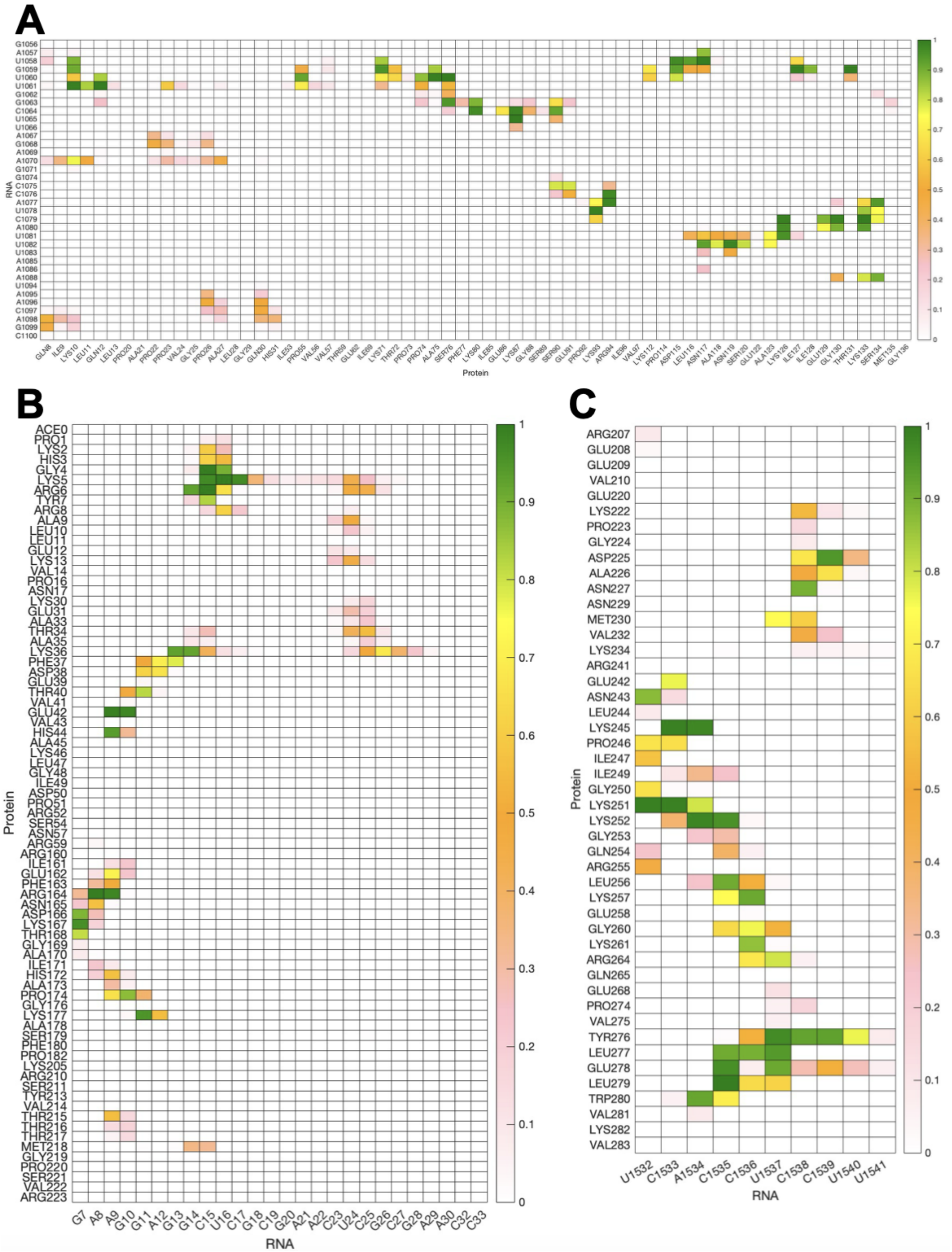


Figure S73: Relative frequency of water-mediated contacts of 1MMS (A), 2VPL (B) and 3IEV (C).

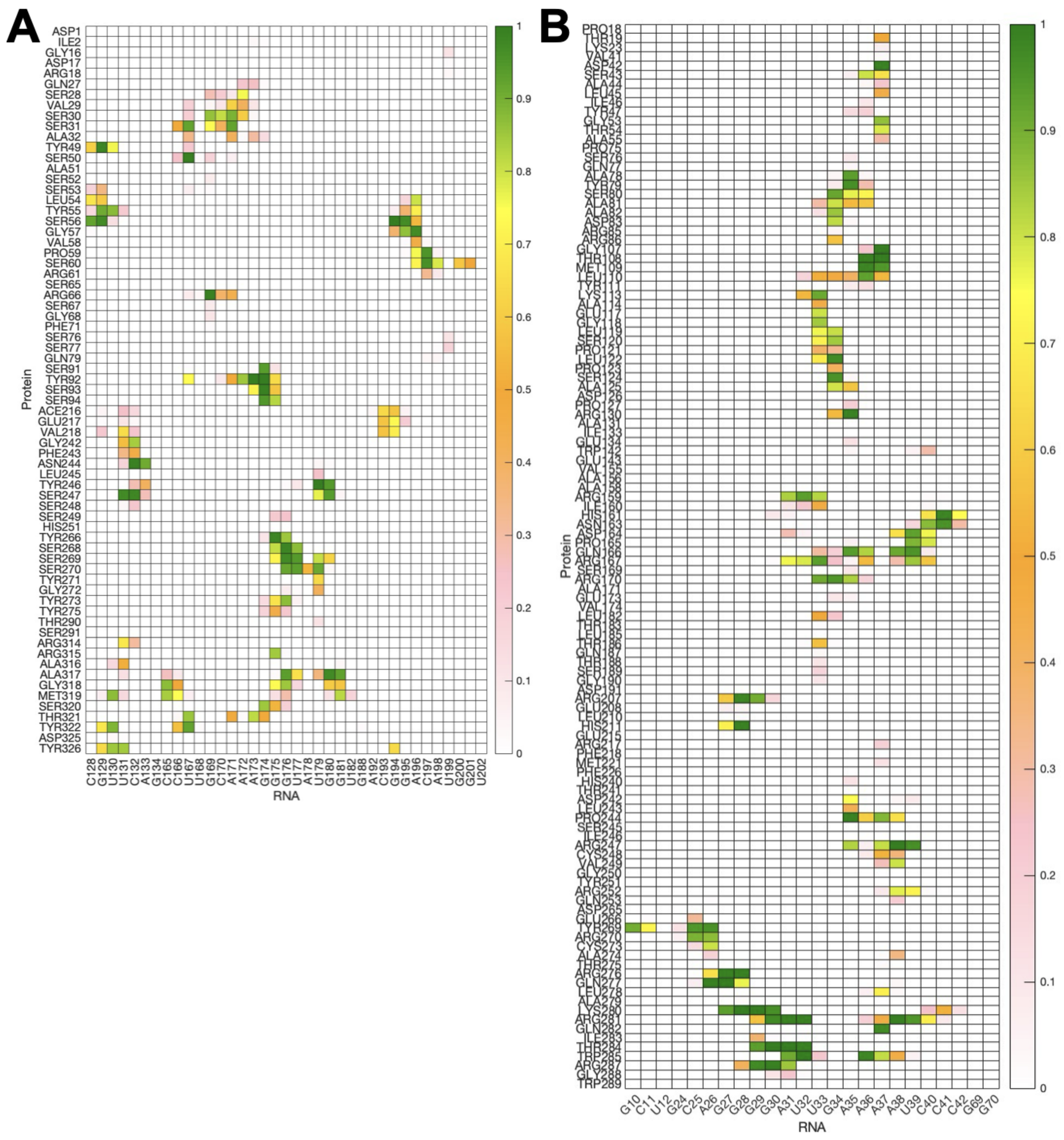

Figure S74: Relative frequency of water-mediated contacts of 2R8S (A) and 2ZM5 (B).
